# Lasy-Seq: a high-throughput library preparation method for RNA-Seq and its application in the analysis of plant responses to fluctuating temperatures

**DOI:** 10.1101/463596

**Authors:** Mari Kamitani, Makoto Kashima, Ayumi Tezuka, Atsushi J. Nagano

## Abstract

RNA-Seq is a whole-transcriptome analysis method used to research biological mechanisms and functions; its use in large-scale experiments is limited by costs and labour. In this study, we established a high-throughput and cost effective RNA-Seq library preparation method that did not require mRNA enrichment. The method adds unique index sequences to samples during reverse transcription (RT) that is conducted at a higher temperature (≥62°C) to suppress RT of A-rich sequences in rRNA, and then pools all samples into a single tube. Both single-read and paired end sequencing of libraries is enabled. We found that the pooled RT products contained large amounts of RNA, mainly rRNA, and caused over-estimations of the quantity of DNA, resulting in unstable tagmentation results. Degradation of RNA before tagmentation was necessary for the stable preparation of libraries. We named this protocol low-cost and easy RNA-Seq (Lasy-Seq), and used it to investigate temperature responses in *Arabidopsis thaliana*. We analysed how sub-ambient temperatures (10–30°C) affected the plant transcriptomes, using time-courses of RNA-Seq from plants grown in randomly fluctuating temperature conditions. Our results suggest that there are diverse mechanisms behind plant temperature responses at different time scales.

## Introduction

RNA-Seq enables us to analyse transcriptomes, the comprehensive expression profile of the genome, and has been used for a variety of analyses, such as the effects of mutations ^1,2^, stress responses ^3-5^, chemical biosynthesis pathways ^6^ and plant-pathogen interactions ^7,8^. However, large scale experiments have been limited due to the large costs required for library preparation and sequencing. Recently, with the rise of single cell RNA-Seq technology, an increasing number of methods for high-throughput RNA-Seq have been reported ^9^. In conventional RNA-Seq methods, enrichment of mRNA occurs at the first step of library preparation, with oligo-dT beads or enzymatic digestion of rRNA in samples ^10^. The preparation of a large number of libraries is cost- and labour-intensive and can result in high variance of the quality and quantity of samples. Previous studies of single cell RNA-Seq have developed a method that adds unique index sequences to each sample during the reverse transcription (RT) step, the first step of library preparation, by adding unique cell-barcodes located in Oligo-dT RT primers ^11,12^. The index-added samples can be pooled into a tube and all remaining reactions conducted in single tube. In applying sample pooling at an early step in a library preparation, concern about false-assignment among samples has been reported ^13^. The rate of false-assignment caused by sequencing (index-hopping) was reported to reach 2% in sequencing with sequencers with patterned flow-cell such as NextSeq, HiSeq 4000 and HiSeq X ^14^. Although the rate was small, in sequencers with non-patterned flow cells such as MiSeq and Hiseq 2500, false-assignment could also be caused by excessive PCR amplification of the library during its preparations, at rates reported to reach 0.4% ^13^. Reducing the steps in library preparation is expected to reduce sample loss caused by insufficient reaction or purification steps. To reduce the steps and amount of time taken for library preparation, previous studies have employed tagmentation with a Tn5 transposase ^15-17^. Efficiency of tagmentation by transposase was reported to be largely affected by the amount of input DNA, resulting in changes in the distributions of insert length ^9^.

In plants, RNA-Seq has been used to analyse various environmental-responses. Plants detect environmental changes, such as ambient temperature fluctuations, with high sensitivity and subsequently alter their growth and/ or architecture ^18,19^. For example a 10% reduction in rice yield and strong inhibition of lettuce seed germination were caused by an increase of only 1°C in ambient temperature ^20,21^. In *Arabidopsis*, high ambient-temperatures cause spindly growth and early flowering of plants, while low ambient-temperatures repress flowering ^22-24^. Molecular mechanisms of ambient-temperature responses are starting to be identified ^25,26^. Furthermore, several studies have indicated that plants refer to past temperatures, such as the existence of heat shock memory ^27,28^. Moreover, it has also been reported that sub-lethal heat stress of plants can result in acquired tolerance to subsequent higher heat stress events, known as heat acclimation. Heat stress memories are stored for longer intervals, this is different from acute tolerance known as a heat shock response ^29-31^. Because majority of these previous studies were conducted under a few constant-temperature conditions, less is known about how long or how much plants refer past temperature.

In this study, we have developed a high-throughput and cost-effective RNA-Seq library preparation method with RT indexing of total-RNA samples, which let us skip the process of mRNA enrichment and pools all samples into a single tube at an early stage of library preparation. Using this method, we have revealed the ambient temperatures and durations of exposure to them, by randomly changing the growth temperatures from 10°C to 30°C every other day, that affected the transcriptomes of *A*. *thaliana*.

## Results

### Optimization of RNA-Seq library preparation methods for high-throughput processing

To develop a high-throughput and cost-effective RNA-Seq preparation method, we applied methods used for single cell RNA-Seq (scRNA-Seq) in previous studies. In the scRNA-Seq method, the amount of input RNA was small, therefore all samples were pooled after being indexed by an index-added primer during RT step. Furthermore, previous studies employed tagmentation with transposase (Nextera TDE1 enzyme) after the second strand synthesis ^32^. As transposase fragmentizes dsDNA by inserting adapters, the tagmentation step can replace fragmentation, end-repair, dA-tailing and adapter ligation steps from the conventional RNA-Seq methods applied in TruSeq ^17^. The pooling and tagmentation steps result in reduced financial costs and labour, allowing us to develop a high-throughput and cost-effective method for RNA-Seq. Initially we simply applied the method from the previous study, hereafter referred to as the small-input method (SI-method), into bulk RNA-Seq, using larger amounts of input RNA than scRNA-Seq (Fig. 1) ^16^. However, due to several problems discussed in the proceeding, we decided to optimize the SI-method for bulk RNA-Seq thus developing a new method; method for large-input (LI-method) (Fig. 1), named low-cost and easy RNA-Seq (Lasy-Seq). Examination of Lasy-Seq was conducted using RNA from *Oryza sativa*.

**Figure 1.**
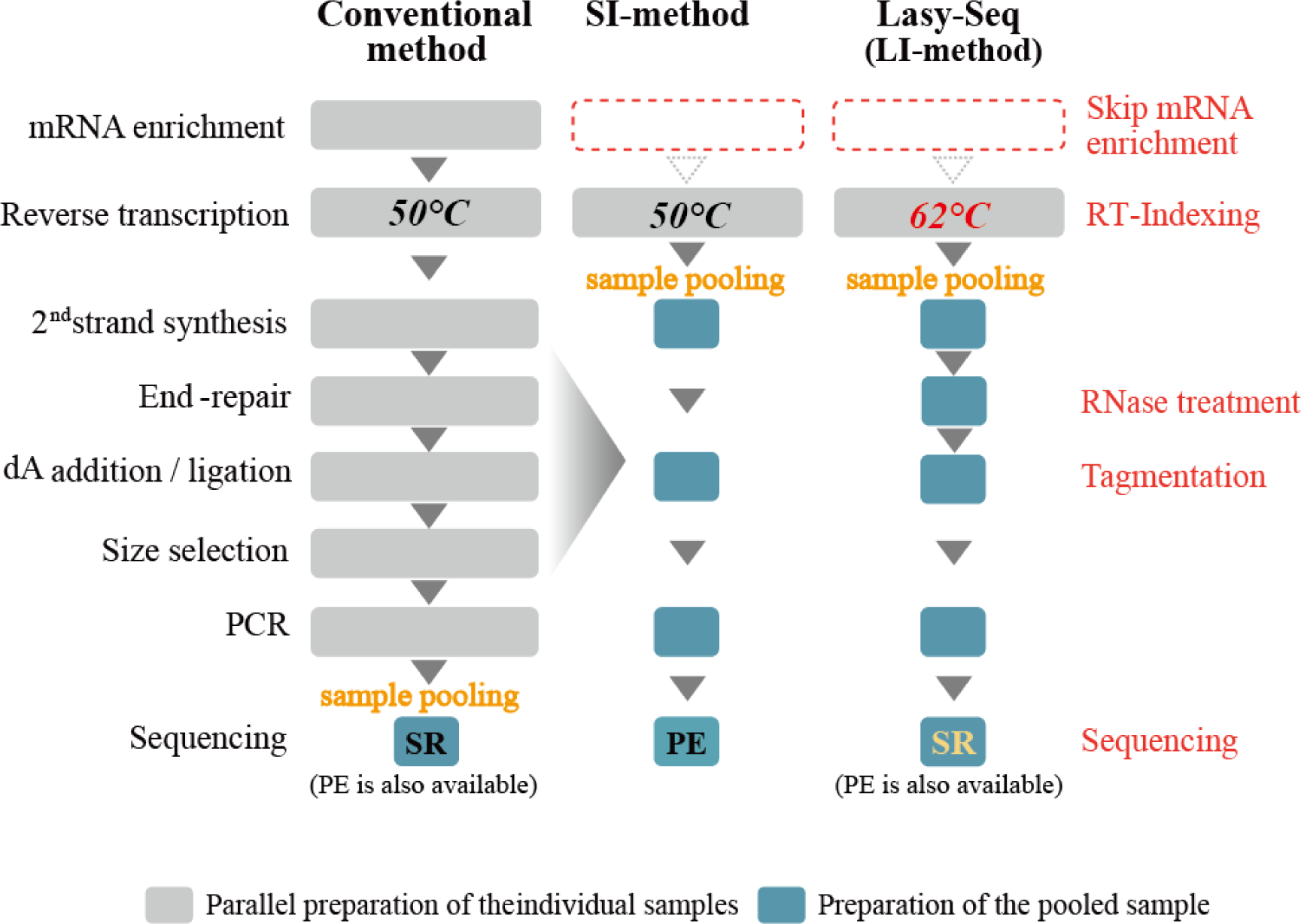
Comparison of the RNA-Seq library preparation methods. Steps modified in Lasy-Seq are shown on the right with red characters. In the conventional method (left), the high-throughput RNA-Seq required parallel preparation of all individual samples throughout all experimental steps. In Lasy-Seq, enrichment of mRNA was not required, and all samples were pooled into a single tube after the RT step, by adding unique index sequencing to each sample at the RT step. SI and LI indicate small-input and large-input total RNA, respectively. SR and PE indicate single-read sequencing and paired-end sequencing, respectively.

We found three main difficulties in applying the SI-method to bulk RNA-Seq. First, we detected large amounts of non-poly-A reads such as rRNA in our bulk RNA-Seq data. In the SI method, we could skip the process of mRNA-enrichment and RT was conducted directly from the total RNA. We found that not only mRNA but also rRNA was transcribed from their internal A-rich regions in rRNAs (Fig. 2A). This phenomenon was also observed in previous studies ^33^. To avoid consumption of sequence reads by rRNA, we tried to supress RT for rRNA by increasing the RT reaction temperature. We set the RT temperature at 50°C (the original temperature with Superscript IV reverse transcriptase), 56°C and 62°C. The number of reads of rRNA was drastically decreased in RT at 62°C (Fig. 2A). In addition, the amount of cDNA of non-poly-A genes other than rRNAs and poly-A genes were quantified by qPCR. The amount of cDNA from poly-A RNA was similar all temperatures, while the amount of cDNA from non-poly-A RNA was reduced (Cp value was increased) at 62°C (Fig. 2B); therefore, we concluded that RT at temperatures greater than 62°C could suppress the transcription of rRNA.

**Figure 2.**
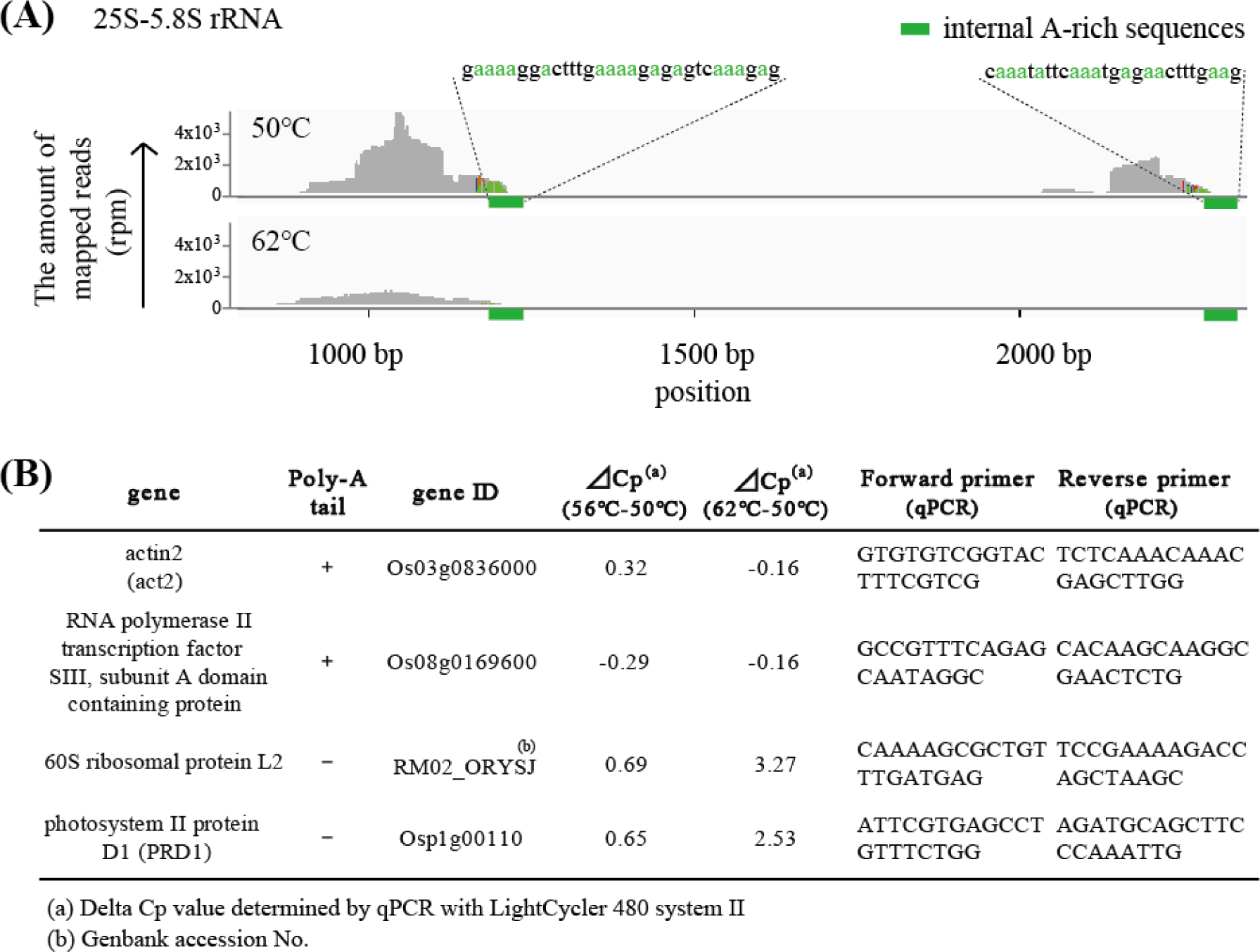
High temperatures suppress RT of non-poly-A RNA. (A) Comparison of the distribution of the reads mapped on 25S-5.8S rRNA reverse transcribed at 50 °C and 62 °C. RT of non-poly-A tailed RNAs were observed from internal A-rich regions. RT at higher temperature suppressed the RT from internal A-rich regions of non-poly-A tailed RNA. **(B)** List of the delta-Cp values in RT-qPCR on genes with and without poly-A tails.

Second, we found that the results of tagmentation were unstable, although the same amounts of input DNA were used. The cause was determined to be RNA-carryover that was also quantified as DNA (Fig. 3), causing over estimations of the quantity of DNA. This could affect the length-distribution of the tagmentation product, as the frequency of tagmentation by transposase was determined from the stoichiometry of DNA and transposase ^17^. This difficulty was solved by adding an RNase treatment step before quantification of DNA for the tagmentation step. We found that the RNase A (or RNase T1) reaction at 37°C for 5 min was enough to remove the RNA in our protocol. In conventional bulk RNA-Seq the problem of RNA-carryover does not occur, as the enrichment step of mRNA was included in the protocols before RT ^10^. It may not be a problem in scRNA-Seq because the procedure uses minute quantities of input RNA and pre-amplification. The degradation step for RNA was necessary with the bulk RNA-Seq without mRNA enrichment.

**Figure 3.**
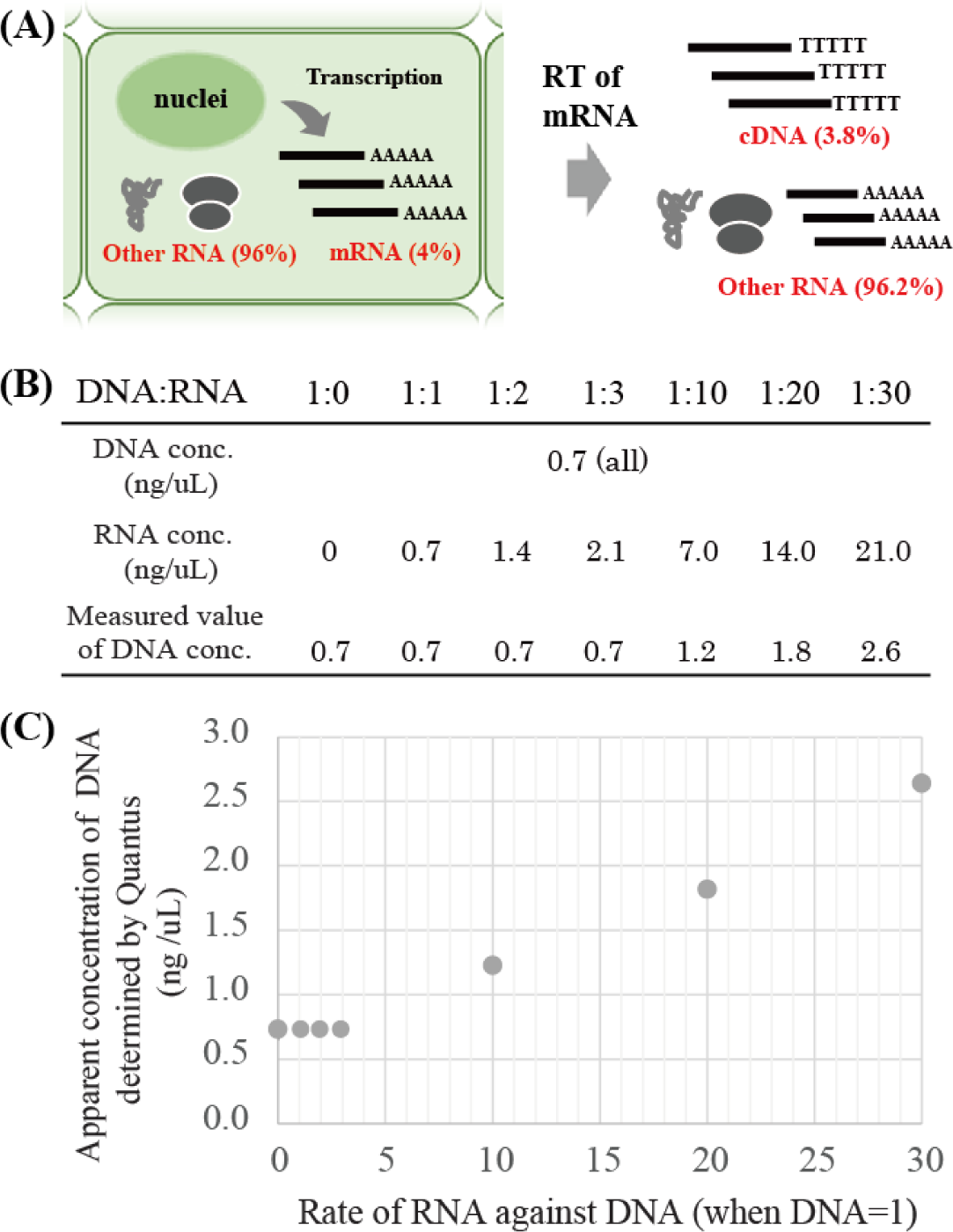
Effect of RNA additions on DNA quantification. **A)** The amount of RNA in reaction solutions after RT. If mRNA occupied 4% of the total amount of RNA in a cell, the rate RNA remained after RT of total RNA became 25 times larger than cDNA. **(B)** Effect of RNA on the measurement of DNA concentrations. Concentrations of DNA were determined for samples with constant amounts of DNA (0.7 ng) and different amounts of RNA (from 0 to 30 times larger than the DNA quantity). DNA concentration was over-estimated in RNA-added samples which contained RNA concentrations more than 10 times larger those of DNA. In the table, “DNA conc.” and “RNA conc.” indicate true concentration of measured liquids. “Measured value of DNA conc.” means the concentration determined by QuantiFluor dsDNA System and Quantus Fluorometer. **(C)** The plotted concentrations and DNA:RNA rates.

Finally, the SI-method required paired-end sequencing, the cost of which is greater than that for single-read sequencing. At first we prepared RT primers for paired-end sequencing based on a previous study (PE78 RT-primer and PE 60 RT primer in Supplementary Fig. S1)^16^. After confirming that these primers worked well using *O*. *sativa* RNA, primers were designed for single-read sequencing of Lasy-Seq (Supplementary Fig. S1). The library constructed by the Lasy-Seq method can be sequenced by not only single-read sequencing, but also by paired-read sequencing from which information of unique molecular identifiers (UMI) is available.

### Rate of false-assignment among the pooled samples

In order to estimate the false-assignment among samples during the PCR and sequencing steps, we prepared samples with and without ERCC-controls and quantified the number of ERCC-control reads detected in samples without ERCC-control. Early-pooled sets were pooled before the library amplification step and late-pooled sets were pooled before the sequencing step. RT primers of different lengths (60 mer and 78 mer) were used and a total of eight samples were prepared (Fig. 4). The technical replicates showed high correlation with each other (Pearson’s correlation coefficients of 0.986 and 0.998 for each of the two RT primer set in Fig. 4A, respectively).

**Figure 4.**
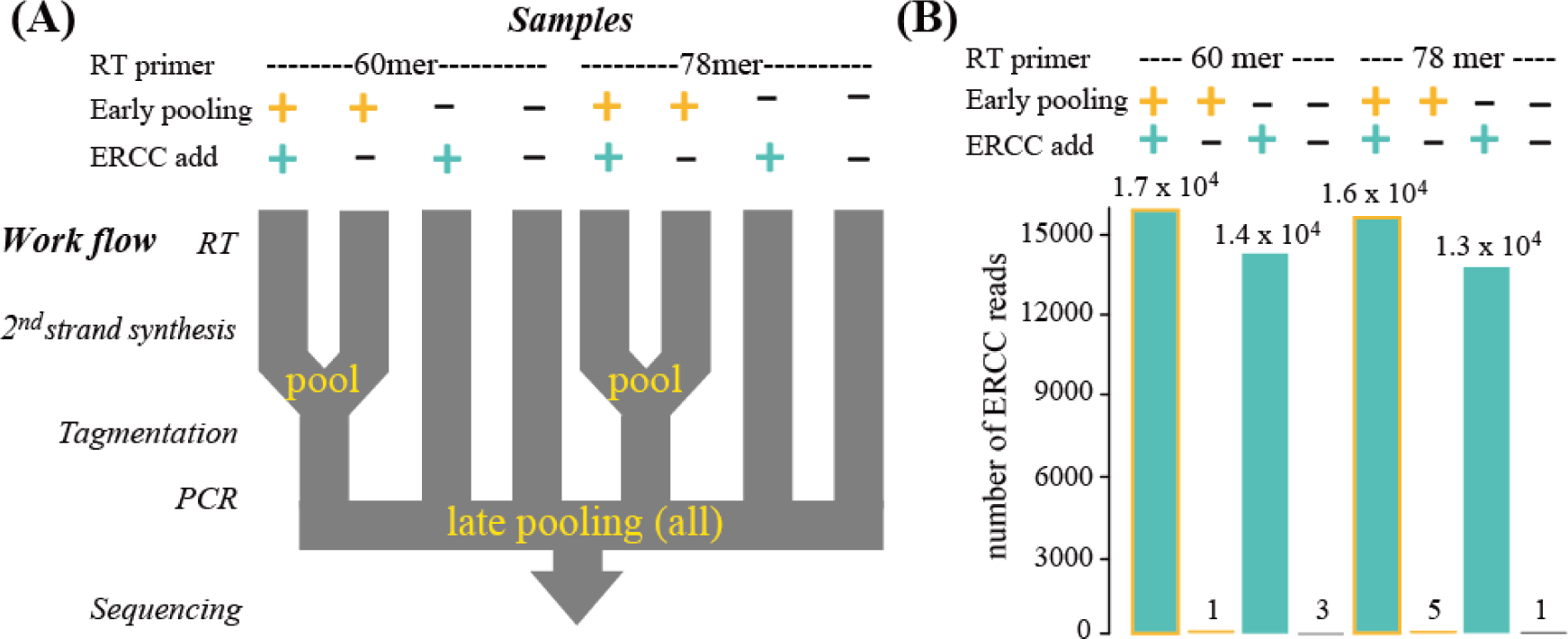
Evaluation of the false-assignment rates associated with sample pooling at early stages. **(A)** Flow of the library preparation for evaluation of false-assignment rates among samples. Early-pooled sets were pooled before the tagmentation step, while late-pooled sets were individually prepared until purification after PCR. All samples were pooled prior to sequencing. **(B)** Number of ERCC reads detected in each sample. Numbers shown above the bar-plot indicate the read number of ERCC. Conditions of the experiment for each sample were shown by the colours of the bar and indicated over the bar-plot.

In late-pooled samples, among randomly selected 10^5^ reads, 1.4 x 10^4^ and 1.3 x 10^4^ reads were mapped on samples with ERCC-control for each RT primer, while 3 and 1 reads were detected in samples without ERCC-control (Fig. 4). These reads could be derived from other samples with ERCC-control sequenced together (in total 6.0 x 10^4^ ERCC-control reads), therefore the false-assignment rate of this lane during sequencing was 0.027% (Supplementary Fig. S2). In early-pooled samples, 1.7 x 10^4^ and 1.6 x 10^4^ reads were mapped on samples with ERCC-control. The number of reads obtained from samples without the ERCC-controls were 3 and 5, which occupied 0.031% and 0.018% of the paired-pooled samples for each RT primer (Fig. 4). These rates include the false-assignment rates caused by sequencing. Therefore, according to rough estimates, the difference between early-pooled and late-pooled samples could be regarded as the false-assignment rate during PCR. The rates of the subtractions (1 and 3 reads) against the ERCC reads in the paired samples were 0.0060% and 0.019% of the paired-pooled samples, respectively (Supplementary Fig. S2). By considering these data, we regarded that false-assignments among samples were almost the same as the rates reported by previous studies (Supplementary table S3). We have concluded that the rates were at an acceptable level for both the RT primer sets when using optimal PCR cycles in the amplification of libraries.

### Correlation between plant transcriptomes and past temperatures

We applied this method to investigate the effect of sub-ambient temperature changes on gene expression of *A*. *thaliana*. Analyses on the correlation between the plant transcriptome and temperatures on the sampling day or previous days were conducted. Plants were cultivated under temperatures randomly fluctuating between 10°C and 30°C each day (Fig. 5). Samples were collected every day at noon for 8 days and were analysed with Lasy-Seq. For each of the 45 samples, from 5.8 x 10^5^ to 6.2 x 10^6^ reads were obtained by sequencing. The rate of reads mapped to the reference sequences were from 93.7% to 95.8% of the total reads. Correlations were calculated between the transcriptomes and the growth temperature on the sampling day and 1,2 and 3 days prior to sampling (Fig. 6). We confirmed that there were no correlations between temperatures on these days (Fig. 5C). The number of genes significantly correlated with each temperature were 2921, 435, 351 and 8 genes for the sampling day and 1, 2 and 3 days prior to sampling, respectively (adjusted p <0.1, correlation coefficients >0.05, red points in Fig. 6, Supplementary table S2). The effect of temperature on gene expression was largest on the sampling day, and then decreased with the lapse of time (Fig. 6).

**Figure 5.**
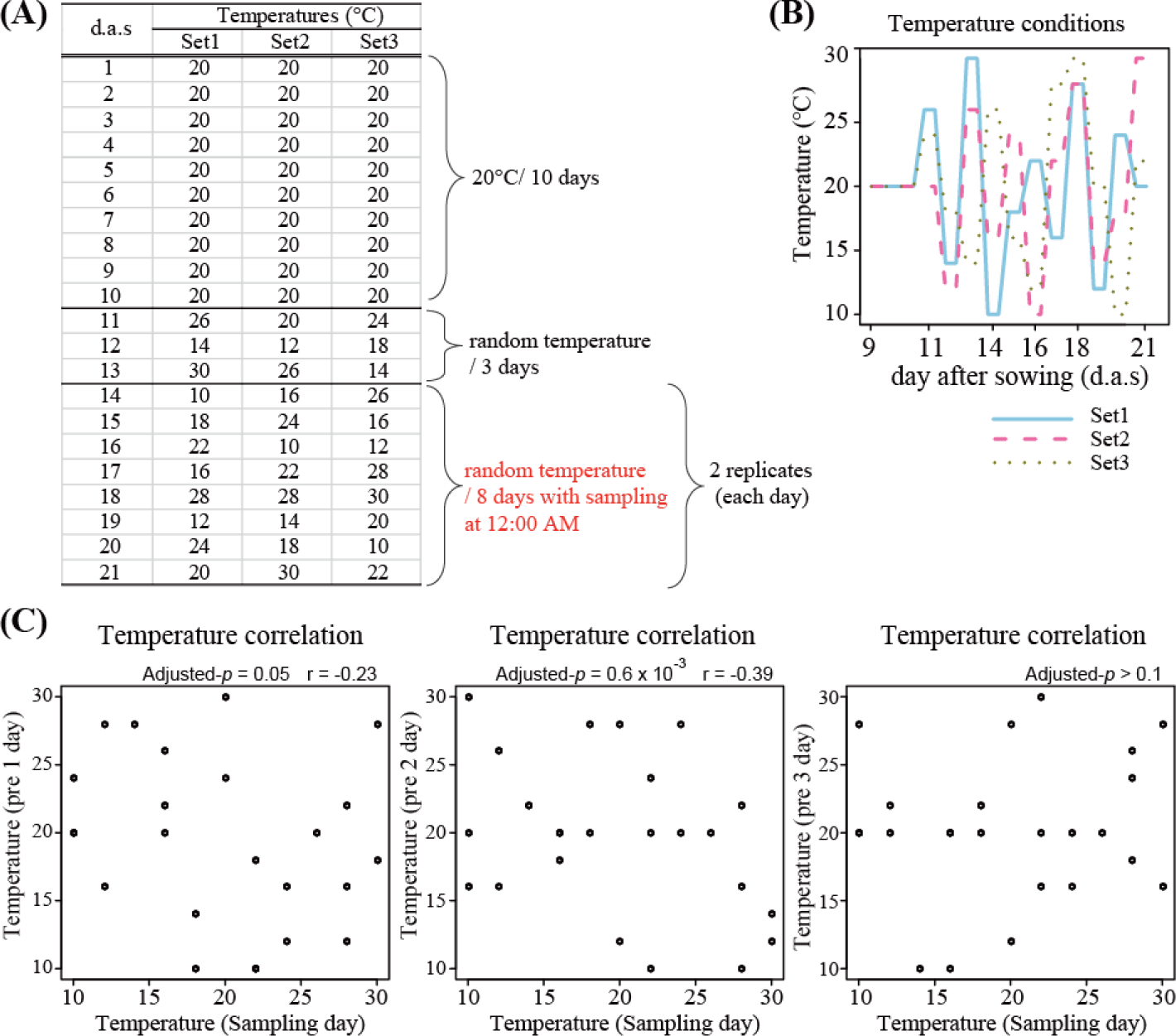
Temperature settings in the temperature response experiment. **(A)** The three sets of temperature conditions. Plants were grown at 20°C for 8 days and then at changing temperature conditions for 3 days. Sampling was conducted from 14 to 21 day after sowing (d.a.s.), indicated by red characters. **(B)** Diagram of the temperatures of the three sets from 8 d.a.s to 21 d.a.s. **(C)** Correlation of the temperature between sampling day and the days prior to sampling. Horizontal axis shows temperature (°C) on the sampling day and vertical axis indicates the temperatures 1,2 and 3 days prior to sampling (from left to right, respectively). The “Adjusted-*p*” indicated adjusted p-value (FDR) and “r” indicated Pearson’s correlation coefficients.

**Figure 6.**
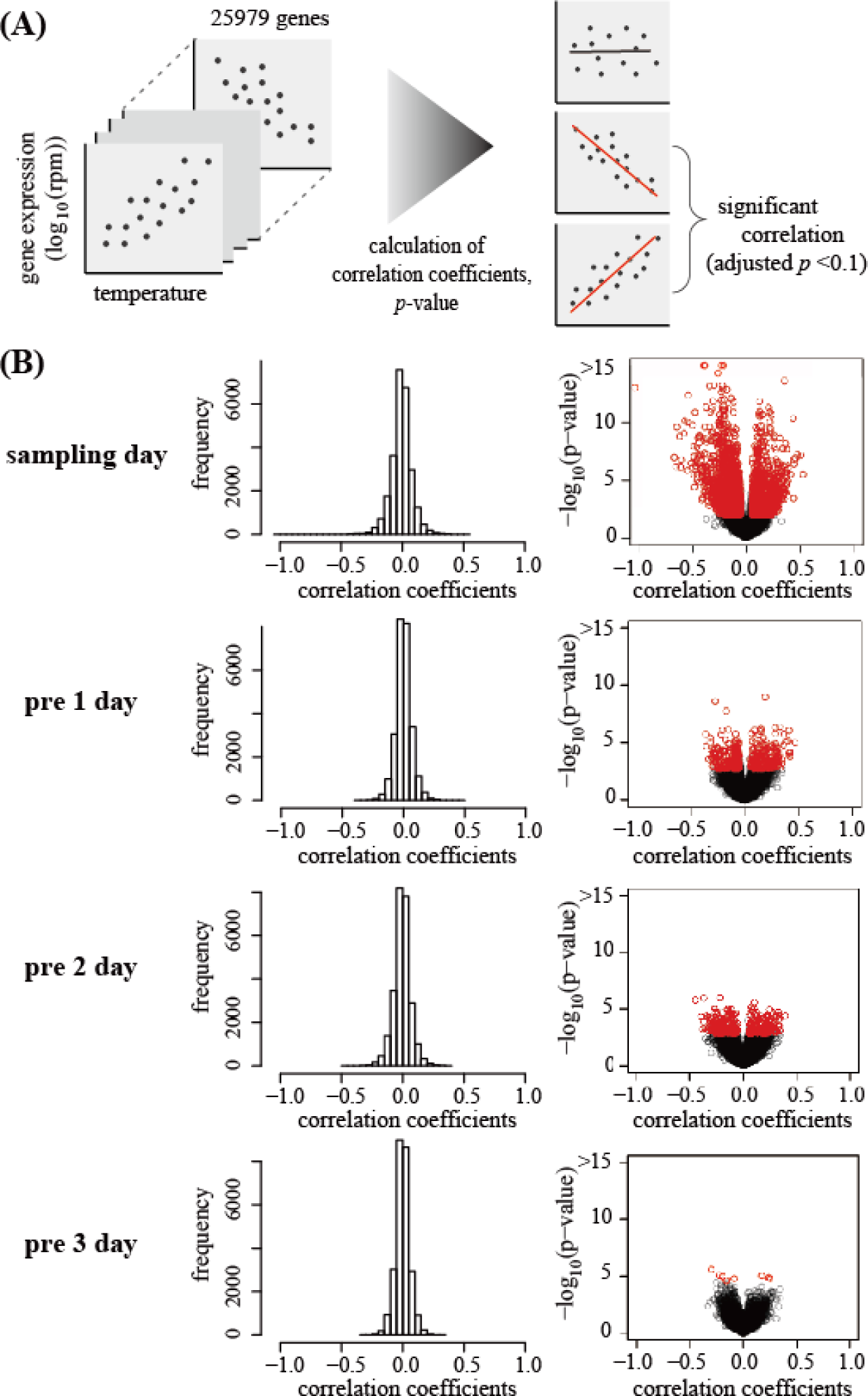
The correlation between the transcriptome and the temperature. **(A)** Flow of the analysis of the correlation between gene expressions and temperatures. **(B)** Distribution of the correlation coefficients for each gene between gene expression levels and the temperature of the sampling day and 1,2 and 3 day prior to sampling (from top panel to bottom panel, respectively). Each circle indicates each gene. Red and orange circles indicate genes for which significant relationships between the expression and the temperature (adjusted *p*-value<0.1) were detected. Red circles represent genes with correlation coefficients of more than 0.05.

The expression of *GIGANTEA* (*GI*) and *PHYTOCLOCK 1* (*PCL1*, synonym: *LUX ARRHYTHMO, LUX*) were negatively correlated with the temperature on sampling day (Fig. 7). These two genes have been related to circadian rhythms ^34^. The amplitudes of the circadian oscillations of *GI* and *PCL1* expression became larger with the increase of temperature, even in the ambient temperature ranges ^35^. All samples were collected at 12:00 (AM) to detect snapshots of the transcriptome, so the increase of the amplitude must be interpreted as a decrease in expression in this study (Fig. 7B). Another example, expression of *LEAFY* (*LFY*) was positively correlated with the temperature on sampling day (Fig. 7). *LFY* is a floral meristem identity gene, which triggers the transition from vegetative to reproductive phases ^36^. Similar temperature-response patterns were observed in *MYB33* and *PUCHI*, which were reported to be positive regulators of *LFY* ^37-39^. *MYB33* mediates gibberellin (GA)-dependent activation of LFY ^37^. *PUCHI*, an AP2/EREBP family gene, plays important roles in floral fate determination and bract suppression ^38^. High correlation suggested that expression of these genes was changed by ambient temperature changes. The opposite pattern was observed for the temperature response of *embryonic flower 1* (*EMF1*) and *apetala 3* (*AP3*). The expression pattern of *EMF1* could be explained by the function of *LFY* as the repressor, reported by previous studies ^40,41^. On the other hand, LFY was reported to be an activator of *AP3* ^36^. *AP3* was reportedly involved in petal and stamen formation ^42^. LFY was known to bind to *AP3* promoter sequences directly and activate *AP3* transcription with other factors ^43^. Most of these previous experiments analysed the developmental processes of plants grown under constant temperature conditions, therefore, different gene-regulatory mechanisms might be working in the temperature response under fluctuating temperature conditions. Some genes had higher correlation to the temperatures from days prior to sampling. For example, *Calcineurin B-like protein 6* (*CBL6*), *AT hook motif DNA-binding family protein* (*AHL6*) and *nucleolin 2* (*NUC2*) showed significant correlations between their expression and the temperature 1 day prior to sampling (Fig. 8), while the relationships were not significant on sampling day. The expression of *CBL6* was decreased with increased temperatures the day prior to sampling (Fig. 8). *CBL6* has been reported to be involved in cold tolerance in *Stipa purpurea* ^44^. Our results detected ambient-cold-temperature responses of this gene which might occur after relatively delays of 1 day. Another gene, *AHL6*, showed similar expression patterns as *CBL6* (Fig. 8), this gene is involved in regulating hypocotyl growth in seedlings ^45^. The *NUC2* gene is one of the most abundant nucleolar proteins, plays multiple roles in the nucleolus and is involved in several steps of ribosome biogenesis. *NUC2* was also reported to be implicated in DNA replication, methylation, recombination, repair and chromatin organization of rDNA ^46,47^. The temperature responses of *AHL6* and *NUC2* were less known, but our results suggest that their responses to ambient temperatures occur approximately one day post exposure (Fig. 8).

**Figure 7.**
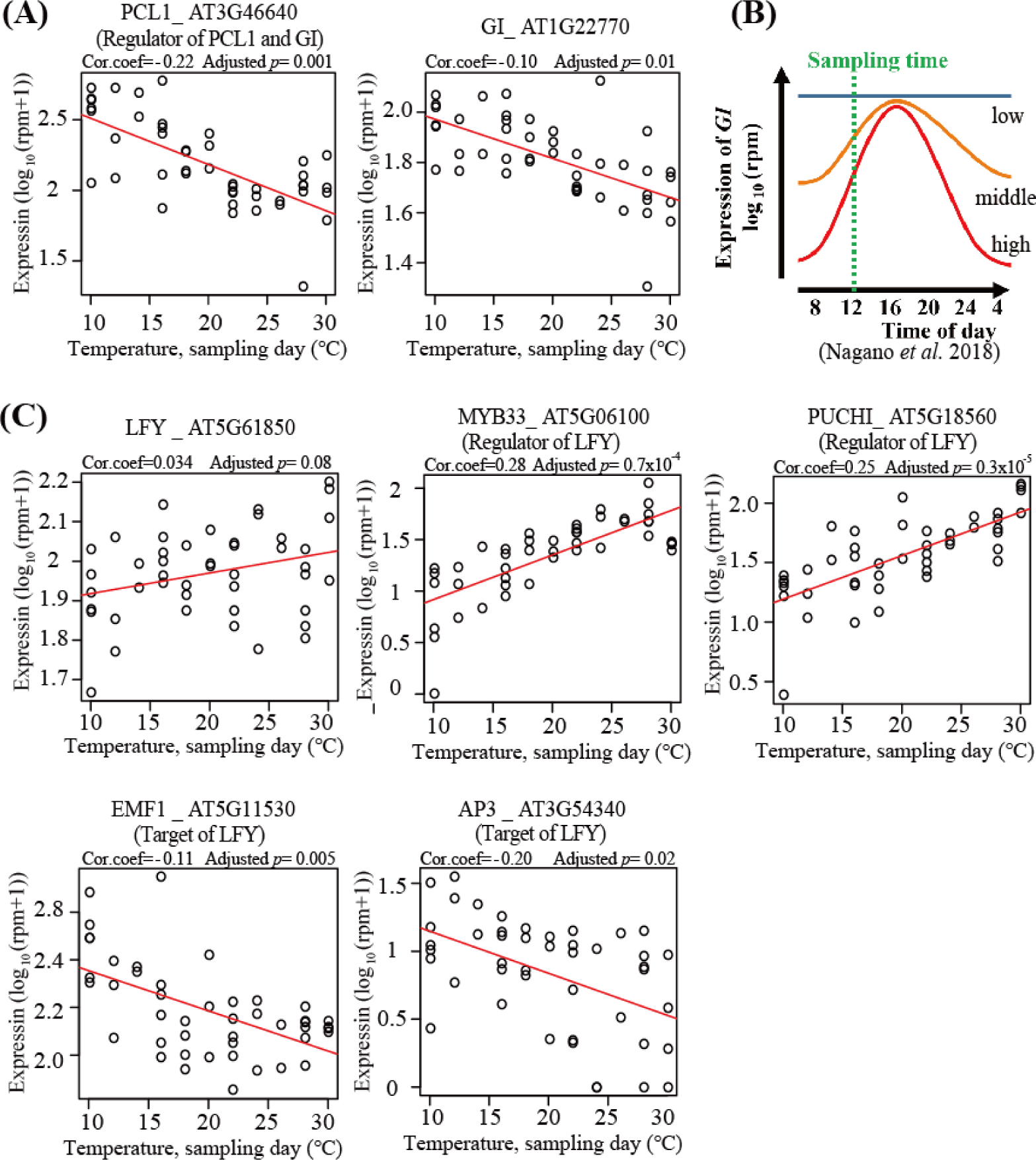
Genes correlated to temperature on sampling day. **(A)** Expression of *PCL1* and *GI* genes. The horizontal axis indicates temperature settings for each sample on sampling day. The vertical axis indicates expression of each gene by log_10_ (rpm+1). Each circle indicates each sample (*n* = 45) and the red lines are regression lines. “Cor.coef” indicates correlation coefficients. **(B)** Schematic diagram of the changes in amplitudes of the circadian oscillations of *GI* correlated to temperature changes reported in a previous study. The lines with “high”, “middle” and “low” represent the circadian oscillations of *GI* under each temperature condition (Nagano *et al*. 2018). A green broken line indicates sampling times in the present study and expression of *GI* at the time became smaller at higher temperatures. **(C)** Expression of *LFY* and the regulator or target genes of *LFY*. Horizontal axis and vertical axis are same as (A).

**Figure 8.**
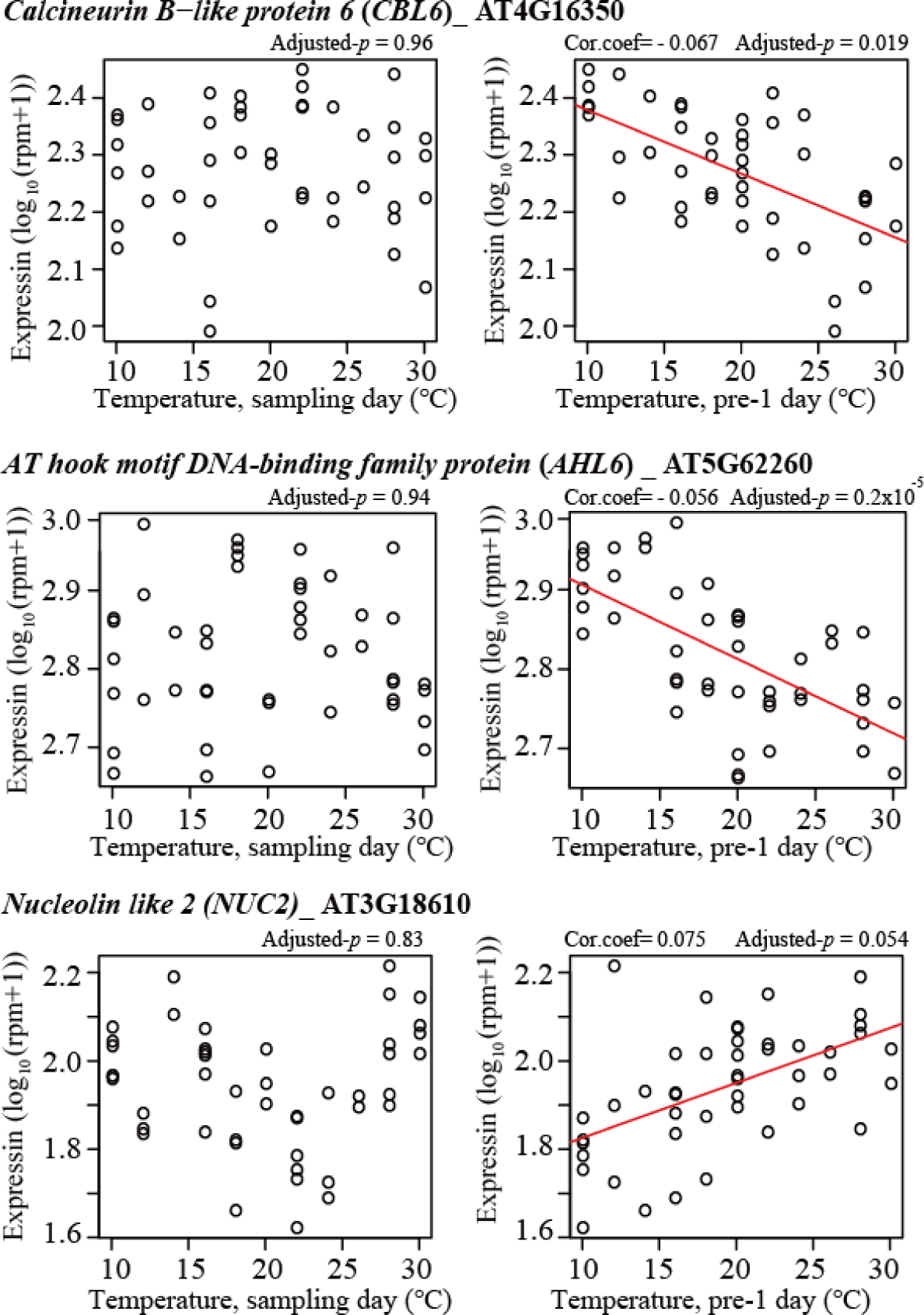
Genes that responded to temperatures one day prior to sampling. Expression levels of *CBL6* (top panel), *FPGS1* (middle panel), and *NUC2* (bottom panel) were plotted. The horizontal axis indicates temperature settings for each sample on each sampling day (left three panels) and one day prior to sampling (right three panels). The vertical axis means show expression for each gene by log_10_ (rpm+1). Each circle indicates each sample (*n* = 45) and red lines are regression lines (drawn only in case of adjusted *p*-value <0.1).

GO enrichment analysis of these temperature-responded in genes revealed genes with GO terms of “intracellular membrane-bounded organelle”, “membrane-bounded organelle”, “intracellular organelle”, “organelle”, “intracellular”, “intracellular part”, “cell” and “cell part” were significantly enriched on the sampling day. One day prior to sampling, “intracellular”, “intracellular part”, “cell” and “cell part” were enriched and no significant enrichments were detected 2 and 3 days prior to sampling. We detected only general GO terms. Genes that we observed in this study may be responding to mild changes in temperature that would not trigger stress-responses.

## Discussion

In this study, we developed a high-throughput RNA-Seq method by simplifying the experimental procedures. By pooling samples after the RT step, Lasy-Seq reduced cost and time compared with those required with previously used methods ^48^. We prepared 192 RT-primers with unique index sequences which enabled sequencing to be conducted in one lane (Supplementary note 1). To pool the more than 192 samples, 2^nd^ index sequences can be added to the libraries by inserting 2^nd^ index sequences into reverse PCR primers, between P5 and Nextera adapter sequences (Supplementary Fig. S1, C). The false assignment rates associated with sample-pooling and caused by pooled-PCR and sequencing were like those reported in previous studies (Supplementary table S3). The false-assignment rates will be affected by the number of PCR cycles; over amplification of libraries is expected to cause higher false-assignment rates. Optimizing PCR cycles is thus necessary for suppressing false-assignment among samples. False-assignment means false detection of reads in a sample from another samples. Considering the false-assignment rates observed in this study (maximum 0.031%), differences in gene expression of larger than approximately 3,000-fold theoretically cannot be detected, because 0.031% of reads from other samples were falsely-assigned. In other words, if 10,000 reads were detected for a gene in a sample, 3.1 reads for the same gene are expected to be falsely-assigned in the other samples sequenced in the same lane. Fasle-assignment cause limitations of dynamic range. For example, the detectable difference of gene expression between samples becomes less than 3225-fold (10,000/3.1). Usually this limit of sensitivity is enough to analyse gene expression changes in the same tissues or plants. However, this sensitivity might be problem in determination of infection by plant viruses, which can produce large amount of reads which exceeds the amount of host total mRNA) in infected samples, and no reads in un-infected samples ^8^. Furthermore, in Lasy-Seq, degradation of RNA-carryover was essential for precise quantification of DNA. Even after RNase treatment, we observed libraries with different length distributions were produced from the same input DNA as from different plant species (data not shown). Therefore, we have recommended to include the optimization step of the input amount for tagmentation. The reason why the length of libraries was different among sample from different species is that GC content of genome or intrinsic--inhibitors of tagmentation may be affecting the reaction.

We applied Lasy-Seq to *A*. *thaliana* to analyse the temperature responses to validate this method, and successfully detected thousands of genes responding to the temperature fluctuations examined in this study. Previous studies reported that phenotypes of mutants can be changed by ambient temperatures. For example in *LFY*, phenotypes of the *lfy-5* mutants became enhanced at 16°C compared with 25°C ^49^. In our study expression of *LFY* and its upstream activators, *MYB33* and *PUCHI*, were positively correlated with the temperature on sampling day and relatively low at lower temperature conditions. Therefore, the low expression levels of *LFY* may result from the low expression levels of these activators, caused by low temperature. To examine responses in gene expression under various temperature-conditions is important to understand plant environmental adaptations. For instance, in our study, genes which responded to temperatures experienced prior to sampling day were also identified by conducting time-course analysis of plants grown under fluctuating-temperature conditions. The correlation between gene expression and past temperatures detected in this study suggests various mechanisms of plant temperature responses with different time scales.

Large–scale transcriptome analysis has recently started and provided new insights into various topics. A previous study analysed transcriptomes of 1,203 samples from 998 accessions of *A*. *thaliana*, and methylomes of 1,107 samples from 1,028 accessions ^50^. Between relict and non-relict accessions, 5,725 differently-expressed genes were determined. Relationships between epialleles and gene expression was analysed and geographic origins were found to be major predictors of altered gene expression caused by the epialleles. Another study conducted transcriptome analyses on 1,785 samples from 7 tissues of 299 maize lines ^51^. They revealed effects of rare genetic alleles on high variance in gene expressions and correlated the variance to fitness ^51^. Their results provided a new insight into evolutionary bottleneck during domestications. In another previous study on plants in natural environments, transcriptome analysis from weekly-samples for 2 years and bihourly-diurnal samples on the four equinoxes/solstices of *A*. *halleri* (873 samples) was conducted ^35^. They identified 2,879 and 7,185 seasonally-and diurnally-oscillating genes, respectively. By shifting the phase of oscillations between temperature and day length, they found that fitness became highest in phase-combinations of natural conditions compared with un-natural conditions. Their results revealed environmental cues that plants actually used for their adaptation to seasonal changes. These studies are cutting edge in this field, and Lasy-Seq will accelerate and generalize large-scale analyses across diverse research topics.

## Methods

### Culture conditions of *Oryza sativa* and *Arabidopsis thaliana*

*Oryza sativa L. japonica* ‘Nipponbare’ was grown for use in the development of our RNA-Seq library preparation method; seeds were sown in germination boxes and approximately one month after germination, fully expanded leaf blades were collected. The leaf samples were immediately frozen by liquid nitrogen and stored at −80 °C until RNA extraction.

*Arabidopsis thaliana* (Col-0, CS70000) was grown for the analysis of temperature responses. Seeds of *A. thaliana* were sown on 1/2 Murashige and Skoog medium with 0.5% gellan gum, incubated for 7 days at 4 °C in the dark, then cultivated for 10 days at 20 °C under 12 hr light / 8 hr dark cycles and a relative humidity of 60%. For the following 11 days, the temperature of the incubator was changed every day, following the designed temperature sets (see Fig. 5 and Results section). Three temperature sets were designed by random sampling from even-numbered temperatures between 10 - 30 °C using a sample function in R 3.4.3 software ^52^. Two replicates of 2 or 3 plant individuals were sampled at 12:00 from the 3rd to 11th day after starting the temperature change (14th to 21st day after sowing). In total 45 samples were collected (Supplementary table S1, see also Fig. 4). Whole plant individuals were collected, immediately frozen by liquid nitrogen and stored at −80 °C until RNA extraction.

### RNA extraction

Samples were ground with zirconia beads (YTZ-4, AS-ONE, Japan), using the TissueLyser II (QIAGEN, MD, USA) with the pre-chilled adapters at −80 °C. Total RNA was extracted by Maxwell 16 LEV Plant RNA Kit (Promega, WI, USA) according to the manufacturer’s instructions. The amount of RNA was determined using Quant-iT RNA Assay Kit broad range (Thermo Fisher Scientific, MA, USA) and Tecan plate reader Infinite 200 PRO (Tecan, Switzerland). The quality was assessed using a Bioanalyzer with Agilent RNA 6000 nano Kit (Agilent Technologies, CA, USA). RNA (5 μg and 500 ng) per sample was used for the library preparations of *O. sativa* and *A. thaliana*, respectively.

### RNA-Seq library preparation

Reverse transcription (RT) of total RNA was performed with oligo-dT primers including index sequences to add a unique index to each sample (RT-indexing, Fig. 1). The RT-indexing primers for single-read sequencing (SR RT-primer in Supplementary Fig. S1.) were designed by modifying RT-primers for paired-end sequencing from a previous study ^16^. RT reactions of the total RNA were conducted with 5.0 μL of RNA in nuclease-free water, 1 μL of 2 μM RT primer, 0.4 μL of 25 mM dNTP (Thermo Fisher Scientific, USA), 4.0 μL of 5X SSIV Buffer (Thermo Fisher Scientific), 2.0 μL of 100 mM DTT (Thermo Fisher Scientific), 0.1 μL of SuperScript IV reverse transcriptase (200 U/μL, Thermo Fisher Scientific), 0.5 μL of RNasin Plus (Ribonuclease Inhibitor, Promega) and nuclease-free water (7.0 μL) to make up the volume to 20 μL. Reverse transcription was carried out at 62°C for 50 min (or 65°C for 10 min for more severe suppression of RT of rRNA), then incubated at 80°C for 15 min to inactivate the enzyme. All indexed samples were then pooled and purified with the same volume of AMPure XP beads (Beckman Coulter, USA) or column purification with Zymo spin column I (Zymo Research) and Membrane Binding Solution (Promega). If the number of samples was large, pooling of the RT products could be conducted by centrifuging the reaction plate set on a one well reservoir as described in a previous study ^15^. The purified cDNA was dissolved in 10 μL (depending on number of pooled-samples) of nuclease-free water.

Second strand synthesis was conducted on the pooled samples (10 μL) with 2 μL of 10X blue buffer (enzymatics, USA), 1 μL of 2.5 mM dNTP (Takara Bio, Japan), 0.5 μL of 100 mM DTT, 0.5 μL of RNaseH (5 U/μL, enzymatics), 1.0 μL of DNA polymerase I (10 U/μL, enzymatics) and nuclease-free water (5 μL) to make up the volume to 20 μL. Reactions were conducted at 16 °C for 2 hours and kept at 4°C until the next reaction. To avoid the carryover of large amounts of RNA, RNase T1 treatments were conducted on the double-stranded DNA with 1 µL of RNase T1 (more than 1 U/µL, Thermo Fisher Scientific, MA, UK). The reaction was conducted at 37°C for 30 min, 95°C for 10 min, gradual-decreases in temperature from 95°C to 45°C (−0.1 °C/s), 25°C for 30 min and 4°C until the next reaction. Alternatively, reactions of 37°C for 5 min with mixtures of RNaseA (10 μg/mL) and RNaseT (1 U/µL) were enough to remove RNA in the samples. The DNA was purified with 20 µL AMPure XP beads and eluted with 10 µL nuclease free water. Alternatively, for many samples, the AMPure bead purification could be replaced by column purification using a Zymo spin column I (Zymo Research) and Membrane Binding Solution (Promega). The DNA was then quantified by QuantiFluor dsDNA System and Quantus Fluorometer (Promega).

Tagmentation by transposases was conducted on the purified DNA, using 5 μL Nextera TD buffer and 0.5 μL TDE1 enzyme (Nextera DNA Sample Preparation kit, Illumina). The optimization of the amount of input DNA (usually between 3 ng and 8 ng) should be conducted for each pooled-sample to construct libraries with an average length of 500 bp; 4 ng, 6 ng, and 8 ng were tested here. Because in libraries with shorter size distribution, sequencing-reads were reached to poly-A sequences at the 3’ end of the insert, which were not informative for quantification of gene expression. Library distributions from 200 bp to 1500 bp with an average length of 500 bp efficiently avoid to read poly-A sequences. Reactions were carried out at 55 °C for 5 min, then stopped by adding 12 μL DNA binding buffer in DNA clean & concentrator kit (Zymo Research). The tagmented library was immediately purified using a Zymo spin column II (Zymo Research) following the manufacturer’s instructions. This purification with Zymo spin column II cannot be replaced by purification with AMPure XP beads or NucleoSpin Gel and PCR Clean-up (Takara Bio), in which final yield of the library was largely decreased.

To determine an optimal number of cycles for the amplification, 2 μL of the tagmented DNA was amplified using KAPA Real-time Library Amplification Kit (KAPA), conducted with 2 μL of the RNA with 5 µL of 2x KAPA HiFi HotStart Real-time PCR Master Mix, 0.5 μL of 10 μM PCR forward-primer, 0.5 μL of 10 μM PCR reverse-primer (Supplementary Fig. S1) and 2 μL of water to make it up to 10 µL. Reactions were carried out at 95°C for 5 min, 30 cycles of 98°C for 20 sec, 60°C for 15 sec, 72°C for 40 sec, followed by 72°C for 3 min, then held at 4 °C. Samples (10 µL) of standards were analysed together and optimal cycles were determined following the manufacturer’s instruction.

The optimized PCR cycles were used for the amplification of the library with 2 μL of the tagmented DNA. Sufficient quantity and diversity of libraries for sequencing was achieved with 2 or 3 replicates of PCR that were pooled after the amplification. The libraries were purified twice with the same volume of AMPure XP beads and dissolved in 20 μL of nuclease-free water. Quantification of the library was conducted using QuantiFluor dsDNA System and Quantus Fluorometer (Promega). The size distribution of the libraries were analysed by the Bioanalyzer with high sensitivity DNA kits (Agilent Technologies, CA, USA) and optimal input amount of DNA were determined. Tagmentation reactions with the optimized input amounts of DNA were conducted in triplicate to reduce PCR cycles in library amplification. The tagmented DNA was eluted in 15 μL of nuclease-free water. All three reaction solutions were pooled after the purification.

To construct libraries for paired-end sequencing, the required modifications were as follows. Reverse transcription should be carried out with the RT-indexing primers for paired-end sequencing (Supplementary Fig. S1). Library amplification was carried out with primers for paired-end sequencing libraries (Supplementary Fig. S1). Temperatures for PCR reactions were same as described above. The protocol with detailed notes is summarised in Supplemental note 1.

### Sequencing

Libraries of *O. sativa* for development of the RNA-Seq library preparation method were constructed with the protocol for paired-end sequencing described above. The libraries were sequenced by PE 75 sequencing with MiSeq with MiSeq Reagent Kit v3 (150 cycles, Illumina). Libraries of *A. thaliana* for analysis of temperature responses were prepared with the protocol for single-read sequencing. Single-read 50 bases and index sequencing were conducted for the libraries using HiSeq 2500 (Illumina) with the TruSeq SBS kit v3 platform conducted by Macrogen Japan Co. For sequencing of libraries prepared by the methods described in this study, we recommend the use of the Illumina platform with non-patterned flow cell such as HiSeq 2500 or MiSeq sequencer (Illumina). The concentration of the libraries produced with Lasy-Seq were sometimes over-estimated, smaller inputs of libraries than the manufacture recommends can improve results.

### Mapping and quantification of short-read sequences

Details of the pre-processing, mapping and quantification processes were described previously (Supplementary Fig. S3) ^8^. FASTQ files from RNA-Seq were pre-processed by removing adapter sequences and low-quality bases using trimmomatic-0.32 as described in previous works ^8,53^. The reference transcriptome sequences of *A. thaliana* and *O. sativa* were prepared from the Arabidopsis Information Portal (Araport 11) and The Rice Annotation Project database ^54,55^. In addition, External RNA Controls Consortium spike-in control (ERCC-control) sequences (92 genes, Thermo Fisher Scientific) were also used as reference sequences. The pre-processed sequences were mapped on each reference and quantified using RSEM-1.2.15 as described in previous work ^8,56^. We subtracted 0.05% of the total reads to avoid false assignment caused by the Illumina platforms analyser as described in a previous study ^8^. This subtraction was not conducted for the analysis on false-assignment rates shown in Fig. 4.

### Analysis of false-assignment rates among pooled samples

To estimate the false-assignment rates, which may be caused by early pooling of libraries, we prepared 5 μg of *O. sativa* RNA samples with and without 40 ng ERCC-control. We prepared in total 8 RNA samples from *O. sativa*. Four of them were reverse transcribed with PE60 RT-primer (60 mer primer sets in Supplementary Fig. S1) and the other four were reverse transcribed with PE78 RT-primer (78 mer primer sets in Supplementary Fig. S1) for paired-end sequencing (Fig. 4 and Supplementary Fig. S1). For each primer set, samples with and without ERCC-control were pooled before amplification (early-pooled sets) and sequencing (late-pooled sets) to estimate the false-assignment rate caused by PCR and sequencing (Fig. 4). Until the pooling steps, samples were separately prepared and all 8 samples were pooled before sequencing. After sequencing, the number of ERCC-control reads in each sample were determined as described above.

Uniquely mapped reads with a mapping quality value of ≥4 were generated using samtools and 5.0 x 10^5^ reads were used for the following analysis. The rates of false-assignment caused by pooled-PCR or sequencing steps were calculated from the numbers of ERCC-control reads in samples with and without ERCC-control (Supplementary Fig. S2). Briefly, ERCC reads detected in the late-pooled samples (without ERCC addition) were regarded as false-assignments caused by sequencing of each sample. Therefore, the rate of total false-assignment reads in all eight samples against total ERCC reads in the lane was estimated to be the false-assignment rate caused by sequencing (Supplementary Fig. S2). The false-assignment rate caused by pooled-PCR was estimated from the ERCC-reads number detected in early-pooled samples (without ERCC addition), as explained in Supplementary Fig. S2.

### Estimate deviation between technical replicates in Lasy-Seq

Correlation coefficient between the early-pooled samples were calculated using rpm except for ERCC-controls to estimate deviation between technical replicates. Pearson’s correlation coefficiency was calculated with cor function in R version 3.5.0^52^.

### Analysis of temperature response in *A*. *thaliana*

Samples with fewer than 10^5^ reads and genes on which fewer than 1 read were mapped on average were excluded from the analysis. For the remaining genes (26,082 genes in 45 samples), single regression analyses were conducted on gene expression (number of normalized-reads, rpm) and temperatures for each day; sampling day, 1 day before the sampling day (pre-1 day), 2 days before the sampling day (pre-2 day) and 3 days before the sampling day (pre-3 day). Correlations were tested with lm function in R. Multiple testing corrections were performed by setting the False Discovery Rate (FDR) using the p.adjust function with BH (FDR) method in R ^57^. Genes with adjusted-*p* value of less than 0.1 were thought to have significant correlation to each temperature. Gene Ontology annotations were obtained from The Arabidopsis Information Resource (TAIR) 10 ^58^. Existence of significant enrichment of particular GO terms were tested (Fisher’s exact test). Multiple testing corrections were performed by p.adjust functions with BH (FDR) method in R.

## Supporting information

Supplementary informations

## Acknowledgements

We thank N. Yamaguchi for his comment on temperature response in *A. thaliana*. We also thank M. Mihara, H. Ooshima, K. Iwayama, Y. Kurita, Y. Hashida and F. Kobayashi for their support on data analysis and material preparations. This work was supported by JSPP KAKENHI Grant Numbers 16H06171, JP16H01473, JST CREST Grant Number JPMJCR15O2 and JST ACCEL Grant Number JPMJAC1403 to AJN.

## Author contribution

M. Kamitani and A.J.N designed the research. M. Kamitani M. Kashima and A.T. conducted the laboratory experiment. M. Kamitani, M. Kashima and A.J.N. conducted the data analysis. M. Kamitani wrote the manuscript. All authors discussed the results and approved the manuscript.

## Data availability

Sequence data from RNA-Seq were deposited in Sequence Read Archive (SRA). The accession numbers are PRJNA508267 (*O. sativa* and *A. thaliana*).

## List of Supplementary Information

Supplementary note 1 The protocol of Lasy-Seq with detailed notes

Supplementary Fig. S1 Primers used in the present study.

Supplementary Fig. S2 Method for calculation of false-assignment rates

Supplementary Fig. S3 Overview of the analysis of RNA-Seq data

Supplementary table S1 Information on the samples collected in this study (*n* = 45)

Supplementary table S2 List of genes significantly correlated to temperature on each day

Supplementary table S3 Summary of the false-assignment rates reported by previous studies

