## Supplementary informations for "Lasy-Seq: a high-throughput library preparation method for RNA-Seq and its application in the analysis of plant responses to fluctuating temperatures"

for

This PDF file includes:

Supplementary note 1　The protocol of Lasy-Seq with detailed notes

Supplementary Fig S1. Primers used in the present study.

Supplementary Fig S2. Overview of the analysis of RNA-Seq data

Supplementary Fig S3. Method for calculation of false-assignment rates

Supplementary table S1 Information on the samples collected in this study ($n$ = 45)

Supplementary table S2 List of genes significantly correlated to temperature on each day

Supplementary table S3 Summary of the false-assignment rates reported by previous studies

### Supplementary note 1: The protocol of Lasy-Seq with detailed notes

#### <u>Things to know before you start</u>

- ✓ This is a protocol for high-throughput analysis. For a smaller number of samples, optimization is required for the input-amount of RNA and volume of elution-solution in each reaction.
- ✓ This protocol does not require an mRNA-enrichment step. The RT reaction is started from total RNA, after second strand synthesis, RNase treatment is required to remove the large amounts of RNA.
- ✓ We usually conduct single-read sequencing for the quantification of gene expression. Paired-end sequencing can also be used for more precision analysis, because information of UMI is available.
- ✓ The concentration of the libraries produced with Lasy-Seq are sometimes over estimated. In such cases, smaller inputs of libraries into sequencing than Illumina recommends improves results.

This protocol will be updated at the following website as required.

https://sites.google.com/view/lasy-seq/home

#### 1. Reverse Transcription

1-1. Prepare total RNA samples in the PCR plate.

| | For each sample | Master mix (for example, *n*=96) |
|---|---|---|
| ➢ RNA 500 ng/sample | X μL | |
| 1-2. Assemble the master mix as follows. | For each sample | If using 5 μL of 100 ng/μL RNA, |
| ➢ 5X Superscript IV First-Strand Buffer | 4.0 μL | 384 μL |
| ➢ RNasin Plus | 0.5 μL | 48 μL |
| ➢ DTT (100 mM) | 2.0 μL | 192 μL |
| ➢ dNTP (25 mM each) | 0.4 μL | 38.4 μL |
| ➢ SuperScript IV reverse transcriptase | 0.1 μL | 9.6 μL |
| ➢ Nuclease-free water | 12 - X μL | 672 μL |

(Master mix: Dispense 14 μL of master mix to each sample)

1-3. Add 19- X μL of master mix to each RNA sample.

1-4. Add 1μL of 2μM SE RT-primer (with index-sequences) to each sample.

1-5. Incubate the plate at 65 °C for 10 min, 80 °C for 15 min and keep the sample at 4°C until the next step.

> ***Note:*** *Total input RNA (pooled from all samples) is recommended to be more than 10 μg, for example, we successfully constructed libraries with 20 samples of 500 ng RNA for each sample. If the number of samples is small, larger amounts of input RNA should be prepared and vice versa.*

#### 2. Purification

2-1. Pool all of the solutions into a tube.

2-2. Place the RT reaction plate on the one well reservoir (Sasagawa et al. 2018. *Genome Biol*). Centrifuge the plate and collect the solution into the one well reservoir as described in the previous study.

2-3. Collect the solution into a 5mL tube and add 2x Membrane binding solution (Promega) and mix well with voltex mixer. Set the Zymo-Spin Column I (Zymo research) on the Vac-Man Laboratory Vacuum Manifold (Promega) and filtrate the collected solution. Wash twice with 700 μL of wash buffer, mixture of ethanol (final conc. 80%) and Tris-HCl (pH 7.6, final conc. 100 mM). Put the column on a new tube and centrifuge it at 16,000 g at 4 °C for 2 min to remove the wash buffer. Put the column on a new tube and elute the RNA/cDNA hybrid by adding 100 μL of nuclease-free water.

*This step can be replaced by AMpure XP beads purification.*

#### 3. Second Strand Synthesis

3-1. Assemble the following mix.

| | | |
|---|---|---|
| ➢ RNA/cDNA hybrid | 30.0 μL | (Store the rest 70μL at -20°C) |
| ➢ 10x Blue buffer | 4.0 μL | |
| ➢ dNTP mix (2. 5 mM each) | 2.0 μL | |
| ➢ DTT (100 mM) | 1.0 μL | |
| ➢ RNase H | 1.0 μL | |
| ➢ DNA polymerase I | 2.0 μL | |

3-2. Incubate at 16 °C for 2 h and keep at 4°C until the next step.

#### 4. RNase treatment

4-1. Add 2 µL of mixture of 1/1000x RNase T1 (1000 U/µL) and RNase A (10 mg/mL).

4-2. Incubate at 37 °C for 5 min and keep at 4°C until the next step.

4-3. Bind the RNA/cDNA hybrid using 0.8x volume of AMPure XP beads. Purify them following to the manufacturer's instructions. Elute the RNA/cDNA hybrid by adding 10 µL of nuclease-free water.
*This purification with AMPure XP beads can be replaced by column purification using Zymo spin column I and Membrane Binding Solution.*

#### 5. Quantification of dsDNA

5-1. Quantify the cDNA. (For example with QuantiFluor dsDNA System and Quantus Fluorometer (Promega)).

#### 6. Optimization of tagmentation of dsDNA

6-1. Assemble the following mix.

- dsDNA — X µL (Test 3 concentration between 3 and 8 ng, for example, 4ng, 6ng and 8ng)
- Tagment DNA buffer (TD) — 5.0 µL
- Tagment DNA Enzyme (TDE1) — 0.5 µL
- Nuclease-free water — 4.5-X µL

6-2. Incubate at 55 °C for EXACT 5 min.

6-3. Immediately add 50 µL (5x volumes) of DNA Binding Buffer to the cDNA samples. Mix briefly by vortexing
and transfer mixture to a Zymo-Spin Column II set on a new collection tube. Centrifuge at 14,000g at room temperature for 30 sec. Discard the flow-through.

6-4. Add 200 µl DNA Wash Buffer to each column. Centrifuge at 14,000g at room temperature for 30 sec. Repeat this wash step again.

6-5. Transfer the column to a 1.5 ml tube Add 18 µL water directly to the column matrix and incubate at room temperature for 1 minute. Centrifuge at 14,000g at room temperature for 30 sec to elute the dsDNA.

> ***Note:*** *This optimization step of input-cDNA amounts is necessary, because in libraries with shorter size distributions, sequencing-reads were reached to poly-A sequences at the 3' end of the insert. Libraries distributed from 200 bp to 1500 bp with the average length of 500 bp were efficient enough to decrease the amount of poly-A sequences in data reads.*
> *The purification step after tagmentation cannot be replaced by purification with AMPure XP beads or NucleoSpin Gel and PCR Clean-up (Takara bio, Japan), in which final yields of the library were largely decreased.*

#### 7. Optimization of the number of PCR cycles

7-1. Assemble the following mix.

- Tagmented DNA — 4.0 µL (for each 3 templates prepared above)

- HiFi HotStart Ready MIX 5.0 µL (for each 3 templates prepared above)
- SE PCR forward-primer (10 µM) 0.5 µL
- SE PCR reverse-primer (10 µM) 0.5 µL

7-2. Add two replicate standards (10 µL) to the wells in the PCR plate following the manufacturer's instructions.

7-3. Incubate at 95°C for 5 min, 30 cycles of 98°C for 20 sec, 60°C for 15 sec, 72°C for 40 sec, followed by 72°C for 3 min and hold at 4 °C until the next step.

7-4. Determine the optimal cycle number by comparing to standards. We usually select the cycle number of 2 or 3 cycles smaller than the cycle at the center of the amplification curve (red-brake line in the figure below).

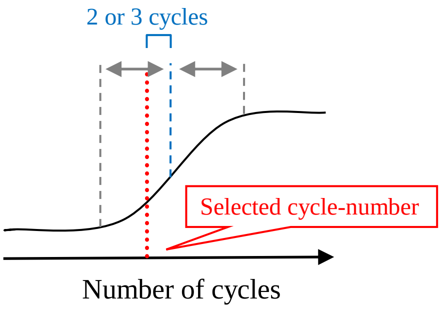


#### 8. Amplification of libraries with optimized PCR cycles

8-1. Assemble the following mix.

- Tagmented DNA 12.0 µL (for each 3 templates prepared above)
- 2x KAPA-HiFi HS Ready MIX 15.0 µL
- SE PCR forward-primer (10 µM) 1.5 µL
- SE PCR reverse-primer (10 µM) 1.5 µL

8-2. Incubate at 95°C for 5 min, optimized cycles of 98°C for 20 sec, 60°C for 15 sec, 72°C for 40 sec followed by 72°C for 3 min and hold at 4 °C until the next step.

8-3. Bind the RNA/cDNA hybrid using same volume of AMPure XP beads. Purify them following the manufacturer’s instructions. Elute the RNA/cDNA hybrid by adding 8 µL of nuclease-free water.

#### 9. Analysis on length-distribution of libraries

9-1. Analyze the length-distribution of three purified-libraries by Bioanalyzer with a high sensitivity DNA kit (Agilent Technologies, CA, USA). Determine the amount of cDNA input (ng) that produced libraries distributed from 200bp to 1500bp with the average length of 500bp, and select this as an optimized condition. A representative of recommended library distribution is shown in the figure below.

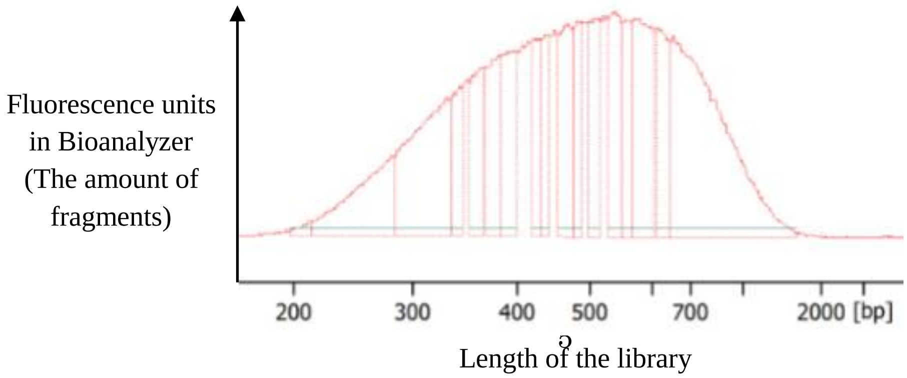

#### 10. Tagmentation of ds-cDNA with optimized input cDNA

10-1. Assemble the following mix **in triplicate**.

- dsDNA X µL (Use optimized amount of input dsDNA)
- Tagment DNA buffer (TD) 5.0 µL
- Tagment DNA Enzyme (TDE1) 0.5 µL
- Nuclease-free water 4.5-X µL

10-2. Incubate at 55 °C for EXACTLY 5 min.

10-3. Immediately add 50 µL (5x volumes) of DNA Binding Buffer to the cDNA samples and mix briefly by vortexing. Pool the three reaction solutions into a tube. Purify the tagmented DNA with a Zymo-Spin Column II as mentioned above. Elute the dsDNA with 54 µL of nuclease-free water.

#### 11. Amplification of libraries with optimized PCR cycles

11-1. Assemble the following mix **in triplicate**.

- Tagmented DNA 16.0 µL (for each 3 templates prepared above)
- 2x KAPA-HiFi HS Ready MIX 20.0 µL (for each 3 templates prepared above)
- SE PCR forward-primer (10 µM) 2.0 µL
- SE PCR reverse-primer (10 µM) 2.0 µL

11-2. Incubate at 95°C for 5minutes, optimized cycle-number of 98°C for 20 sec, 60°C for 15 sec, 72°C for 40 sec, followed by 72°C for 3 min and hold at 4 °C until the next step.

11-3. Bind the RNA/cDNA hybrid using 0.8x volume of AMPure XP beads. Purify them following to the manufacturer's instructions. Elute the RNA/cDNA hybrid by adding 20 µL of nuclease-free water. Pool the triplicated samples.

#### 12. Analysis on length-distribution and concentration of libraries

12-1. Analyze the length-distribution of three purified-libraries by Bioanalyzer with high sensitivity DNA kit (Agilent Technologies, CA, USA).
Quantify the library using QuantiFluor dsDNA System and Quantus Fluorometer (Promega).

12-2. Sequencing of the library. We recommend the use of the Illumina platform with non-patterned flow cells such as HiSeq 2500 or MiSeq sequencer (Illumina) to suppress false-assignment among samples.

### Primer sequences used in this protocol

#### PCR primers

| | |
|---|---|
| SE PCR forward-primer | CAAGCAGAAGACGGCATACGAGAT |
| SE PCR reverse-primer | AATGATACGGCGACCACCGAGATCTACACTCGTCGGCAGCGTC |

#### SE RT-primers (5’-3’) Index1~96, Index101~196

“n” in the sequence indicates unique molecular identifier.

| | |
|---|---|
| Index 001 | CAGAAGACGGCATACGAGATAACTTTAGTGACTGGAGTTCAGACGTGTGCTCTTCCGATCNNNNNNTTTTTTTTTTTTTTTTTTV |
| Index 002 | CAGAAGACGGCATACGAGATCATAGAGTGTGACTGGAGTTCAGACGTGTGCTCTTCCGATCNNNNNNTTTTTTTTTTTTTTTTTTV |
| Index 003 | CAGAAGACGGCATACGAGATTGCTGGAGGTGACTGGAGTTCAGACGTGTGCTCTTCCGATCNNNNNNTTTTTTTTTTTTTTTTTTV |
| Index 004 | CAGAAGACGGCATACGAGATTTGGTGACGTGACTGGAGTTCAGACGTGTGCTCTTCCGATCNNNNNNTTTTTTTTTTTTTTTTTTV |
| Index 005 | CAGAAGACGGCATACGAGATGTCCACGTGTGACTGGAGTTCAGACGTGTGCTCTTCCGATCNNNNNNTTTTTTTTTTTTTTTTTTV |
| Index 006 | CAGAAGACGGCATACGAGATGAGACGGAGTGACTGGAGTTCAGACGTGTGCTCTTCCGATCNNNNNNTTTTTTTTTTTTTTTTTTV |
| Index 007 | CAGAAGACGGCATACGAGATCCAGTATCGTGACTGGAGTTCAGACGTGTGCTCTTCCGATCNNNNNNTTTTTTTTTTTTTTTTTTV |
| Index 008 | CAGAAGACGGCATACGAGATCCGATCTTGTGACTGGAGTTCAGACGTGTGCTCTTCCGATCNNNNNNTTTTTTTTTTTTTTTTTTV |
| Index 009 | CAGAAGACGGCATACGAGATTGCGAGCCGTGACTGGAGTTCAGACGTGTGCTCTTCCGATCNNNNNNTTTTTTTTTTTTTTTTTTV |
| Index 010 | CAGAAGACGGCATACGAGATATTCGTTAGTGACTGGAGTTCAGACGTGTGCTCTTCCGATCNNNNNNTTTTTTTTTTTTTTTTTTV |
| Index 011 | CAGAAGACGGCATACGAGATTATTCGTAGTGACTGGAGTTCAGACGTGTGCTCTTCCGATCNNNNNNTTTTTTTTTTTTTTTTTTV |
| Index 012 | CAGAAGACGGCATACGAGATCGGCCCAGGTGACTGGAGTTCAGACGTGTGCTCTTCCGATCNNNNNNTTTTTTTTTTTTTTTTTTV |
| Index 013 | CAGAAGACGGCATACGAGATAATTGGCGGTGACTGGAGTTCAGACGTGTGCTCTTCCGATCNNNNNNTTTTTTTTTTTTTTTTTTV |
| Index 014 | CAGAAGACGGCATACGAGATCGCCAGGGGTGACTGGAGTTCAGACGTGTGCTCTTCCGATCNNNNNNTTTTTTTTTTTTTTTTTTV |
| Index 015 | CAGAAGACGGCATACGAGATAGATTCCGGTGACTGGAGTTCAGACGTGTGCTCTTCCGATCNNNNNNTTTTTTTTTTTTTTTTTTV |
| Index 016 | CAGAAGACGGCATACGAGATTATCGAGAGTGACTGGAGTTCAGACGTGTGCTCTTCCGATCNNNNNNTTTTTTTTTTTTTTTTTTV |
| Index 017 | CAGAAGACGGCATACGAGATTCCATGACGTGACTGGAGTTCAGACGTGTGCTCTTCCGATCNNNNNNTTTTTTTTTTTTTTTTTTV |
| Index 018 | CAGAAGACGGCATACGAGATACCACATCGTGACTGGAGTTCAGACGTGTGCTCTTCCGATCNNNNNNTTTTTTTTTTTTTTTTTTV |
| Index 019 | CAGAAGACGGCATACGAGATTAGTGTAAGTGACTGGAGTTCAGACGTGTGCTCTTCCGATCNNNNNNTTTTTTTTTTTTTTTTTTV |
| Index 020 | CAGAAGACGGCATACGAGATCCGGGCGTGTGACTGGAGTTCAGACGTGTGCTCTTCCGATCNNNNNNTTTTTTTTTTTTTTTTTTV |
| Index 021 | CAGAAGACGGCATACGAGATACCTGACGGTGACTGGAGTTCAGACGTGTGCTCTTCCGATCNNNNNNTTTTTTTTTTTTTTTTTTV |
| Index 022 | CAGAAGACGGCATACGAGATCTAGGCAGGTGACTGGAGTTCAGACGTGTGCTCTTCCGATCNNNNNNTTTTTTTTTTTTTTTTTTV |
| Index 023 | CAGAAGACGGCATACGAGATGGAACGCCGTGACTGGAGTTCAGACGTGTGCTCTTCCGATCNNNNNNTTTTTTTTTTTTTTTTTTV |
| Index 024 | CAGAAGACGGCATACGAGATGGCATTTTGTGACTGGAGTTCAGACGTGTGCTCTTCCGATCNNNNNNTTTTTTTTTTTTTTTTTTV |
| Index 025 | CAGAAGACGGCATACGAGATCGTGGATAGTGACTGGAGTTCAGACGTGTGCTCTTCCGATCNNNNNNTTTTTTTTTTTTTTTTTTV |
| Index 026 | CAGAAGACGGCATACGAGATCCGACATGGTGACTGGAGTTCAGACGTGTGCTCTTCCGATCNNNNNNTTTTTTTTTTTTTTTTTTV |
| Index 027 | CAGAAGACGGCATACGAGATTCCGCTTAGTGACTGGAGTTCAGACGTGTGCTCTTCCGATCNNNNNNTTTTTTTTTTTTTTTTTTV |
| Index 028 | CAGAAGACGGCATACGAGATATTGGAGGGTGACTGGAGTTCAGACGTGTGCTCTTCCGATCNNNNNNTTTTTTTTTTTTTTTTTTV |

| | |
|---|---|
| Index 029 | CAGAAGACGGCATACGAGATTTGACAGCGTGACTGGAGTTCAGACGTGTGCTCTTCCGATCNNNNNNTTTTTTTTTTTTTTTTTTV |
| Index 030 | CAGAAGACGGCATACGAGATCCCAATCGGTGACTGGAGTTCAGACGTGTGCTCTTCCGATCNNNNNNTTTTTTTTTTTTTTTTTTV |
| Index 031 | CAGAAGACGGCATACGAGATTAGAAGCCGTGACTGGAGTTCAGACGTGTGCTCTTCCGATCNNNNNNTTTTTTTTTTTTTTTTTTV |
| Index 032 | CAGAAGACGGCATACGAGATAGAAGTGAGTGACTGGAGTTCAGACGTGTGCTCTTCCGATCNNNNNNTTTTTTTTTTTTTTTTTTV |
| Index 033 | CAGAAGACGGCATACGAGATACATCATAGTGACTGGAGTTCAGACGTGTGCTCTTCCGATCNNNNNNTTTTTTTTTTTTTTTTTTV |
| Index 034 | CAGAAGACGGCATACGAGATGGTTAAAAGTGACTGGAGTTCAGACGTGTGCTCTTCCGATCNNNNNNTTTTTTTTTTTTTTTTTTV |
| Index 035 | CAGAAGACGGCATACGAGATCAGTGGCCGTGACTGGAGTTCAGACGTGTGCTCTTCCGATCNNNNNNTTTTTTTTTTTTTTTTTTV |
| Index 036 | CAGAAGACGGCATACGAGATATTTAGGCGTGACTGGAGTTCAGACGTGTGCTCTTCCGATCNNNNNNTTTTTTTTTTTTTTTTTTV |
| Index 037 | CAGAAGACGGCATACGAGATGAGGGCCCGTGACTGGAGTTCAGACGTGTGCTCTTCCGATCNNNNNNTTTTTTTTTTTTTTTTTTV |
| Index 038 | CAGAAGACGGCATACGAGATCTCGTATTGTGACTGGAGTTCAGACGTGTGCTCTTCCGATCNNNNNNTTTTTTTTTTTTTTTTTTV |
| Index 039 | CAGAAGACGGCATACGAGATGGATCGTGGTGACTGGAGTTCAGACGTGTGCTCTTCCGATCNNNNNNTTTTTTTTTTTTTTTTTTV |
| Index 040 | CAGAAGACGGCATACGAGATCACTAATAGTGACTGGAGTTCAGACGTGTGCTCTTCCGATCNNNNNNTTTTTTTTTTTTTTTTTTV |
| Index 041 | CAGAAGACGGCATACGAGATCCTTCGCTGTGACTGGAGTTCAGACGTGTGCTCTTCCGATCNNNNNNTTTTTTTTTTTTTTTTTTV |
| Index 042 | CAGAAGACGGCATACGAGATAGATGGTCGTGACTGGAGTTCAGACGTGTGCTCTTCCGATCNNNNNNTTTTTTTTTTTTTTTTTTV |
| Index 043 | CAGAAGACGGCATACGAGATGATAGGTAGTGACTGGAGTTCAGACGTGTGCTCTTCCGATCNNNNNNTTTTTTTTTTTTTTTTTTV |
| Index 044 | CAGAAGACGGCATACGAGATAGCCGCTAGTGACTGGAGTTCAGACGTGTGCTCTTCCGATCNNNNNNTTTTTTTTTTTTTTTTTTV |
| Index 045 | CAGAAGACGGCATACGAGATCTTTGTCAGTGACTGGAGTTCAGACGTGTGCTCTTCCGATCNNNNNNTTTTTTTTTTTTTTTTTTV |
| Index 046 | CAGAAGACGGCATACGAGATACGCCACTGTGACTGGAGTTCAGACGTGTGCTCTTCCGATCNNNNNNTTTTTTTTTTTTTTTTTTV |
| Index 047 | CAGAAGACGGCATACGAGATTAGACTGTGTGACTGGAGTTCAGACGTGTGCTCTTCCGATCNNNNNNTTTTTTTTTTTTTTTTTTV |
| Index 048 | CAGAAGACGGCATACGAGATGTTCTCAAGTGACTGGAGTTCAGACGTGTGCTCTTCCGATCNNNNNNTTTTTTTTTTTTTTTTTTV |
| Index 049 | CAGAAGACGGCATACGAGATAGCGTGACGTGACTGGAGTTCAGACGTGTGCTCTTCCGATCNNNNNNTTTTTTTTTTTTTTTTTTV |
| Index 050 | CAGAAGACGGCATACGAGATGCGTCTACGTGACTGGAGTTCAGACGTGTGCTCTTCCGATCNNNNNNTTTTTTTTTTTTTTTTTTV |
| Index 051 | CAGAAGACGGCATACGAGATTTACCAGGGTGACTGGAGTTCAGACGTGTGCTCTTCCGATCNNNNNNTTTTTTTTTTTTTTTTTTV |
| Index 052 | CAGAAGACGGCATACGAGATGCCAGTTGGTGACTGGAGTTCAGACGTGTGCTCTTCCGATCNNNNNNTTTTTTTTTTTTTTTTTTV |
| Index 053 | CAGAAGACGGCATACGAGATGGTCACGGGTGACTGGAGTTCAGACGTGTGCTCTTCCGATCNNNNNNTTTTTTTTTTTTTTTTTTV |
| Index 054 | CAGAAGACGGCATACGAGATGACAAGATGTGACTGGAGTTCAGACGTGTGCTCTTCCGATCNNNNNNTTTTTTTTTTTTTTTTTTV |
| Index 055 | CAGAAGACGGCATACGAGATGCCAGAAAGTGACTGGAGTTCAGACGTGTGCTCTTCCGATCNNNNNNTTTTTTTTTTTTTTTTTTV |
| Index 056 | CAGAAGACGGCATACGAGATACAGAGGCGTGACTGGAGTTCAGACGTGTGCTCTTCCGATCNNNNNNTTTTTTTTTTTTTTTTTTV |
| Index 057 | CAGAAGACGGCATACGAGATCGAGAGTCGTGACTGGAGTTCAGACGTGTGCTCTTCCGATCNNNNNNTTTTTTTTTTTTTTTTTTV |
| Index 058 | CAGAAGACGGCATACGAGATCTCACGTAGTGACTGGAGTTCAGACGTGTGCTCTTCCGATCNNNNNNTTTTTTTTTTTTTTTTTTV |
| Index 059 | CAGAAGACGGCATACGAGATCTTATTACGTGACTGGAGTTCAGACGTGTGCTCTTCCGATCNNNNNNTTTTTTTTTTTTTTTTTTV |
| Index 060 | CAGAAGACGGCATACGAGATCGCTATTTGTGACTGGAGTTCAGACGTGTGCTCTTCCGATCNNNNNNTTTTTTTTTTTTTTTTTTV |
| Index 061 | CAGAAGACGGCATACGAGATGTTGTCCTGTGACTGGAGTTCAGACGTGTGCTCTTCCGATCNNNNNNTTTTTTTTTTTTTTTTTTV |
| Index 062 | CAGAAGACGGCATACGAGATTCAACTCTGTGACTGGAGTTCAGACGTGTGCTCTTCCGATCNNNNNNTTTTTTTTTTTTTTTTTTV |
| Index 063 | CAGAAGACGGCATACGAGATATACTTACGTGACTGGAGTTCAGACGTGTGCTCTTCCGATCNNNNNNTTTTTTTTTTTTTTTTTTV |
| Index 064 | CAGAAGACGGCATACGAGATTGTGTGGCGTGACTGGAGTTCAGACGTGTGCTCTTCCGATCNNNNNNTTTTTTTTTTTTTTTTTTV |
| Index 065 | CAGAAGACGGCATACGAGATCGACTCTTGTGACTGGAGTTCAGACGTGTGCTCTTCCGATCNNNNNNTTTTTTTTTTTTTTTTTTV |
| Index 066 | CAGAAGACGGCATACGAGATTCGCTCGCGTGACTGGAGTTCAGACGTGTGCTCTTCCGATCNNNNNNTTTTTTTTTTTTTTTTTTV |

| | |
|---|---|
| Index 067 | CAGAAGACGGCATACGAGATACACGAACGTGACTGGAGTTCAGACGTGTGCTCTTCCGATCNNNNNNTTTTTTTTTTTTTTTTTTV |
| Index 068 | CAGAAGACGGCATACGAGATTTAAATGCGTGACTGGAGTTCAGACGTGTGCTCTTCCGATCNNNNNNTTTTTTTTTTTTTTTTTTV |
| Index 069 | CAGAAGACGGCATACGAGATCCTCGGCGGTGACTGGAGTTCAGACGTGTGCTCTTCCGATCNNNNNNTTTTTTTTTTTTTTTTTTV |
| Index 070 | CAGAAGACGGCATACGAGATTATATTAGGTGACTGGAGTTCAGACGTGTGCTCTTCCGATCNNNNNNTTTTTTTTTTTTTTTTTTV |
| Index 071 | CAGAAGACGGCATACGAGATGCTGGCTTGTGACTGGAGTTCAGACGTGTGCTCTTCCGATCNNNNNNTTTTTTTTTTTTTTTTTTV |
| Index 072 | CAGAAGACGGCATACGAGATTGGATGTAGTGACTGGAGTTCAGACGTGTGCTCTTCCGATCNNNNNNTTTTTTTTTTTTTTTTTTV |
| Index 073 | CAGAAGACGGCATACGAGATTTTAGCACGTGACTGGAGTTCAGACGTGTGCTCTTCCGATCNNNNNNTTTTTTTTTTTTTTTTTTV |
| Index 074 | CAGAAGACGGCATACGAGATCAGTACGGGTGACTGGAGTTCAGACGTGTGCTCTTCCGATCNNNNNNTTTTTTTTTTTTTTTTTTV |
| Index 075 | CAGAAGACGGCATACGAGATATCCTCCAGTGACTGGAGTTCAGACGTGTGCTCTTCCGATCNNNNNNTTTTTTTTTTTTTTTTTTV |
| Index 076 | CAGAAGACGGCATACGAGATGTCACTACGTGACTGGAGTTCAGACGTGTGCTCTTCCGATCNNNNNNTTTTTTTTTTTTTTTTTTV |
| Index 077 | CAGAAGACGGCATACGAGATTGGTACTCGTGACTGGAGTTCAGACGTGTGCTCTTCCGATCNNNNNNTTTTTTTTTTTTTTTTTTV |
| Index 078 | CAGAAGACGGCATACGAGATCCGAGTTAGTGACTGGAGTTCAGACGTGTGCTCTTCCGATCNNNNNNTTTTTTTTTTTTTTTTTTV |
| Index 079 | CAGAAGACGGCATACGAGATGCTCTGCCGTGACTGGAGTTCAGACGTGTGCTCTTCCGATCNNNNNNTTTTTTTTTTTTTTTTTTV |
| Index 080 | CAGAAGACGGCATACGAGATGCCTCGATGTGACTGGAGTTCAGACGTGTGCTCTTCCGATCNNNNNNTTTTTTTTTTTTTTTTTTV |
| Index 081 | CAGAAGACGGCATACGAGATTATCTCTCGTGACTGGAGTTCAGACGTGTGCTCTTCCGATCNNNNNNTTTTTTTTTTTTTTTTTTV |
| Index 082 | CAGAAGACGGCATACGAGATGCTGGTAAGTGACTGGAGTTCAGACGTGTGCTCTTCCGATCNNNNNNTTTTTTTTTTTTTTTTTTV |
| Index 083 | CAGAAGACGGCATACGAGATCAGGTGCTGTGACTGGAGTTCAGACGTGTGCTCTTCCGATCNNNNNNTTTTTTTTTTTTTTTTTTV |
| Index 084 | CAGAAGACGGCATACGAGATGAACCAGCGTGACTGGAGTTCAGACGTGTGCTCTTCCGATCNNNNNNTTTTTTTTTTTTTTTTTTV |
| Index 085 | CAGAAGACGGCATACGAGATGGTGCATTGTGACTGGAGTTCAGACGTGTGCTCTTCCGATCNNNNNNTTTTTTTTTTTTTTTTTTV |
| Index 086 | CAGAAGACGGCATACGAGATCTCGCTCAGTGACTGGAGTTCAGACGTGTGCTCTTCCGATCNNNNNNTTTTTTTTTTTTTTTTTTV |
| Index 087 | CAGAAGACGGCATACGAGATTGGAAGAGGTGACTGGAGTTCAGACGTGTGCTCTTCCGATCNNNNNNTTTTTTTTTTTTTTTTTTV |
| Index 088 | CAGAAGACGGCATACGAGATAATCGGGCGTGACTGGAGTTCAGACGTGTGCTCTTCCGATCNNNNNNTTTTTTTTTTTTTTTTTTV |
| Index 089 | CAGAAGACGGCATACGAGATATGGTTTCGTGACTGGAGTTCAGACGTGTGCTCTTCCGATCNNNNNNTTTTTTTTTTTTTTTTTTV |
| Index 090 | CAGAAGACGGCATACGAGATGCAGACCAGTGACTGGAGTTCAGACGTGTGCTCTTCCGATCNNNNNNTTTTTTTTTTTTTTTTTTV |
| Index 091 | CAGAAGACGGCATACGAGATGCACACTTGTGACTGGAGTTCAGACGTGTGCTCTTCCGATCNNNNNNTTTTTTTTTTTTTTTTTTV |
| Index 092 | CAGAAGACGGCATACGAGATAAGAGTTCGTGACTGGAGTTCAGACGTGTGCTCTTCCGATCNNNNNNTTTTTTTTTTTTTTTTTTV |
| Index 093 | CAGAAGACGGCATACGAGATCATTTATGGTGACTGGAGTTCAGACGTGTGCTCTTCCGATCNNNNNNTTTTTTTTTTTTTTTTTTV |
| Index 094 | CAGAAGACGGCATACGAGATCTGATGAGGTGACTGGAGTTCAGACGTGTGCTCTTCCGATCNNNNNNTTTTTTTTTTTTTTTTTTV |
| Index 095 | CAGAAGACGGCATACGAGATATAGAGAGGTGACTGGAGTTCAGACGTGTGCTCTTCCGATCNNNNNNTTTTTTTTTTTTTTTTTTV |
| Index 096 | CAGAAGACGGCATACGAGATGGAGGTATGTGACTGGAGTTCAGACGTGTGCTCTTCCGATCNNNNNNTTTTTTTTTTTTTTTTTTV |

| | |
|---|---|
| Index 101 | CAGAAGACGGCATACGAGATATCGTTGTGTGACTGGAGTTCAGACGTGTGCTCTTCCGATCNNNNNNTTTTTTTTTTTTTTTTTTV |
| Index 102 | CAGAAGACGGCATACGAGATTATACACAGTGACTGGAGTTCAGACGTGTGCTCTTCCGATCNNNNNNTTTTTTTTTTTTTTTTTTV |
| Index 103 | CAGAAGACGGCATACGAGATCCAGGGCCGTGACTGGAGTTCAGACGTGTGCTCTTCCGATCNNNNNNTTTTTTTTTTTTTTTTTTV |
| Index 104 | CAGAAGACGGCATACGAGATAAAGGAGCGTGACTGGAGTTCAGACGTGTGCTCTTCCGATCNNNNNNTTTTTTTTTTTTTTTTTTV |
| Index 105 | CAGAAGACGGCATACGAGATAAACTCCTGTGACTGGAGTTCAGACGTGTGCTCTTCCGATCNNNNNNTTTTTTTTTTTTTTTTTTV |
| Index 106 | CAGAAGACGGCATACGAGATCCACCGGGGTGACTGGAGTTCAGACGTGTGCTCTTCCGATCNNNNNNTTTTTTTTTTTTTTTTTTV |
| Index 107 | CAGAAGACGGCATACGAGATGTATTAGAGTGACTGGAGTTCAGACGTGTGCTCTTCCGATCNNNNNNTTTTTTTTTTTTTTTTTTV |

| | |
|---|---|
| Index 108 | CAGAAGACGGCATACGAGATACAACCATGTGACTGGAGTTCAGACGTGTGCTCTTCCGATCNNNNNNTTTTTTTTTTTTTTTTTV |
| Index 109 | CAGAAGACGGCATACGAGATTGGGTTCCGTGACTGGAGTTCAGACGTGTGCTCTTCCGATCNNNNNNTTTTTTTTTTTTTTTTTV |
| Index 110 | CAGAAGACGGCATACGAGATCGGCATAAGTGACTGGAGTTCAGACGTGTGCTCTTCCGATCNNNNNNTTTTTTTTTTTTTTTTTV |
| Index 111 | CAGAAGACGGCATACGAGATCTCCTTTAGTGACTGGAGTTCAGACGTGTGCTCTTCCGATCNNNNNNTTTTTTTTTTTTTTTTTV |
| Index 112 | CAGAAGACGGCATACGAGATGATACTAAGTGACTGGAGTTCAGACGTGTGCTCTTCCGATCNNNNNNTTTTTTTTTTTTTTTTTV |
| Index 113 | CAGAAGACGGCATACGAGATCCTAATTCGTGACTGGAGTTCAGACGTGTGCTCTTCCGATCNNNNNNTTTTTTTTTTTTTTTTTV |
| Index 114 | CAGAAGACGGCATACGAGATTCACTACGGTGACTGGAGTTCAGACGTGTGCTCTTCCGATCNNNNNNTTTTTTTTTTTTTTTTTV |
| Index 115 | CAGAAGACGGCATACGAGATTAGCGTGCGTGACTGGAGTTCAGACGTGTGCTCTTCCGATCNNNNNNTTTTTTTTTTTTTTTTTV |
| Index 116 | CAGAAGACGGCATACGAGATCTCCAAGCGTGACTGGAGTTCAGACGTGTGCTCTTCCGATCNNNNNNTTTTTTTTTTTTTTTTTV |
| Index 117 | CAGAAGACGGCATACGAGATCTTCATATGTGACTGGAGTTCAGACGTGTGCTCTTCCGATCNNNNNNTTTTTTTTTTTTTTTTTV |
| Index 118 | CAGAAGACGGCATACGAGATTGGCGCCGGTGACTGGAGTTCAGACGTGTGCTCTTCCGATCNNNNNNTTTTTTTTTTTTTTTTTV |
| Index 119 | CAGAAGACGGCATACGAGATCCACATCTGTGACTGGAGTTCAGACGTGTGCTCTTCCGATCNNNNNNTTTTTTTTTTTTTTTTTV |
| Index 120 | CAGAAGACGGCATACGAGATCGAGACCTGTGACTGGAGTTCAGACGTGTGCTCTTCCGATCNNNNNNTTTTTTTTTTTTTTTTTV |
| Index 121 | CAGAAGACGGCATACGAGATGTGAGGCAGTGACTGGAGTTCAGACGTGTGCTCTTCCGATCNNNNNNTTTTTTTTTTTTTTTTTV |
| Index 122 | CAGAAGACGGCATACGAGATTCTCGTGTGTGACTGGAGTTCAGACGTGTGCTCTTCCGATCNNNNNNTTTTTTTTTTTTTTTTTV |
| Index 123 | CAGAAGACGGCATACGAGATTTTACGATGTGACTGGAGTTCAGACGTGTGCTCTTCCGATCNNNNNNTTTTTTTTTTTTTTTTTV |
| Index 124 | CAGAAGACGGCATACGAGATATAACGTCGTGACTGGAGTTCAGACGTGTGCTCTTCCGATCNNNNNNTTTTTTTTTTTTTTTTTV |
| Index 125 | CAGAAGACGGCATACGAGATACACGCTGGTGACTGGAGTTCAGACGTGTGCTCTTCCGATCNNNNNNTTTTTTTTTTTTTTTTTV |
| Index 126 | CAGAAGACGGCATACGAGATTTAAGACCGTGACTGGAGTTCAGACGTGTGCTCTTCCGATCNNNNNNTTTTTTTTTTTTTTTTTV |
| Index 127 | CAGAAGACGGCATACGAGATTTCCCATCGTGACTGGAGTTCAGACGTGTGCTCTTCCGATCNNNNNNTTTTTTTTTTTTTTTTTV |
| Index 128 | CAGAAGACGGCATACGAGATGTGACCCCGTGACTGGAGTTCAGACGTGTGCTCTTCCGATCNNNNNNTTTTTTTTTTTTTTTTTV |
| Index 129 | CAGAAGACGGCATACGAGATGACCGCGCGTGACTGGAGTTCAGACGTGTGCTCTTCCGATCNNNNNNTTTTTTTTTTTTTTTTTV |
| Index 130 | CAGAAGACGGCATACGAGATGTCGCAAAGTGACTGGAGTTCAGACGTGTGCTCTTCCGATCNNNNNNTTTTTTTTTTTTTTTTTV |
| Index 131 | CAGAAGACGGCATACGAGATGCGATCAAGTGACTGGAGTTCAGACGTGTGCTCTTCCGATCNNNNNNTTTTTTTTTTTTTTTTTV |
| Index 132 | CAGAAGACGGCATACGAGATTAGGCTAGGTGACTGGAGTTCAGACGTGTGCTCTTCCGATCNNNNNNTTTTTTTTTTTTTTTTTV |
| Index 133 | CAGAAGACGGCATACGAGATGTCATATAGTGACTGGAGTTCAGACGTGTGCTCTTCCGATCNNNNNNTTTTTTTTTTTTTTTTTV |
| Index 134 | CAGAAGACGGCATACGAGATAGTGTATCGTGACTGGAGTTCAGACGTGTGCTCTTCCGATCNNNNNNTTTTTTTTTTTTTTTTTV |
| Index 135 | CAGAAGACGGCATACGAGATGTCTTGTGGTGACTGGAGTTCAGACGTGTGCTCTTCCGATCNNNNNNTTTTTTTTTTTTTTTTTV |
| Index 136 | CAGAAGACGGCATACGAGATGACGTTATGTGACTGGAGTTCAGACGTGTGCTCTTCCGATCNNNNNNTTTTTTTTTTTTTTTTTV |
| Index 137 | CAGAAGACGGCATACGAGATCATGATCCGTGACTGGAGTTCAGACGTGTGCTCTTCCGATCNNNNNNTTTTTTTTTTTTTTTTTV |
| Index 138 | CAGAAGACGGCATACGAGATCAGCAAGTGTGACTGGAGTTCAGACGTGTGCTCTTCCGATCNNNNNNTTTTTTTTTTTTTTTTTV |
| Index 139 | CAGAAGACGGCATACGAGATGGCTTAATGTGACTGGAGTTCAGACGTGTGCTCTTCCGATCNNNNNNTTTTTTTTTTTTTTTTTV |
| Index 140 | CAGAAGACGGCATACGAGATAACGACGAGTGACTGGAGTTCAGACGTGTGCTCTTCCGATCNNNNNNTTTTTTTTTTTTTTTTTV |
| Index 141 | CAGAAGACGGCATACGAGATGCAAGCGGGTGACTGGAGTTCAGACGTGTGCTCTTCCGATCNNNNNNTTTTTTTTTTTTTTTTTV |
| Index 142 | CAGAAGACGGCATACGAGATAAGAGCGTGTGACTGGAGTTCAGACGTGTGCTCTTCCGATCNNNNNNTTTTTTTTTTTTTTTTTV |
| Index 143 | CAGAAGACGGCATACGAGATAGGCTCGAGTGACTGGAGTTCAGACGTGTGCTCTTCCGATCNNNNNNTTTTTTTTTTTTTTTTTV |
| Index 144 | CAGAAGACGGCATACGAGATGCAGTCGTGTGACTGGAGTTCAGACGTGTGCTCTTCCGATCNNNNNNTTTTTTTTTTTTTTTTTV |
| Index 145 | CAGAAGACGGCATACGAGATTTCACCTTGTGACTGGAGTTCAGACGTGTGCTCTTCCGATCNNNNNNTTTTTTTTTTTTTTTTTV |

| | |
|---|---|
| Index 146 | CAGAAGACGGCATACGAGATTCCTGTCCGTGACTGGAGTTCAGACGTGTGCTCTTCCGATCNNNNNNTTTTTTTTTTTTTTTTTTV |
| Index 147 | CAGAAGACGGCATACGAGATACCCCGTAGTGACTGGAGTTCAGACGTGTGCTCTTCCGATCNNNNNNTTTTTTTTTTTTTTTTTTV |
| Index 148 | CAGAAGACGGCATACGAGATAAAGAGTTGTGACTGGAGTTCAGACGTGTGCTCTTCCGATCNNNNNNTTTTTTTTTTTTTTTTTTV |
| Index 149 | CAGAAGACGGCATACGAGATGATGAAATGTGACTGGAGTTCAGACGTGTGCTCTTCCGATCNNNNNNTTTTTTTTTTTTTTTTTTV |
| Index 150 | CAGAAGACGGCATACGAGATGACCATAAGTGACTGGAGTTCAGACGTGTGCTCTTCCGATCNNNNNNTTTTTTTTTTTTTTTTTTV |
| Index 151 | CAGAAGACGGCATACGAGATAGCAAGTAGTGACTGGAGTTCAGACGTGTGCTCTTCCGATCNNNNNNTTTTTTTTTTTTTTTTTTV |
| Index 152 | CAGAAGACGGCATACGAGATAAACTGAGGTGACTGGAGTTCAGACGTGTGCTCTTCCGATCNNNNNNTTTTTTTTTTTTTTTTTTV |
| Index 153 | CAGAAGACGGCATACGAGATGAGCGATAGTGACTGGAGTTCAGACGTGTGCTCTTCCGATCNNNNNNTTTTTTTTTTTTTTTTTTV |
| Index 154 | CAGAAGACGGCATACGAGATGTTCTTCGGTGACTGGAGTTCAGACGTGTGCTCTTCCGATCNNNNNNTTTTTTTTTTTTTTTTTTV |
| Index 155 | CAGAAGACGGCATACGAGATCCTGAGACGTGACTGGAGTTCAGACGTGTGCTCTTCCGATCNNNNNNTTTTTTTTTTTTTTTTTTV |
| Index 156 | CAGAAGACGGCATACGAGATAGGTGAACGTGACTGGAGTTCAGACGTGTGCTCTTCCGATCNNNNNNTTTTTTTTTTTTTTTTTTV |
| Index 157 | CAGAAGACGGCATACGAGATCACATCTGGTGACTGGAGTTCAGACGTGTGCTCTTCCGATCNNNNNNTTTTTTTTTTTTTTTTTTV |
| Index 158 | CAGAAGACGGCATACGAGATTCTCAATCGTGACTGGAGTTCAGACGTGTGCTCTTCCGATCNNNNNNTTTTTTTTTTTTTTTTTTV |
| Index 159 | CAGAAGACGGCATACGAGATCACAACGTGTGACTGGAGTTCAGACGTGTGCTCTTCCGATCNNNNNNTTTTTTTTTTTTTTTTTTV |
| Index 160 | CAGAAGACGGCATACGAGATCACATACAGTGACTGGAGTTCAGACGTGTGCTCTTCCGATCNNNNNNTTTTTTTTTTTTTTTTTTV |
| Index 161 | CAGAAGACGGCATACGAGATAGTTATCCGTGACTGGAGTTCAGACGTGTGCTCTTCCGATCNNNNNNTTTTTTTTTTTTTTTTTTV |
| Index 162 | CAGAAGACGGCATACGAGATAAGCCTTTGTGACTGGAGTTCAGACGTGTGCTCTTCCGATCNNNNNNTTTTTTTTTTTTTTTTTTV |
| Index 163 | CAGAAGACGGCATACGAGATTAAAGCCGGTGACTGGAGTTCAGACGTGTGCTCTTCCGATCNNNNNNTTTTTTTTTTTTTTTTTTV |
| Index 164 | CAGAAGACGGCATACGAGATTCTAGACTGTGACTGGAGTTCAGACGTGTGCTCTTCCGATCNNNNNNTTTTTTTTTTTTTTTTTTV |
| Index 165 | CAGAAGACGGCATACGAGATAGTCCACAGTGACTGGAGTTCAGACGTGTGCTCTTCCGATCNNNNNNTTTTTTTTTTTTTTTTTTV |
| Index 166 | CAGAAGACGGCATACGAGATGCACGTCGGTGACTGGAGTTCAGACGTGTGCTCTTCCGATCNNNNNNTTTTTTTTTTTTTTTTTTV |
| Index 167 | CAGAAGACGGCATACGAGATGGCCTTGGGTGACTGGAGTTCAGACGTGTGCTCTTCCGATCNNNNNNTTTTTTTTTTTTTTTTTTV |
| Index 168 | CAGAAGACGGCATACGAGATTTGTCCGTGTGACTGGAGTTCAGACGTGTGCTCTTCCGATCNNNNNNTTTTTTTTTTTTTTTTTTV |
| Index 169 | CAGAAGACGGCATACGAGATGATGCACGGTGACTGGAGTTCAGACGTGTGCTCTTCCGATCNNNNNNTTTTTTTTTTTTTTTTTTV |
| Index 170 | CAGAAGACGGCATACGAGATGACGTGCCGTGACTGGAGTTCAGACGTGTGCTCTTCCGATCNNNNNNTTTTTTTTTTTTTTTTTTV |
| Index 171 | CAGAAGACGGCATACGAGATTTAGGTACGTGACTGGAGTTCAGACGTGTGCTCTTCCGATCNNNNNNTTTTTTTTTTTTTTTTTTV |
| Index 172 | CAGAAGACGGCATACGAGATACTCTATGGTGACTGGAGTTCAGACGTGTGCTCTTCCGATCNNNNNNTTTTTTTTTTTTTTTTTTV |
| Index 173 | CAGAAGACGGCATACGAGATGGATTTTCGTGACTGGAGTTCAGACGTGTGCTCTTCCGATCNNNNNNTTTTTTTTTTTTTTTTTTV |
| Index 174 | CAGAAGACGGCATACGAGATGGATAGACGTGACTGGAGTTCAGACGTGTGCTCTTCCGATCNNNNNNTTTTTTTTTTTTTTTTTTV |
| Index 175 | CAGAAGACGGCATACGAGATCTCTTAAAGTGACTGGAGTTCAGACGTGTGCTCTTCCGATCNNNNNNTTTTTTTTTTTTTTTTTTV |
| Index 176 | CAGAAGACGGCATACGAGATGCATAGCTGTGACTGGAGTTCAGACGTGTGCTCTTCCGATCNNNNNNTTTTTTTTTTTTTTTTTTV |
| Index 177 | CAGAAGACGGCATACGAGATTGCCCAAGGTGACTGGAGTTCAGACGTGTGCTCTTCCGATCNNNNNNTTTTTTTTTTTTTTTTTTV |
| Index 178 | CAGAAGACGGCATACGAGATCGTTACGAGTGACTGGAGTTCAGACGTGTGCTCTTCCGATCNNNNNNTTTTTTTTTTTTTTTTTTV |
| Index 179 | CAGAAGACGGCATACGAGATAAGGTGAAGTGACTGGAGTTCAGACGTGTGCTCTTCCGATCNNNNNNTTTTTTTTTTTTTTTTTTV |
| Index 180 | CAGAAGACGGCATACGAGATTTCTAACAGTGACTGGAGTTCAGACGTGTGCTCTTCCGATCNNNNNNTTTTTTTTTTTTTTTTTTV |
| Index 181 | CAGAAGACGGCATACGAGATGTGCAACCGTGACTGGAGTTCAGACGTGTGCTCTTCCGATCNNNNNNTTTTTTTTTTTTTTTTTTV |
| Index 182 | CAGAAGACGGCATACGAGATACTGCAGCGTGACTGGAGTTCAGACGTGTGCTCTTCCGATCNNNNNNTTTTTTTTTTTTTTTTTTV |
| Index 183 | CAGAAGACGGCATACGAGATGGTCGATCGTGACTGGAGTTCAGACGTGTGCTCTTCCGATCNNNNNNTTTTTTTTTTTTTTTTTTV |

| | |
|---|---|
| Index 184 | CAGAAGACGGCATACGAGATGTGTGATCGTGACTGGAGTTCAGACGTGTGCTCTTCCGATCNNNNNNTTTTTTTTTTTTTTTTTTV |
| Index 185 | CAGAAGACGGCATACGAGATCAGCCTCGGTGACTGGAGTTCAGACGTGTGCTCTTCCGATCNNNNNNTTTTTTTTTTTTTTTTTTV |
| Index 186 | CAGAAGACGGCATACGAGATTGACTAGAGTGACTGGAGTTCAGACGTGTGCTCTTCCGATCNNNNNNTTTTTTTTTTTTTTTTTTV |
| Index 187 | CAGAAGACGGCATACGAGATAACAGCACGTGACTGGAGTTCAGACGTGTGCTCTTCCGATCNNNNNNTTTTTTTTTTTTTTTTTTV |
| Index 188 | CAGAAGACGGCATACGAGATTGCGATAGGTGACTGGAGTTCAGACGTGTGCTCTTCCGATCNNNNNNTTTTTTTTTTTTTTTTTTV |
| Index 189 | CAGAAGACGGCATACGAGATCCAGATAAGTGACTGGAGTTCAGACGTGTGCTCTTCCGATCNNNNNNTTTTTTTTTTTTTTTTTTV |
| Index 190 | CAGAAGACGGCATACGAGATGGGATCTGGTGACTGGAGTTCAGACGTGTGCTCTTCCGATCNNNNNNTTTTTTTTTTTTTTTTTTV |
| Index 191 | CAGAAGACGGCATACGAGATTATTTCGGGTGACTGGAGTTCAGACGTGTGCTCTTCCGATCNNNNNNTTTTTTTTTTTTTTTTTTV |
| Index 192 | CAGAAGACGGCATACGAGATAGCCGTAGGTGACTGGAGTTCAGACGTGTGCTCTTCCGATCNNNNNNTTTTTTTTTTTTTTTTTTV |
| Index 193 | CAGAAGACGGCATACGAGATGTACCTTGGTGACTGGAGTTCAGACGTGTGCTCTTCCGATCNNNNNNTTTTTTTTTTTTTTTTTTV |
| Index 194 | CAGAAGACGGCATACGAGATCCAAGTGCGTGACTGGAGTTCAGACGTGTGCTCTTCCGATCNNNNNNTTTTTTTTTTTTTTTTTTV |
| Index 195 | CAGAAGACGGCATACGAGATTCTCCTTGGTGACTGGAGTTCAGACGTGTGCTCTTCCGATCNNNNNNTTTTTTTTTTTTTTTTTTV |
| Index 196 | CAGAAGACGGCATACGAGATAGCTTCAAGTGACTGGAGTTCAGACGTGTGCTCTTCCGATCNNNNNNTTTTTTTTTTTTTTTTTTV |

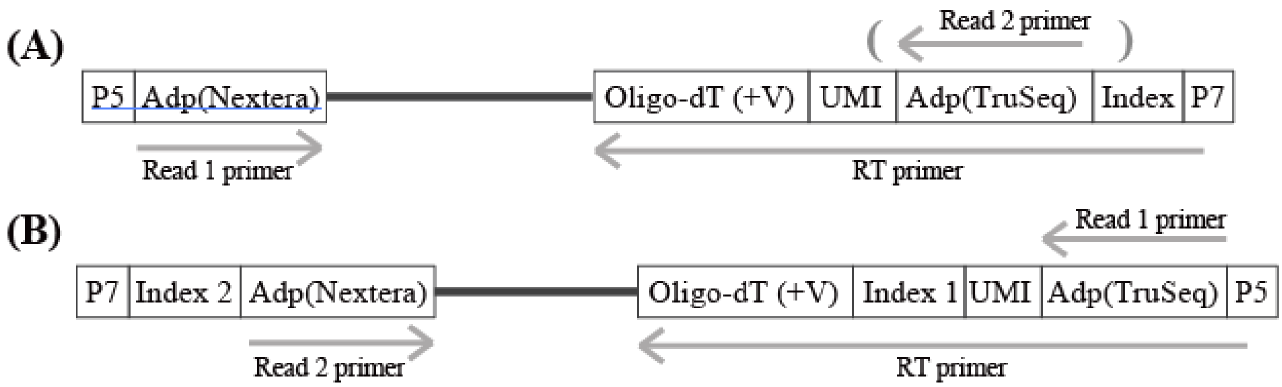


**(C)**

| Primer | Sequence(5'-3') |
|---|---|
| SR RT-primer (SR) | CAGAAGACGGCATACGAGATxxxxxxxx GTGACTGGAGTTCAGACGTGTGCTCTTCCGATCNNNNNNTTTTTTTTTTTTTTTTTTV |
| SR PCR forward-primer (SR) | CAAGCAGAAGACGGCATACGAGAT (for 5' part of every RT primers) |
| SR PCR reverse-primer (SR) | AATGATACGGCGACCACCGAGATCT(2nd index can be inserted here)ACACTCGTCGGCAGCGTC (for the Nextera tagmentation adapter sequence) |
| PE60 RT-primer (PE) | CCCTACACGACGCTCTTCCGATCTNNNNNNNxxxxxxxxxxTTTTTTTTTTTTTTTTTTVN |
| PE78 RT-primer (PE) | CCCTACACGACGCTCTTCCGATCTNNNNNNNNNNNNNxxxxxxxxxxTTTTTTTTTTTTTTTTTTTT TTTTTTTTTTVN |
| PE PCR forward-primer (PE) | AATGATACGGCGACCACCGAGATCTACACACACTCTTTCCCTACACGACGCTCTTCCGATCT (for 5' part of every RT primers) |
| PE PCR reverse-primer (PE) | Mixture of 8 primers with different index sequences. CAAGCAGAAGACGGCATACGAGAT********GTCTCGTGGGCTCGG (example ******** : ccgagggt, tcagctgt, cttgctgt, gtagacgc, agtgacac, ggtaattc, ttggccca, ctgtgttg) (for the Nextera tagmentation adapter sequence) |

**Supplementary figure 1　Primers used in the present study.**

**(A)** Schematic drawing of the library for the single-read sequencing (SR). P7 and P5 indicates sequences which are required for the hybridization to the flowcell (Illumina). Adp(TruSeq) and Adp(Nextera) means the adapter sequences used in the TruSeq and Nextera kit, respectively. **(B)** Schematic drawing of the library for the paired-end sequencing. In paired-end sequencing, quality of the latter part of the read 1 becomes low because of sequencing the polyA sequences, only index1 and UMI in read1 and read 2 sequences can be used for the analysis. **(C)** List of the sequences of the primers used in the present study. The "x" and "N" indicate index sequences and random bases (UMI). "V" meands bases of either "A", "C" or "G" (IDT). RT-indexing primer with 78mer in length was constructed depending on a previous study (PE78 RT-primer, Cao et al. 2017). Shorter RT primer (PE60 RT-primer) were designed to save cost of preparing primers. Diversity of index sequences were required for Illumina platforms (PE PCR reverse-primer).

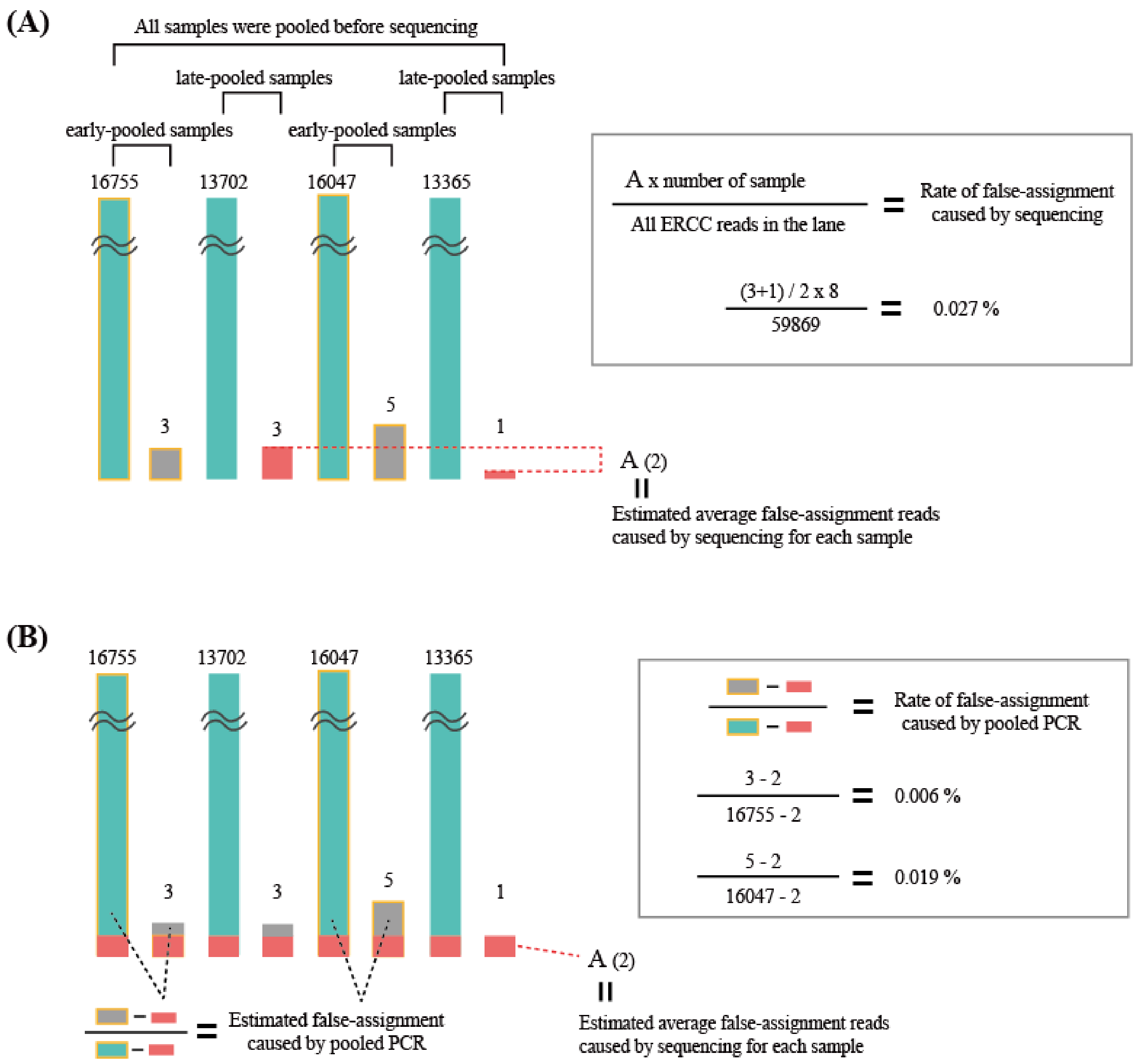


**Supplement figure 2　Method for calculation of false-assignment rates**

Method to calculate false-assignment rate caused by sequencing and pooled PCR used in the present study. (A) Method for calculation of false-assignment rate caused by sequencing. (B) Method for calculation of false-assignment rate caused by pooled PCR. Detaile of each sample was indicated in Fig 2 and Material and Method section.

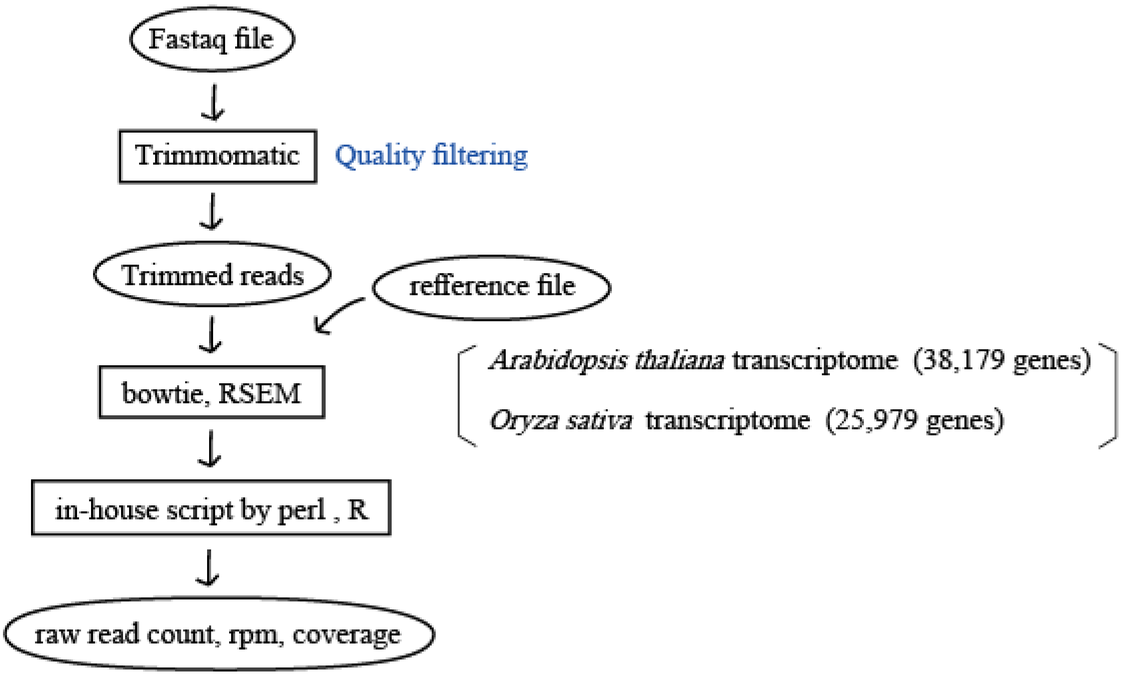


**Supplement figure 3 Overview of the analysis of RNA-Seq data.**
The FASTQ files obtained from RNA-Seq were filtered by trimomatic. Then the trimmed reads were mapped onto the plant transcriptome sequence and ERCC sequences. The produced bam files were analyzed by RSEM to calculate expected read count and in-house script in perl and R to calculate rpm, and coverage calculation. Detailed protocols are described in our previous study (Kamitani et al. 2016 FEMS Microbial Ecology, open access).

**Table S1 Information of the samples collected in this study (*n* = 45)**

| Sample ID | Index ID for RNA-Seq | Temperature of sampling day | Temperature of pre-1 day | Temperature of pre-2 day | Temperature of pre-3 day | Genotype | Day after sowing |
|---|---|---|---|---|---|---|---|
| 1 | 1 | 10 | 20 | 20 | 20 | Col-0 | 14 |
| 2 | 2 | 10 | 20 | 20 | 20 | Col-0 | 14 |
| 3 | 3 | 18 | 10 | 20 | 20 | Col-0 | 15 |
| 4 | 4 | 18 | 10 | 20 | 20 | Col-0 | 15 |
| 5 | 5 | 22 | 18 | 10 | 20 | Col-0 | 16 |
| 6 | 6 | 22 | 18 | 10 | 20 | Col-0 | 16 |
| 7 | 7 | 16 | 22 | 18 | 10 | Col-0 | 17 |
| 8 | 8 | 16 | 22 | 18 | 10 | Col-0 | 17 |
| 9 | 9 | 28 | 16 | 22 | 18 | Col-0 | 18 |
| 10 | 10 | 28 | 16 | 22 | 18 | Col-0 | 18 |
| 11 | 11 | 12 | 28 | 16 | 22 | Col-0 | 19 |
| 12 | 12 | 24 | 12 | 28 | 16 | Col-0 | 20 |
| 13 | 13 | 20 | 24 | 12 | 28 | Col-0 | 21 |
| 14 | 30 | 16 | 20 | 20 | 20 | Col-0 | 14 |
| 15 | 31 | 16 | 20 | 20 | 20 | Col-0 | 14 |
| 16 | 32 | 24 | 16 | 20 | 20 | Col-0 | 15 |
| 17 | 33 | 24 | 16 | 20 | 20 | Col-0 | 15 |
| 18 | 34 | 10 | 24 | 16 | 20 | Col-0 | 16 |
| 19 | 35 | 10 | 24 | 16 | 20 | Col-0 | 16 |
| 20 | 36 | 22 | 10 | 24 | 16 | Col-0 | 17 |
| 21 | 37 | 22 | 10 | 24 | 16 | Col-0 | 17 |
| 22 | 38 | 28 | 22 | 10 | 24 | Col-0 | 18 |
| 23 | 39 | 28 | 22 | 10 | 24 | Col-0 | 18 |
| 24 | 40 | 14 | 28 | 22 | 10 | Col-0 | 19 |
| 25 | 41 | 14 | 28 | 22 | 10 | Col-0 | 19 |
| 26 | 42 | 18 | 14 | 28 | 22 | Col-0 | 20 |
| 27 | 43 | 18 | 14 | 28 | 22 | Col-0 | 20 |
| 28 | 44 | 30 | 18 | 14 | 28 | Col-0 | 21 |
| 29 | 45 | 30 | 18 | 14 | 28 | Col-0 | 21 |
| 30 | 62 | 26 | 20 | 20 | 20 | Col-0 | 14 |
| 31 | 63 | 26 | 20 | 20 | 20 | Col-0 | 14 |
| 32 | 64 | 16 | 26 | 20 | 20 | Col-0 | 15 |
| 33 | 65 | 16 | 26 | 20 | 20 | Col-0 | 15 |
| 34 | 66 | 12 | 16 | 26 | 20 | Col-0 | 16 |
| 35 | 67 | 12 | 16 | 26 | 20 | Col-0 | 16 |
| 36 | 68 | 28 | 12 | 16 | 26 | Col-0 | 17 |
| 37 | 69 | 28 | 12 | 16 | 26 | Col-0 | 17 |
| 38 | 70 | 30 | 28 | 12 | 16 | Col-0 | 18 |
| 39 | 71 | 30 | 28 | 12 | 16 | Col-0 | 18 |
| 40 | 72 | 20 | 30 | 28 | 12 | Col-0 | 19 |
| 41 | 73 | 20 | 30 | 28 | 12 | Col-0 | 19 |
| 42 | 74 | 10 | 20 | 30 | 28 | Col-0 | 20 |
| 43 | 75 | 10 | 20 | 30 | 28 | Col-0 | 20 |
| 44 | 76 | 22 | 10 | 20 | 30 | Col-0 | 21 |
| 45 | 77 | 22 | 10 | 20 | 30 | Col-0 | 21 |

**Table S2 List of genes significantly correlated to temperature on each day (adjusted *p* -value <0.1)**

| Gene ID | Short description of gene | Abbreviation | Day | correlation coefficient |
|---|---|---|---|---|
| AT2G21390.1 | Coatomer, alpha subunit | - | Sampling day | 0.14 |
| AT2G09135.1 | | - | Sampling day | 0.20 |
| AT1G64560.1 | NULL | - | Sampling day | 0.25 |
| AT4G29180.3 | root hair specific 16 | RHS16 | Sampling day | 0.13 |
| AT4G23200.1 | cysteine-rich RLK (RECEPTOR-like protein kinase) 12 | CRK12 | Sampling day | 0.13 |
| AT1G05250.1 | Peroxidase superfamily protein | AtPRX1,PRX2 | Sampling day | 0.22 |
| AT2G23160.1 | F-box family protein | - | Sampling day | 0.19 |
| AT4G29940.1 | pathogenesis related homeodomain protein A | PRHA | Sampling day | 0.32 |
| AT4G27395.1 | pre-tRNA | - | Sampling day | 0.23 |
| AT2G31865.3 | poly(ADP-ribose) glycohydrolase 2 | PARG2 | Sampling day | 0.08 |
| AT3G07115.1 | pre-tRNA | - | Sampling day | 0.10 |
| AT2G21570.1 | pre-tRNA | - | Sampling day | 0.12 |
| AT1G07310.1 | Calcium-dependent lipid-binding (CaLB domain) family protein | - | Sampling day | 0.05 |
| AT5G20300.4 | Avirulence induced gene (AIG1) family protein | Toc90 | Sampling day | 0.10 |
| AT4G29580.2 | Cytidine/deoxycytidylate deaminase family protein | - | Sampling day | 0.12 |
| AT1G09735.2 | | - | Sampling day | 0.07 |
| AT1G12430.1 | armadillo repeat kinesin 3 | ARK3 | Sampling day | 0.08 |
| AT1G50880.1 | F-box and associated interaction domains-containing protein | - | Sampling day | 0.18 |
| AT1G04820.1 | tubulin alpha-4 chain | TUA4 | Sampling day | 0.17 |
| AT2G30750.1 | cytochrome P450, family 71, subfamily A, polypeptide 12 | CYP71A12 | Sampling day | 0.21 |
| AT2G37010.1 | non-intrinsic ABC protein 12 | NAP12 | Sampling day | 0.21 |
| AT1G36035.1 | transposable element gene | - | Sampling day | 0.07 |
| AT5G52900.1 | NULL | MAKR6 | Sampling day | 0.14 |
| AT1G33055.1 | NULL | - | Sampling day | 0.16 |
| AT2G40380.1 | prenylated RAB acceptor 1.B2 | PRA1.B2 | Sampling day | 0.18 |
| AT4G30060.1 | Core-2/I-branching beta-1,6-N-acetylglucosaminyltransferase family | - | Sampling day | 0.13 |
| AT1G06040.1 | B-box zinc finger family protein | STO | Sampling day | 0.13 |
| AT3G61280.2 | Arabidopsis thaliana protein of unknown function (DUF821) | - | Sampling day | 0.24 |
| AT3G29525.1 | transposable element gene | - | Sampling day | 0.14 |
| AT3G18960.1 | AP2/B3-like transcriptional factor family protein | - | Sampling day | 0.21 |
| AT4G14750.3 | IQ-domain 19 | IQD19 | Sampling day | 0.09 |
| ATCG01060.1 | iron-sulfur cluster binding;electron carriers;4 iron, 4 sulfur cluster | PSAC | Sampling day | 0.12 |
| AT4G16935.1 | transposable element gene | - | Sampling day | 0.35 |
| AT3G32310.1 | NULL | - | Sampling day | 0.07 |
| AT1G65770.1 | ascorbic acid mannose pathway regulator 1 | AMR1 | Sampling day | 0.15 |
| AT2G37570.1 | HSP20-like chaperones superfamily protein | SLT1 | Sampling day | 0.15 |
| AT1G56050.1 | GTP-binding protein-related | EngD-2 | Sampling day | 0.19 |
| AT2G22155.1 | NULL | - | Sampling day | 0.21 |
| AT1G21430.1 | Flavin-binding monooxygenase family protein | YUC11 | Sampling day | 0.33 |
| AT5G40660.1 | ATP12 protein-related | - | Sampling day | 0.22 |
| AT5G38800.1 | basic leucine-zipper 43 | bZIP43 | Sampling day | 0.18 |
| AT2G05195.1 | | - | Sampling day | 0.17 |
| AT2G20930.1 | SNARE-like superfamily protein | - | Sampling day | 0.33 |
| AT3G55980.2 | salt-inducible zinc finger 1 | SZF1 | Sampling day | 0.06 |
| AT1G46984.1 | F-box family protein | - | Sampling day | 0.23 |
| AT1G09157.1 | Protein of unknown function (DUF679) | - | Sampling day | 0.22 |
| AT1G63240.1 | NULL | - | Sampling day | 0.29 |
| AT5G61490.1 | Uncharacterised conserved protein (UCP012943) | - | Sampling day | 0.21 |
| AT2G12640.1 | transposable element gene | - | Sampling day | 0.05 |
| AT5G26130.1 | CAP (Cysteine-rich secretory proteins, Antigen 5, and Pathogenesis-related 1 protein) superfamily protein | - | Sampling day | 0.16 |
| AT4G34975.1 | pre-tRNA | - | Sampling day | 0.09 |
| AT1G06023.1 | | - | Sampling day | 0.11 |
| AT4G25390.1 | Protein kinase superfamily protein | - | Sampling day | 0.20 |
| AT2G38030.1 | pre-tRNA | - | Sampling day | 0.17 |
| AT3G52000.1 | serine carboxypeptidase-like 36 | scpl36 | Sampling day | 0.19 |
| AT3G32028.1 | transposable element gene | - | Sampling day | 0.19 |
| AT3G04525.1 | pre-tRNA | - | Sampling day | 0.37 |
| AT1G05800.1 | alpha/beta-Hydrolases superfamily protein | DGL | Sampling day | 0.19 |
| AT5G03345.2 | NULL | - | Sampling day | 0.10 |
| AT4G38050.1 | Xanthine/uracil permease family protein | - | Sampling day | 0.10 |
| AT5G50365.1 | | - | Sampling day | 0.37 |
| AT2G12580.1 | transposable element gene | - | Sampling day | 0.18 |
| AT5G18200.1 | UTP:galactose-1-phosphate uridylyltransferases;ribose-5-phosphate adenylyltransferases | - | Sampling day | 0.17 |
| AT5G01280.2 | NULL | BPP3 | Sampling day | 0.21 |
| AT5G42000.2 | ORMDL family protein | - | Sampling day | 0.13 |
| AT3G21481.1 | NULL | - | Sampling day | 0.14 |
| AT2G35384.1 | snoRNA | - | Sampling day | 0.10 |
| AT4G05595.1 | transposable element gene | - | Sampling day | 0.11 |
| AT5G57710.1 | Double Clp-N motif-containing P-loop nucleoside triphosphate hydrolases superfamily protein | SMAX1 | Sampling day | 0.08 |
| AT1G28050.1 | B-box type zinc finger protein with CCT domain | BBX13 | Sampling day | 0.13 |

| | | | | |
|---|---|---|---|---|
| AT1G07900.1 | LOB domain-containing protein 1 | LBD1 | Sampling day | 0.10 |
| AT2G04285.1 | | - | Sampling day | 0.18 |
| AT5G16910.1 | cellulose-synthase like D2 | CSLD2 | Sampling day | 0.23 |
| AT2G35160.3 | SU(VAR)3-9 homolog 5 | SUVH5 | Sampling day | 0.21 |
| AT1G05930.1 | Domain of unknown function (DUF313) | - | Sampling day | 0.20 |
| AT4G33430.2 | BRI1-associated receptor kinase | BAK1 | Sampling day | 0.17 |
| AT2G34780.1 | maternal effect embryo arrest 22 | MEE22 | Sampling day | 0.17 |
| AT1G60170.1 | pre-mRNA processing ribonucleoprotein binding region-containing | emb1220 | Sampling day | 0.30 |
| AT4G08868.1 | NULL | - | Sampling day | 0.19 |
| AT5G38560.1 | Protein kinase superfamily protein | PERK8 | Sampling day | 0.11 |
| AT5G11450.2 | Mog1/PsbP/DUF1795-like photosystem II reaction center PsbP family | PPD5 | Sampling day | 0.14 |
| AT5G26667.4 | P-loop containing nucleoside triphosphate hydrolases superfamily | PYR6 | Sampling day | 0.30 |
| AT4G01980.1 | transposable element gene | - | Sampling day | 0.19 |
| AT3G09100.2 | mRNA capping enzyme family protein | - | Sampling day | 0.07 |
| AT1G56330.1 | secretion-associated RAS 1B | SAR1B | Sampling day | 0.31 |
| AT5G07295.1 | | - | Sampling day | 0.29 |
| AT5G28970.1 | transposable element gene | - | Sampling day | 0.28 |
| AT4G16870.1 | transposable element gene | - | Sampling day | 0.20 |
| AT5G16990.1 | Zinc-binding dehydrogenase family protein | - | Sampling day | 0.15 |
| AT2G11420.1 | transposable element gene | - | Sampling day | 0.31 |
| AT1G67220.1 | histone acetyltransferase of the CBP family 2 | HAC2 | Sampling day | 0.05 |
| AT2G01900.1 | DNAse I-like superfamily protein | - | Sampling day | 0.13 |
| AT1G67330.1 | Protein of unknown function (DUF579) | - | Sampling day | 0.13 |
| AT3G03410.1 | EF hand calcium-binding protein family | - | Sampling day | 0.14 |
| AT1G43725.1 | transposable element gene | - | Sampling day | 0.18 |
| AT2G44750.2 | thiamin pyrophosphokinase 2 | TPK2 | Sampling day | 0.09 |
| AT5G57950.2 | 26S proteasome regulatory subunit, putative | - | Sampling day | 0.21 |
| AT2G12230.1 | NULL | - | Sampling day | 0.20 |
| AT4G07375.1 | | - | Sampling day | 0.05 |
| AT4G36120.1 | Plant protein of unknown function (DUF869) | - | Sampling day | 0.11 |
| AT5G61495.1 | NULL | - | Sampling day | 0.20 |
| AT2G00490.1 | | - | Sampling day | 0.23 |
| AT3G47650.1 | DnaJ/Hsp40 cysteine-rich domain superfamily protein | - | Sampling day | 0.10 |
| AT3G28970.1 | Domain of unknown function (DUF298) | AAR3 | Sampling day | 0.10 |
| AT4G06539.1 | transposable element gene | - | Sampling day | 0.16 |
| AT3G05345.1 | Chaperone DnaJ-domain superfamily protein | - | Sampling day | 0.19 |
| AT2G25110.1 | stromal cell-derived factor 2-like protein precursor | SDF2 | Sampling day | 0.19 |
| AT3G29460.1 | transposable element gene | - | Sampling day | 0.25 |
| AT1G35940.1 | transposable element gene | - | Sampling day | 0.18 |
| AT5G17420.1 | Cellulose synthase family protein | IRX3 | Sampling day | 0.07 |
| AT5G26910.4 | NULL | TRM8 | Sampling day | 0.22 |
| AT5G33438.1 | transposable element gene | - | Sampling day | 0.14 |
| AT3G07005.1 | low-molecular-weight cysteine-rich 43 | LCR43 | Sampling day | 0.26 |
| AT5G08725.1 | | - | Sampling day | 0.07 |
| AT3G25940.1 | TFIIB zinc-binding protein | - | Sampling day | 0.35 |
| AT3G27050.1 | NULL | - | Sampling day | 0.10 |
| AT2G40530.1 | NULL | - | Sampling day | 0.30 |
| AT4G22380.1 | Ribosomal protein L7Ae/L30e/S12e/Gadd45 family protein | - | Sampling day | 0.19 |
| AT5G42240.1 | serine carboxypeptidase-like 42 | scpl42 | Sampling day | 0.14 |
| AT5G40540.2 | Protein kinase superfamily protein | - | Sampling day | 0.11 |
| AT3G02750.1 | Protein phosphatase 2C family protein | - | Sampling day | 0.09 |
| AT2G45330.1 | RNA 2'-phosphotransferase, Tpt1 / KptA family | emb1067 | Sampling day | 0.12 |
| AT3G31996.1 | transposable element gene | - | Sampling day | 0.10 |
| AT5G38790.1 | NULL | - | Sampling day | 0.12 |
| AT3G13672.1 | TRAF-like superfamily protein | - | Sampling day | 0.22 |
| AT4G17250.1 | NULL | - | Sampling day | 0.32 |
| AT1G01720.1 | NAC (No Apical Meristem) domain transcriptional regulator superfamily protein | ATAF1 | Sampling day | 0.11 |
| AT1G64618.1 | other RNA | - | Sampling day | 0.22 |
| AT5G55040.1 | DNA-binding bromodomain-containing protein | - | Sampling day | 0.20 |
| AT2G25770.2 | Polyketide cyclase/dehydrase and lipid transport superfamily protein | - | Sampling day | 0.20 |
| AT1G62680.2 | Pentatricopeptide repeat (PPR) superfamily protein | - | Sampling day | 0.18 |
| AT4G07917.1 | transposable element gene | - | Sampling day | 0.17 |
| AT4G28860.1 | casein kinase I-like 4 | ckl4 | Sampling day | 0.12 |
| AT2G36830.1 | gamma tonoplast intrinsic protein | GAMMA-TIP | Sampling day | 0.15 |
| AT5G04110.3 | DNA GYRASE B3 | GYRB3 | Sampling day | 0.22 |
| AT3G47120.1 | RNA recognition motif (RRM)-containing protein | - | Sampling day | 0.13 |
| AT2G47000.2 | ATP binding cassette subfamily B4 | ABCB4 | Sampling day | 0.09 |
| AT5G16505.1 | NULL | MUG4 | Sampling day | 0.16 |
| AT2G17700.1 | ACT-like protein tyrosine kinase family protein | STY8 | Sampling day | 0.11 |
| AT2G10690.1 | transposable element gene | - | Sampling day | 0.06 |
| AT4G03960.1 | Phosphotyrosine protein phosphatases superfamily protein | PFA-DSP4 | Sampling day | 0.07 |
| AT3G00330.1 | | - | Sampling day | 0.09 |
| AT5G29571.1 | transposable element gene | - | Sampling day | 0.19 |
| AT1G20930.1 | cyclin-dependent kinase B2;2 | CDKB2%3B2 | Sampling day | 0.07 |
| AT4G03760.1 | transposable element gene | - | Sampling day | 0.14 |

| | | | | |
|---|---|---|---|---|
| AT5G46874.1 | Putative membrane lipoprotein | - | Sampling day | 0.23 |
| AT3G50700.1 | indeterminate(ID)-domain 2 | IDD2 | Sampling day | 0.20 |
| AT1G29330.1 | ER lumen protein retaining receptor family protein | ERD2 | Sampling day | 0.20 |
| AT2G41420.1 | proline-rich family protein | WIH2 | Sampling day | 0.17 |
| AT1G57010.1 | pre-tRNA | - | Sampling day | 0.13 |
| AT2G33730.1 | P-loop containing nucleoside triphosphate hydrolases superfamily | - | Sampling day | 0.06 |
| AT1G68765.1 | Putative membrane lipoprotein | IDA | Sampling day | 0.07 |
| AT1G72860.1 | Disease resistance protein (TIR-NBS-LRR class) family | - | Sampling day | 0.14 |
| AT3G31460.1 | transposable element gene | - | Sampling day | 0.13 |
| AT4G07205.1 | | - | Sampling day | 0.17 |
| AT5G53815.1 | transposable element gene | - | Sampling day | 0.06 |
| AT1G21510.1 | NULL | - | Sampling day | 0.13 |
| AT1G42718.1 | transposable element gene | - | Sampling day | 0.09 |
| AT3G08105.1 | | - | Sampling day | 0.36 |
| AT2G43320.1 | S-adenosyl-L-methionine-dependent methyltransferases superfamily | - | Sampling day | 0.05 |
| AT3G42254.1 | transposable element gene | - | Sampling day | 0.09 |
| AT1G09287.1 | | - | Sampling day | 0.21 |
| AT3G04895.1 | | - | Sampling day | 0.07 |
| AT4G17914.1 | NULL | - | Sampling day | 0.10 |
| AT5G27220.1 | Frigida-like protein | - | Sampling day | 0.22 |
| AT1G12130.1 | Flavin-binding monooxygenase family protein | - | Sampling day | 0.32 |
| AT1G04600.1 | myosin XI A | XIA | Sampling day | 0.08 |
| AT5G01895.1 | | - | Sampling day | 0.23 |
| AT1G07743.1 | | - | Sampling day | 0.12 |
| AT5G24430.1 | Calcium-dependent protein kinase (CDPK) family protein | - | Sampling day | 0.27 |
| AT4G39530.1 | Tetratricopeptide repeat (TPR)-like superfamily protein | - | Sampling day | 0.08 |
| AT2G05300.2 | NULL | - | Sampling day | 0.13 |
| AT2G46130.1 | WRKY DNA-binding protein 43 | WRKY43 | Sampling day | 0.07 |
| AT3G06280.1 | F-box associated ubiquitination effector family protein | - | Sampling day | 0.15 |
| AT2G14150.1 | transposable element gene | - | Sampling day | 0.37 |
| AT1G24180.1 | Thiamin diphosphate-binding fold (THDP-binding) superfamily protein | IAR4 | Sampling day | 0.26 |
| AT3G53130.1 | Cytochrome P450 superfamily protein | LUT1 | Sampling day | 0.16 |
| AT5G67420.2 | LOB domain-containing protein 37 | LBD37 | Sampling day | 0.31 |
| AT3G16890.1 | pentatricopeptide (PPR) domain protein 40 | PPR40 | Sampling day | 0.16 |
| AT5G34860.1 | transposable element gene | - | Sampling day | 0.13 |
| AT5G09635.1 | | - | Sampling day | 0.25 |
| AT5G32017.1 | pre-tRNA | - | Sampling day | 0.24 |
| AT1G58300.1 | heme oxygenase 4 | HO4 | Sampling day | 0.31 |
| AT5G48680.1 | Sterile alpha motif (SAM) domain-containing protein | - | Sampling day | 0.26 |
| AT4G18340.2 | Glycosyl hydrolase superfamily protein | - | Sampling day | 0.23 |
| AT5G28671.1 | transposable element gene | - | Sampling day | 0.07 |
| AT1G34822.1 | | - | Sampling day | 0.08 |
| AT4G24990.2 | Ubiquitin family protein | ATGP4 | Sampling day | 0.17 |
| AT2G21040.1 | Calcium-dependent lipid-binding (CaLB domain) family protein | - | Sampling day | 0.21 |
| AT4G17820.1 | transposable element gene | - | Sampling day | 0.20 |
| AT2G22370.1 | NULL | - | Sampling day | 0.10 |
| AT2G44140.1 | Peptidase family C54 protein | - | Sampling day | 0.16 |
| AT5G57420.1 | indole-3-acetic acid inducible 33 | IAA33 | Sampling day | 0.19 |
| AT1G33800.1 | Protein of unknown function (DUF579) | GXMT1 | Sampling day | 0.19 |
| AT5G58950.1 | Protein kinase superfamily protein | - | Sampling day | 0.15 |
| AT5G25625.1 | pre-tRNA | - | Sampling day | 0.09 |
| AT1G04650.1 | NULL | - | Sampling day | 0.24 |
| AT1G17250.1 | receptor like protein 3 | RLP3 | Sampling day | 0.11 |
| AT4G32490.1 | early nodulin-like protein 4 | ENODL4 | Sampling day | 0.14 |
| AT1G60760.1 | Plant invertase/pectin methylesterase inhibitor superfamily protein | - | Sampling day | 0.26 |
| AT5G47120.1 | BAX inhibitor 1 | BI1 | Sampling day | 0.19 |
| AT4G15440.1 | hydroperoxide lyase 1 | HPL1 | Sampling day | 0.12 |
| AT5G39155.1 | transposable element gene | - | Sampling day | 0.13 |
| AT4G06770.1 | | - | Sampling day | 0.07 |
| ATMG00920.1 | NULL | ORF215B | Sampling day | 0.05 |
| AT3G47550.2 | RING/FYVE/PHD zinc finger superfamily protein | - | Sampling day | 0.27 |
| AT3G43442.1 | transposable element gene | - | Sampling day | 0.15 |
| AT2G36792.2 | other RNA | - | Sampling day | 0.14 |
| AT1G30030.1 | transposable element gene | - | Sampling day | 0.22 |
| AT4G17720.1 | RNA-binding (RRM/RBD/RNP motifs) family protein | - | Sampling day | 0.21 |
| AT1G07277.1 | | - | Sampling day | 0.16 |
| AT1G54870.2 | NAD(P)-binding Rossmann-fold superfamily protein | - | Sampling day | 0.13 |
| AT1G51340.2 | MATE efflux family protein | - | Sampling day | 0.26 |
| AT5G08795.1 | | - | Sampling day | 0.20 |
| AT2G25312.1 | ECA1 gametogenesis related family protein | - | Sampling day | 0.13 |
| AT2G05910.1 | Protein of unknown function (DUF567) | - | Sampling day | 0.24 |
| AT4G24652.1 | NULL | - | Sampling day | 0.18 |
| AT1G27160.1 | valyl-tRNA synthetase / valine--tRNA ligase-related | - | Sampling day | 0.18 |
| AT3G19690.1 | CAP (Cysteine-rich secretory proteins, Antigen 5, and Pathogenesis-related 1 protein) superfamily protein | - | Sampling day | 0.07 |
| AT1G48560.1 | NULL | - | Sampling day | 0.23 |

| | | | | |
|---|---|---|---|---|
| AT4G14630.1 | germin-like protein 9 | GLP9 | Sampling day | 0.11 |
| AT2G09215.1 | | - | Sampling day | 0.13 |
| AT5G54130.2 | Calcium-binding endonuclease/exonuclease/phosphatase family | - | Sampling day | 0.07 |
| AT1G74960.2 | fatty acid biosynthesis 1 | FAB1 | Sampling day | 0.15 |
| AT4G00295.1 | | - | Sampling day | 0.18 |
| AT2G30615.1 | NULL | - | Sampling day | 0.17 |
| AT5G09465.1 | | - | Sampling day | 0.19 |
| AT2G09925.1 | | - | Sampling day | 0.15 |
| AT1G08870.1 | pre-tRNA | - | Sampling day | 0.07 |
| AT5G47700.1 | 60S acidic ribosomal protein family | - | Sampling day | 0.24 |
| AT1G25370.2 | Protein of unknown function (DUF1639) | - | Sampling day | 0.12 |
| AT4G06145.1 | | - | Sampling day | 0.43 |
| AT2G31380.1 | salt tolerance homologue | STH | Sampling day | 0.06 |
| AT4G06845.1 | | - | Sampling day | 0.26 |
| AT1G80830.1 | natural resistance-associated macrophage protein 1 | NRAMP1 | Sampling day | 0.18 |
| AT3G11500.2 | Small nuclear ribonucleoprotein family protein | - | Sampling day | 0.12 |
| AT4G04255.1 | transposable element gene | - | Sampling day | 0.14 |
| AT3G01145.1 | | - | Sampling day | 0.22 |
| AT1G54030.2 | GDSL-like Lipase/Acylhydrolase superfamily protein | MVP1 | Sampling day | 0.14 |
| AT1G09233.1 | | - | Sampling day | 0.26 |
| AT2G03900.1 | NULL | - | Sampling day | 0.15 |
| AT2G29485.1 | | - | Sampling day | 0.15 |
| AT2G21270.2 | ubiquitin fusion degradation 1 | UFD1 | Sampling day | 0.11 |
| AT1G04033.1 | | - | Sampling day | 0.22 |
| AT2G07760.1 | Zinc knuckle (CCHC-type) family protein | - | Sampling day | 0.16 |
| AT3G19380.1 | plant U-box 25 | PUB25 | Sampling day | 0.18 |
| AT5G56190.6 | Transducin/WD40 repeat-like superfamily protein | - | Sampling day | 0.08 |
| AT4G25900.1 | Galactose mutarotase-like superfamily protein | - | Sampling day | 0.18 |
| AT1G62780.1 | NULL | - | Sampling day | 0.10 |
| AT3G03665.1 | | - | Sampling day | 0.13 |
| AT1G09523.1 | | - | Sampling day | 0.31 |
| AT4G04595.1 | | - | Sampling day | 0.19 |
| AT1G13670.1 | NULL | - | Sampling day | 0.16 |
| AT2G06303.1 | transposable element gene | - | Sampling day | 0.06 |
| AT1G73655.1 | FKBP-like peptidyl-prolyl cis-trans isomerase family protein | - | Sampling day | 0.10 |
| AT4G26870.1 | Class II aminoacyl-tRNA and biotin synthetases superfamily protein | - | Sampling day | 0.09 |
| AT3G62760.1 | Glutathione S-transferase family protein | ATGSTF13 | Sampling day | 0.21 |
| AT4G27910.1 | SET domain protein 16 | SDG16 | Sampling day | 0.29 |
| AT1G69830.1 | alpha-amylase-like 3 | AMY3 | Sampling day | 0.07 |
| AT3G03330.1 | NAD(P)-binding Rossmann-fold superfamily protein | - | Sampling day | 0.12 |
| AT3G63215.1 | | - | Sampling day | 0.19 |
| AT3G60966.1 | RING/U-box superfamily protein | - | Sampling day | 0.13 |
| AT3G25620.1 | ABC-2 type transporter family protein | ABCG21 | Sampling day | 0.20 |
| AT1G54000.1 | GDSL-like Lipase/Acylhydrolase superfamily protein | GLL22 | Sampling day | 0.12 |
| AT5G65970.1 | Seven transmembrane MLO family protein | MLO10 | Sampling day | 0.20 |
| AT1G09693.1 | | - | Sampling day | 0.08 |
| AT3G16050.1 | pyridoxine biosynthesis 1.2 | PDX1.2 | Sampling day | 0.19 |
| AT5G62080.1 | Bifunctional inhibitor/lipid-transfer protein/seed storage 2S albumin superfamily protein | - | Sampling day | 0.23 |
| AT3G54860.1 | Sec1/munc18-like (SM) proteins superfamily | ATVPS33 | Sampling day | 0.08 |
| AT1G07460.2 | Concanavalin A-like lectin family protein | - | Sampling day | 0.18 |
| AT5G54930.1 | AT hook motif-containing protein | - | Sampling day | 0.13 |
| AT2G05010.1 | transposable element gene | - | Sampling day | 0.12 |
| AT2G17940.1 | Plant protein of unknown function (DUF827) | - | Sampling day | 0.18 |
| AT1G30540.2 | Actin-like ATPase superfamily protein | - | Sampling day | 0.23 |
| AT5G16220.1 | Octicosapeptide/Phox/Bem1p family protein | - | Sampling day | 0.21 |
| AT3G44769.1 | | - | Sampling day | 0.09 |
| AT4G10970.2 | NULL | - | Sampling day | 0.44 |
| AT5G24655.1 | response to low sulfur 4 | LSU4 | Sampling day | 0.23 |
| AT1G06063.1 | | - | Sampling day | 0.05 |
| AT1G29520.1 | AWPM-19-like family protein | - | Sampling day | 0.38 |
| AT5G09200.1 | | - | Sampling day | 0.17 |
| AT2G23600.2 | acetone-cyanohydrin lyase | ACL | Sampling day | 0.05 |
| AT5G56555.1 | | - | Sampling day | 0.21 |
| AT1G76070.1 | NULL | - | Sampling day | 0.21 |
| AT5G24860.2 | flowering promoting factor 1 | FPF1 | Sampling day | 0.08 |
| AT1G64070.1 | Disease resistance protein (TIR-NBS-LRR class) family | RLM1 | Sampling day | 0.13 |
| AT3G56530.1 | NAC domain containing protein 64 | NAC064 | Sampling day | 0.25 |
| AT2G43550.1 | Scorpion toxin-like knottin superfamily protein | - | Sampling day | 0.19 |
| AT3G08205.1 | | - | Sampling day | 0.10 |
| AT4G06519.1 | transposable element gene | - | Sampling day | 0.12 |
| AT3G13430.2 | RING/U-box superfamily protein | - | Sampling day | 0.14 |
| AT2G22090.2 | RNA-binding (RRM/RBD/RNP motifs) family protein | UBA1A | Sampling day | 0.15 |
| AT2G01860.1 | Tetratricopeptide repeat (TPR)-like superfamily protein | EMB975 | Sampling day | 0.21 |
| AT5G36340.1 | ECA1 gametogenesis related family protein | - | Sampling day | 0.12 |
| AT4G07420.1 | transposable element gene | - | Sampling day | 0.10 |

| | | | | |
|---|---|---|---|---|
| AT3G07690.1 | 6-phosphogluconate dehydrogenase family protein | - | Sampling day | 0.09 |
| AT1G68862.4 | NULL | - | Sampling day | 0.12 |
| AT5G44160.2 | C2H2-like zinc finger protein | NUC | Sampling day | 0.18 |
| AT2G01650.3 | plant UBX domain-containing protein 2 | PUX2 | Sampling day | 0.29 |
| AT5G47240.2 | nudix hydrolase homolog 8 | NUDT8 | Sampling day | 0.38 |
| AT5G56110.1 | myb domain protein 103 | MYB80 | Sampling day | 0.07 |
| AT5G39680.1 | Pentatricopeptide repeat (PPR) superfamily protein | EMB2744 | Sampling day | 0.33 |
| AT4G27270.4 | Quinone reductase family protein | - | Sampling day | 0.23 |
| AT1G57780.1 | heavy-metal-associated domain-containing protein | - | Sampling day | 0.08 |
| AT3G43425.1 | transposable element gene | - | Sampling day | 0.24 |
| AT5G67360.1 | Subtilase family protein | ARA12 | Sampling day | 0.14 |
| AT5G04130.2 | DNA GYRASE B2 | GYRB2 | Sampling day | 0.27 |
| AT5G57270.5 | Core-2/I-branching beta-1,6-N-acetylglucosaminyltransferase family | - | Sampling day | 0.38 |
| AT1G62690.2 | NULL | - | Sampling day | 0.08 |
| AT1G09460.1 | Carbohydrate-binding X8 domain superfamily protein | - | Sampling day | 0.09 |
| AT1G04930.2 | hydroxyproline-rich glycoprotein family protein | - | Sampling day | 0.14 |
| AT5G18270.1 | Arabidopsis NAC domain containing protein 87 | ANAC087 | Sampling day | 0.11 |
| AT3G10010.2 | demeter-like 2 | DML2 | Sampling day | 0.27 |
| AT2G38480.1 | Uncharacterised protein family (UPF0497) | - | Sampling day | 0.11 |
| AT5G19290.1 | alpha/beta-Hydrolases superfamily protein | - | Sampling day | 0.10 |
| AT4G34332.1 | other RNA | - | Sampling day | 0.08 |
| AT4G10540.1 | Subtilase family protein | - | Sampling day | 0.21 |
| AT5G67240.1 | small RNA degrading nuclease 3 | SDN3 | Sampling day | 0.13 |
| AT5G13320.3 | Auxin-responsive GH3 family protein | PBS3 | Sampling day | 0.18 |
| AT3G43586.1 | transposable element gene | - | Sampling day | 0.09 |
| AT3G02035.1 | | - | Sampling day | 0.24 |
| AT1G69580.1 | Homeodomain-like superfamily protein | - | Sampling day | 0.11 |
| AT1G66430.1 | pfkB-like carbohydrate kinase family protein | - | Sampling day | 0.11 |
| AT4G10230.1 | NULL | - | Sampling day | 0.22 |
| AT4G08602.1 | transposable element gene | - | Sampling day | 0.08 |
| AT3G53770.2 | late embryogenesis abundant 3 (LEA3) family protein | - | Sampling day | 0.21 |
| AT1G22050.1 | membrane-anchored ubiquitin-fold protein 6 precursor | MUB6 | Sampling day | 0.11 |
| AT3G29648.1 | transposable element gene | - | Sampling day | 0.07 |
| AT3G46720.1 | UDP-Glycosyltransferase superfamily protein | - | Sampling day | 0.09 |
| AT3G10360.1 | pumilio 4 | PUM4 | Sampling day | 0.06 |
| AT3G17640.1 | Leucine-rich repeat (LRR) family protein | - | Sampling day | 0.10 |
| AT2G07415.1 | | - | Sampling day | 0.09 |
| AT1G74456.1 | snoRNA | - | Sampling day | 0.09 |
| AT1G29270.1 | NULL | - | Sampling day | 0.21 |
| AT2G32800.1 | protein kinase family protein | AP4.3A | Sampling day | 0.15 |
| AT3G28240.1 | transposable element gene | - | Sampling day | 0.05 |
| AT1G41827.1 | transposable element gene | - | Sampling day | 0.18 |
| AT2G03470.1 | ELM2 domain-containing protein | - | Sampling day | 0.28 |
| AT5G03885.1 | | - | Sampling day | 0.29 |
| AT3G59420.1 | crinkly4 | CR4 | Sampling day | 0.24 |
| AT5G10530.1 | Concanavalin A-like lectin protein kinase family protein | - | Sampling day | 0.10 |
| AT5G39205.1 | transposable element gene | - | Sampling day | 0.15 |
| AT2G43130.1 | P-loop containing nucleoside triphosphate hydrolases superfamily | ARA4 | Sampling day | 0.10 |
| AT5G24670.3 | Cytidine/deoxycytidylate deaminase family protein | EMB2820 | Sampling day | 0.21 |
| AT5G27150.1 | Na+/H+ exchanger 1 | NHX1 | Sampling day | 0.19 |
| AT2G09610.1 | | - | Sampling day | 0.20 |
| AT1G53830.1 | pectin methylesterase 2 | PME2 | Sampling day | 0.24 |
| AT5G18610.2 | Protein kinase superfamily protein | - | Sampling day | 0.12 |
| AT1G67910.2 | NULL | - | Sampling day | 0.22 |
| AT2G30575.1 | los glycosyltransferase 5 | LGT5 | Sampling day | 0.15 |
| AT5G18560.1 | Integrase-type DNA-binding superfamily protein | PUCHI | Sampling day | 0.26 |
| AT1G07490.1 | ROTUNDIFOLIA like 3 | RTFL3 | Sampling day | 0.07 |
| AT1G02400.2 | gibberellin 2-oxidase 6 | GA2OX6 | Sampling day | 0.09 |
| AT5G65650.1 | Protein of unknown function (DUF1195) | - | Sampling day | 0.11 |
| AT3G50030.1 | ARM-repeat/Tetratricopeptide repeat (TPR)-like protein | - | Sampling day | 0.07 |
| AT5G01995.1 | | - | Sampling day | 0.23 |
| AT5G26970.1 | NULL | - | Sampling day | 0.17 |
| AT1G05863.1 | | - | Sampling day | 0.12 |
| AT3G03370.2 | NULL | - | Sampling day | 0.25 |
| AT1G49970.1 | CLP protease proteolytic subunit 1 | CLPR1 | Sampling day | 0.13 |
| AT5G48470.1 | NULL | - | Sampling day | 0.07 |
| AT3G16870.1 | GATA transcription factor 17 | GATA17 | Sampling day | 0.08 |
| AT5G63440.1 | Protein of unknown function (DUF167) | - | Sampling day | 0.16 |
| AT3G18320.1 | F-box and associated interaction domains-containing protein | - | Sampling day | 0.15 |
| AT1G40115.1 | transposable element gene | - | Sampling day | 0.27 |
| AT3G24927.1 | NULL | - | Sampling day | 0.13 |
| AT4G10820.1 | F-box family protein | - | Sampling day | 0.15 |
| AT4G06235.1 | | - | Sampling day | 0.17 |
| AT5G11210.6 | glutamate receptor 2.5 | GLR2.5 | Sampling day | 0.10 |
| AT4G06020.1 | | - | Sampling day | 0.30 |
| AT1G27630.3 | cyclin T 1;3 | CYCT1%3B3 | Sampling day | 0.08 |

| | | | | |
|---|---|---|---|---|
| AT1G30580.1 | GTP binding | - | Sampling day | 0.07 |
| AT5G46900.1 | Bifunctional inhibitor/lipid-transfer protein/seed storage 2S albumin superfamily protein | - | Sampling day | 0.10 |
| AT4G07324.1 | transposable element gene | - | Sampling day | 0.14 |
| AT5G20110.2 | Dynein light chain type 1 family protein | - | Sampling day | 0.17 |
| AT2G36050.1 | ovate family protein 15 | OFP15 | Sampling day | 0.15 |
| AT2G12305.1 | transposable element gene | - | Sampling day | 0.18 |
| AT2G28860.1 | cytochrome P450, family 710, subfamily A, polypeptide 4 | CYP710A4 | Sampling day | 0.31 |
| AT1G11595.1 | NULL | - | Sampling day | 0.19 |
| AT2G43870.1 | Pectin lyase-like superfamily protein | - | Sampling day | 0.14 |
| AT3G14640.1 | cytochrome P450, family 72, subfamily A, polypeptide 10 | CYP72A10 | Sampling day | 0.07 |
| AT5G35375.1 | NULL | - | Sampling day | 0.05 |
| AT5G09600.1 | succinate dehydrogenase 3-1 | SDH3-1 | Sampling day | 0.06 |
| AT4G19045.1 | Mob1/phocein family protein | - | Sampling day | 0.15 |
| AT4G07380.1 | NULL | - | Sampling day | 0.14 |
| AT1G35200.1 | NULL | - | Sampling day | 0.08 |
| AT2G08150.1 | | - | Sampling day | 0.25 |
| AT2G42395.1 | NULL | - | Sampling day | 0.13 |
| AT4G38830.1 | cysteine-rich RLK (RECEPTOR-like protein kinase) 26 | CRK26 | Sampling day | 0.31 |
| AT1G53570.6 | mitogen-activated protein kinase kinase kinase 3 | MAP3KA | Sampling day | 0.11 |
| AT3G07755.1 | | - | Sampling day | 0.21 |
| AT2G33490.1 | hydroxyproline-rich glycoprotein family protein | - | Sampling day | 0.07 |
| AT1G19500.1 | NULL | - | Sampling day | 0.16 |
| AT1G54690.1 | gamma histone variant H2AX | GAMMA-H2AX | Sampling day | 0.33 |
| AT5G44495.1 | ECA1 gametogenesis related family protein | - | Sampling day | 0.27 |
| AT4G11610.1 | C2 calcium/lipid-binding plant phosphoribosyltransferase family protein | - | Sampling day | 0.12 |
| AT4G07868.1 | NULL | - | Sampling day | 0.16 |
| AT3G28890.2 | receptor like protein 43 | RLP43 | Sampling day | 0.31 |
| AT2G05710.1 | aconitase 3 | ACO3 | Sampling day | 0.07 |
| AT2G38900.2 | Serine protease inhibitor, potato inhibitor I-type family protein | - | Sampling day | 0.10 |
| AT1G28300.1 | AP2/B3-like transcriptional factor family protein | LEC2 | Sampling day | 0.16 |
| AT1G55620.2 | chloride channel F | CLC-F | Sampling day | 0.12 |
| AT3G58700.1 | Ribosomal L5P family protein | - | Sampling day | 0.08 |
| AT1G72520.1 | PLAT/LH2 domain-containing lipoxygenase family protein | LOX4 | Sampling day | 0.07 |
| AT2G47140.1 | NAD(P)-binding Rossmann-fold superfamily protein | SDR5 | Sampling day | 0.23 |
| AT4G31700.1 | ribosomal protein S6 | RPS6 | Sampling day | 0.06 |
| AT3G09050.1 | NULL | - | Sampling day | 0.09 |
| AT2G40050.1 | Cysteine/Histidine-rich C1 domain family protein | - | Sampling day | 0.19 |
| AT3G54200.1 | Late embryogenesis abundant (LEA) hydroxyproline-rich glycoprotein | - | Sampling day | 0.31 |
| AT2G11320.1 | transposable element gene | - | Sampling day | 0.24 |
| AT1G27280.1 | Paired amphipathic helix (PAH2) superfamily protein | - | Sampling day | 0.21 |
| AT5G53470.1 | acyl-CoA binding protein 1 | ACBP1 | Sampling day | 0.20 |
| AT2G48030.2 | DNAse I-like superfamily protein | - | Sampling day | 0.19 |
| AT3G12915.1 | Ribosomal protein S5/Elongation factor G/III/V family protein | - | Sampling day | 0.08 |
| AT5G28570.1 | transposable element gene | - | Sampling day | 0.38 |
| AT4G14960.2 | Tubulin/FtsZ family protein | TUA6 | Sampling day | 0.15 |
| AT2G04125.1 | | - | Sampling day | 0.29 |
| AT2G24590.1 | RNA recognition motif and CCHC-type zinc finger domains containing | RSZ22a | Sampling day | 0.16 |
| AT1G09245.1 | Plant self-incompatibility protein S1 family | - | Sampling day | 0.18 |
| AT1G54035.1 | NULL | - | Sampling day | 0.05 |
| AT5G56870.1 | beta-galactosidase 4 | BGAL4 | Sampling day | 0.19 |
| AT1G24510.2 | TCP-1/cpn60 chaperonin family protein | - | Sampling day | 0.06 |
| AT3G43123.1 | transposable element gene | - | Sampling day | 0.06 |
| AT2G08560.1 | | - | Sampling day | 0.07 |
| AT1G76370.1 | Protein kinase superfamily protein | - | Sampling day | 0.14 |
| AT1G14610.1 | valyl-tRNA synthetase / valine--tRNA ligase (VALRS) | TWN2 | Sampling day | 0.14 |
| AT5G58600.2 | Plant protein of unknown function (DUF828) | PMR5 | Sampling day | 0.15 |
| AT5G48485.1 | Bifunctional inhibitor/lipid-transfer protein/seed storage 2S albumin superfamily protein | DIR1 | Sampling day | 0.26 |
| AT1G32810.2 | RING/FYVE/PHD zinc finger superfamily protein | - | Sampling day | 0.18 |
| AT5G42920.2 | THO complex, subunit 5 | THO5 | Sampling day | 0.12 |
| AT2G16320.1 | transposable element gene | - | Sampling day | 0.10 |
| AT1G24050.1 | RNA-processing, Lsm domain | - | Sampling day | 0.09 |
| AT2G07749.1 | Mitovirus RNA-dependent RNA polymerase | - | Sampling day | 0.51 |
| AT5G13890.1 | Family of unknown function (DUF716) | - | Sampling day | 0.17 |
| AT1G59730.1 | thioredoxin H-type 7 | TH7 | Sampling day | 0.23 |
| AT5G05025.1 | NULL | - | Sampling day | 0.16 |
| AT1G58290.1 | Glutamyl-tRNA reductase family protein | HEMA1 | Sampling day | 0.16 |
| AT1G09213.1 | | - | Sampling day | 0.34 |
| AT1G18570.1 | myb domain protein 51 | MYB51 | Sampling day | 0.09 |
| AT3G20400.1 | F-box associated ubiquitination effector family protein | EMB2743 | Sampling day | 0.12 |
| AT5G09340.1 | Ubiquitin family protein | - | Sampling day | 0.06 |
| AT1G11510.1 | DNA-binding storekeeper protein-related transcriptional regulator | - | Sampling day | 0.33 |
| AT5G03035.1 | | - | Sampling day | 0.05 |
| AT3G07620.1 | Exostosin family protein | - | Sampling day | 0.17 |
| AT4G15765.2 | FAD/NAD(P)-binding oxidoreductase family protein | - | Sampling day | 0.13 |

| | | | | |
|---|---|---|---|---|
| AT2G07965.1 | | - | Sampling day | 0.41 |
| AT5G45745.1 | pre-tRNA | - | Sampling day | 0.16 |
| AT1G56820.1 | pre-tRNA | - | Sampling day | 0.26 |
| AT3G46110.2 | Domain of unknown function (DUF966) | - | Sampling day | 0.17 |
| AT2G27120.2 | DNA polymerase epsilon catalytic subunit | TIL2 | Sampling day | 0.16 |
| AT1G32390.1 | transposable element gene | - | Sampling day | 0.26 |
| AT4G07285.1 | | - | Sampling day | 0.09 |
| AT3G12900.1 | 2-oxoglutarate (2OG) and Fe(II)-dependent oxygenase superfamily | - | Sampling day | 0.17 |
| AT1G16225.2 | Target SNARE coiled-coil domain protein | - | Sampling day | 0.10 |
| AT1G31170.1 | sulfiredoxin | SRX | Sampling day | 0.13 |
| AT1G32530.1 | RING/U-box superfamily protein | - | Sampling day | 0.16 |
| AT3G26000.1 | Ribonuclease inhibitor | - | Sampling day | 0.16 |
| AT2G18820.1 | transposable element gene | - | Sampling day | 0.07 |
| AT5G41530.1 | NULL | - | Sampling day | 0.05 |
| AT5G28330.1 | NULL | - | Sampling day | 0.08 |
| AT1G22940.1 | thiamin biosynthesis protein, putative | TH1 | Sampling day | 0.10 |
| AT1G65590.1 | beta-hexosaminidase 3 | HEXO3 | Sampling day | 0.25 |
| AT3G13750.1 | beta galactosidase 1 | BGAL1 | Sampling day | 0.13 |
| AT2G23410.1 | cis-prenyltransferase | CPT | Sampling day | 0.08 |
| AT1G46768.1 | related to AP2 1 | RAP2.1 | Sampling day | 0.09 |
| AT2G14900.1 | Gibberellin-regulated family protein | - | Sampling day | 0.07 |
| AT4G08155.1 | | - | Sampling day | 0.26 |
| AT1G28230.1 | purine permease 1 | PUP1 | Sampling day | 0.08 |
| AT3G30844.1 | transposable element gene | - | Sampling day | 0.22 |
| AT1G61540.1 | Galactose oxidase/kelch repeat superfamily protein | - | Sampling day | 0.18 |
| AT5G27050.1 | AGAMOUS-like 101 | AGL101 | Sampling day | 0.17 |
| AT1G23930.1 | transposable element gene | - | Sampling day | 0.22 |
| AT5G56160.2 | Sec14p-like phosphatidylinositol transfer family protein | - | Sampling day | 0.24 |
| AT4G24175.2 | NULL | - | Sampling day | 0.13 |
| AT2G32790.1 | Ubiquitin-conjugating enzyme family protein | - | Sampling day | 0.21 |
| AT1G75360.1 | NULL | - | Sampling day | 0.09 |
| AT2G07731.1 | NULL | - | Sampling day | 0.19 |
| AT4G13257.1 | transposable element gene | - | Sampling day | 0.20 |
| AT3G22961.1 | Paired amphipathic helix (PAH2) superfamily protein | - | Sampling day | 0.24 |
| AT2G10380.1 | transposable element gene | - | Sampling day | 0.19 |
| AT1G04780.1 | Ankyrin repeat family protein | - | Sampling day | 0.17 |
| AT4G01480.1 | pyrophosphorylase 5 | PPa5 | Sampling day | 0.19 |
| AT2G32590.1 | NULL | EMB2795 | Sampling day | 0.21 |
| AT5G35740.1 | Carbohydrate-binding X8 domain superfamily protein | - | Sampling day | 0.17 |
| AT1G53780.4 | peptidyl-prolyl cis-trans isomerases;hydrolases;nucleoside-triphosphatases;ATP binding;nucleotide binding;ATPases | - | Sampling day | 0.09 |
| AT2G07213.1 | other RNA | - | Sampling day | 0.07 |
| AT2G17180.1 | C2H2-like zinc finger protein | DAZ1 | Sampling day | 0.15 |
| AT1G07645.1 | dessication-induced 1VOC superfamily protein | DSI-1VOC | Sampling day | 0.20 |
| AT4G20070.1 | allantoate amidohydrolase | AAH | Sampling day | 0.10 |
| AT1G09433.1 | | - | Sampling day | 0.18 |
| AT3G51580.1 | NULL | - | Sampling day | 0.10 |
| AT4G11670.2 | Protein of unknown function (DUF810) | - | Sampling day | 0.19 |
| AT1G66210.1 | Subtilisin-like serine endopeptidase family protein | - | Sampling day | 0.19 |
| AT2G38770.1 | P-loop containing nucleoside triphosphate hydrolases superfamily | EMB2765 | Sampling day | 0.07 |
| AT1G35645.1 | transposable element gene | - | Sampling day | 0.12 |
| AT3G43415.1 | | - | Sampling day | 0.11 |
| AT5G36655.1 | transposable element gene | - | Sampling day | 0.15 |
| AT5G64813.1 | Ras-related small GTP-binding family protein | LIP1 | Sampling day | 0.22 |
| AT2G33110.2 | vesicle-associated membrane protein 723 | VAMP723 | Sampling day | 0.12 |
| AT1G13230.1 | Leucine-rich repeat (LRR) family protein | - | Sampling day | 0.12 |
| AT1G57450.1 | pre-tRNA | - | Sampling day | 0.10 |
| AT3G09455.1 | | - | Sampling day | 0.10 |
| AT4G03930.1 | Plant invertase/pectin methylesterase inhibitor superfamily | - | Sampling day | 0.12 |
| AT5G01960.1 | RING/U-box superfamily protein | - | Sampling day | 0.05 |
| AT5G62310.1 | AGC (cAMP-dependent, cGMP-dependent and protein kinase C) kinase family protein | IRE | Sampling day | 0.11 |
| AT4G31150.3 | endonuclease V family protein | - | Sampling day | 0.28 |
| AT5G32517.1 | transposable element gene | - | Sampling day | 0.14 |
| AT2G37220.1 | RNA-binding (RRM/RBD/RNP motifs) family protein | - | Sampling day | 0.25 |
| AT4G13262.1 | NULL | - | Sampling day | 0.19 |
| AT3G31317.1 | transposable element gene | - | Sampling day | 0.30 |
| AT5G38930.1 | RmlC-like cupins superfamily protein | - | Sampling day | 0.15 |
| AT5G08670.1 | ATP synthase alpha/beta family protein | - | Sampling day | 0.11 |
| AT3G06405.1 | | - | Sampling day | 0.20 |
| AT2G01910.1 | Microtubule associated protein (MAP65/ASE1) family protein | ATMAP65-6 | Sampling day | 0.17 |
| AT5G46330.1 | Leucine-rich receptor-like protein kinase family protein | FLS2 | Sampling day | 0.19 |
| AT5G17070.1 | NULL | - | Sampling day | 0.30 |
| AT2G20298.1 | NULL | - | Sampling day | 0.22 |
| AT1G62225.1 | NULL | - | Sampling day | 0.12 |
| AT1G43387.1 | transposable element gene | - | Sampling day | 0.20 |

| | | | | |
|---|---|---|---|---|
| AT3G60565.1 | transposable element gene | - | Sampling day | 0.13 |
| AT3G05650.1 | receptor like protein 32 | RLP32 | Sampling day | 0.33 |
| AT5G25630.3 | Tetratricopeptide repeat (TPR)-like superfamily protein | - | Sampling day | 0.19 |
| AT5G06900.1 | cytochrome P450, family 93, subfamily D, polypeptide 1 | CYP93D1 | Sampling day | 0.10 |
| AT4G29540.3 | bacterial transferase hexapeptide repeat-containing protein | LpxA | Sampling day | 0.07 |
| AT3G01060.1 | NULL | - | Sampling day | 0.13 |
| AT4G27900.1 | CCT motif family protein | - | Sampling day | 0.06 |
| AT2G13750.1 | transposable element gene | - | Sampling day | 0.19 |
| AT5G05995.1 | | - | Sampling day | 0.15 |
| AT1G28420.1 | homeobox-1 | HB-1 | Sampling day | 0.08 |
| AT1G79520.3 | Cation efflux family protein | - | Sampling day | 0.22 |
| AT2G10860.1 | transposable element gene | - | Sampling day | 0.18 |
| AT5G52360.1 | actin depolymerizing factor 10 | ADF10 | Sampling day | 0.22 |
| AT1G61490.4 | S-locus lectin protein kinase family protein | - | Sampling day | 0.18 |
| AT4G13080.1 | xyloglucan endotransglucosylase/hydrolase 1 | XTH1 | Sampling day | 0.22 |
| AT5G32420.2 | transposable element gene | - | Sampling day | 0.10 |
| AT4G04380.1 | transposable element gene | - | Sampling day | 0.30 |
| AT3G28760.2 | NULL | - | Sampling day | 0.28 |
| AT3G07080.1 | EamA-like transporter family | - | Sampling day | 0.08 |
| AT1G43745.1 | transposable element gene | - | Sampling day | 0.14 |
| AT5G41370.1 | homolog of xeroderma pigmentosum complementation group B 1 | XPB1 | Sampling day | 0.15 |
| AT5G30102.1 | transposable element gene | - | Sampling day | 0.17 |
| AT4G35690.1 | Arabidopsis protein of unknown function (DUF241) | - | Sampling day | 0.10 |
| AT3G06075.1 | | - | Sampling day | 0.07 |
| AT2G11380.1 | transposable element gene | - | Sampling day | 0.17 |
| AT3G62850.1 | zinc finger protein-related | - | Sampling day | 0.19 |
| AT3G03260.2 | homeodomain GLABROUS 8 | HDG8 | Sampling day | 0.17 |
| AT2G41990.1 | NULL | - | Sampling day | 0.15 |
| AT3G30388.1 | NULL | - | Sampling day | 0.10 |
| AT2G22300.3 | signal responsive 1 | SR1 | Sampling day | 0.15 |
| AT2G05565.1 | | - | Sampling day | 0.12 |
| AT1G06110.1 | SKP1/ASK-interacting protein 16 | SKIP16 | Sampling day | 0.31 |
| AT2G37120.1 | S1FA-like DNA-binding protein | - | Sampling day | 0.06 |
| AT1G14270.2 | CAAX amino terminal protease family protein | - | Sampling day | 0.05 |
| AT2G42160.1 | zinc finger (ubiquitin-hydrolase) domain-containing protein | BRIZ1 | Sampling day | 0.15 |
| AT5G44410.1 | FAD-binding Berberine family protein | - | Sampling day | 0.06 |
| AT4G20910.2 | double-stranded RNA binding protein-related / DsRBD protein-related | HEN1 | Sampling day | 0.15 |
| AT1G71950.1 | Proteinase inhibitor, propeptide | - | Sampling day | 0.28 |
| AT5G41340.2 | ubiquitin conjugating enzyme 4 | UBC4 | Sampling day | 0.08 |
| AT1G13090.1 | cytochrome P450, family 71, subfamily B, polypeptide 28 | CYP71B28 | Sampling day | 0.06 |
| AT3G05685.2 | Cystatin/monellin superfamily protein | - | Sampling day | 0.17 |
| AT3G43684.1 | transposable element gene | - | Sampling day | 0.06 |
| AT2G04690.3 | Pyridoxamine 5'-phosphate oxidase family protein | - | Sampling day | 0.09 |
| AT1G74650.2 | myb domain protein 31 | MYB31 | Sampling day | 0.18 |
| AT1G40101.1 | transposable element gene | - | Sampling day | 0.17 |
| AT5G46915.1 | transcriptional factor B3 family protein | - | Sampling day | 0.12 |
| AT4G29460.1 | Phospholipase A2 family protein | PLA2-GAMMA | Sampling day | 0.05 |
| AT1G23935.1 | NULL | - | Sampling day | 0.12 |
| ATMG09450.1 | | - | Sampling day | 0.06 |
| AT3G12502.1 | other RNA | - | Sampling day | 0.05 |
| AT3G07855.1 | | - | Sampling day | 0.13 |
| AT4G04780.1 | mediator 21 | MED21 | Sampling day | 0.44 |
| AT1G48355.1 | | - | Sampling day | 0.10 |
| AT5G16110.1 | NULL | - | Sampling day | 0.34 |
| AT1G10875.1 | | - | Sampling day | 0.08 |
| AT4G11910.1 | NULL | - | Sampling day | 0.08 |
| AT4G06537.1 | transposable element gene | - | Sampling day | 0.16 |
| AT5G17890.1 | DA1-related protein 4 | DAR4 | Sampling day | 0.21 |
| AT5G33253.1 | transposable element gene | - | Sampling day | 0.12 |
| AT5G08340.1 | Nucleotidylyl transferase superfamily protein | - | Sampling day | 0.09 |
| AT5G01555.1 | | - | Sampling day | 0.12 |
| AT2G03320.1 | NULL | - | Sampling day | 0.12 |
| AT2G23790.1 | Protein of unknown function (DUF607) | - | Sampling day | 0.16 |
| AT3G50330.1 | basic helix-loop-helix (bHLH) DNA-binding superfamily protein | HEC2 | Sampling day | 0.15 |
| AT5G41763.1 | | - | Sampling day | 0.06 |
| AT5G50443.1 | | - | Sampling day | 0.14 |
| AT3G06920.2 | Tetratricopeptide repeat (TPR)-like superfamily protein | - | Sampling day | 0.11 |
| AT1G63660.1 | GMP synthase (glutamine-hydrolyzing), putative / glutamine amidotransferase, putative | - | Sampling day | 0.14 |
| AT4G27570.1 | UDP-Glycosyltransferase superfamily protein | - | Sampling day | 0.15 |
| AT5G60820.1 | RING/U-box superfamily protein | - | Sampling day | 0.14 |
| AT4G31695.1 | pre-tRNA | - | Sampling day | 0.09 |
| AT5G39490.1 | F-box family protein | - | Sampling day | 0.10 |
| AT4G30380.2 | Barwin-related endoglucanase | - | Sampling day | 0.12 |
| AT4G05205.1 | | - | Sampling day | 0.05 |
| AT3G08780.2 | NULL | - | Sampling day | 0.12 |

| | | | | |
|---|---|---|---|---|
| AT3G17140.1 | Plant invertase/pectin methylesterase inhibitor superfamily protein | - | Sampling day | 0.16 |
| AT2G37390.1 | Chloroplast-targeted copper chaperone protein | NAKR2 | Sampling day | 0.19 |
| AT4G25310.1 | 2-oxoglutarate (2OG) and Fe(II)-dependent oxygenase superfamily | - | Sampling day | 0.10 |
| AT5G24790.1 | Protein of unknown function, DUF599 | - | Sampling day | 0.27 |
| AT5G22820.1 | ARM repeat superfamily protein | - | Sampling day | 0.28 |
| AT3G31390.1 | transposable element gene | - | Sampling day | 0.21 |
| AT1G34640.1 | peptidases | - | Sampling day | 0.12 |
| AT1G50310.1 | sugar transporter 9 | STP9 | Sampling day | 0.20 |
| AT3G02640.1 | NULL | - | Sampling day | 0.23 |
| AT2G08960.1 | | - | Sampling day | 0.16 |
| AT1G43180.1 | NULL | - | Sampling day | 0.22 |
| AT4G05360.1 | Zinc knuckle (CCHC-type) family protein | - | Sampling day | 0.22 |
| AT4G28365.1 | early nodulin-like protein 3 | ENODL3 | Sampling day | 0.30 |
| AT3G44718.1 | Plant thionin family protein | - | Sampling day | 0.08 |
| AT3G04355.1 | | - | Sampling day | 0.21 |
| AT3G48231.1 | low-molecular-weight cysteine-rich 48 | LCR48 | Sampling day | 0.16 |
| ATCG00590.1 | electron carriers | ORF31 | Sampling day | 0.05 |
| AT3G01410.1 | Polynucleotidyl transferase, ribonuclease H-like superfamily protein | - | Sampling day | 0.15 |
| AT3G11150.1 | 2-oxoglutarate (2OG) and Fe(II)-dependent oxygenase superfamily | - | Sampling day | 0.28 |
| AT3G03540.1 | non-specific phospholipase C5 | NPC5 | Sampling day | 0.33 |
| AT3G09940.1 | monodehydroascorbate reductase | MDHAR | Sampling day | 0.21 |
| AT5G46960.1 | Plant invertase/pectin methylesterase inhibitor superfamily protein | - | Sampling day | 0.21 |
| AT1G33300.1 | transposable element gene | - | Sampling day | 0.25 |
| AT4G33870.1 | Peroxidase superfamily protein | - | Sampling day | 0.15 |
| AT1G07433.1 | | - | Sampling day | 0.20 |
| AT5G36925.1 | NULL | - | Sampling day | 0.27 |
| AT2G32910.1 | DCD (Development and Cell Death) domain protein | - | Sampling day | 0.12 |
| AT5G50560.1 | Protein of unknown function (DUF 3339) | - | Sampling day | 0.14 |
| AT1G66960.2 | Terpenoid cyclases family protein | LUP5 | Sampling day | 0.20 |
| AT2G24700.1 | Transcriptional factor B3 family protein | - | Sampling day | 0.21 |
| AT3G01880.1 | Plant protein of unknown function (DUF946) | - | Sampling day | 0.19 |
| AT1G04037.1 | | - | Sampling day | 0.10 |
| AT3G15530.1 | S-adenosyl-L-methionine-dependent methyltransferases superfamily | - | Sampling day | 0.06 |
| AT3G10282.1 | NULL | - | Sampling day | 0.08 |
| AT1G07407.1 | | - | Sampling day | 0.18 |
| AT3G13210.1 | crooked neck protein, putative / cell cycle protein, putative | - | Sampling day | 0.13 |
| AT2G23130.2 | arabinogalactan protein 17 | AGP17 | Sampling day | 0.10 |
| AT3G62529.1 | NULL | - | Sampling day | 0.17 |
| AT1G08465.1 | Plant-specific transcription factor YABBY family protein | YAB2 | Sampling day | 0.37 |
| AT4G07650.1 | transposable element gene | - | Sampling day | 0.13 |
| AT2G17350.1 | NULL | - | Sampling day | 0.07 |
| AT5G08435.1 | | - | Sampling day | 0.20 |
| AT4G14430.1 | indole-3-butyric acid response 10 | IBR10 | Sampling day | 0.16 |
| AT1G50390.1 | pfkB-like carbohydrate kinase family protein | - | Sampling day | 0.20 |
| AT4G07120.1 | | U1-2 | Sampling day | 0.26 |
| AT3G21050.1 | transposable element gene | - | Sampling day | 0.14 |
| AT1G08133.1 | | - | Sampling day | 0.16 |
| AT3G43526.1 | transposable element gene | - | Sampling day | 0.18 |
| AT2G03915.1 | transposable element gene | - | Sampling day | 0.22 |
| AT3G01775.1 | | - | Sampling day | 0.08 |
| AT1G68907.1 | Defensin-like (DEFL) family protein | - | Sampling day | 0.17 |
| AT2G36040.1 | transposable element gene | - | Sampling day | 0.09 |
| AT2G07195.1 | | - | Sampling day | 0.20 |
| AT4G06395.1 | | - | Sampling day | 0.06 |
| AT5G61850.2 | floral meristem identity control protein LEAFY (LFY) | LFY | Sampling day | 0.05 |
| AT2G24900.1 | transposable element gene | - | Sampling day | 0.10 |
| AT4G06180.2 | | - | Sampling day | 0.11 |
| AT1G61550.1 | S-locus lectin protein kinase family protein | - | Sampling day | 0.26 |
| AT5G61880.3 | Protein Transporter, Pam16 | - | Sampling day | 0.13 |
| AT5G48010.2 | thalianol synthase 1 | THAS1 | Sampling day | 0.19 |
| AT3G28785.1 | | - | Sampling day | 0.26 |
| AT2G06650.1 | transposable element gene | - | Sampling day | 0.27 |
| AT5G26865.1 | | - | Sampling day | 0.11 |
| AT2G28940.1 | Protein kinase superfamily protein | - | Sampling day | 0.11 |
| AT3G31314.1 | transposable element gene | - | Sampling day | 0.19 |
| AT5G16870.1 | Peptidyl-tRNA hydrolase II (PTH2) family protein | - | Sampling day | 0.06 |
| AT4G27290.2 | S-locus lectin protein kinase family protein | - | Sampling day | 0.22 |
| AT4G14260.1 | Protein of unknown function (DUF295) | - | Sampling day | 0.18 |
| AT5G43010.1 | regulatory particle triple-A ATPase 4A | RPT4A | Sampling day | 0.14 |
| AT1G58265.1 | Cytochrome P450 superfamily protein | - | Sampling day | 0.23 |
| AT1G55964.1 | NULL | - | Sampling day | 0.17 |
| AT5G60930.2 | P-loop containing nucleoside triphosphate hydrolases superfamily | - | Sampling day | 0.09 |
| AT1G41893.1 | transposable element gene | - | Sampling day | 0.13 |
| AT4G30360.1 | cyclic nucleotide-gated channel 17 | CNGC17 | Sampling day | 0.20 |
| AT4G09200.1 | SPla/RYanodine receptor (SPRY) domain-containing protein | - | Sampling day | 0.23 |
| AT2G41250.1 | Haloacid dehalogenase-like hydrolase (HAD) superfamily protein | - | Sampling day | 0.13 |

| | | | | |
|---|---|---|---|---|
| AT4G08740.1 | NULL | - | Sampling day | 0.17 |
| AT2G05615.1 | | - | Sampling day | 0.10 |
| AT5G06100.4 | myb domain protein 33 | MYB33 | Sampling day | 0.26 |
| AT2G45180.1 | Bifunctional inhibitor/lipid-transfer protein/seed storage 2S albumin superfamily protein | - | Sampling day | 0.06 |
| AT5G29624.1 | Cysteine/Histidine-rich C1 domain family protein | - | Sampling day | 0.06 |
| AT5G11440.2 | CTC-interacting domain 5 | CID5 | Sampling day | 0.19 |
| AT1G16520.1 | NULL | - | Sampling day | 0.12 |
| AT1G51680.1 | 4-coumarate:CoA ligase 1 | 4CL1 | Sampling day | 0.16 |
| AT1G48710.1 | transposable element gene | - | Sampling day | 0.05 |
| AT1G55610.2 | BRI1 like | BRL1 | Sampling day | 0.11 |
| AT1G07400.1 | HSP20-like chaperones superfamily protein | - | Sampling day | 0.16 |
| AT1G57835.1 | NULL | - | Sampling day | 0.14 |
| AT2G05110.1 | transposable element gene | - | Sampling day | 0.33 |
| AT5G24215.1 | | - | Sampling day | 0.23 |
| AT3G44267.1 | transposable element gene | - | Sampling day | 0.06 |
| AT2G46494.1 | RING/U-box superfamily protein | - | Sampling day | 0.18 |
| AT3G50301.1 | NULL | - | Sampling day | 0.17 |
| AT5G34843.1 | transposable element gene | - | Sampling day | 0.19 |
| AT2G20060.1 | Ribosomal protein L4/L1 family | - | Sampling day | 0.08 |
| AT5G62930.3 | SGNH hydrolase-type esterase superfamily protein | - | Sampling day | 0.11 |
| AT1G11050.1 | Protein kinase superfamily protein | - | Sampling day | 0.37 |
| AT5G27495.1 | Defensin-like (DEFL) family protein | - | Sampling day | 0.08 |
| AT5G19960.1 | RNA-binding (RRM/RBD/RNP motifs) family protein | - | Sampling day | 0.10 |
| AT2G28671.1 | NULL | - | Sampling day | 0.17 |
| AT4G09105.1 | | - | Sampling day | 0.07 |
| AT5G09385.1 | | - | Sampling day | 0.10 |
| AT3G57220.1 | Glycosyl transferase family 4 protein | - | Sampling day | 0.07 |
| AT4G01930.1 | Cysteine/Histidine-rich C1 domain family protein | - | Sampling day | 0.10 |
| AT5G36907.1 | | - | Sampling day | 0.09 |
| AT4G27380.1 | NULL | - | Sampling day | 0.08 |
| AT1G16630.1 | NULL | - | Sampling day | 0.20 |
| AT3G60420.1 | Phosphoglycerate mutase family protein | - | Sampling day | 0.19 |
| AT1G19100.1 | Histidine kinase-, DNA gyrase B-, and HSP90-like ATPase family | DMS11 | Sampling day | 0.08 |
| AT1G20515.1 | other RNA | - | Sampling day | 0.17 |
| AT4G13400.2 | 2-oxoglutarate (2OG) and Fe(II)-dependent oxygenase superfamily | - | Sampling day | 0.10 |
| AT2G43920.4 | S-adenosyl-L-methionine-dependent methyltransferases superfamily | HOL2 | Sampling day | 0.25 |
| AT1G32010.1 | myosin heavy chain-related | - | Sampling day | 0.27 |
| AT5G10560.1 | Glycosyl hydrolase family protein | - | Sampling day | 0.10 |
| AT5G02855.1 | | - | Sampling day | 0.12 |
| AT4G38860.1 | SAUR-like auxin-responsive protein family | - | Sampling day | 0.11 |
| AT2G09435.1 | | - | Sampling day | 0.08 |
| AT1G09377.1 | | - | Sampling day | 0.19 |
| AT2G31690.1 | alpha/beta-Hydrolases superfamily protein | - | Sampling day | 0.08 |
| AT3G22337.1 | | - | Sampling day | 0.06 |
| AT2G07681.1 | Cytochrome C assembly protein | ABCI4 | Sampling day | 0.10 |
| AT3G04155.1 | | - | Sampling day | 0.30 |
| AT4G04295.1 | | - | Sampling day | 0.19 |
| AT3G21120.1 | F-box and associated interaction domains-containing protein | - | Sampling day | 0.14 |
| AT1G65480.2 | PEBP (phosphatidylethanolamine-binding protein) family protein | FT | Sampling day | 0.11 |
| AT3G27390.2 | NULL | - | Sampling day | 0.09 |
| AT3G07510.1 | NULL | - | Sampling day | 0.17 |
| ATMG00880.1 | NULL | ORF187 | Sampling day | 0.08 |
| AT3G32172.1 | transposable element gene | - | Sampling day | 0.10 |
| AT1G55500.4 | evolutionarily conserved C-terminal region 4 | ECT4 | Sampling day | 0.07 |
| AT5G01235.1 | | - | Sampling day | 0.09 |
| AT4G04170.1 | transposable element gene | - | Sampling day | 0.06 |
| AT5G08785.1 | | - | Sampling day | 0.36 |
| AT4G19360.1 | SCD6 protein-related | - | Sampling day | 0.10 |
| AT5G31087.1 | transposable element gene | - | Sampling day | 0.18 |
| AT1G05880.3 | RING/U-box superfamily protein | ARI12 | Sampling day | 0.16 |
| AT2G05950.1 | transposable element gene | - | Sampling day | 0.05 |
| AT2G38290.1 | ammonium transporter 2 | AMT2 | Sampling day | 0.12 |
| AT1G35240.1 | auxin response factor 20 | ARF20 | Sampling day | 0.12 |
| AT1G23250.2 | Caleosin-related family protein | - | Sampling day | 0.24 |
| ATCG00220.1 | photosystem II reaction center protein M | PSBM | Sampling day | 0.11 |
| AT2G12480.3 | serine carboxypeptidase-like 43 | SCPL43 | Sampling day | 0.07 |
| AT4G07862.1 | transposable element gene | - | Sampling day | 0.06 |
| AT5G08200.1 | peptidoglycan-binding LysM domain-containing protein | - | Sampling day | 0.17 |
| AT2G09990.1 | Ribosomal protein S5 domain 2-like superfamily protein | - | Sampling day | 0.10 |
| AT1G05837.1 | | - | Sampling day | 0.20 |
| AT3G19430.2 | late embryogenesis abundant protein-related / LEA protein-related | - | Sampling day | 0.05 |
| AT3G58420.1 | TRAF-like superfamily protein | - | Sampling day | 0.11 |
| AT1G03390.1 | HXXXD-type acyl-transferase family protein | - | Sampling day | 0.18 |
| AT5G14960.1 | DP-E2F-like 2 | DEL2 | Sampling day | 0.07 |
| AT1G27930.1 | Protein of unknown function (DUF579) | - | Sampling day | 0.11 |

| | | | | |
|---|---|---|---|---|
| AT1G55475.2 | NULL | - | Sampling day | 0.08 |
| AT2G47210.2 | myb-like transcription factor family protein | - | Sampling day | 0.24 |
| AT3G14460.1 | LRR and NB-ARC domains-containing disease resistance protein | - | Sampling day | 0.08 |
| AT5G05990.1 | Mitochondrial glycoprotein family protein | - | Sampling day | 0.07 |
| AT5G08470.2 | peroxisome 1 | PEX1 | Sampling day | 0.24 |
| AT3G59820.3 | LETM1-like protein | LETM1 | Sampling day | 0.05 |
| AT4G09425.1 | transposable element gene | - | Sampling day | 0.19 |
| AT3G49880.1 | glycosyl hydrolase family protein 43 | - | Sampling day | 0.10 |
| AT3G25970.1 | Pentatricopeptide repeat (PPR) superfamily protein | - | Sampling day | 0.13 |
| AT2G09335.1 | | - | Sampling day | 0.12 |
| AT1G58440.1 | FAD/NAD(P)-binding oxidoreductase family protein | XF1 | Sampling day | 0.11 |
| AT2G17540.2 | NULL | - | Sampling day | 0.23 |
| AT5G47080.3 | casein kinase II beta chain 1 | CKB1 | Sampling day | 0.14 |
| AT4G08116.1 | MIR401; miRNA | MIR401 | Sampling day | 0.20 |
| AT2G15980.1 | Tetratricopeptide repeat (TPR)-like superfamily protein | - | Sampling day | 0.19 |
| AT3G04985.1 | | - | Sampling day | 0.28 |
| AT1G80660.3 | H(+)-ATPase 9 | HA9 | Sampling day | 0.12 |
| AT1G48320.1 | Thioesterase superfamily protein | DHNAT1 | Sampling day | 0.12 |
| AT3G02625.1 | | - | Sampling day | 0.09 |
| AT3G45680.1 | Major facilitator superfamily protein | - | Sampling day | 0.12 |
| AT5G26720.1 | NULL | - | Sampling day | 0.09 |
| AT3G57180.1 | P-loop containing nucleoside triphosphate hydrolases superfamily | BPG2 | Sampling day | 0.09 |
| AT5G32800.1 | transposable element gene | - | Sampling day | 0.22 |
| AT5G05435.1 | other RNA | - | Sampling day | 0.35 |
| AT1G69325.1 | Remorin family protein | - | Sampling day | 0.15 |
| AT1G04393.1 | | - | Sampling day | 0.36 |
| AT1G19530.1 | NULL | - | Sampling day | 0.23 |
| AT2G07215.1 | NULL | - | Sampling day | 0.13 |
| AT2G44230.1 | Plant protein of unknown function (DUF946) | - | Sampling day | 0.15 |
| AT5G27440.1 | NULL | - | Sampling day | 0.13 |
| AT1G42170.1 | transposable element gene | - | Sampling day | 0.22 |
| AT4G33770.3 | Inositol 1,3,4-trisphosphate 5/6-kinase family protein | - | Sampling day | 0.06 |
| AT3G11710.1 | lysyl-tRNA synthetase 1 | ATKRS-1 | Sampling day | 0.14 |
| AT2G17295.1 | snoRNA | - | Sampling day | 0.16 |
| AT4G28210.1 | embryo defective 1923 | EMB1923 | Sampling day | 0.17 |
| AT3G08465.1 | | - | Sampling day | 0.08 |
| AT3G63430.2 | NULL | TRM5 | Sampling day | 0.09 |
| AT4G05587.1 | transposable element gene | - | Sampling day | 0.10 |
| AT5G48810.1 | cytochrome B5 isoform D | CB5-D | Sampling day | 0.21 |
| AT2G08895.1 | | - | Sampling day | 0.27 |
| AT4G02480.1 | AAA-type ATPase family protein | - | Sampling day | 0.06 |
| AT3G30833.1 | transposable element gene | - | Sampling day | 0.20 |
| AT5G25870.1 | NULL | - | Sampling day | 0.19 |
| AT1G24140.1 | Matrixin family protein | - | Sampling day | 0.07 |
| AT1G80370.1 | Cyclin A2;4 | CYCA2%3B4 | Sampling day | 0.08 |
| AT2G21780.1 | NULL | - | Sampling day | 0.22 |
| AT4G23110.1 | insulin-like growth factor binding | - | Sampling day | 0.23 |
| AT4G06510.1 | transposable element gene | - | Sampling day | 0.14 |
| AT5G59350.1 | NULL | - | Sampling day | 0.21 |
| AT2G28340.1 | GATA transcription factor 13 | GATA13 | Sampling day | 0.18 |
| AT5G43035.1 | transposable element gene | - | Sampling day | 0.23 |
| AT3G55940.1 | Phosphoinositide-specific phospholipase C family protein | - | Sampling day | 0.52 |
| AT1G06890.1 | nodulin MtN21 /EamA-like transporter family protein | - | Sampling day | 0.18 |
| AT5G55200.1 | Co-chaperone GrpE family protein | MGE1 | Sampling day | 0.14 |
| AT1G38131.1 | O-fucosyltransferase family protein | - | Sampling day | 0.05 |
| AT3G06650.2 | ATP-citrate lyase B-1 | ACLB-1 | Sampling day | 0.07 |
| AT3G06780.1 | glycine-rich protein | - | Sampling day | 0.27 |
| AT4G04105.1 | transposable element gene | - | Sampling day | 0.13 |
| AT2G04395.3 | Nucleic acid-binding, OB-fold-like protein | - | Sampling day | 0.22 |
| AT1G07853.1 | | - | Sampling day | 0.11 |
| AT1G62330.1 | O-fucosyltransferase family protein | - | Sampling day | 0.17 |
| AT4G22305.2 | alpha/beta-Hydrolases superfamily protein | - | Sampling day | 0.29 |
| AT3G30415.1 | NULL | - | Sampling day | 0.17 |
| AT3G22136.1 | transposable element gene | - | Sampling day | 0.19 |
| AT3G42803.1 | transposable element gene | - | Sampling day | 0.27 |
| AT1G19485.2 | Transducin/WD40 repeat-like superfamily protein | - | Sampling day | 0.26 |
| AT3G08725.1 | | - | Sampling day | 0.07 |
| AT3G59640.2 | glycine-rich protein | - | Sampling day | 0.09 |
| AT1G54170.1 | CTC-interacting domain 3 | CID3 | Sampling day | 0.15 |
| AT4G02190.1 | Cysteine/Histidine-rich C1 domain family protein | - | Sampling day | 0.13 |
| AT3G11505.1 | pre-tRNA | - | Sampling day | 0.17 |
| AT4G24615.1 | | - | Sampling day | 0.18 |
| AT1G08737.1 | | U5-2 | Sampling day | 0.07 |
| AT1G80550.1 | Pentatricopeptide repeat (PPR) superfamily protein | - | Sampling day | 0.21 |
| AT2G06290.1 | transposable element gene | - | Sampling day | 0.20 |
| AT3G10113.1 | Homeodomain-like superfamily protein | - | Sampling day | 0.28 |

| | | | | |
|---|---|---|---|---|
| AT4G29090.1 | Ribonuclease H-like superfamily protein | - | Sampling day | 0.16 |
| AT1G76600.1 | NULL | - | Sampling day | 0.16 |
| AT2G18160.1 | basic leucine-zipper 2 | bZIP2 | Sampling day | 0.10 |
| AT5G03775.1 | pre-tRNA | - | Sampling day | 0.14 |
| AT3G56150.1 | eukaryotic translation initiation factor 3C | EIF3C | Sampling day | 0.09 |
| AT5G07415.1 | | U1-13p | Sampling day | 0.14 |
| AT1G01725.1 | NULL | - | Sampling day | 0.25 |
| AT1G14770.2 | RING/FYVE/PHD zinc finger superfamily protein | - | Sampling day | 0.07 |
| AT4G20090.1 | Pentatricopeptide repeat (PPR) superfamily protein | EMB1025 | Sampling day | 0.08 |
| AT1G68640.1 | bZIP transcription factor family protein | PAN | Sampling day | 0.06 |
| AT3G17220.1 | pectin methylesterase inhibitor 2 | PMEI2 | Sampling day | 0.20 |
| AT5G02980.1 | Galactose oxidase/kelch repeat superfamily protein | - | Sampling day | 0.23 |
| AT2G42090.1 | actin 9 | ACT9 | Sampling day | 0.10 |
| AT3G55200.1 | Cleavage and polyadenylation specificity factor (CPSF) A subunit | SAP130a | Sampling day | 0.47 |
| AT4G24972.1 | tapetum determinant 1 | TPD1 | Sampling day | 0.19 |
| AT1G26210.1 | SOB five-like 1 | SOFL1 | Sampling day | 0.33 |
| AT3G12560.2 | TRF-like 9 | TRFL9 | Sampling day | 0.11 |
| AT1G54200.1 | NULL | - | Sampling day | 0.11 |
| AT4G00520.4 | Acyl-CoA thioesterase family protein | - | Sampling day | 0.06 |
| AT3G32275.1 | transposable element gene | - | Sampling day | 0.10 |
| AT2G10000.1 | transposable element gene | - | Sampling day | 0.19 |
| AT3G05655.1 | | - | Sampling day | 0.09 |
| AT4G07664.1 | transposable element gene | - | Sampling day | 0.11 |
| AT2G02100.1 | low-molecular-weight cysteine-rich 69 | LCR69 | Sampling day | 0.19 |
| AT4G28680.6 | L-tyrosine decarboxylase | TYRDC | Sampling day | 0.12 |
| AT3G22690.2 | NULL | - | Sampling day | 0.10 |
| AT1G05530.1 | UDP-glucosyl transferase 75B2 | UGT75B2 | Sampling day | 0.19 |
| AT1G18630.1 | glycine-rich RNA-binding protein 6 | GR-RBP6 | Sampling day | 0.38 |
| AT2G05550.1 | transposable element gene | - | Sampling day | 0.16 |
| AT4G28570.1 | Long-chain fatty alcohol dehydrogenase family protein | - | Sampling day | 0.20 |
| AT2G31080.1 | transposable element gene | - | Sampling day | 0.22 |
| AT2G40160.1 | Plant protein of unknown function (DUF828) | TBL30 | Sampling day | 0.06 |
| AT5G06610.1 | Protein of unknown function (DUF620) | - | Sampling day | 0.23 |
| AT2G22470.1 | arabinogalactan protein 2 | AGP2 | Sampling day | 0.10 |
| AT4G07738.1 | transposable element gene | - | Sampling day | 0.27 |
| AT1G54280.3 | ATPase E1-E2 type family protein / haloacid dehalogenase-like hydrolase family protein | - | Sampling day | 0.17 |
| AT3G57400.1 | NULL | - | Sampling day | 0.10 |
| AT1G71110.1 | NULL | - | Sampling day | 0.10 |
| AT3G52090.2 | DNA-directed RNA polymerase, RBP11-like | NRPB11 | Sampling day | 0.16 |
| AT2G42750.1 | DNAJ heat shock N-terminal domain-containing protein | - | Sampling day | 0.24 |
| AT4G32320.1 | ascorbate peroxidase 6 | APX6 | Sampling day | 0.09 |
| AT3G23720.1 | transposable element gene | - | Sampling day | 0.20 |
| AT1G57290.1 | pre-tRNA | - | Sampling day | 0.19 |
| AT4G18940.1 | RNA ligase/cyclic nucleotide phosphodiesterase family protein | - | Sampling day | 0.17 |
| AT1G71440.1 | tubulin folding cofactor E / Pfifferling (PFI) | PFI | Sampling day | 0.11 |
| AT4G06676.2 | NULL | - | Sampling day | 0.13 |
| AT4G20430.3 | Subtilase family protein | - | Sampling day | 0.25 |
| AT5G35600.1 | histone deacetylase7 | HDA7 | Sampling day | 0.17 |
| AT4G03255.1 | | - | Sampling day | 0.15 |
| AT3G27540.1 | beta-1,4-N-acetylglucosaminyltransferase family protein | - | Sampling day | 0.24 |
| AT5G35923.1 | transposable element gene | - | Sampling day | 0.12 |
| AT5G59380.1 | methyl-CPG-binding domain 6 | MBD6 | Sampling day | 0.12 |
| AT3G13870.1 | Root hair defective 3 GTP-binding protein (RHD3) | RHD3 | Sampling day | 0.15 |
| AT2G07799.1 | NULL | - | Sampling day | 0.10 |
| AT5G16640.1 | Pentatricopeptide repeat (PPR) superfamily protein | - | Sampling day | 0.22 |
| AT4G33560.1 | Wound-responsive family protein | - | Sampling day | 0.13 |
| AT3G52360.1 | NULL | - | Sampling day | 0.12 |
| AT3G17010.1 | AP2/B3-like transcriptional factor family protein | REM22 | Sampling day | 0.14 |
| AT1G58520.5 | lipases;hydrolases, acting on ester bonds | RXW8 | Sampling day | 0.22 |
| AT1G68825.1 | ROTUNDIFOLIA like 15 | RTFL15 | Sampling day | 0.17 |
| AT1G43040.2 | SAUR-like auxin-responsive protein family | - | Sampling day | 0.20 |
| AT5G63780.2 | RING/FYVE/PHD zinc finger superfamily protein | SHA1 | Sampling day | 0.22 |
| AT4G07090.1 | | - | Sampling day | 0.07 |
| AT2G06667.1 | NULL | - | Sampling day | 0.08 |
| AT4G25235.1 | NULL | - | Sampling day | 0.20 |
| AT3G01615.1 | | - | Sampling day | 0.12 |
| AT2G32250.5 | FAR1-related sequence 2 | FRS2 | Sampling day | 0.44 |
| AT4G07250.1 | | - | Sampling day | 0.15 |
| AT4G08114.1 | transposable element gene | - | Sampling day | 0.18 |
| AT1G73340.1 | Cytochrome P450 superfamily protein | - | Sampling day | 0.09 |
| AT2G46710.1 | Rho GTPase activating protein with PAK-box/P21-Rho-binding domain | ROPGAP3 | Sampling day | 0.10 |
| AT1G31640.1 | AGAMOUS-like 92 | AGL92 | Sampling day | 0.07 |
| AT2G19500.1 | cytokinin oxidase 2 | CKX2 | Sampling day | 0.07 |
| AT5G07015.1 | | - | Sampling day | 0.13 |
| AT5G50120.1 | Transducin/WD40 repeat-like superfamily protein | - | Sampling day | 0.08 |

| | | | | |
|---|---|---|---|---|
| AT1G19920.1 | Pseudouridine synthase/archaeosine transglycosylase-like family protein | APS2 | Sampling day | 0.20 |
| AT4G02900.2 | ERD (early-responsive to dehydration stress) family protein | - | Sampling day | 0.06 |
| AT1G09747.1 | | - | Sampling day | 0.12 |
| AT1G77060.1 | Phosphoenolpyruvate carboxylase family protein | - | Sampling day | 0.24 |
| AT5G24460.1 | NULL | - | Sampling day | 0.12 |
| AT4G32120.1 | Galactosyltransferase family protein | - | Sampling day | 0.15 |
| AT3G48130.1 | NULL | RSU1 | Sampling day | 0.22 |
| AT1G42615.1 | | - | Sampling day | 0.22 |
| AT2G08765.1 | | - | Sampling day | 0.08 |
| AT1G06737.1 | | - | Sampling day | 0.09 |
| AT2G09370.1 | | - | Sampling day | 0.19 |
| AT2G43240.1 | Nucleotide-sugar transporter family protein | - | Sampling day | 0.24 |
| AT2G37680.2 | NULL | - | Sampling day | 0.31 |
| AT2G07235.1 | | - | Sampling day | 0.16 |
| AT3G61310.1 | AT hook motif DNA-binding family protein | - | Sampling day | 0.15 |
| AT5G06935.1 | | - | Sampling day | 0.20 |
| AT5G32572.1 | transposable element gene | - | Sampling day | 0.09 |
| AT4G35210.1 | Arabidopsis protein of unknown function (DUF241) | - | Sampling day | 0.14 |
| AT1G03310.2 | debranching enzyme 1 | DBE1 | Sampling day | 0.26 |
| AT5G13610.1 | Protein of unknown function (DUF155) | - | Sampling day | 0.14 |
| AT2G01660.2 | plasmodesmata-located protein 6 | PDLP6 | Sampling day | 0.24 |
| AT5G30440.1 | transposable element gene | - | Sampling day | 0.24 |
| AT5G04060.1 | S-adenosyl-L-methionine-dependent methyltransferases superfamily | - | Sampling day | 0.15 |
| AT5G47950.1 | HXXXD-type acyl-transferase family protein | - | Sampling day | 0.26 |
| AT5G40810.2 | Cytochrome C1 family | - | Sampling day | 0.26 |
| AT3G43570.1 | GDSL-like Lipase/Acylhydrolase superfamily protein | - | Sampling day | 0.10 |
| AT1G07993.1 | | - | Sampling day | 0.06 |
| AT1G77480.2 | Eukaryotic aspartyl protease family protein | - | Sampling day | 0.18 |
| AT1G35143.1 | transposable element gene | - | Sampling day | 0.20 |
| AT5G58540.3 | Protein kinase superfamily protein | - | Sampling day | 0.18 |
| AT2G16700.3 | actin depolymerizing factor 5 | ADF5 | Sampling day | 0.11 |
| AT2G44065.3 | Ribosomal protein L2 family | - | Sampling day | 0.10 |
| AT3G01360.1 | Family of unknown function (DUF716) | - | Sampling day | 0.12 |
| AT5G53110.1 | RING/U-box superfamily protein | - | Sampling day | 0.14 |
| AT5G44230.2 | Pentatricopeptide repeat (PPR) superfamily protein | - | Sampling day | 0.09 |
| AT1G61130.1 | serine carboxypeptidase-like 32 | SCPL32 | Sampling day | 0.25 |
| AT5G08860.1 | | - | Sampling day | 0.14 |
| AT2G07780.1 | transposable element gene | - | Sampling day | 0.11 |
| AT2G14310.1 | transposable element gene | - | Sampling day | 0.27 |
| AT4G08065.1 | | - | Sampling day | 0.25 |
| AT5G18500.1 | Protein kinase superfamily protein | - | Sampling day | 0.08 |
| AT4G22560.1 | NULL | - | Sampling day | 0.10 |
| AT5G15025.1 | | - | Sampling day | 0.11 |
| AT1G20450.1 | Dehydrin family protein | ERD10 | Sampling day | 0.08 |
| AT2G02023.1 | NULL | - | Sampling day | 0.30 |
| AT4G25300.2 | 2-oxoglutarate (2OG) and Fe(II)-dependent oxygenase superfamily | - | Sampling day | 0.08 |
| AT4G32870.1 | Polyketide cyclase/dehydrase and lipid transport superfamily protein | - | Sampling day | 0.09 |
| AT5G04510.3 | 3'-phosphoinositide-dependent protein kinase 1 | PDK1 | Sampling day | 0.06 |
| AT4G20520.1 | RNA binding;RNA-directed DNA polymerases | - | Sampling day | 0.22 |
| AT3G05670.2 | RING/U-box protein | - | Sampling day | 0.12 |
| AT5G54320.1 | Protein of unknown function (DUF295) | - | Sampling day | 0.11 |
| AT1G16640.1 | AP2/B3-like transcriptional factor family protein | - | Sampling day | 0.22 |
| AT3G48201.1 | MIR861a; miRNA | MIR861A | Sampling day | 0.17 |
| AT3G03990.1 | alpha/beta-Hydrolases superfamily protein | - | Sampling day | 0.26 |
| AT3G04105.1 | | - | Sampling day | 0.13 |
| AT5G18820.2 | TCP-1/cpn60 chaperonin family protein | Cpn60alpha2 | Sampling day | 0.23 |
| AT3G57680.2 | Peptidase S41 family protein | - | Sampling day | 0.22 |
| AT2G20465.1 | Molecular chaperone Hsp40/DnaJ family protein | - | Sampling day | 0.06 |
| AT1G60835.1 | NULL | - | Sampling day | 0.09 |
| AT5G34770.1 | transposable element gene | - | Sampling day | 0.19 |
| AT1G05680.1 | Uridine diphosphate glycosyltransferase 74E2 | UGT74E2 | Sampling day | 0.23 |
| AT3G29290.1 | Pentatricopeptide repeat (PPR) superfamily protein | emb2076 | Sampling day | 0.13 |
| AT5G07955.1 | | - | Sampling day | 0.19 |
| AT4G03100.1 | Rho GTPase activating protein with PAK-box/P21-Rho-binding domain | - | Sampling day | 0.13 |
| AT4G27660.1 | NULL | - | Sampling day | 0.12 |
| AT3G28685.1 | pre-tRNA | - | Sampling day | 0.21 |
| AT1G27860.1 | Protein of unknown function (DUF626) | - | Sampling day | 0.28 |
| AT3G04440.2 | Plasma-membrane choline transporter family protein | - | Sampling day | 0.34 |
| AT2G27650.1 | Ubiquitin carboxyl-terminal hydrolase-related protein | - | Sampling day | 0.12 |
| AT3G04620.1 | Alba DNA/RNA-binding protein | DAN1 | Sampling day | 0.30 |
| AT5G03370.1 | acylphosphatase family | - | Sampling day | 0.05 |
| AT4G28410.2 | Tyrosine transaminase family protein | - | Sampling day | 0.22 |
| AT1G62170.2 | Serine protease inhibitor (SERPIN) family protein | - | Sampling day | 0.18 |
| AT5G24200.2 | alpha/beta-Hydrolases superfamily protein | - | Sampling day | 0.26 |
| AT1G02800.1 | cellulase 2 | CEL2 | Sampling day | 0.12 |
| AT4G04740.4 | calcium-dependent protein kinase 23 | CPK23 | Sampling day | 0.05 |

| | | | | |
|---|---|---|---|---|
| AT5G48640.4 | Cyclin family protein | - | Sampling day | 0.29 |
| AT4G09589.1 | | - | Sampling day | 0.10 |
| AT3G02105.1 | | - | Sampling day | 0.06 |
| AT5G16500.1 | Protein kinase superfamily protein | - | Sampling day | 0.16 |
| AT3G49960.1 | Peroxidase superfamily protein | - | Sampling day | 0.11 |
| AT1G68850.1 | Peroxidase superfamily protein | - | Sampling day | 0.26 |
| AT1G23200.1 | Plant invertase/pectin methylesterase inhibitor superfamily | - | Sampling day | 0.09 |
| AT1G09733.1 | | - | Sampling day | 0.25 |
| AT4G37685.1 | NULL | - | Sampling day | 0.14 |
| AT3G09045.1 | | - | Sampling day | 0.23 |
| AT1G03475.1 | Coproporphyrinogen III oxidase | LIN2 | Sampling day | 0.39 |
| AT1G09105.2 | | - | Sampling day | 0.11 |
| AT2G25490.1 | EIN3-binding F box protein 1 | EBF1 | Sampling day | 0.10 |
| AT2G20270.1 | Thioredoxin superfamily protein | - | Sampling day | 0.16 |
| AT1G60780.1 | Clathrin adaptor complexes medium subunit family protein | HAP13 | Sampling day | 0.17 |
| AT5G03944.1 | NULL | - | Sampling day | 0.25 |
| AT2G33840.1 | Tyrosyl-tRNA synthetase, class Ib, bacterial/mitochondrial | - | Sampling day | 0.22 |
| AT2G04180.1 | transposable element gene | - | Sampling day | 0.29 |
| AT3G03920.1 | H/ACA ribonucleoprotein complex, subunit Gar1/Naf1 protein | - | Sampling day | 0.21 |
| AT1G73885.1 | NULL | - | Sampling day | 0.35 |
| AT1G51030.1 | NULL | - | Sampling day | 0.34 |
| AT4G00955.1 | NULL | - | Sampling day | 0.16 |
| AT1G77910.1 | NULL | - | Sampling day | 0.23 |
| AT1G36270.1 | transposable element gene | - | Sampling day | 0.36 |
| AT3G07995.1 | | - | Sampling day | 0.21 |
| AT4G14420.1 | HR-like lesion-inducing protein-related | - | Sampling day | 0.07 |
| AT1G08797.1 | | - | Sampling day | 0.13 |
| AT1G07843.1 | | - | Sampling day | 0.27 |
| AT5G02640.1 | NULL | - | Sampling day | 0.21 |
| AT3G62410.1 | CP12 domain-containing protein 2 | CP12-2 | Sampling day | 0.10 |
| AT2G16145.1 | MIR416; miRNA | MIR416 | Sampling day | 0.18 |
| AT1G04443.1 | | - | Sampling day | 0.20 |
| AT4G21610.2 | lsd one like 2 | LOL2 | Sampling day | 0.10 |
| AT3G30885.1 | NULL | - | Sampling day | 0.09 |
| AT4G17430.1 | O-fucosyltransferase family protein | - | Sampling day | 0.09 |
| AT5G04495.1 | | - | Sampling day | 0.14 |
| AT5G07315.1 | pre-tRNA | - | Sampling day | 0.07 |
| AT1G23720.2 | Proline-rich extensin-like family protein | - | Sampling day | 0.07 |
| AT3G28020.1 | NULL | - | Sampling day | 0.28 |
| AT5G46750.1 | ARF-GAP domain 9 | AGD9 | Sampling day | 0.07 |
| AT2G43040.1 | tetratricopeptide repeat (TPR)-containing protein | NPG1 | Sampling day | 0.12 |
| AT4G07460.1 | transposable element gene | - | Sampling day | 0.08 |
| AT4G22890.1 | PGR5-LIKE A | PGR5-LIKE A | Sampling day | 0.06 |
| AT5G04710.1 | Zn-dependent exopeptidases superfamily protein | - | Sampling day | 0.19 |
| AT2G41720.3 | Tetratricopeptide repeat (TPR)-like superfamily protein | EMB2654 | Sampling day | 0.16 |
| AT1G27660.1 | basic helix-loop-helix (bHLH) DNA-binding superfamily protein | - | Sampling day | 0.11 |
| AT5G39240.1 | NULL | - | Sampling day | 0.23 |
| AT3G18827.1 | MIR868a; miRNA | MIR868A | Sampling day | 0.16 |
| AT3G50290.1 | HXXXD-type acyl-transferase family protein | - | Sampling day | 0.13 |
| AT4G02055.1 | pre-tRNA | - | Sampling day | 0.06 |
| AT2G39730.2 | rubisco activase | RCA | Sampling day | 0.20 |
| AT1G63300.1 | Myosin heavy chain-related protein | - | Sampling day | 0.08 |
| AT5G56605.1 | transposable element gene | - | Sampling day | 0.08 |
| AT2G07665.1 | | - | Sampling day | 0.17 |
| AT2G43890.1 | Pectin lyase-like superfamily protein | - | Sampling day | 0.08 |
| AT1G28591.1 | NULL | - | Sampling day | 0.18 |
| AT4G34440.1 | Protein kinase superfamily protein | PERK5 | Sampling day | 0.08 |
| AT1G29410.2 | phosphoribosylanthranilate isomerase 3 | PAI3 | Sampling day | 0.06 |
| AT4G02400.1 | U3 ribonucleoprotein (Utp) family protein | - | Sampling day | 0.09 |
| AT1G09470.2 | NULL | - | Sampling day | 0.06 |
| AT1G23970.2 | Protein of unknown function (DUF626) | - | Sampling day | 0.21 |
| AT2G34270.1 | NULL | - | Sampling day | 0.11 |
| AT5G08890.1 | | - | Sampling day | 0.07 |
| AT5G48690.3 | NULL | - | Sampling day | 0.09 |
| AT5G22150.1 | NULL | - | Sampling day | 0.09 |
| AT5G20830.1 | sucrose synthase 1 | SUS1 | Sampling day | 0.17 |
| AT4G39840.1 | NULL | - | Sampling day | 0.16 |
| AT4G00590.2 | N-terminal nucleophile aminohydrolases (Ntn hydrolases) superfamily | - | Sampling day | 0.08 |
| AT4G20330.1 | Transcription initiation factor TFIIE, beta subunit | - | Sampling day | 0.10 |
| AT4G29430.1 | ribosomal protein S15A E | rps15ae | Sampling day | 0.07 |
| AT1G15030.1 | Protein of unknown function (DUF789) | - | Sampling day | 0.13 |
| ATMG00950.1 | NULL | TRNQ | Sampling day | 0.20 |
| AT2G05000.1 | transposable element gene | - | Sampling day | 0.21 |
| AT4G09795.1 | low-molecular-weight cysteine-rich 13 | LCR13 | Sampling day | 0.14 |
| AT5G37872.1 | transposable element gene | - | Sampling day | 0.16 |
| AT1G64170.1 | cation/H+ exchanger 16 | CHX16 | Sampling day | 0.10 |

| | | | | |
|---|---|---|---|---|
| AT2G45600.1 | alpha/beta-Hydrolases superfamily protein | - | Sampling day | 0.18 |
| AT1G50600.2 | scarecrow-like 5 | SCL5 | Sampling day | 0.17 |
| AT5G47260.1 | ATP binding;GTP binding;nucleotide binding;nucleoside- | - | Sampling day | 0.07 |
| AT4G13340.1 | Leucine-rich repeat (LRR) family protein | LRX3 | Sampling day | 0.20 |
| AT2G28815.1 | | - | Sampling day | 0.29 |
| AT2G04955.1 | | - | Sampling day | 0.06 |
| AT5G02505.1 | pre-tRNA | - | Sampling day | 0.20 |
| AT2G29930.1 | F-box/RNI-like superfamily protein | - | Sampling day | 0.08 |
| AT1G16350.1 | Aldolase-type TIM barrel family protein | - | Sampling day | 0.36 |
| AT3G28360.1 | P-glycoprotein 16 | ABCB16 | Sampling day | 0.20 |
| AT4G24200.2 | Transcription elongation factor (TFIIS) family protein | - | Sampling day | 0.11 |
| AT4G17005.1 | transposable element gene | - | Sampling day | 0.23 |
| AT4G08333.1 | transposable element gene | - | Sampling day | 0.13 |
| AT2G47030.1 | Plant invertase/pectin methylesterase inhibitor superfamily | VGDH1 | Sampling day | 0.12 |
| AT2G44740.1 | cyclin p4;1 | CYCP4%3B1 | Sampling day | 0.23 |
| AT5G45905.1 | NULL | - | Sampling day | 0.14 |
| AT2G25440.1 | receptor like protein 20 | RLP20 | Sampling day | 0.21 |
| AT2G29735.1 | | - | Sampling day | 0.20 |
| AT1G26225.1 | | - | Sampling day | 0.06 |
| AT4G08165.1 | | - | Sampling day | 0.06 |
| AT4G07700.1 | transposable element gene | - | Sampling day | 0.18 |
| AT2G33860.1 | Transcriptional factor B3 family protein / auxin-responsive factor AUX/IAA-related | ETT | Sampling day | 0.12 |
| AT1G30500.2 | nuclear factor Y, subunit A7 | NF-YA7 | Sampling day | 0.22 |
| AT5G11370.1 | FBD / Leucine Rich Repeat domains containing protein | - | Sampling day | 0.11 |
| AT3G42590.1 | transposable element gene | - | Sampling day | 0.08 |
| AT3G24300.1 | ammonium transporter 1;3 | AMT1%3B3 | Sampling day | 0.18 |
| AT2G40860.3 | protein kinase family protein / protein phosphatase 2C ( PP2C) family | - | Sampling day | 0.14 |
| AT2G06925.1 | Phospholipase A2 family protein | PLA2-ALPHA | Sampling day | 0.26 |
| AT2G36490.1 | demeter-like 1 | DML1 | Sampling day | 0.12 |
| AT3G33069.1 | transposable element gene | - | Sampling day | 0.18 |
| AT5G42720.1 | Glycosyl hydrolase family 17 protein | - | Sampling day | 0.26 |
| AT2G33030.1 | receptor like protein 25 | RLP25 | Sampling day | 0.18 |
| AT5G49800.1 | Polyketide cyclase/dehydrase and lipid transport superfamily protein | - | Sampling day | 0.28 |
| AT2G26910.1 | pleiotropic drug resistance 4 | ABCG32 | Sampling day | 0.15 |
| AT5G43930.3 | Transducin family protein / WD-40 repeat family protein | - | Sampling day | 0.07 |
| ATCG00130.1 | ATPase, F0 complex, subunit B/B', bacterial/chloroplast | ATPF | Sampling day | 0.20 |
| AT4G19770.1 | Glycosyl hydrolase family protein with chitinase insertion domain | - | Sampling day | 0.18 |
| AT1G12590.1 | pre-tRNA | - | Sampling day | 0.16 |
| AT4G14020.1 | Rapid alkalinization factor (RALF) family protein | - | Sampling day | 0.10 |
| AT3G30580.1 | NULL | - | Sampling day | 0.07 |
| AT1G72440.1 | CCAAT-binding factor | EDA25 | Sampling day | 0.07 |
| AT5G24000.1 | Protein of unknown function (DUF819) | - | Sampling day | 0.09 |
| ATCG01200.1 | NULL | TRNI.3 | Sampling day | 0.11 |
| AT1G53600.1 | Tetratricopeptide repeat (TPR)-like superfamily protein | - | Sampling day | 0.20 |
| AT1G67785.1 | NULL | - | Sampling day | 0.28 |
| AT1G02530.2 | P-glycoprotein 12 | ABCB12 | Sampling day | 0.10 |
| AT3G15640.1 | Rubredoxin-like superfamily protein | - | Sampling day | 0.30 |
| AT5G66360.2 | Ribosomal RNA adenine dimethylase family protein | DIM1B | Sampling day | 0.13 |
| AT5G20750.1 | transposable element gene | - | Sampling day | 0.12 |
| AT3G02350.1 | galacturonosyltransferase 9 | GAUT9 | Sampling day | 0.09 |
| AT3G08410.1 | | - | Sampling day | 0.11 |
| AT1G07051.1 | MIR847a; miRNA | MIR847A | Sampling day | 0.14 |
| AT3G45650.2 | nitrate excretion transporter1 | NAXT1 | Sampling day | 0.06 |
| AT1G68930.1 | pentatricopeptide (PPR) repeat-containing protein | - | Sampling day | 0.24 |
| AT4G11580.1 | RNI-like superfamily protein | - | Sampling day | 0.11 |
| AT3G04175.1 | | - | Sampling day | 0.15 |
| AT5G57200.1 | ENTH/ANTH/VHS superfamily protein | - | Sampling day | 0.29 |
| AT3G02095.1 | | - | Sampling day | 0.23 |
| AT5G56070.1 | NULL | - | Sampling day | 0.10 |
| AT3G44470.2 | transposable element gene | - | Sampling day | 0.09 |
| AT1G17820.1 | Putative integral membrane protein conserved region (DUF2404) | - | Sampling day | 0.07 |
| AT3G22740.1 | homocysteine S-methyltransferase 3 | HMT3 | Sampling day | 0.09 |
| AT4G05580.1 | transposable element gene | - | Sampling day | 0.14 |
| AT3G46180.2 | UDP-galactose transporter 5 | UTR5 | Sampling day | 0.14 |
| AT4G21595.1 | MIR169G; miRNA | MIR169G | Sampling day | 0.18 |
| AT5G11490.3 | adaptin family protein | - | Sampling day | 0.15 |
| AT1G13040.2 | Pentatricopeptide repeat (PPR-like) superfamily protein | - | Sampling day | 0.34 |
| AT4G18593.2 | dual specificity protein phosphatase-related | - | Sampling day | 0.15 |
| AT3G26170.1 | cytochrome P450, family 71, subfamily B, polypeptide 19 | CYP71B19 | Sampling day | 0.08 |
| AT4G14450.1 | NULL | - | Sampling day | 0.13 |
| AT3G05325.1 | | - | Sampling day | 0.16 |
| AT3G17265.1 | F-box and associated interaction domains-containing protein | - | Sampling day | 0.11 |
| AT1G79190.1 | ARM repeat superfamily protein | - | Sampling day | 0.11 |
| AT4G24320.1 | Ubiquitin carboxyl-terminal hydrolase family protein | - | Sampling day | 0.12 |
| AT5G22140.1 | FAD/NAD(P)-binding oxidoreductase family protein | - | Sampling day | 0.06 |

| | | | | |
|---|---|---|---|---|
| AT1G60810.2 | ATP-citrate lyase A-2 | ACLA-2 | Sampling day | 0.18 |
| AT1G50960.1 | gibberellin 2-oxidase 7 | GA2OX7 | Sampling day | 0.21 |
| AT1G37012.1 | transposable element gene | - | Sampling day | 0.11 |
| AT1G02065.2 | squamosa promoter binding protein-like 8 | SPL8 | Sampling day | 0.08 |
| AT1G14190.2 | Glucose-methanol-choline (GMC) oxidoreductase family protein | - | Sampling day | 0.45 |
| AT3G25070.2 | RPM1 interacting protein 4 | RIN4 | Sampling day | 0.09 |
| AT4G19070.1 | Putative membrane lipoprotein | - | Sampling day | 0.15 |
| AT4G06630.1 | transposable element gene | - | Sampling day | 0.28 |
| AT5G39995.1 | NULL | - | Sampling day | 0.09 |
| AT3G03830.1 | SAUR-like auxin-responsive protein family | SAUR28 | Sampling day | 0.08 |
| AT1G04190.1 | Tetratricopeptide repeat (TPR)-like superfamily protein | TPR3 | Sampling day | 0.09 |
| AT2G36950.1 | Heavy metal transport/detoxification superfamily protein | - | Sampling day | 0.27 |
| AT2G03270.1 | DNA-binding protein, putative | - | Sampling day | 0.15 |
| AT3G18310.1 | NULL | - | Sampling day | -0.46 |
| AT5G46240.1 | potassium channel in Arabidopsis thaliana 1 | KAT1 | Sampling day | -0.05 |
| AT2G38310.1 | PYR1-like 4 | PYL4 | Sampling day | -0.08 |
| AT5G05185.1 | | - | Sampling day | -0.10 |
| AT1G09943.1 | | - | Sampling day | -0.16 |
| AT3G13820.1 | F-box and associated interaction domains-containing protein | - | Sampling day | -0.18 |
| AT2G09180.1 | | - | Sampling day | -0.15 |
| AT5G05445.1 | | - | Sampling day | -0.15 |
| AT1G58010.1 | NULL | - | Sampling day | -0.14 |
| AT1G53903.1 | Protein of unknown function (DUF581) | - | Sampling day | -0.14 |
| AT1G05597.1 | | - | Sampling day | -0.05 |
| AT2G07719.1 | Putative membrane lipoprotein | - | Sampling day | -0.07 |
| AT3G52760.1 | Integral membrane Yip1 family protein | - | Sampling day | -0.40 |
| AT5G52750.2 | Heavy metal transport/detoxification superfamily protein | - | Sampling day | -0.10 |
| AT3G08315.1 | | - | Sampling day | -0.06 |
| AT2G36060.1 | MMS ZWEI homologue 3 | MMZ3 | Sampling day | -0.22 |
| AT3G02135.1 | | - | Sampling day | -0.14 |
| AT3G36659.1 | Plant invertase/pectin methylesterase inhibitor superfamily protein | - | Sampling day | -0.22 |
| AT4G03635.1 | NULL | - | Sampling day | -0.17 |
| AT4G22320.2 | NULL | - | Sampling day | -0.38 |
| AT3G62620.4 | sucrose-phosphatase-related | - | Sampling day | -0.22 |
| AT1G06827.1 | | - | Sampling day | -0.09 |
| AT4G34970.1 | actin depolymerizing factor 9 | ADF9 | Sampling day | -0.10 |
| AT5G34839.1 | transposable element gene | - | Sampling day | -0.07 |
| AT4G32050.1 | neurochondrin family protein | - | Sampling day | -0.05 |
| AT1G17140.1 | interactor of constitutive active rops 1 | ICR1 | Sampling day | -0.06 |
| AT5G26030.1 | ferrochelatase 1 | FC1 | Sampling day | -0.16 |
| AT3G51880.1 | high mobility group B1 | HMGB1 | Sampling day | -0.46 |
| AT2G47470.1 | thioredoxin family protein | UNE5 | Sampling day | -0.17 |
| AT5G49240.1 | pseudo-response regulator 4 | APRR4 | Sampling day | -0.22 |
| AT3G27500.1 | Cysteine/Histidine-rich C1 domain family protein | - | Sampling day | -0.14 |
| AT2G03620.2 | magnesium transporter 3 | MGT3 | Sampling day | -0.19 |
| AT3G50840.3 | Phototropic-responsive NPH3 family protein | - | Sampling day | -0.22 |
| AT5G05985.1 | pre-tRNA | - | Sampling day | -0.23 |
| AT1G08357.1 | | - | Sampling day | -0.13 |
| AT1G23160.1 | Auxin-responsive GH3 family protein | - | Sampling day | -0.14 |
| AT5G36310.1 | ECA1 gametogenesis related family protein | - | Sampling day | -0.12 |
| AT5G23395.1 | Cox19-like CHCH family protein | MIA40 | Sampling day | -0.06 |
| AT4G32070.1 | Octicosapeptide/Phox/Bem1p (PB1) domain-containing protein / tetratricopeptide repeat (TPR)-containing protein | Phox4 | Sampling day | -0.60 |
| AT3G51370.1 | Protein phosphatase 2C family protein | - | Sampling day | -0.06 |
| AT1G01740.3 | Protein kinase protein with tetratricopeptide repeat domain | BSK4 | Sampling day | -0.12 |
| AT1G65830.1 | pre-tRNA | - | Sampling day | -0.22 |
| AT4G23150.1 | cysteine-rich RLK (RECEPTOR-like protein kinase) 7 | CRK7 | Sampling day | -0.39 |
| AT3G27480.1 | Cysteine/Histidine-rich C1 domain family protein | - | Sampling day | -0.23 |
| AT1G68490.1 | NULL | - | Sampling day | -0.15 |
| AT1G18970.1 | germin-like protein 4 | GLP4 | Sampling day | -0.24 |
| AT3G43723.1 | transposable element gene | - | Sampling day | -0.18 |
| AT1G22210.1 | Haloacid dehalogenase-like hydrolase (HAD) superfamily protein | TPPC | Sampling day | -0.26 |
| AT5G28920.1 | NULL | - | Sampling day | -0.10 |
| AT4G26790.1 | GDSL-like Lipase/Acylhydrolase superfamily protein | - | Sampling day | -0.14 |
| AT4G07498.1 | transposable element gene | - | Sampling day | -0.16 |
| AT5G59720.1 | heat shock protein 18.2 | HSP18.2 | Sampling day | -0.13 |
| AT3G26250.1 | Cysteine/Histidine-rich C1 domain family protein | - | Sampling day | -0.24 |
| AT5G33433.1 | transposable element gene | - | Sampling day | -0.10 |
| AT1G04817.1 | | - | Sampling day | -0.12 |
| AT1G62720.1 | Pentatricopeptide repeat (PPR-like) superfamily protein | NG1 | Sampling day | -0.10 |
| AT1G02570.1 | NULL | - | Sampling day | -0.09 |
| AT1G26790.1 | Dof-type zinc finger DNA-binding family protein | - | Sampling day | -0.05 |
| AT1G01350.1 | Zinc finger (CCCH-type/C3HC4-type RING finger) family protein | - | Sampling day | -0.06 |
| AT3G20541.1 | NULL | - | Sampling day | -0.15 |
| AT3G58930.1 | F-box/RNI-like superfamily protein | - | Sampling day | -0.07 |
| AT1G71830.1 | somatic embryogenesis receptor-like kinase 1 | SERK1 | Sampling day | -0.06 |

| | | | | |
|---|---|---|---|---|
| AT5G05970.2 | Transducin/WD40 repeat-like superfamily protein | NEDD1 | Sampling day | -0.14 |
| AT1G17710.1 | Pyridoxal phosphate phosphatase-related protein | PEPC1 | Sampling day | -0.27 |
| AT5G19880.1 | Peroxidase superfamily protein | - | Sampling day | -0.24 |
| AT5G06515.1 | | - | Sampling day | -0.18 |
| AT2G32415.3 | Polynucleotidyl transferase, ribonuclease H fold protein with HRDC | - | Sampling day | -0.07 |
| AT1G03120.2 | NULL | RAB28 | Sampling day | -0.15 |
| AT1G43130.2 | like COV 2 | LCV2 | Sampling day | -0.36 |
| AT2G35310.1 | Transcriptional factor B3 family protein | - | Sampling day | -0.13 |
| AT5G64760.3 | regulatory particle non-ATPase subunit 5B | RPN5B | Sampling day | -0.55 |
| AT2G34655.1 | NULL | - | Sampling day | -0.12 |
| AT4G40042.1 | Microsomal signal peptidase 12 kDa subunit (SPC12) | - | Sampling day | -0.10 |
| AT5G42620.1 | metalloendopeptidases;zinc ion binding | - | Sampling day | -0.06 |
| AT2G23850.1 | NULL | - | Sampling day | -0.07 |
| AT2G24850.1 | tyrosine aminotransferase 3 | TAT3 | Sampling day | -0.19 |
| AT4G03555.1 | | - | Sampling day | -0.21 |
| AT5G43935.1 | flavonol synthase 6 | FLS6 | Sampling day | -0.22 |
| ATMG01190.1 | ATP synthase subunit 1 | ATP1 | Sampling day | -0.18 |
| AT1G64160.1 | Disease resistance-responsive (dirigent-like protein) family protein | DIR5 | Sampling day | -0.19 |
| AT1G51600.1 | ZIM-LIKE 2 | ZML2 | Sampling day | -0.44 |
| AT2G34010.2 | NULL | - | Sampling day | -0.23 |
| AT3G49750.1 | receptor like protein 44 | RLP44 | Sampling day | -0.26 |
| AT1G35040.1 | NULL | - | Sampling day | -0.16 |
| AT3G04815.1 | | - | Sampling day | -0.13 |
| AT1G74250.1 | DNAJ heat shock N-terminal domain-containing protein | - | Sampling day | -0.07 |
| AT1G32510.1 | NAC domain containing protein 11 | NAC011 | Sampling day | -0.09 |
| AT5G63905.1 | NULL | - | Sampling day | -0.29 |
| AT3G05955.1 | | - | Sampling day | -0.12 |
| AT4G37340.1 | cytochrome P450, family 81, subfamily D, polypeptide 3 | CYP81D3 | Sampling day | -0.10 |
| AT3G42190.1 | transposable element gene | - | Sampling day | -0.22 |
| AT3G57930.2 | NULL | - | Sampling day | -0.13 |
| AT5G36030.1 | transposable element gene | - | Sampling day | -0.26 |
| AT1G17280.6 | ubiquitin-conjugating enzyme 34 | UBC34 | Sampling day | -0.19 |
| AT3G42083.1 | transposable element gene | - | Sampling day | -0.14 |
| AT5G21150.1 | Argonaute family protein | AGO9 | Sampling day | -0.26 |
| AT5G03545.1 | NULL | - | Sampling day | -0.19 |
| AT2G47680.1 | zinc finger (CCCH type) helicase family protein | - | Sampling day | -0.11 |
| AT5G61040.1 | NULL | - | Sampling day | -0.19 |
| AT2G08205.1 | | - | Sampling day | -0.07 |
| AT1G08955.1 | | - | Sampling day | -0.17 |
| AT5G05240.1 | Uncharacterised conserved protein (UCP030365) | - | Sampling day | -0.08 |
| AT5G25950.1 | Protein of Unknown Function (DUF239) | - | Sampling day | -0.13 |
| AT1G02180.1 | ferredoxin-related | - | Sampling day | -0.17 |
| AT1G49370.1 | NULL | - | Sampling day | -0.12 |
| AT1G09610.1 | Protein of unknown function (DUF579) | GXM3 | Sampling day | -0.05 |
| AT1G09823.1 | | - | Sampling day | -0.10 |
| AT1G21600.1 | plastid transcriptionally active 6 | PTAC6 | Sampling day | -0.41 |
| AT4G00280.1 | NULL | - | Sampling day | -0.20 |
| AT4G18280.1 | glycine-rich cell wall protein-related | - | Sampling day | -0.43 |
| AT2G05335.1 | SCR-like 15 | SCRL15 | Sampling day | -0.24 |
| AT2G13270.1 | transposable element gene | - | Sampling day | -0.18 |
| AT2G47585.1 | MIR164/MIR164A; miRNA | MIR164A | Sampling day | -0.12 |
| AT4G32960.1 | NULL | - | Sampling day | -0.42 |
| AT5G65170.1 | VQ motif-containing protein | - | Sampling day | -0.37 |
| AT4G36515.1 | NULL | - | Sampling day | -0.08 |
| AT2G33170.1 | Leucine-rich repeat receptor-like protein kinase family protein | - | Sampling day | -0.20 |
| AT5G53920.1 | ribosomal protein L11 methyltransferase-related | - | Sampling day | -0.12 |
| AT1G53345.1 | NULL | - | Sampling day | -0.14 |
| AT5G00455.1 | | - | Sampling day | -0.09 |
| AT5G03325.1 | | - | Sampling day | -0.06 |
| AT2G35600.3 | BREVIS RADIX-like 1 | BRXL1 | Sampling day | -0.10 |
| AT5G55460.1 | Bifunctional inhibitor/lipid-transfer protein/seed storage 2S albumin superfamily protein | - | Sampling day | -0.15 |
| AT3G21160.2 | alpha-mannosidase 2 | MNS2 | Sampling day | -0.08 |
| AT1G76065.1 | LYR family of Fe/S cluster biogenesis protein | - | Sampling day | -0.24 |
| AT1G73970.2 | NULL | - | Sampling day | -0.11 |
| AT4G24805.1 | S-adenosyl-L-methionine-dependent methyltransferases superfamily | - | Sampling day | -0.21 |
| AT2G34570.1 | PIN domain-like family protein | MEE21 | Sampling day | -0.24 |
| AT5G17760.2 | P-loop containing nucleoside triphosphate hydrolases superfamily | - | Sampling day | -0.23 |
| AT2G37050.4 | Leucine-rich repeat protein kinase family protein | - | Sampling day | -0.13 |
| AT3G10114.1 | NULL | - | Sampling day | -0.21 |
| AT4G12570.1 | ubiquitin protein ligase 5 | UPL5 | Sampling day | -0.11 |
| AT3G33595.1 | transposable element gene | - | Sampling day | -0.22 |
| AT5G47300.1 | F-box and associated interaction domains-containing protein | - | Sampling day | -0.15 |
| AT3G01595.1 | | - | Sampling day | -0.11 |
| AT3G62710.1 | Glycosyl hydrolase family protein | - | Sampling day | -0.17 |
| AT2G37450.3 | nodulin MtN21 /EamA-like transporter family protein | UMAMIT13 | Sampling day | -0.18 |

| | | | | |
|---|---|---|---|---|
| AT3G59960.2 | histone-lysine N-methyltransferase ASHH4 | ASHH4 | Sampling day | -0.29 |
| AT1G23300.1 | MATE efflux family protein | - | Sampling day | -0.12 |
| AT3G44757.1 | NULL | - | Sampling day | -0.17 |
| AT4G27550.1 | trehalose-6-phosphatase synthase S4 | TPS4 | Sampling day | -0.08 |
| AT5G55270.1 | Protein of unknown function (DUF295) | - | Sampling day | -0.13 |
| AT2G35030.1 | Pentatricopeptide repeat (PPR) superfamily protein | - | Sampling day | -0.32 |
| AT5G31637.1 | transposable element gene | - | Sampling day | -0.11 |
| AT1G54180.2 | BREVIS RADIX-like 3 | BRX-LIKE3 | Sampling day | -0.21 |
| AT2G09953.1 | transposable element gene | - | Sampling day | -0.28 |
| AT4G07470.1 | transposable element gene | - | Sampling day | -0.14 |
| AT4G33050.5 | calmodulin-binding family protein | EDA39 | Sampling day | -0.12 |
| AT2G04715.1 | | - | Sampling day | -0.17 |
| AT3G27473.1 | Cysteine/Histidine-rich C1 domain family protein | - | Sampling day | -0.14 |
| AT1G72630.1 | ELF4-like 2 | ELF4-L2 | Sampling day | -0.18 |
| AT4G24820.1 | 26S proteasome, regulatory subunit Rpn7;Proteasome component (PCI) domain | - | Sampling day | -0.12 |
| AT4G24690.1 | ubiquitin-associated (UBA)/TS-N domain-containing protein / octicosapeptide/Phox/Bemp1 (PB1) domain-containing protein | NBR1 | Sampling day | -0.08 |
| AT4G32285.2 | ENTH/ANTH/VHS superfamily protein | - | Sampling day | -0.31 |
| AT5G36659.1 | ECA1 gametogenesis related family protein | - | Sampling day | -0.19 |
| AT4G20700.2 | Protein of unknown function (DUF1204) | - | Sampling day | -0.08 |
| AT3G03140.1 | Tudor/PWWP/MBT superfamily protein | - | Sampling day | -0.24 |
| AT5G07320.1 | Mitochondrial substrate carrier family protein | APC3 | Sampling day | -0.06 |
| AT1G36495.1 | transposable element gene | - | Sampling day | -0.10 |
| AT4G06915.1 | | - | Sampling day | -0.46 |
| AT2G02690.2 | Cysteine/Histidine-rich C1 domain family protein | - | Sampling day | -0.10 |
| AT1G16025.1 | NULL | - | Sampling day | -0.21 |
| AT4G21240.1 | F-box and associated interaction domains-containing protein | - | Sampling day | -0.28 |
| AT1G20390.1 | transposable element gene | - | Sampling day | -0.15 |
| AT5G03195.1 | | - | Sampling day | -0.24 |
| AT3G18090.3 | nuclear RNA polymerase D2B | NRPD2B | Sampling day | -0.09 |
| AT1G09813.1 | | - | Sampling day | -0.12 |
| AT2G21800.3 | essential meiotic endonuclease 1A | EME1A | Sampling day | -0.23 |
| AT3G25830.1 | terpene synthase-like sequence-1,8-cineole | TPS-CIN | Sampling day | -0.19 |
| AT1G80520.1 | Sterile alpha motif (SAM) domain-containing protein | - | Sampling day | -0.18 |
| AT3G05135.1 | | - | Sampling day | -0.22 |
| AT1G23560.1 | Domain of unknown function (DUF220) | - | Sampling day | -0.14 |
| ATMG00340.1 | NULL | TRNY.1 | Sampling day | -0.08 |
| AT2G29165.1 | transposable element gene | - | Sampling day | -0.22 |
| AT1G54930.2 | GRF zinc finger / Zinc knuckle protein | - | Sampling day | -0.29 |
| AT1G45190.1 | downregulated in DIF1 18 | DD18 | Sampling day | -0.16 |
| AT3G14730.1 | Pentatricopeptide repeat (PPR) superfamily protein | - | Sampling day | -0.24 |
| AT3G23980.1 | BLISTER | BLI | Sampling day | -0.10 |
| AT5G02450.1 | Ribosomal protein L36e family protein | - | Sampling day | -0.10 |
| AT2G05675.1 | | - | Sampling day | -0.17 |
| AT4G21710.1 | DNA-directed RNA polymerase family protein | NRPB2 | Sampling day | -0.17 |
| AT2G26270.1 | NULL | - | Sampling day | -0.19 |
| AT5G48820.1 | inhibitor/interactor with cyclin-dependent kinase | ICK6 | Sampling day | -0.14 |
| AT5G39863.1 | NULL | - | Sampling day | -0.19 |
| AT4G39200.1 | Ribosomal protein S25 family protein | - | Sampling day | -0.18 |
| AT2G07530.1 | transposable element gene | - | Sampling day | -0.19 |
| AT1G70190.1 | Ribosomal protein L7/L12, oligomerisation;Ribosomal protein L7/L12, C-terminal/adaptor protein ClpS-like | - | Sampling day | -0.09 |
| AT5G02605.1 | | - | Sampling day | -0.21 |
| AT2G13145.1 | NULL | - | Sampling day | -0.17 |
| AT1G08753.1 | | - | Sampling day | -0.18 |
| AT1G24170.1 | Nucleotide-diphospho-sugar transferases superfamily protein | LGT9 | Sampling day | -0.17 |
| AT4G04435.1 | | - | Sampling day | -0.14 |
| AT3G26310.1 | cytochrome P450, family 71, subfamily B, polypeptide 35 | CYP71B35 | Sampling day | -0.18 |
| AT5G52355.1 | pre-tRNA | - | Sampling day | -0.10 |
| AT1G13210.1 | autoinhibited Ca2+/ATPase II | ACA.l | Sampling day | -0.19 |
| AT5G43140.2 | Peroxisomal membrane 22 kDa (Mpv17/PMP22) family protein | - | Sampling day | -0.07 |
| AT3G52742.1 | other RNA | - | Sampling day | -0.19 |
| AT3G10030.2 | aspartate/glutamate/uridylate kinase family protein | - | Sampling day | -0.21 |
| AT5G46850.2 | NULL | - | Sampling day | -0.10 |
| AT1G61240.6 | Protein of unknown function (DUF707) | - | Sampling day | -0.18 |
| AT5G33226.1 | transposable element gene | - | Sampling day | -0.11 |
| AT3G09860.1 | NULL | - | Sampling day | -0.23 |
| AT4G14400.1 | ankyrin repeat family protein | ACD6 | Sampling day | -0.13 |
| AT1G51405.1 | myosin-related | - | Sampling day | -0.20 |
| AT1G05697.1 | | - | Sampling day | -0.10 |
| AT5G66550.2 | Maf-like protein | - | Sampling day | -0.22 |
| AT5G47090.1 | NULL | - | Sampling day | -0.21 |
| AT3G08055.1 | | - | Sampling day | -0.38 |
| AT5G12235.1 | CLAVATA3/ESR-RELATED 22 | CLE22 | Sampling day | -0.12 |
| AT4G21630.1 | Subtilase family protein | - | Sampling day | -0.18 |

| | | | | |
|---|---|---|---|---|
| AT3G63530.2 | RING/U-box superfamily protein | BB | Sampling day | -0.06 |
| AT5G06030.1 | Plant self-incompatibility protein S1 family | - | Sampling day | -0.19 |
| AT5G64400.1 | NULL | - | Sampling day | -0.21 |
| AT1G09065.1 | | - | Sampling day | -0.17 |
| AT5G53120.6 | spermidine synthase 3 | SPDS3 | Sampling day | -0.06 |
| AT4G23885.1 | NULL | - | Sampling day | -0.13 |
| AT5G41491.1 | NULL | - | Sampling day | -0.06 |
| AT5G56470.1 | FAD-dependent oxidoreductase family protein | GulLO7 | Sampling day | -0.13 |
| AT5G47360.1 | Tetratricopeptide repeat (TPR)-like superfamily protein | - | Sampling day | -0.13 |
| AT2G37910.1 | cation/hydrogen exchanger, putative (CHX21) | - | Sampling day | -0.14 |
| AT4G29990.1 | Leucine-rich repeat transmembrane protein kinase protein | - | Sampling day | -0.08 |
| ATMG01100.1 | NULL | ORF105A | Sampling day | -0.18 |
| AT3G14130.2 | Aldolase-type TIM barrel family protein | HAOX1 | Sampling day | -0.07 |
| AT1G52600.1 | Peptidase S24/S26A/S26B/S26C family protein | - | Sampling day | -0.11 |
| AT3G54340.1 | K-box region and MADS-box transcription factor family protein | AP3 | Sampling day | -0.19 |
| AT2G36840.1 | ACT-like superfamily protein | ACR10 | Sampling day | -0.24 |
| AT1G05493.1 | | - | Sampling day | -0.20 |
| AT3G10572.2 | 3-phosphoinositide-dependent protein kinase-1, putative | APEM9 | Sampling day | -0.14 |
| AT4G29010.1 | Enoyl-CoA hydratase/isomerase family | AIM1 | Sampling day | -0.19 |
| AT5G09370.2 | Bifunctional inhibitor/lipid-transfer protein/seed storage 2S albumin superfamily protein | - | Sampling day | -0.11 |
| AT3G41762.1 | NULL | - | Sampling day | -0.17 |
| AT4G11945.1 | transposable element gene | - | Sampling day | -0.11 |
| AT5G33389.1 | transposable element gene | - | Sampling day | -0.28 |
| AT3G55795.1 | pre-tRNA | - | Sampling day | -0.17 |
| AT1G65070.3 | DNA mismatch repair protein MutS, type 2 | - | Sampling day | -0.20 |
| AT1G29195.1 | NULL | - | Sampling day | -0.14 |
| AT1G62570.1 | flavin-monooxygenase glucosinolate S-oxygenase 4 | FMO GS-OX4 | Sampling day | -0.22 |
| AT1G16970.1 | KU70 homolog | KU70 | Sampling day | -0.09 |
| AT4G33880.1 | ROOT HAIR DEFECTIVE 6-LIKE 2 | RSL2 | Sampling day | -0.12 |
| AT1G33260.1 | Protein kinase superfamily protein | - | Sampling day | -0.10 |
| AT5G03925.1 | | - | Sampling day | -0.11 |
| AT5G55260.2 | protein phosphatase X 2 | PPX2 | Sampling day | -0.14 |
| AT4G11030.1 | AMP-dependent synthetase and ligase family protein | - | Sampling day | -0.17 |
| AT4G21400.8 | cysteine-rich RLK (RECEPTOR-like protein kinase) 28 | CRK28 | Sampling day | -0.20 |
| AT5G61120.2 | NULL | - | Sampling day | -0.12 |
| AT5G28390.1 | RNA-binding (RRM/RBD/RNP motifs) family protein | - | Sampling day | -0.11 |
| AT3G62060.2 | Pectinacetylesterase family protein | - | Sampling day | -0.10 |
| AT3G59580.2 | Plant regulator RWP-RK family protein | - | Sampling day | -0.48 |
| AT5G36730.1 | F-box and associated interaction domains-containing protein | - | Sampling day | -0.20 |
| AT1G35146.1 | transposable element gene | - | Sampling day | -0.15 |
| AT2G36610.1 | homeobox protein 22 | HB22 | Sampling day | -0.22 |
| AT5G43790.1 | Pentatricopeptide repeat (PPR) superfamily protein | - | Sampling day | -0.07 |
| AT2G35350.1 | poltergeist like 1 | PLL1 | Sampling day | -0.20 |
| AT5G01990.1 | Auxin efflux carrier family protein | - | Sampling day | -0.22 |
| AT1G64583.2 | Tetratricopeptide repeat (TPR)-like superfamily protein | - | Sampling day | -0.32 |
| AT5G59600.1 | Tetratricopeptide repeat (TPR)-like superfamily protein | - | Sampling day | -0.17 |
| AT1G27490.1 | F-box and associated interaction domains-containing protein | - | Sampling day | -0.13 |
| AT1G66770.1 | Nodulin MtN3 family protein | SWEET6 | Sampling day | -0.13 |
| AT4G03640.1 | transposable element gene | - | Sampling day | -0.11 |
| AT1G06287.1 | | - | Sampling day | -0.18 |
| AT5G39930.1 | CLP1-similar protein 5 | CLPS5 | Sampling day | -0.19 |
| AT5G23690.1 | Polynucleotide adenylyltransferase family protein | - | Sampling day | -0.07 |
| AT3G56750.1 | NULL | - | Sampling day | -0.18 |
| AT5G52250.1 | Transducin/WD40 repeat-like superfamily protein | RUP1 | Sampling day | -0.10 |
| AT3G52285.1 | pre-tRNA | - | Sampling day | -0.23 |
| AT1G57370.1 | pre-tRNA | - | Sampling day | -0.15 |
| AT1G61760.1 | Late embryogenesis abundant (LEA) hydroxyproline-rich glycoprotein | - | Sampling day | -0.08 |
| ATCG00740.1 | RNA polymerase subunit alpha | RPOA | Sampling day | -0.11 |
| AT1G63820.1 | CCT motif family protein | - | Sampling day | -0.09 |
| AT2G18070.1 | NULL | - | Sampling day | -0.12 |
| AT2G08110.1 | | - | Sampling day | -0.23 |
| AT1G08125.2 | S-adenosyl-L-methionine-dependent methyltransferases superfamily | - | Sampling day | -0.22 |
| AT1G47657.1 | | - | Sampling day | -0.11 |
| AT1G30570.1 | hercules receptor kinase 2 | HERK2 | Sampling day | -0.20 |
| AT3G30846.1 | transposable element gene | - | Sampling day | -0.20 |
| AT2G09150.1 | | - | Sampling day | -0.12 |
| AT2G13690.1 | PRLI-interacting factor, putative | - | Sampling day | -0.27 |
| AT2G47270.1 | sequence-specific DNA binding transcription factors;transcription | UPB1 | Sampling day | -0.18 |
| AT1G04990.3 | Zinc finger C-x8-C-x5-C-x3-H type family protein | - | Sampling day | -0.17 |
| AT1G48440.1 | B-cell receptor-associated 31-like | - | Sampling day | -0.27 |
| AT5G16080.1 | carboxyesterase 17 | CXE17 | Sampling day | -0.16 |
| AT3G08370.1 | | - | Sampling day | -0.09 |
| AT5G05665.1 | | - | Sampling day | -0.17 |
| AT2G33780.1 | VQ motif-containing protein | - | Sampling day | -0.41 |
| AT2G36580.1 | Pyruvate kinase family protein | - | Sampling day | -0.05 |

| | | | | |
|---|---|---|---|---|
| AT1G72500.2 | NULL | - | Sampling day | -0.05 |
| AT4G14840.1 | NULL | - | Sampling day | -0.12 |
| AT3G02955.1 | | - | Sampling day | -0.22 |
| AT5G35650.1 | transposable element gene | - | Sampling day | -0.15 |
| AT3G49760.1 | basic leucine-zipper 5 | bZIP5 | Sampling day | -0.32 |
| AT3G02510.2 | Regulator of chromosome condensation (RCC1) family protein | - | Sampling day | -0.09 |
| AT1G78950.2 | Terpenoid cyclases family protein | BAS | Sampling day | -0.12 |
| AT3G05970.1 | long-chain acyl-CoA synthetase 6 | LACS6 | Sampling day | -0.26 |
| AT1G10455.1 | NULL | - | Sampling day | -0.12 |
| AT1G62790.1 | Bifunctional inhibitor/lipid-transfer protein/seed storage 2S albumin superfamily protein | - | Sampling day | -0.23 |
| AT5G58930.1 | Protein of unknown function (DUF740) | - | Sampling day | -0.21 |
| AT1G15810.1 | S15/NS1, RNA-binding protein | - | Sampling day | -0.15 |
| AT3G53720.2 | cation/H+ exchanger 20 | CHX20 | Sampling day | -0.08 |
| AT5G04025.1 | | - | Sampling day | -0.16 |
| AT5G25750.1 | NULL | - | Sampling day | -0.67 |
| AT5G35920.1 | cytochrome P450, family 79, subfamily A, polypeptide 4 pseudogene | CYP79A4P | Sampling day | -0.23 |
| AT1G05823.1 | | - | Sampling day | -0.25 |
| AT2G04940.1 | scramblase-related | - | Sampling day | -0.20 |
| AT4G13245.1 | snoRNA | - | Sampling day | -0.23 |
| AT1G50530.1 | NULL | - | Sampling day | -0.13 |
| AT5G51050.1 | Mitochondrial substrate carrier family protein | APC2 | Sampling day | -0.21 |
| AT5G09670.1 | loricrin-related | - | Sampling day | -0.27 |
| AT5G50690.2 | hydroxysteroid dehydrogenase 7 | HSD7 | Sampling day | -0.17 |
| AT4G18750.1 | Pentatricopeptide repeat (PPR) superfamily protein | DOT4 | Sampling day | -0.06 |
| AT1G16410.1 | cytochrome p450 79f1 | CYP79F1 | Sampling day | -0.06 |
| AT5G45740.1 | Ubiquitin domain-containing protein | - | Sampling day | -0.22 |
| AT5G60410.1 | DNA-binding protein with MIZ/SP-RING zinc finger, PHD-finger and SAP domain | SIZ1 | Sampling day | -0.15 |
| AT5G48650.1 | Nuclear transport factor 2 (NTF2) family protein with RNA binding (RRM-RBD-RNP motifs) domain | - | Sampling day | -0.11 |
| AT5G40390.1 | Raffinose synthase family protein | SIP1 | Sampling day | -0.21 |
| AT4G04110.1 | Toll-Interleukin-Resistance (TIR) domain family protein | - | Sampling day | -0.08 |
| AT2G42470.1 | TRAF-like family protein | - | Sampling day | -0.11 |
| AT2G46150.1 | Late embryogenesis abundant (LEA) hydroxyproline-rich glycoprotein | - | Sampling day | -0.15 |
| AT2G39170.1 | NULL | - | Sampling day | -0.14 |
| AT4G19310.1 | transposable element gene | - | Sampling day | -0.07 |
| AT2G46950.2 | cytochrome P450, family 709, subfamily B, polypeptide 2 | CYP709B2 | Sampling day | -0.17 |
| AT5G51520.1 | Plant invertase/pectin methylesterase inhibitor superfamily protein | - | Sampling day | -0.08 |
| AT5G34686.1 | transposable element gene | - | Sampling day | -0.09 |
| AT5G11325.1 | pre-tRNA | - | Sampling day | -0.28 |
| AT4G35130.1 | Tetratricopeptide repeat (TPR)-like superfamily protein | - | Sampling day | -0.22 |
| AT2G00370.1 | | - | Sampling day | -0.12 |
| AT4G08033.1 | transposable element gene | - | Sampling day | -0.17 |
| AT4G09295.1 | | - | Sampling day | -0.42 |
| AT2G10205.1 | transposable element gene | - | Sampling day | -0.33 |
| AT1G02140.1 | mago nashi family protein | MAGO | Sampling day | -0.27 |
| AT5G07925.1 | | - | Sampling day | -0.32 |
| AT5G46590.1 | NAC domain containing protein 96 | NAC096 | Sampling day | -0.21 |
| AT5G11790.2 | N-MYC downregulated-like 2 | NDL2 | Sampling day | -0.07 |
| AT3G15604.1 | NULL | - | Sampling day | -0.08 |
| AT4G17900.2 | PLATZ transcription factor family protein | - | Sampling day | -0.16 |
| AT5G19675.1 | | - | Sampling day | -0.45 |
| AT5G04690.2 | Ankyrin repeat family protein | - | Sampling day | -0.09 |
| AT2G01031.1 | transposable element gene | - | Sampling day | -0.06 |
| AT5G63290.1 | Radical SAM superfamily protein | - | Sampling day | -0.28 |
| AT4G12010.1 | Disease resistance protein (TIR-NBS-LRR class) family | - | Sampling day | -0.31 |
| AT5G23940.1 | HXXXD-type acyl-transferase family protein | PEL3 | Sampling day | -0.06 |
| AT5G22950.1 | SNF7 family protein | VPS24.1 | Sampling day | -0.21 |
| AT1G71870.1 | MATE efflux family protein | - | Sampling day | -0.22 |
| AT1G07483.1 | | - | Sampling day | -0.34 |
| AT5G54380.1 | protein kinase family protein | THE1 | Sampling day | -0.07 |
| AT3G47820.1 | PLANT U-BOX 39 | PUB39 | Sampling day | -0.19 |
| AT1G23070.1 | Protein of unknown function (DUF300) | - | Sampling day | -0.49 |
| AT3G27660.1 | oleosin 4 | OLEO4 | Sampling day | -0.16 |
| AT4G02405.2 | S-adenosyl-L-methionine-dependent methyltransferases superfamily | - | Sampling day | -0.10 |
| AT4G12670.2 | Homeodomain-like superfamily protein | - | Sampling day | -0.39 |
| AT3G22110.1 | 20S proteasome alpha subunit C1 | PAC1 | Sampling day | -0.08 |
| AT3G46487.1 | transposable element gene | - | Sampling day | -0.30 |
| AT1G49080.1 | transposable element gene | - | Sampling day | -0.11 |
| AT3G16340.1 | pleiotropic drug resistance 1 | ABCG29 | Sampling day | -0.16 |
| AT1G19140.1 | NULL | - | Sampling day | -0.20 |
| AT3G51080.1 | GATA transcription factor 6 | GATA6 | Sampling day | -0.40 |
| AT5G15010.1 | Tetratricopeptide repeat (TPR)-like superfamily protein | - | Sampling day | -0.21 |
| AT1G07673.1 | | - | Sampling day | -0.17 |
| AT3G24420.1 | alpha/beta-Hydrolases superfamily protein | - | Sampling day | -0.18 |

| | | | | |
|---|---|---|---|---|
| AT3G55935.1 | | - | Sampling day | -0.17 |
| AT1G53220.1 | pre-tRNA | - | Sampling day | -0.07 |
| AT1G12400.3 | Nucleotide excision repair, TFIIH, subunit TTDA | - | Sampling day | -0.14 |
| AT5G19170.1 | Protein of Unknown Function (DUF239) | - | Sampling day | -0.25 |
| AT2G08975.1 | | - | Sampling day | -0.18 |
| AT1G48760.2 | delta-adaptin | delta-ADR | Sampling day | -0.14 |
| AT2G20480.1 | NULL | - | Sampling day | -0.18 |
| AT1G68875.1 | NULL | - | Sampling day | -0.10 |
| AT4G06684.1 | transposable element gene | - | Sampling day | -0.14 |
| AT1G22770.1 | gigantea protein (GI) | GI | Sampling day | -0.09 |
| ATMG00070.1 | NADH dehydrogenase subunit 9 | NAD9 | Sampling day | -0.20 |
| AT2G47380.2 | Cytochrome c oxidase subunit Vc family protein | - | Sampling day | -0.17 |
| AT5G53420.5 | CCT motif family protein | - | Sampling day | -0.17 |
| AT4G39235.1 | NULL | - | Sampling day | -0.11 |
| AT1G09207.1 | | - | Sampling day | -0.12 |
| AT5G26740.2 | Protein of unknown function (DUF300) | - | Sampling day | -0.18 |
| AT2G29560.1 | cytosolic enolase | ENOC | Sampling day | -0.18 |
| AT1G60987.1 | SCR-like 5 | SCRL5 | Sampling day | -0.21 |
| AT3G06460.1 | GNS1/SUR4 membrane protein family | - | Sampling day | -0.19 |
| AT2G33775.1 | ralf-like 19 | RALFL19 | Sampling day | -0.44 |
| AT1G09035.1 | | - | Sampling day | -0.17 |
| AT1G35570.1 | transposable element gene | - | Sampling day | -0.26 |
| AT3G43681.1 | transposable element gene | - | Sampling day | -0.24 |
| AT3G08640.1 | Protein of unknown function (DUF3411) | - | Sampling day | -0.23 |
| AT4G20645.1 | | - | Sampling day | -0.13 |
| AT3G31560.1 | | - | Sampling day | -0.14 |
| AT1G22920.1 | COP9 signalosome 5A | CSN5A | Sampling day | -0.21 |
| AT2G09125.1 | | - | Sampling day | -0.17 |
| AT3G47040.3 | Glycosyl hydrolase family protein | - | Sampling day | -0.07 |
| AT3G42203.1 | transposable element gene | - | Sampling day | -0.16 |
| AT3G21590.2 | Senescence/dehydration-associated protein-related | - | Sampling day | -0.09 |
| AT1G04507.1 | | - | Sampling day | -0.21 |
| AT4G36925.2 | NULL | - | Sampling day | -0.17 |
| AT1G48470.1 | glutamine synthetase 1;5 | GLN1%3B5 | Sampling day | -0.20 |
| ATMG00516.1 | NADH dehydrogenase 1C | NAD1C | Sampling day | -0.25 |
| AT1G07230.1 | non-specific phospholipase C1 | NPC1 | Sampling day | -0.28 |
| AT2G40020.2 | Nucleolar histone methyltransferase-related protein | - | Sampling day | -0.09 |
| AT3G13662.1 | Disease resistance-responsive (dirigent-like protein) family protein | - | Sampling day | -0.23 |
| AT3G44330.1 | NULL | - | Sampling day | -0.05 |
| AT3G04810.3 | NIMA-related kinase 2 | NEK2 | Sampling day | -0.07 |
| AT1G75330.1 | ornithine carbamoyltransferase | OTC | Sampling day | -0.09 |
| AT2G23390.2 | NULL | - | Sampling day | -0.22 |
| AT1G74050.1 | Ribosomal protein L6 family protein | - | Sampling day | -0.05 |
| AT1G32760.1 | Glutaredoxin family protein | - | Sampling day | -0.19 |
| AT3G00540.1 | | - | Sampling day | -0.24 |
| AT4G20150.1 | NULL | - | Sampling day | -0.14 |
| AT3G49410.2 | Transcription factor IIIC, subunit 5 | - | Sampling day | -0.28 |
| AT2G37290.2 | Ypt/Rab-GAP domain of gyp1p superfamily protein | - | Sampling day | -0.27 |
| AT1G08547.1 | | - | Sampling day | -0.26 |
| AT1G28870.1 | pre-tRNA | - | Sampling day | -0.28 |
| AT3G44117.1 | NULL | - | Sampling day | -0.16 |
| AT4G00780.1 | TRAF-like family protein | - | Sampling day | -0.21 |
| AT4G35810.2 | 2-oxoglutarate (2OG) and Fe(II)-dependent oxygenase superfamily | - | Sampling day | -0.21 |
| AT5G05870.1 | UDP-glucosyl transferase 76C1 | UGT76C1 | Sampling day | -0.15 |
| AT4G03113.1 | NULL | - | Sampling day | -0.23 |
| AT1G63570.2 | Receptor-like protein kinase-related family protein | - | Sampling day | -0.24 |
| AT2G19825.1 | transposable element gene | - | Sampling day | -0.27 |
| AT1G37035.1 | transposable element gene | - | Sampling day | -0.18 |
| AT5G36260.1 | Eukaryotic aspartyl protease family protein | - | Sampling day | -0.19 |
| AT4G33160.2 | F-box family protein | - | Sampling day | -0.30 |
| AT1G65870.1 | Disease resistance-responsive (dirigent-like protein) family protein | - | Sampling day | -0.10 |
| AT4G00270.1 | DNA-binding storekeeper protein-related transcriptional regulator | - | Sampling day | -0.13 |
| AT5G62880.2 | RAC-like 10 | RAC10 | Sampling day | -0.18 |
| AT1G55205.1 | NULL | - | Sampling day | -0.42 |
| AT1G23201.2 | NULL | - | Sampling day | -0.18 |
| AT5G18450.1 | Integrase-type DNA-binding superfamily protein | - | Sampling day | -0.17 |
| AT1G42700.1 | NULL | - | Sampling day | -0.16 |
| AT1G21310.1 | extensin 3 | EXT3 | Sampling day | -0.20 |
| AT2G36540.1 | Haloacid dehalogenase-like hydrolase (HAD) superfamily protein | - | Sampling day | -0.15 |
| AT1G45215.1 | Protein of unknown function (DUF784) | - | Sampling day | -0.23 |
| AT1G05103.1 | | - | Sampling day | -0.06 |
| AT3G15360.1 | thioredoxin M-type 4 | TRX-M4 | Sampling day | -0.06 |
| AT1G09140.3 | SERINE-ARGININE PROTEIN 30 | SR30 | Sampling day | -0.07 |
| AT4G13560.1 | Late embryogenesis abundant protein (LEA) family protein | UNE15 | Sampling day | -0.11 |
| AT3G03000.1 | EF hand calcium-binding protein family | - | Sampling day | -0.21 |
| AT1G22970.2 | NULL | - | Sampling day | -0.29 |

| | | | | |
|---|---|---|---|---|
| AT5G67320.1 | WD-40 repeat family protein | HOS15 | Sampling day | -0.18 |
| AT2G02710.4 | PAS/LOV protein B | PLPB | Sampling day | -0.10 |
| AT3G59030.1 | MATE efflux family protein | TT12 | Sampling day | -0.20 |
| AT1G06613.1 | | - | Sampling day | -0.25 |
| AT1G19740.1 | ATP-dependent protease La (LON) domain protein | - | Sampling day | -0.18 |
| AT2G26480.1 | UDP-glucosyl transferase 76D1 | UGT76D1 | Sampling day | -0.11 |
| AT1G24010.1 | Polyketide cyclase/dehydrase and lipid transport superfamily protein | - | Sampling day | -0.15 |
| AT5G06940.1 | Leucine-rich repeat receptor-like protein kinase family protein | - | Sampling day | -0.21 |
| AT1G56300.1 | Chaperone DnaJ-domain superfamily protein | - | Sampling day | -0.06 |
| AT5G65730.1 | xyloglucan endotransglucosylase/hydrolase 6 | XTH6 | Sampling day | -0.06 |
| AT1G75070.1 | pre-tRNA | - | Sampling day | -0.16 |
| AT3G60760.1 | NULL | - | Sampling day | -0.15 |
| AT1G08660.2 | MALE GAMETOPHYTE DEFECTIVE 2 | MGP2 | Sampling day | -0.06 |
| AT2G02650.1 | Ribonuclease H-like superfamily protein | - | Sampling day | -0.21 |
| AT1G05627.1 | | - | Sampling day | -0.29 |
| AT4G27990.1 | YGGT family protein | YLMG1-2 | Sampling day | -0.23 |
| AT4G36860.2 | LIM domain-containing protein | - | Sampling day | -0.08 |
| AT3G47470.1 | light-harvesting chlorophyll-protein complex I subunit A4 | LHCA4 | Sampling day | -0.17 |
| AT2G14610.1 | pathogenesis-related gene 1 | PR1 | Sampling day | -0.11 |
| AT4G37410.1 | cytochrome P450, family 81, subfamily F, polypeptide 4 | CYP81F4 | Sampling day | -0.18 |
| AT4G24150.3 | growth-regulating factor 8 | GRF8 | Sampling day | -0.23 |
| AT3G06575.1 | | - | Sampling day | -0.17 |
| AT5G06260.1 | TLD-domain containing nucleolar protein | - | Sampling day | -0.21 |
| AT3G15980.4 | Coatomer, beta' subunit | - | Sampling day | -0.14 |
| AT4G18700.1 | CBL-interacting protein kinase 12 | CIPK12 | Sampling day | -0.09 |
| AT1G17410.3 | Nucleoside diphosphate kinase family protein | - | Sampling day | -0.27 |
| AT5G07175.1 | | - | Sampling day | -0.08 |
| AT5G50915.4 | basic helix-loop-helix (bHLH) DNA-binding superfamily protein | - | Sampling day | -0.14 |
| AT5G49760.1 | Leucine-rich repeat protein kinase family protein | - | Sampling day | -0.19 |
| AT4G20600.1 | Protein with domains of unknown function (DUF26 and DUF1204) | - | Sampling day | -0.36 |
| AT3G17520.1 | Late embryogenesis abundant protein (LEA) family protein | - | Sampling day | -0.16 |
| AT4G37650.1 | GRAS family transcription factor | SHR | Sampling day | -0.06 |
| AT3G42622.1 | transposable element gene | - | Sampling day | -0.18 |
| AT3G33133.1 | transposable element gene | - | Sampling day | -0.25 |
| ATMG09720.1 | | - | Sampling day | -0.13 |
| AT5G59830.3 | NULL | - | Sampling day | -0.29 |
| AT1G14010.1 | emp24/gp25L/p24 family/GOLD family protein | - | Sampling day | -0.16 |
| AT3G06950.1 | Pseudouridine synthase family protein | - | Sampling day | -0.18 |
| AT1G26300.2 | BSD domain-containing protein | - | Sampling day | -0.08 |
| AT5G35340.1 | transposable element gene | - | Sampling day | -0.19 |
| AT1G04487.1 | | - | Sampling day | -0.13 |
| AT2G13965.1 | | - | Sampling day | -0.12 |
| AT3G08200.1 | | - | Sampling day | -0.18 |
| AT3G47780.1 | ABC2 homolog 6 | ABCA7 | Sampling day | -0.20 |
| AT2G32273.1 | MIR417 (MICRORNA417); miRNA | MIR417 | Sampling day | -0.24 |
| AT2G43450.1 | NULL | - | Sampling day | -0.09 |
| AT2G24255.1 | NULL | - | Sampling day | -0.10 |
| AT2G25240.1 | Serine protease inhibitor (SERPIN) family protein | CCP3 | Sampling day | -0.21 |
| AT1G01530.1 | AGAMOUS-like 28 | AGL28 | Sampling day | -0.07 |
| AT4G00040.1 | Chalcone and stilbene synthase family protein | - | Sampling day | -0.12 |
| AT4G06485.1 | transposable element gene | - | Sampling day | -0.14 |
| AT1G24000.1 | Polyketide cyclase/dehydrase and lipid transport superfamily protein | - | Sampling day | -0.15 |
| AT5G03200.1 | RING/U-box superfamily protein | LUL1 | Sampling day | -0.16 |
| AT3G30751.1 | transposable element gene | - | Sampling day | -0.15 |
| AT5G54070.1 | heat shock transcription factor A9 | HSFA9 | Sampling day | -0.09 |
| AT3G20840.1 | Integrase-type DNA-binding superfamily protein | PLT1 | Sampling day | -0.09 |
| AT2G01818.1 | PLATZ transcription factor family protein | - | Sampling day | -0.17 |
| AT5G14260.1 | Rubisco methyltransferase family protein | - | Sampling day | -0.10 |
| AT2G45970.1 | cytochrome P450, family 86, subfamily A, polypeptide 8 | CYP86A8 | Sampling day | -0.09 |
| AT1G63540.1 | hydroxyproline-rich glycoprotein family protein | - | Sampling day | -0.15 |
| AT1G47570.1 | RING/U-box superfamily protein | - | Sampling day | -0.07 |
| AT5G32267.1 | transposable element gene | - | Sampling day | -0.14 |
| AT1G11265.1 | transposable element gene | - | Sampling day | -0.25 |
| AT3G22010.1 | Receptor-like protein kinase-related family protein | - | Sampling day | -0.22 |
| AT1G14518.1 | other RNA | - | Sampling day | -0.11 |
| AT3G42656.1 | transposable element gene | - | Sampling day | -0.18 |
| AT3G15900.1 | NULL | - | Sampling day | -0.12 |
| AT1G04883.1 | | - | Sampling day | -0.11 |
| AT3G29618.1 | transposable element gene | - | Sampling day | -0.17 |
| AT4G07840.1 | transposable element gene | - | Sampling day | -0.54 |
| AT3G53520.1 | UDP-glucuronic acid decarboxylase 1 | UXS1 | Sampling day | -0.13 |
| AT5G53900.1 | Serine/threonine-protein kinase WNK (With No Lysine)-related | - | Sampling day | -0.19 |
| AT2G42455.1 | | - | Sampling day | -0.09 |
| AT1G15050.1 | indole-3-acetic acid inducible 34 | IAA34 | Sampling day | -0.13 |
| AT5G37170.2 | O-methyltransferase family protein | - | Sampling day | -0.27 |
| AT5G26270.2 | NULL | - | Sampling day | -0.07 |

| | | | | |
|---|---|---|---|---|
| AT3G48490.1 | NULL | - | Sampling day | -0.44 |
| AT5G33624.1 | transposable element gene | - | Sampling day | -0.19 |
| AT3G25880.1 | NAD(P)-binding Rossmann-fold superfamily protein | - | Sampling day | -0.11 |
| AT3G11810.1 | NULL | - | Sampling day | -0.15 |
| AT3G29105.1 | NULL | - | Sampling day | -0.14 |
| AT4G06654.1 | transposable element gene | - | Sampling day | -0.27 |
| AT2G24681.1 | AP2/B3-like transcriptional factor family protein | - | Sampling day | -0.19 |
| AT5G58510.1 | NULL | - | Sampling day | -0.33 |
| AT1G72030.1 | Acyl-CoA N-acyltransferases (NAT) superfamily protein | - | Sampling day | -0.27 |
| AT5G53270.1 | Seed maturation protein | - | Sampling day | -0.09 |
| AT3G58410.1 | TRAF-like family protein | - | Sampling day | -0.17 |
| AT1G52850.1 | transposable element gene | - | Sampling day | -0.07 |
| AT1G14300.2 | ARM repeat superfamily protein | - | Sampling day | -0.10 |
| AT1G24580.1 | RING/U-box superfamily protein | - | Sampling day | -0.11 |
| AT3G57200.2 | NULL | - | Sampling day | -0.19 |
| AT2G34480.2 | Ribosomal protein L18ae/LX family protein | - | Sampling day | -0.23 |
| AT3G13500.1 | NULL | - | Sampling day | -0.21 |
| AT1G44160.1 | HSP40/DnaJ peptide-binding protein | - | Sampling day | -0.12 |
| AT2G45560.1 | cytochrome P450, family 76, subfamily C, polypeptide 1 | CYP76C1 | Sampling day | -0.16 |
| AT3G15610.1 | Transducin/WD40 repeat-like superfamily protein | - | Sampling day | -0.24 |
| AT5G07385.1 | | - | Sampling day | -0.10 |
| AT5G63160.3 | BTB and TAZ domain protein 1 | BT1 | Sampling day | -0.10 |
| AT5G18300.1 | NAC domain containing protein 88 | NAC088 | Sampling day | -0.21 |
| AT5G33370.1 | GDSL-like Lipase/Acylhydrolase superfamily protein | - | Sampling day | -0.12 |
| AT4G16480.1 | inositol transporter 4 | INT4 | Sampling day | -0.25 |
| AT1G65470.1 | chromatin assembly factor-1 (FASCIATA1) (FAS1) | FAS1 | Sampling day | -0.17 |
| AT5G62740.1 | SPFH/Band 7/PHB domain-containing membrane-associated protein | HIR1 | Sampling day | -0.18 |
| AT5G54830.1 | DOMON domain-containing protein / dopamine beta-monooxygenase N-terminal domain-containing protein | - | Sampling day | -0.21 |
| AT5G32228.1 | transposable element gene | - | Sampling day | -0.16 |
| AT2G09850.1 | transposable element gene | - | Sampling day | -0.19 |
| AT1G13470.1 | Protein of unknown function (DUF1262) | - | Sampling day | -0.17 |
| AT2G03934.1 | NULL | - | Sampling day | -0.08 |
| AT5G25450.1 | Cytochrome bd ubiquinol oxidase, 14kDa subunit | - | Sampling day | -0.19 |
| AT5G37473.1 | Defensin-like (DEFL) family protein | - | Sampling day | -0.19 |
| AT4G21170.1 | Tetratricopeptide repeat (TPR)-like superfamily protein | - | Sampling day | -0.38 |
| AT3G58230.1 | Ubiquitin-specific protease family C19-related protein | - | Sampling day | -0.10 |
| AT1G69000.1 | pre-tRNA | - | Sampling day | -0.20 |
| AT1G53470.1 | mechanosensitive channel of small conductance-like 4 | MSL4 | Sampling day | -0.27 |
| AT3G33075.1 | transposable element gene | - | Sampling day | -0.08 |
| AT3G08835.1 | | - | Sampling day | -0.09 |
| AT5G52045.1 | | - | Sampling day | -0.14 |
| AT3G08545.1 | | - | Sampling day | -0.17 |
| AT5G24800.1 | basic leucine zipper 9 | BZIP9 | Sampling day | -0.23 |
| AT2G18890.3 | Protein kinase superfamily protein | - | Sampling day | -0.06 |
| AT1G20400.1 | Protein of unknown function (DUF1204) | - | Sampling day | -0.23 |
| AT1G26540.1 | Agenet domain-containing protein | - | Sampling day | -0.23 |
| AT3G24093.1 | Paired amphipathic helix (PAH2) superfamily protein | - | Sampling day | -0.22 |
| AT5G07520.1 | glycine-rich protein 18 | GRP18 | Sampling day | -0.16 |
| AT3G22053.1 | NULL | - | Sampling day | -0.12 |
| AT1G46554.1 | other RNA | - | Sampling day | -0.20 |
| AT3G17765.1 | | - | Sampling day | -0.09 |
| AT5G15070.3 | Phosphoglycerate mutase-like family protein | - | Sampling day | -0.11 |
| AT5G50640.1 | CBS / octicosapeptide/Phox/Bemp1 (PB1) domains-containing protein | - | Sampling day | -0.21 |
| AT1G09215.1 | | - | Sampling day | -0.14 |
| AT5G44283.1 | pre-tRNA | - | Sampling day | -0.11 |
| AT3G60980.1 | Tetratricopeptide repeat (TPR)-like superfamily protein | - | Sampling day | -0.10 |
| AT3G46240.1 | NULL | - | Sampling day | -0.11 |
| AT1G61200.1 | homeobox-leucine zipper protein-related | - | Sampling day | -0.14 |
| AT5G15980.1 | Pentatricopeptide repeat (PPR) superfamily protein | - | Sampling day | -0.22 |
| AT1G71720.1 | Nucleic acid-binding proteins superfamily | PDE338 | Sampling day | -0.09 |
| AT1G33980.1 | Smg-4/UPF3 family protein | UPF3 | Sampling day | -0.16 |
| AT2G22780.1 | peroxisomal NAD-malate dehydrogenase 1 | PMDH1 | Sampling day | -0.19 |
| AT5G07505.1 | transposable element gene | - | Sampling day | -0.08 |
| AT5G50770.1 | hydroxysteroid dehydrogenase 6 | HSD6 | Sampling day | -0.21 |
| AT4G07810.1 | transposable element gene | - | Sampling day | -0.11 |
| AT3G54160.1 | RNI-like superfamily protein | - | Sampling day | -0.20 |
| AT1G72050.4 | transcription factor IIIA | TFIIIA | Sampling day | -0.11 |
| AT1G77520.1 | O-methyltransferase family protein | - | Sampling day | -0.22 |
| AT1G51402.1 | NULL | - | Sampling day | -0.05 |
| AT5G35900.1 | LOB domain-containing protein 35 | LBD35 | Sampling day | -0.14 |
| AT1G47630.1 | NULL | CYP96A7 | Sampling day | -0.22 |
| AT3G11964.2 | RNA binding;RNA binding | - | Sampling day | -0.23 |
| AT4G18450.1 | Integrase-type DNA-binding superfamily protein | - | Sampling day | -0.11 |
| AT1G12672.2 | NULL | - | Sampling day | -0.24 |
| AT2G21910.1 | cytochrome P450, family 96, subfamily A, polypeptide 5 | CYP96A5 | Sampling day | -0.18 |

| | | | | |
|---|---|---|---|---|
| AT5G04480.1 | UDP-Glycosyltransferase superfamily protein | - | Sampling day | -0.21 |
| AT2G28420.1 | Lactoylglutathione lyase / glyoxalase I family protein | GLYI8 | Sampling day | -0.10 |
| AT1G07873.1 | | - | Sampling day | -0.13 |
| AT4G34020.1 | Class I glutamine amidotransferase-like superfamily protein | DJ1C | Sampling day | -0.23 |
| AT3G61750.2 | Cytochrome b561/ferric reductase transmembrane with DOMON related domain | - | Sampling day | -0.21 |
| AT3G30370.1 | NULL | - | Sampling day | -0.14 |
| AT4G06699.1 | transposable element gene | - | Sampling day | -0.12 |
| AT3G04090.1 | small and basic intrinsic protein 1A | SIP1A | Sampling day | -0.06 |
| AT1G58090.1 | F-box and associated interaction domains-containing protein | - | Sampling day | -0.16 |
| AT5G07270.1 | XB3 ortholog 3 in Arabidopsis thaliana | XBAT33 | Sampling day | -0.15 |
| AT5G02315.1 | | - | Sampling day | -0.26 |
| AT3G43610.2 | Spc97 / Spc98 family of spindle pole body (SBP) component | - | Sampling day | -0.07 |
| AT4G36945.1 | PLC-like phosphodiesterases superfamily protein | - | Sampling day | -0.23 |
| AT3G01320.1 | SIN3-like 1 | SNL1 | Sampling day | -0.09 |
| AT2G08385.1 | | - | Sampling day | -0.39 |
| AT5G05835.1 | | - | Sampling day | -0.07 |
| AT3G02985.1 | | - | Sampling day | -0.15 |
| AT2G09910.1 | transposable element gene | - | Sampling day | -0.22 |
| AT1G14800.1 | Nucleic acid-binding, OB-fold-like protein | - | Sampling day | -0.13 |
| AT3G44380.1 | Late embryogenesis abundant (LEA) hydroxyproline-rich glycoprotein | - | Sampling day | -0.11 |
| AT1G12930.1 | ARM repeat superfamily protein | - | Sampling day | -0.43 |
| AT5G43770.1 | proline-rich family protein | - | Sampling day | -0.08 |
| AT5G10450.3 | G-box regulating factor 6 | GRF6 | Sampling day | -0.17 |
| AT3G42313.1 | transposable element gene | - | Sampling day | -0.12 |
| AT2G31957.1 | low-molecular-weight cysteine-rich 75 | LCR75 | Sampling day | -0.06 |
| AT4G39150.3 | DNAJ heat shock N-terminal domain-containing protein | - | Sampling day | -0.22 |
| AT5G20820.1 | SAUR-like auxin-responsive protein family | - | Sampling day | -0.08 |
| AT2G05280.1 | transposable element gene | - | Sampling day | -0.14 |
| AT2G01470.1 | SEC12P-like 2 protein | STL2P | Sampling day | -0.13 |
| AT2G39375.1 | | - | Sampling day | -0.14 |
| AT2G00600.1 | | - | Sampling day | -0.10 |
| AT4G21540.1 | sphingosine kinase 1 | SPHK1 | Sampling day | -0.07 |
| AT1G04293.1 | | - | Sampling day | -0.22 |
| AT5G06230.1 | TRICHOME BIREFRINGENCE-LIKE 9 | TBL9 | Sampling day | -0.23 |
| AT4G02100.1 | Heat shock protein DnaJ with tetratricopeptide repeat | - | Sampling day | -0.09 |
| AT3G08330.1 | | - | Sampling day | -0.17 |
| AT4G13572.1 | NULL | - | Sampling day | -0.05 |
| AT2G13463.1 | NULL | - | Sampling day | -0.14 |
| AT4G06015.1 | | - | Sampling day | -0.19 |
| AT1G18460.1 | alpha/beta-Hydrolases superfamily protein | - | Sampling day | -0.18 |
| AT3G05925.1 | | - | Sampling day | -0.09 |
| AT2G02061.1 | Nucleotide-diphospho-sugar transferase family protein | - | Sampling day | -0.16 |
| AT1G27100.1 | Actin cross-linking protein | - | Sampling day | -0.16 |
| AT3G05930.1 | germin-like protein 8 | GLP8 | Sampling day | -0.35 |
| AT1G31040.1 | PLATZ transcription factor family protein | - | Sampling day | -0.09 |
| AT4G24020.1 | NIN like protein 7 | NLP7 | Sampling day | -0.20 |
| AT4G05000.3 | Vacuolar protein sorting-associated protein VPS28 family protein | VPS28-2 | Sampling day | -0.08 |
| AT2G21920.1 | F-box associated ubiquitination effector family protein | - | Sampling day | -0.18 |
| AT5G66790.1 | Protein kinase superfamily protein | - | Sampling day | -0.07 |
| AT3G15570.1 | Phototropic-responsive NPH3 family protein | - | Sampling day | -0.12 |
| AT1G05077.1 | | - | Sampling day | -0.10 |
| AT5G61450.1 | P-loop containing nucleoside triphosphate hydrolases superfamily | - | Sampling day | -0.16 |
| AT4G39345.1 | pre-tRNA | - | Sampling day | -0.07 |
| AT3G18790.1 | NULL | - | Sampling day | -0.19 |
| AT1G08057.1 | | - | Sampling day | -0.38 |
| AT4G05800.1 | | - | Sampling day | -0.09 |
| AT2G10600.1 | transposable element gene | - | Sampling day | -0.06 |
| AT5G28482.1 | transposable element gene | - | Sampling day | -0.14 |
| AT2G03940.1 | transposable element gene | - | Sampling day | -0.22 |
| AT5G40620.1 | NULL | - | Sampling day | -0.23 |
| AT1G71250.1 | GDSL-like Lipase/Acylhydrolase superfamily protein | - | Sampling day | -0.60 |
| AT3G30520.1 | NULL | - | Sampling day | -0.61 |
| AT3G33004.1 | NULL | - | Sampling day | -0.21 |
| AT3G24350.1 | syntaxin of plants 32 | SYP32 | Sampling day | -0.19 |
| AT2G15250.1 | transposable element gene | - | Sampling day | -0.20 |
| AT3G28720.2 | NULL | - | Sampling day | -0.22 |
| AT2G29940.1 | pleiotropic drug resistance 3 | ABCG31 | Sampling day | -0.13 |
| AT5G39750.1 | AGAMOUS-like 81 | AGL81 | Sampling day | -0.33 |
| AT1G51890.2 | Leucine-rich repeat protein kinase family protein | - | Sampling day | -0.16 |
| AT1G02640.1 | beta-xylosidase 2 | BXL2 | Sampling day | -0.08 |
| AT3G22200.2 | Pyridoxal phosphate (PLP)-dependent transferases superfamily protein | POP2 | Sampling day | -0.06 |
| AT5G44280.1 | RING 1A | RING1A | Sampling day | -0.23 |
| AT4G36020.2 | cold shock domain protein 1 | CSDP1 | Sampling day | -0.09 |
| AT2G27600.1 | AAA-type ATPase family protein | SKD1 | Sampling day | -0.23 |
| AT1G36710.1 | NULL | - | Sampling day | -0.14 |

| | | | | |
|---|---|---|---|---|
| AT1G54095.1 | Protein of unknown function (DUF1677) | - | Sampling day | -0.36 |
| AT1G52220.4 | NULL | - | Sampling day | -0.18 |
| AT5G55508.1 | NULL | - | Sampling day | -0.15 |
| AT1G80140.1 | Pectin lyase-like superfamily protein | - | Sampling day | -0.14 |
| AT4G04415.1 | | - | Sampling day | -0.13 |
| AT1G06430.2 | FTSH protease 8 | FTSH8 | Sampling day | -0.17 |
| AT3G42924.1 | transposable element gene | - | Sampling day | -0.17 |
| AT1G77390.1 | CYCLIN A1;2 | CYCA1%3B2 | Sampling day | -0.14 |
| AT3G62960.1 | Thioredoxin superfamily protein | - | Sampling day | -0.08 |
| AT1G33355.1 | | - | Sampling day | -0.08 |
| AT2G09650.1 | | - | Sampling day | -0.12 |
| AT3G55430.1 | O-Glycosyl hydrolases family 17 protein | - | Sampling day | -0.06 |
| AT1G06880.1 | pre-tRNA | - | Sampling day | -0.13 |
| AT4G28040.4 | nodulin MtN21 /EamA-like transporter family protein | UMAMIT33 | Sampling day | -0.20 |
| AT1G34480.1 | Cysteine/Histidine-rich C1 domain family protein | - | Sampling day | -0.22 |
| AT5G00430.1 | | - | Sampling day | -0.19 |
| AT2G13820.2 | Bifunctional inhibitor/lipid-transfer protein/seed storage 2S albumin superfamily protein | XYP2 | Sampling day | -0.25 |
| AT5G28140.1 | transposable element gene | - | Sampling day | -0.18 |
| AT1G31630.1 | AGAMOUS-like 86 | AGL86 | Sampling day | -0.19 |
| AT1G27390.2 | translocase outer membrane 20-2 | TOM20-2 | Sampling day | -0.19 |
| AT5G25130.1 | cytochrome P450, family 71, subfamily B, polypeptide 12 | CYP71B12 | Sampling day | -0.13 |
| AT3G43622.1 | transposable element gene | - | Sampling day | -0.06 |
| AT5G18633.1 | transposable element gene | - | Sampling day | -0.16 |
| AT1G61732.1 | MIR776a; miRNA | MIR776A | Sampling day | -0.24 |
| AT5G27930.3 | Protein phosphatase 2C family protein | - | Sampling day | -0.10 |
| AT1G53050.1 | Protein kinase superfamily protein | - | Sampling day | -0.12 |
| AT3G09162.1 | NULL | - | Sampling day | -0.08 |
| AT1G53400.1 | Ubiquitin domain-containing protein | - | Sampling day | -0.13 |
| AT3G09435.1 | | - | Sampling day | -0.09 |
| AT3G44935.1 | NULL | - | Sampling day | -0.32 |
| AT1G39070.1 | | - | Sampling day | -0.21 |
| AT5G07255.1 | | - | Sampling day | -0.21 |
| AT4G18670.1 | Leucine-rich repeat (LRR) family protein | - | Sampling day | -0.10 |
| AT5G54110.2 | membrane-associated mannitol-induced | MAMI | Sampling day | -0.19 |
| AT1G45244.1 | pre-tRNA | - | Sampling day | -0.09 |
| AT1G20620.4 | catalase 3 | CAT3 | Sampling day | -0.10 |
| AT2G29580.1 | CCCH-type zinc fingerfamily protein with RNA-binding domain | MAC5B | Sampling day | -0.13 |
| AT3G03430.1 | Calcium-binding EF-hand family protein | - | Sampling day | -0.06 |
| AT3G55440.1 | triosephosphate isomerase | TPI | Sampling day | -0.05 |
| AT1G34120.1 | inositol polyphosphate 5-phosphatase I | IP5PI | Sampling day | -0.09 |
| AT5G48270.1 | Plant protein of unknown function (DUF868) | - | Sampling day | -0.09 |
| AT1G17120.1 | cationic amino acid transporter 8 | CAT8 | Sampling day | -0.15 |
| AT3G44274.1 | NULL | - | Sampling day | -0.21 |
| AT2G08870.1 | | - | Sampling day | -0.26 |
| AT4G07941.1 | transposable element gene | - | Sampling day | -0.26 |
| AT2G21290.1 | NULL | - | Sampling day | -0.16 |
| AT4G29654.1 | Protein kinase superfamily protein | - | Sampling day | -0.07 |
| AT3G31945.1 | transposable element gene | - | Sampling day | -0.17 |
| AT1G09043.1 | | - | Sampling day | -0.11 |
| AT4G11200.1 | transposable element gene | - | Sampling day | -0.14 |
| AT4G23440.2 | Disease resistance protein (TIR-NBS class) | - | Sampling day | -0.38 |
| AT2G12450.1 | transposable element gene | - | Sampling day | -0.14 |
| AT4G01020.1 | helicase domain-containing protein / IBR domain-containing protein / zinc finger protein-related | - | Sampling day | -0.09 |
| ATCG00140.1 | ATP synthase subunit C family protein | ATPH | Sampling day | -0.24 |
| AT5G20770.1 | transposable element gene | - | Sampling day | -0.12 |
| AT5G13220.6 | jasmonate-zim-domain protein 10 | JAZ10 | Sampling day | -0.11 |
| AT4G25790.1 | CAP (Cysteine-rich secretory proteins, Antigen 5, and Pathogenesis-related 1 protein) superfamily protein | - | Sampling day | -0.16 |
| AT1G76780.2 | HSP20-like chaperones superfamily protein | - | Sampling day | -0.08 |
| AT1G68460.1 | isopentenyltransferase 1 | IPT1 | Sampling day | -0.11 |
| AT5G24352.2 | Serine/threonine-protein kinase WNK (With No Lysine)-related | - | Sampling day | -0.19 |
| AT1G66360.1 | Calcium-dependent lipid-binding (CaLB domain) family protein | - | Sampling day | -0.17 |
| AT3G02845.1 | | - | Sampling day | -0.14 |
| AT1G27880.2 | DEAD/DEAH box RNA helicase family protein | - | Sampling day | -0.26 |
| AT4G16920.2 | Disease resistance protein (TIR-NBS-LRR class) family | - | Sampling day | -0.37 |
| AT3G03828.1 | NULL | - | Sampling day | -0.18 |
| AT3G62370.1 | heme binding | - | Sampling day | -0.08 |
| AT2G33320.1 | Calcium-dependent lipid-binding (CaLB domain) family protein | - | Sampling day | -0.25 |
| AT2G13580.1 | NULL | - | Sampling day | -0.24 |
| AT2G08570.1 | | - | Sampling day | -0.09 |
| AT2G40475.1 | NULL | ASG8 | Sampling day | -0.29 |
| AT1G69660.1 | TRAF-like family protein | - | Sampling day | -0.24 |
| AT2G32785.1 | NULL | - | Sampling day | -0.16 |
| AT2G37720.1 | TRICHOME BIREFRINGENCE-LIKE 15 | TBL15 | Sampling day | -0.17 |

| | | | | |
|---|---|---|---|---|
| AT4G08120.1 | transposable element gene | - | Sampling day | -0.26 |
| AT4G33370.1 | DEA(D/H)-box RNA helicase family protein | - | Sampling day | -0.24 |
| AT2G22040.1 | Transducin/WD40 repeat-like superfamily protein | LST8-2 | Sampling day | -0.22 |
| AT5G17330.1 | glutamate decarboxylase | GAD | Sampling day | -0.16 |
| AT4G24977.1 | NULL | - | Sampling day | -0.23 |
| AT1G30750.1 | NULL | - | Sampling day | -0.19 |
| AT5G57655.2 | xylose isomerase family protein | - | Sampling day | -0.13 |
| AT1G14990.1 | NULL | - | Sampling day | -0.24 |
| AT4G17616.1 | Pentatricopeptide repeat (PPR) superfamily protein | - | Sampling day | -0.07 |
| AT2G31260.1 | autophagy 9 (APG9) | APG9 | Sampling day | -0.17 |
| AT1G26515.1 | F-box and associated interaction domains-containing protein | - | Sampling day | -0.12 |
| AT3G43151.1 | transposable element gene | - | Sampling day | -0.14 |
| AT1G71980.1 | Protease-associated (PA) RING/U-box zinc finger family protein | - | Sampling day | -0.23 |
| AT2G27820.1 | prephenate dehydratase 1 | PD1 | Sampling day | -0.21 |
| AT1G36650.1 | transposable element gene | - | Sampling day | -0.17 |
| AT5G02615.1 | pre-tRNA | - | Sampling day | -0.17 |
| AT5G16290.1 | VALINE-TOLERANT 1 | VAT1 | Sampling day | -0.16 |
| AT1G63130.1 | Tetratricopeptide repeat (TPR)-like superfamily protein | - | Sampling day | -0.18 |
| AT5G60120.3 | target of early activation tagged (EAT) 2 | TOE2 | Sampling day | -0.15 |
| AT5G57520.1 | zinc finger protein 2 | ZFP2 | Sampling day | -0.18 |
| AT2G35090.1 | Protein of unknown function (DUF1640) | - | Sampling day | -0.21 |
| ATCG00980.1 | NULL | TRNR.2 | Sampling day | -0.22 |
| AT5G24660.1 | response to low sulfur 2 | LSU2 | Sampling day | -0.09 |
| AT1G10400.1 | UDP-Glycosyltransferase superfamily protein | - | Sampling day | -0.33 |
| AT5G42900.2 | cold regulated gene 27 | COR27 | Sampling day | -0.23 |
| AT1G24807.1 | Glutamine amidotransferase type 1 family protein | - | Sampling day | -0.06 |
| AT2G05470.1 | transposable element gene | - | Sampling day | -0.05 |
| AT4G37520.1 | Peroxidase superfamily protein | - | Sampling day | -0.16 |
| AT2G34420.1 | photosystem II light harvesting complex gene B1B2 | LHB1B2 | Sampling day | -0.14 |
| AT3G09290.1 | telomerase activator1 | TAC1 | Sampling day | -0.16 |
| AT3G49850.2 | telomere repeat binding factor 3 | TRB3 | Sampling day | -0.27 |
| AT4G39610.1 | Protein of unknown function, DUF617 | - | Sampling day | -0.13 |
| AT5G01300.2 | PEBP (phosphatidylethanolamine-binding protein) family protein | - | Sampling day | -0.21 |
| AT2G06190.1 | transposable element gene | - | Sampling day | -0.14 |
| AT3G48350.1 | Cysteine proteinases superfamily protein | CEP3 | Sampling day | -0.13 |
| AT1G13448.1 | other RNA | - | Sampling day | -0.09 |
| AT3G01225.1 | | - | Sampling day | -0.17 |
| AT4G40085.1 | other RNA | - | Sampling day | -0.09 |
| AT5G57820.1 | zinc ion binding | - | Sampling day | -0.09 |
| AT5G49746.1 | transposable element gene | - | Sampling day | -0.06 |
| AT1G16310.1 | Cation efflux family protein | - | Sampling day | -0.20 |
| AT5G58820.1 | Subtilisin-like serine endopeptidase family protein | - | Sampling day | -0.08 |
| AT2G10608.1 | NULL | - | Sampling day | -0.21 |
| AT2G26610.1 | Transducin family protein / WD-40 repeat family protein | - | Sampling day | -0.06 |
| AT3G54750.3 | NULL | - | Sampling day | -0.07 |
| AT1G21835.1 | Plant thionin family protein | - | Sampling day | -0.20 |
| AT5G46490.3 | Disease resistance protein (TIR-NBS-LRR class) family | - | Sampling day | -0.16 |
| AT5G17830.2 | Plasma-membrane choline transporter family protein | - | Sampling day | -0.25 |
| AT4G15480.1 | UDP-Glycosyltransferase superfamily protein | UGT84A1 | Sampling day | -0.11 |
| AT5G11720.1 | Glycosyl hydrolases family 31 protein | - | Sampling day | -0.19 |
| AT3G57480.1 | zinc finger (C2H2 type, AN1-like) family protein | - | Sampling day | -0.18 |
| AT4G24265.2 | NULL | - | Sampling day | -0.22 |
| AT5G17680.2 | disease resistance protein (TIR-NBS-LRR class), putative | - | Sampling day | -0.19 |
| AT3G33572.1 | transposable element gene | - | Sampling day | -0.15 |
| AT1G16130.1 | wall associated kinase-like 2 | WAKL2 | Sampling day | -0.16 |
| AT1G68920.1 | basic helix-loop-helix (bHLH) DNA-binding superfamily protein | - | Sampling day | -0.18 |
| AT4G33925.1 | NULL | SSN2 | Sampling day | -0.26 |
| AT2G31981.1 | NULL | - | Sampling day | -0.11 |
| AT3G30831.1 | transposable element gene | - | Sampling day | -0.19 |
| AT3G32377.1 | NULL | - | Sampling day | -0.18 |
| AT4G19865.1 | Galactose oxidase/kelch repeat superfamily protein | - | Sampling day | -0.10 |
| AT3G45540.1 | RING/U-box protein with C6HC-type zinc finger | - | Sampling day | -0.20 |
| AT5G04405.1 | | - | Sampling day | -0.09 |
| AT5G04937.1 | | - | Sampling day | -0.09 |
| AT1G18320.1 | Mitochondrial import inner membrane translocase subunit Tim17/Tim22/Tim23 family protein | - | Sampling day | -0.20 |
| AT5G67400.1 | root hair specific 19 | RHS19 | Sampling day | -0.06 |
| AT5G63790.1 | NAC domain containing protein 102 | NAC102 | Sampling day | -0.22 |
| AT3G05480.1 | cell cycle checkpoint control protein family | RAD9 | Sampling day | -0.18 |
| AT4G05680.1 | | - | Sampling day | -0.16 |
| AT5G19330.1 | ARM repeat protein interacting with ABF2 | ARIA | Sampling day | -0.25 |
| AT1G59600.1 | ZCW7 | ZCW7 | Sampling day | -0.18 |
| AT3G14180.1 | sequence-specific DNA binding transcription factors | ASIL2 | Sampling day | -0.19 |
| AT1G10960.1 | ferredoxin 1 | FD1 | Sampling day | -0.11 |
| AT2G38210.1 | putative PDX1-like protein 4 | PDX1L4 | Sampling day | -0.17 |
| AT1G20940.1 | F-box family protein | - | Sampling day | -0.09 |

| | | | | |
|---|---|---|---|---|
| AT5G13780.1 | Acyl-CoA N-acyltransferases (NAT) superfamily protein | - | Sampling day | -0.22 |
| AT1G60600.2 | UbiA prenyltransferase family protein | ABC4 | Sampling day | -0.09 |
| AT3G15960.1 | mismatched DNA binding;ATP binding | - | Sampling day | -0.15 |
| AT2G14595.1 | transposable element gene | - | Sampling day | -0.10 |
| AT2G11440.1 | NULL | - | Sampling day | -0.18 |
| AT1G72910.1 | Toll-Interleukin-Resistance (TIR) domain-containing protein | - | Sampling day | -0.17 |
| AT2G37820.1 | Cysteine/Histidine-rich C1 domain family protein | - | Sampling day | -0.13 |
| AT1G80040.4 | NULL | - | Sampling day | -0.18 |
| AT1G78230.1 | Outer arm dynein light chain 1 protein | - | Sampling day | -0.20 |
| AT2G10617.1 | transposable element gene | - | Sampling day | -0.15 |
| AT1G57530.1 | pre-tRNA | - | Sampling day | -0.10 |
| AT5G36500.1 | ECA1 gametogenesis related family protein | - | Sampling day | -0.08 |
| AT1G46912.1 | F-box associated ubiquitination effector family protein | - | Sampling day | -0.12 |
| AT4G31351.1 | NULL | - | Sampling day | -0.31 |
| AT2G08745.1 | | - | Sampling day | -0.10 |
| AT3G27140.1 | Protein phosphatase 2C family protein | - | Sampling day | -0.13 |
| AT1G72790.1 | hydroxyproline-rich glycoprotein family protein | - | Sampling day | -0.23 |
| AT4G08871.1 | transposable element gene | - | Sampling day | -0.09 |
| AT5G53150.3 | DNAJ heat shock N-terminal domain-containing protein | - | Sampling day | -0.20 |
| AT2G38040.1 | acetyl Co-enzyme a carboxylase carboxyltransferase alpha subunit | CAC3 | Sampling day | -0.25 |
| AT4G29820.1 | homolog of CFIM-25 | CFIM-25 | Sampling day | -0.15 |
| AT1G09677.1 | | - | Sampling day | -0.20 |
| AT1G36406.1 | transposable element gene | - | Sampling day | -0.11 |
| AT4G20690.1 | NULL | - | Sampling day | -0.09 |
| AT3G21352.1 | NULL | - | Sampling day | -0.20 |
| AT5G19875.1 | NULL | - | Sampling day | -0.13 |
| AT1G29390.2 | cold regulated 314 thylakoid membrane 2 | COR314-TM2 | Sampling day | -0.16 |
| AT3G02420.2 | NULL | - | Sampling day | -0.13 |
| AT5G02665.1 | | - | Sampling day | -0.16 |
| AT1G12294.1 | MIR472a; miRNA | MIR472A | Sampling day | -0.18 |
| AT3G44750.1 | histone deacetylase 3 | HDA3 | Sampling day | -0.18 |
| AT4G14770.2 | TESMIN/TSO1-like CXC 2 | TCX2 | Sampling day | -0.28 |
| AT4G20050.4 | Pectin lyase-like superfamily protein | QRT3 | Sampling day | -0.12 |
| AT1G09983.1 | | - | Sampling day | -0.09 |
| AT2G09755.1 | | - | Sampling day | -0.21 |
| AT5G45576.1 | transposable element gene | - | Sampling day | -0.17 |
| AT3G01105.1 | | - | Sampling day | -0.25 |
| AT1G69770.1 | chromomethylase 3 | CMT3 | Sampling day | -0.20 |
| AT3G28930.2 | AIG2-like (avirulence induced gene) family protein | AIG2 | Sampling day | -0.45 |
| AT1G18871.1 | NULL | - | Sampling day | -0.18 |
| AT1G21610.4 | wound-responsive family protein | - | Sampling day | -0.22 |
| ATMG00440.1 | NULL | ORF152A | Sampling day | -0.11 |
| AT1G09795.1 | ATP phosphoribosyl transferase 2 | ATP-PRT2 | Sampling day | -0.18 |
| AT3G04860.1 | Plant protein of unknown function (DUF868) | - | Sampling day | -0.15 |
| AT1G09307.1 | | - | Sampling day | -0.21 |
| AT1G53645.1 | hydroxyproline-rich glycoprotein family protein | - | Sampling day | -0.22 |
| AT4G34750.1 | SAUR-like auxin-responsive protein family | - | Sampling day | -0.19 |
| AT1G35790.1 | transposable element gene | - | Sampling day | -0.06 |
| AT2G04935.1 | | - | Sampling day | -0.25 |
| AT1G06117.1 | | - | Sampling day | -0.30 |
| AT2G28520.1 | vacuolar proton ATPase A1 | VHA-A1 | Sampling day | -0.24 |
| AT2G16580.1 | SAUR-like auxin-responsive protein family | - | Sampling day | -1.04 |
| AT4G26365.1 | snoRNA | - | Sampling day | -0.20 |
| AT5G04347.2 | Plant self-incompatibility protein S1 family | - | Sampling day | -0.24 |
| AT4G02930.1 | GTP binding Elongation factor Tu family protein | - | Sampling day | -0.34 |
| AT1G04570.1 | Major facilitator superfamily protein | - | Sampling day | -0.27 |
| AT2G04375.1 | | U2-10 | Sampling day | -0.20 |
| AT2G03300.1 | Toll-Interleukin-Resistance (TIR) domain family protein | - | Sampling day | -0.39 |
| AT2G35610.1 | xyloglucanase 113 | XEG113 | Sampling day | -0.10 |
| AT1G04640.2 | lipoyltransferase 2 | LIP2 | Sampling day | -0.12 |
| AT2G12750.1 | transposable element gene | - | Sampling day | -0.17 |
| AT1G15500.1 | TLC ATP/ADP transporter | ATNTT2 | Sampling day | -0.21 |
| AT3G44230.1 | NULL | - | Sampling day | -0.31 |
| AT3G28680.1 | Serine carboxypeptidase S28 family protein | - | Sampling day | -0.06 |
| AT3G14400.1 | ubiquitin-specific protease 25 | UBP25 | Sampling day | -0.13 |
| AT5G32613.1 | Zinc knuckle (CCHC-type) family protein | - | Sampling day | -0.06 |
| AT1G53890.4 | Protein of unknown function (DUF567) | - | Sampling day | -0.08 |
| AT1G36310.2 | S-adenosyl-L-methionine-dependent methyltransferases superfamily | - | Sampling day | -0.24 |
| AT2G13970.1 | transposable element gene | - | Sampling day | -0.08 |
| AT2G28350.1 | auxin response factor 10 | ARF10 | Sampling day | -0.20 |
| ATMG09830.1 | | - | Sampling day | -0.22 |
| AT3G08650.1 | ZIP metal ion transporter family | - | Sampling day | -0.26 |
| AT2G36200.1 | P-loop containing nucleoside triphosphate hydrolases superfamily | - | Sampling day | -0.19 |
| AT2G45760.1 | BON association protein 2 | BAP2 | Sampling day | -0.16 |
| AT5G07540.1 | glycine-rich protein 16 | GRP16 | Sampling day | -0.16 |
| AT1G36370.1 | serine hydroxymethyltransferase 7 | SHM7 | Sampling day | -0.11 |

| | | | | |
|---|---|---|---|---|
| AT3G30817.1 | NULL | - | Sampling day | -0.12 |
| AT2G24800.1 | Peroxidase superfamily protein | - | Sampling day | -0.09 |
| AT5G12236.1 | NULL | - | Sampling day | -0.05 |
| AT1G08933.1 | | - | Sampling day | -0.11 |
| AT3G51150.2 | ATP binding microtubule motor family protein | - | Sampling day | -0.12 |
| AT1G18200.1 | RAB GTPase homolog A6B | RABA6b | Sampling day | -0.19 |
| AT3G12080.1 | GTP-binding family protein | emb2738 | Sampling day | -0.18 |
| AT3G11470.4 | 4'-phosphopantetheinyl transferase superfamily | - | Sampling day | -0.10 |
| AT4G07686.1 | transposable element gene | - | Sampling day | -0.25 |
| AT2G34870.1 | hydroxyproline-rich glycoprotein family protein | MEE26 | Sampling day | -0.11 |
| AT1G48310.1 | chromatin remodeling factor18 | CHR18 | Sampling day | -0.15 |
| AT5G04935.1 | | - | Sampling day | -0.24 |
| AT5G38050.1 | RNA polymerase II transcription elongation factor | - | Sampling day | -0.24 |
| AT2G43330.1 | inositol transporter 1 | INT1 | Sampling day | -0.47 |
| AT2G10180.1 | transposable element gene | - | Sampling day | -0.18 |
| AT3G02820.2 | zinc knuckle (CCHC-type) family protein | - | Sampling day | -0.17 |
| AT2G36895.3 | NULL | - | Sampling day | -0.22 |
| AT5G42830.1 | HXXXD-type acyl-transferase family protein | - | Sampling day | -0.15 |
| AT1G73120.1 | NULL | - | Sampling day | -0.10 |
| AT5G59020.1 | Protein of unknown function (DUF3527) | - | Sampling day | -0.10 |
| AT3G08595.1 | | - | Sampling day | -0.17 |
| AT1G64750.3 | deletion of SUV3 suppressor 1(I) | DSS1(I) | Sampling day | -0.12 |
| AT4G10780.2 | LRR and NB-ARC domains-containing disease resistance protein | - | Sampling day | -0.26 |
| AT5G24490.1 | 30S ribosomal protein, putative | - | Sampling day | -0.20 |
| AT3G44260.1 | Polynucleotidyl transferase, ribonuclease H-like superfamily protein | CAF1a | Sampling day | -0.13 |
| AT4G17090.1 | chloroplast beta-amylase | CT-BMY | Sampling day | -0.15 |
| AT5G48860.1 | NULL | - | Sampling day | -0.17 |
| AT4G18640.1 | Leucine-rich repeat protein kinase family protein | MRH1 | Sampling day | -0.14 |
| AT3G43580.1 | Beta-galactosidase related protein | - | Sampling day | -0.08 |
| AT3G32070.1 | transposable element gene | - | Sampling day | -0.07 |
| AT3G45245.1 | ECA1 gametogenesis related family protein | - | Sampling day | -0.09 |
| AT1G49440.1 | transposable element gene | - | Sampling day | -0.11 |
| AT2G07645.1 | | - | Sampling day | -0.19 |
| AT3G52230.1 | NULL | - | Sampling day | -0.21 |
| AT1G80330.1 | gibberellin 3-oxidase 4 | GA3OX4 | Sampling day | -0.19 |
| AT2G36305.2 | farnesylated protein-converting enzyme 2 | FACE2 | Sampling day | -0.21 |
| AT1G01160.2 | GRF1-interacting factor 2 | GIF2 | Sampling day | -0.12 |
| AT1G70210.1 | CYCLIN D1;1 | CYCD1%3B1 | Sampling day | -0.06 |
| AT1G30757.2 | NULL | - | Sampling day | -0.09 |
| AT5G57010.1 | calmodulin-binding family protein | - | Sampling day | -0.16 |
| AT3G25610.1 | ATPase E1-E2 type family protein / haloacid dehalogenase-like hydrolase family protein | - | Sampling day | -0.07 |
| AT1G69090.1 | Protein of unknown function (DUF295) | - | Sampling day | -0.15 |
| AT5G03405.1 | | - | Sampling day | -0.24 |
| AT2G13260.1 | transposable element gene | - | Sampling day | -0.11 |
| AT1G11690.1 | NULL | - | Sampling day | -0.10 |
| AT3G29370.1 | NULL | P1R3 | Sampling day | -0.22 |
| AT5G19700.1 | MATE efflux family protein | - | Sampling day | -0.06 |
| AT1G20270.2 | 2-oxoglutarate (2OG) and Fe(II)-dependent oxygenase superfamily | - | Sampling day | -0.09 |
| AT3G55370.3 | OBF-binding protein 3 | OBP3 | Sampling day | -0.11 |
| AT2G21500.2 | RING/U-box superfamily protein | - | Sampling day | -0.27 |
| AT2G36010.2 | E2F transcription factor 3 | E2F3 | Sampling day | -0.18 |
| AT2G42000.2 | Plant EC metallothionein-like protein, family 15 | AtMT4a | Sampling day | -0.12 |
| AT1G18340.1 | basal transcription factor complex subunit-related | - | Sampling day | -0.14 |
| AT4G14140.1 | DNA methyltransferase 2 | DMT2 | Sampling day | -0.21 |
| AT4G31740.1 | Sec1/munc18-like (SM) proteins superfamily | - | Sampling day | -0.20 |
| AT5G46160.1 | Ribosomal protein L14p/L23e family protein | - | Sampling day | -0.20 |
| AT3G03345.1 | | - | Sampling day | -0.16 |
| AT2G10610.1 | transposable element gene | - | Sampling day | -0.18 |
| AT2G28780.1 | NULL | - | Sampling day | -0.18 |
| AT2G38970.1 | Zinc finger (C3HC4-type RING finger) family protein | - | Sampling day | -0.49 |
| AT1G39590.1 | transposable element gene | - | Sampling day | -0.10 |
| AT4G30740.1 | NULL | - | Sampling day | -0.58 |
| AT3G32445.1 | NULL | - | Sampling day | -0.14 |
| ATCG00180.1 | DNA-directed RNA polymerase family protein | RPOC1 | Sampling day | -0.32 |
| AT1G54680.1 | NULL | - | Sampling day | -0.11 |
| AT5G47580.1 | NULL | ASG7 | Sampling day | -0.07 |
| AT1G56210.1 | Heavy metal transport/detoxification superfamily protein | - | Sampling day | -0.09 |
| AT4G17610.2 | tRNA/rRNA methyltransferase (SpoU) family protein | - | Sampling day | -0.18 |
| AT5G25020.1 | Protein of unknown function (DUF1336) | - | Sampling day | -0.11 |
| AT5G28580.2 | transposable element gene | - | Sampling day | -0.20 |
| AT1G04343.1 | | - | Sampling day | -0.15 |
| AT4G13455.1 | transposable element gene | - | Sampling day | -0.06 |
| AT1G42280.1 | transposable element gene | - | Sampling day | -0.08 |
| AT1G35780.2 | NULL | - | Sampling day | -0.15 |
| AT3G55600.2 | Membrane fusion protein Use1 | - | Sampling day | -0.15 |

| | | | | |
|---|---|---|---|---|
| AT4G10345.1 | MIR832a; miRNA | MIR832A | Sampling day | -0.21 |
| AT3G43291.1 | NULL | - | Sampling day | -0.21 |
| AT3G59530.3 | Calcium-dependent phosphotriesterase superfamily protein | LAP3 | Sampling day | -0.07 |
| AT1G69380.1 | Protein of unknown function (DUF155) | RRG | Sampling day | -0.13 |
| AT5G48605.1 | Putative membrane lipoprotein | - | Sampling day | -0.06 |
| AT3G06035.1 | Glycoprotein membrane precursor GPI-anchored | - | Sampling day | -0.33 |
| AT5G57060.3 | NULL | - | Sampling day | -0.10 |
| AT5G51110.1 | Transcriptional coactivator/pterin dehydratase | - | Sampling day | -0.06 |
| AT1G77010.1 | Pentatricopeptide repeat (PPR) superfamily protein | - | Sampling day | -0.12 |
| AT1G07170.3 | PHF5-like protein | - | Sampling day | -0.15 |
| AT3G05760.1 | C2H2 and C2HC zinc fingers superfamily protein | - | Sampling day | -0.14 |
| AT5G01880.1 | RING/U-box superfamily protein | - | Sampling day | -0.19 |
| AT5G08310.1 | Tetratricopeptide repeat (TPR)-like superfamily protein | - | Sampling day | -0.08 |
| AT2G48120.1 | pale cress protein (PAC) | PAC | Sampling day | -0.07 |
| AT5G52600.1 | myb domain protein 82 | MYB82 | Sampling day | -0.10 |
| AT5G43070.1 | WPP domain protein 1 | WPP1 | Sampling day | -0.18 |
| AT1G32520.1 | NULL | - | Sampling day | -0.20 |
| AT5G47590.2 | Heat shock protein HSP20/alpha crystallin family | - | Sampling day | -0.54 |
| AT3G13225.1 | WW domain-containing protein | - | Sampling day | -0.13 |
| AT5G33256.1 | transposable element gene | - | Sampling day | -0.22 |
| AT3G28430.1 | NULL | - | Sampling day | -0.08 |
| AT2G03905.1 | | - | Sampling day | -0.17 |
| AT3G28715.1 | ATPase, V0/A0 complex, subunit C/D | - | Sampling day | -0.17 |
| AT5G01345.1 | | - | Sampling day | -0.12 |
| AT5G02550.1 | NULL | - | Sampling day | -0.25 |
| AT3G06500.2 | Plant neutral invertase family protein | A/N-InvC | Sampling day | -0.10 |
| AT1G37036.1 | transposable element gene | - | Sampling day | -0.10 |
| AT2G40490.1 | Uroporphyrinogen decarboxylase | HEME2 | Sampling day | -0.20 |
| AT5G14230.1 | NULL | - | Sampling day | -0.12 |
| AT5G13420.1 | Aldolase-type TIM barrel family protein | TRA2 | Sampling day | -0.09 |
| AT2G07420.1 | transposable element gene | - | Sampling day | -0.14 |
| AT5G23908.1 | NULL | - | Sampling day | -0.07 |
| AT2G23580.1 | methyl esterase 4 | MES4 | Sampling day | -0.38 |
| AT3G60390.1 | homeobox-leucine zipper protein 3 | HAT3 | Sampling day | -0.23 |
| AT5G27765.1 | | - | Sampling day | -0.32 |
| AT5G46025.1 | Ras-related small GTP-binding family protein | - | Sampling day | -0.05 |
| AT5G05520.1 | Outer membrane OMP85 family protein | - | Sampling day | -0.15 |
| AT1G36530.1 | transposable element gene | - | Sampling day | -0.12 |
| AT3G08915.1 | | - | Sampling day | -0.25 |
| AT1G01695.1 | Phosphatidylinositol N-acetyglucosaminlytransferase subunit P-related | TRM33 | Sampling day | -0.11 |
| AT1G15380.1 | Lactoylglutathione lyase / glyoxalase I family protein | GLYI4 | Sampling day | -0.18 |
| AT4G36850.2 | PQ-loop repeat family protein / transmembrane family protein | - | Sampling day | -0.11 |
| AT5G54540.1 | Uncharacterised conserved protein (UCP012943) | - | Sampling day | -0.17 |
| AT5G54740.1 | seed storage albumin 5 | SESA5 | Sampling day | -0.22 |
| AT2G38620.1 | cyclin-dependent kinase B1;2 | CDKB1%3B2 | Sampling day | -0.19 |
| AT4G09225.1 | | - | Sampling day | -0.25 |
| AT3G08905.1 | | - | Sampling day | -0.13 |
| AT4G08872.1 | transposable element gene | - | Sampling day | -0.20 |
| AT3G57900.1 | NULL | - | Sampling day | -0.17 |
| AT1G40112.1 | transposable element gene | - | Sampling day | -0.09 |
| AT5G10590.1 | NULL | - | Sampling day | -0.14 |
| AT3G09950.1 | NULL | - | Sampling day | -0.10 |
| AT1G11160.3 | Transducin/WD40 repeat-like superfamily protein | - | Sampling day | -0.11 |
| AT5G09755.1 | pre-tRNA | - | Sampling day | -0.08 |
| AT3G59830.1 | Integrin-linked protein kinase family | - | Sampling day | -0.14 |
| AT5G61640.2 | peptidemethionine sulfoxide reductase 1 | PMSR1 | Sampling day | -0.21 |
| AT3G26235.1 | NULL | - | Sampling day | -0.06 |
| AT1G15260.1 | NULL | - | Sampling day | -0.06 |
| AT1G50470.1 | F-box associated ubiquitination effector family protein | - | Sampling day | -0.14 |
| AT3G43680.1 | transposable element gene | - | Sampling day | -0.14 |
| AT3G58470.1 | nucleic acid binding;methyltransferases | - | Sampling day | -0.14 |
| AT5G55150.1 | Protein of unknown function (DUF295) | - | Sampling day | -0.07 |
| AT2G13542.1 | NULL | - | Sampling day | -0.14 |
| AT5G59280.1 | pumilio 16 | PUM16 | Sampling day | -0.10 |
| AT5G24260.2 | prolyl oligopeptidase family protein | - | Sampling day | -0.21 |
| AT1G54980.1 | Plant invertase/pectin methylesterase inhibitor superfamily protein | - | Sampling day | -0.20 |
| AT3G30350.2 | NULL | RGF4 | Sampling day | -0.12 |
| AT5G16810.2 | Protein kinase superfamily protein | - | Sampling day | -0.13 |
| AT2G14290.1 | F-box family protein with a domain of unknown function (DUF295) | - | Sampling day | -0.18 |
| AT1G34800.1 | Plant thionin family protein | - | Sampling day | -0.16 |
| AT5G16650.2 | Chaperone DnaJ-domain superfamily protein | - | Sampling day | -0.11 |
| AT5G03440.3 | NULL | - | Sampling day | -0.16 |
| AT5G05480.1 | Peptide-N4-(N-acetyl-beta-glucosaminyl)asparagine amidase A protein | - | Sampling day | -0.18 |
| AT5G28160.1 | Galactose oxidase/kelch repeat superfamily protein | - | Sampling day | -0.21 |
| AT1G17720.2 | Protein phosphatase 2A, regulatory subunit PR55 | ATB BETA | Sampling day | -0.16 |
| AT4G24710.2 | P-loop containing nucleoside triphosphate hydrolases superfamily | - | Sampling day | -0.17 |

| | | | | |
|---|---|---|---|---|
| AT2G04340.1 | NULL | - | Sampling day | -0.07 |
| AT4G11402.1 | | - | Sampling day | -0.24 |
| AT5G32423.1 | transposable element gene | - | Sampling day | -0.21 |
| AT2G46650.1 | cytochrome B5 isoform C | CB5-C | Sampling day | -0.15 |
| AT5G26610.2 | D111/G-patch domain-containing protein | - | Sampling day | -0.19 |
| AT5G12470.1 | Protein of unknown function (DUF3411) | - | Sampling day | -0.19 |
| AT2G07827.2 | NULL | - | Sampling day | -0.20 |
| AT3G26818.1 | MIR169M; miRNA | MIR169M | Sampling day | -0.20 |
| AT3G05850.1 | NULL | MUG7 | Sampling day | -0.12 |
| AT5G09470.1 | dicarboxylate carrier 3 | DIC3 | Sampling day | -0.08 |
| AT5G61060.1 | histone deacetylase 5 | HDA05 | Sampling day | -0.08 |
| AT5G56365.1 | pre-tRNA | - | Sampling day | -0.15 |
| AT3G03960.1 | TCP-1/cpn60 chaperonin family protein | - | Sampling day | -0.23 |
| AT5G27450.2 | mevalonate kinase | MK | Sampling day | -0.19 |
| AT1G07543.1 | | - | Sampling day | -0.16 |
| AT2G09845.1 | | - | Sampling day | -0.24 |
| AT3G46230.1 | heat shock protein 17.4 | HSP17.4 | Sampling day | -0.21 |
| AT2G32350.1 | Ubiquitin-like superfamily protein | - | Sampling day | -0.20 |
| AT4G06585.1 | transposable element gene | - | Sampling day | -0.22 |
| AT1G65032.1 | NULL | - | Sampling day | -0.17 |
| AT5G62090.1 | SEUSS-like 2 | SLK2 | Sampling day | -0.08 |
| AT5G32125.1 | transposable element gene | - | Sampling day | -0.15 |
| AT4G08505.1 | | - | Sampling day | -0.16 |
| AT1G15170.1 | MATE efflux family protein | - | Sampling day | -0.14 |
| AT1G54650.2 | Methyltransferase family protein | - | Sampling day | -0.13 |
| AT1G09280.1 | NULL | - | Sampling day | -0.16 |
| AT2G15750.1 | transposable element gene | - | Sampling day | -0.09 |
| AT1G51805.1 | Leucine-rich repeat protein kinase family protein | - | Sampling day | -0.16 |
| AT5G54690.1 | galacturonosyltransferase 12 | GAUT12 | Sampling day | -0.20 |
| AT1G33102.1 | NULL | - | Sampling day | -0.17 |
| AT2G04900.1 | NULL | - | Sampling day | -0.12 |
| AT5G32875.1 | transposable element gene | - | Sampling day | -0.09 |
| AT3G59150.1 | F-box/RNI-like superfamily protein | - | Sampling day | -0.20 |
| AT2G07673.1 | NULL | - | Sampling day | -0.18 |
| AT4G11745.1 | Galactose oxidase/kelch repeat superfamily protein | - | Sampling day | -0.22 |
| AT4G12950.1 | Fasciclin-like arabinogalactan family protein | - | Sampling day | -0.32 |
| AT3G56410.1 | Protein of unknown function (DUF3133) | - | Sampling day | -0.21 |
| AT3G30440.1 | transposable element gene | - | Sampling day | -0.20 |
| AT2G20840.2 | Secretory carrier membrane protein (SCAMP) family protein | SCAMP3 | Sampling day | -0.05 |
| AT3G23680.1 | F-box associated ubiquitination effector family protein | - | Sampling day | -0.13 |
| AT2G08780.1 | | - | Sampling day | -0.09 |
| AT4G24830.1 | arginosuccinate synthase family | - | Sampling day | -0.09 |
| AT2G38080.1 | Laccase/Diphenol oxidase family protein | IRX12 | Sampling day | -0.25 |
| AT2G42388.1 | other RNA | - | Sampling day | -0.06 |
| AT1G06790.1 | RNA polymerase Rpb7 N-terminal domain-containing protein | - | Sampling day | -0.20 |
| AT5G09475.1 | | - | Sampling day | -0.21 |
| AT5G28885.1 | NULL | - | Sampling day | -0.22 |
| AT4G28180.1 | NULL | - | Sampling day | -0.17 |
| AT1G21830.1 | NULL | - | Sampling day | -0.12 |
| AT5G48335.1 | NULL | - | Sampling day | -0.12 |
| AT1G26190.2 | Phosphoribulokinase / Uridine kinase family | - | Sampling day | -0.14 |
| AT3G02605.1 | | - | Sampling day | -0.08 |
| AT1G05540.2 | Protein of unknown function (DUF295) | - | Sampling day | -0.09 |
| AT3G60955.2 | NULL | CYP76C8P | Sampling day | -0.13 |
| AT3G10020.1 | NULL | - | Sampling day | -0.16 |
| AT1G73810.1 | Core-2/I-branching beta-1,6-N-acetylglucosaminyltransferase family | - | Sampling day | -0.34 |
| AT4G13950.1 | ralf-like 31 | RALFL31 | Sampling day | -0.13 |
| AT1G44970.1 | Peroxidase superfamily protein | - | Sampling day | -0.16 |
| AT3G05295.1 | | - | Sampling day | -0.20 |
| AT5G44306.1 | NULL | - | Sampling day | -0.27 |
| AT1G60500.1 | Dynamin related protein 4C | DRP4C | Sampling day | -0.16 |
| AT1G30590.2 | RNA polymerase I specific transcription initiation factor RRN3 protein | - | Sampling day | -0.12 |
| AT5G03630.1 | Pyridine nucleotide-disulphide oxidoreductase family protein | ATMDAR2 | Sampling day | -0.26 |
| AT4G17450.1 | transposable element gene | - | Sampling day | -0.16 |
| AT3G24575.1 | | - | Sampling day | -0.28 |
| AT1G79530.1 | glyceraldehyde-3-phosphate dehydrogenase of plastid 1 | GAPCP-1 | Sampling day | -0.41 |
| AT1G29290.1 | NULL | - | Sampling day | -0.28 |
| AT5G06190.2 | NULL | - | Sampling day | -0.08 |
| AT4G25210.1 | DNA-binding storekeeper protein-related transcriptional regulator | - | Sampling day | -0.17 |
| AT5G20225.1 | other RNA | - | Sampling day | -0.22 |
| AT2G23140.3 | RING/U-box superfamily protein with ARM repeat domain. | PUB4 | Sampling day | -0.14 |
| AT1G12667.1 | NULL | - | Sampling day | -0.12 |
| AT1G16180.1 | Serinc-domain containing serine and sphingolipid biosynthesis protein | - | Sampling day | -0.17 |
| AT4G03505.1 | NULL | - | Sampling day | -0.07 |
| AT1G69470.1 | NULL | - | Sampling day | -0.10 |
| AT2G33720.1 | AP2/B3-like transcriptional factor family protein | - | Sampling day | -0.26 |

| | | | | |
|---|---|---|---|---|
| AT1G06133.1 | | - | Sampling day | -0.14 |
| AT1G19380.1 | Protein of unknown function (DUF1195) | - | Sampling day | -0.20 |
| AT5G51500.1 | Plant invertase/pectin methylesterase inhibitor superfamily | - | Sampling day | -0.32 |
| AT5G16550.1 | NULL | - | Sampling day | -0.14 |
| AT1G63830.4 | PLAC8 family protein | - | Sampling day | -0.10 |
| AT2G12130.1 | transposable element gene | - | Sampling day | -0.07 |
| AT1G53920.1 | GDSL-motif lipase 5 | GLIP5 | Sampling day | -0.28 |
| AT3G21700.3 | Ras-related small GTP-binding family protein | SGP2 | Sampling day | -0.15 |
| AT1G05927.1 | | - | Sampling day | -0.21 |
| AT4G36990.1 | heat shock factor 4 | HSF4 | Sampling day | -1.14 |
| AT1G50700.1 | calcium-dependent protein kinase 33 | CPK33 | Sampling day | -0.14 |
| AT2G46800.3 | zinc transporter of Arabidopsis thaliana | ZAT | Sampling day | -0.14 |
| AT1G67580.1 | Protein kinase superfamily protein | - | Sampling day | -0.30 |
| AT4G10120.3 | Sucrose-phosphate synthase family protein | ATSPS4F | Sampling day | -0.46 |
| AT5G31873.1 | transposable element gene | - | Sampling day | -0.18 |
| AT2G08945.1 | | - | Sampling day | -0.08 |
| ATCG00210.1 | electron transporter, transferring electrons within cytochrome b6/f complex of photosystem IIs | YCF6 | Sampling day | -0.19 |
| AT1G63730.1 | Disease resistance protein (TIR-NBS-LRR class) family | - | Sampling day | -0.11 |
| AT1G33590.2 | Leucine-rich repeat (LRR) family protein | - | Sampling day | -0.10 |
| AT3G32385.1 | transposable element gene | - | Sampling day | -0.22 |
| AT3G04345.1 | | - | Sampling day | -0.10 |
| AT4G14930.2 | Survival protein SurE-like phosphatase/nucleotidase | - | Sampling day | -0.11 |
| AT1G63270.1 | non-intrinsic ABC protein 10 | ABCI1 | Sampling day | -0.13 |
| AT3G47050.1 | Glycosyl hydrolase family protein | - | Sampling day | -0.15 |
| AT2G17972.1 | NULL | - | Sampling day | -0.09 |
| AT3G44730.2 | kinesin-like protein 1 | KP1 | Sampling day | -0.32 |
| AT2G09060.1 | | - | Sampling day | -0.11 |
| AT4G04450.2 | WRKY family transcription factor | WRKY42 | Sampling day | -0.24 |
| AT3G42837.1 | transposable element gene | - | Sampling day | -0.15 |
| AT1G53360.2 | F-box associated ubiquitination effector family protein | - | Sampling day | -0.07 |
| AT2G19840.1 | transposable element gene | - | Sampling day | -0.06 |
| AT5G53500.1 | Transducin/WD40 repeat-like superfamily protein | - | Sampling day | -0.26 |
| AT3G52290.1 | IQ-domain 3 | IQD3 | Sampling day | -0.07 |
| AT3G28180.1 | Cellulose-synthase-like C4 | CSLC04 | Sampling day | -0.10 |
| AT5G42232.1 | Defensin-like (DEFL) family protein | - | Sampling day | -0.09 |
| AT1G22400.1 | UDP-Glycosyltransferase superfamily protein | UGT85A1 | Sampling day | -0.19 |
| AT5G12210.1 | RAB geranylgeranyl transferase beta subunit 1 | RGTB1 | Sampling day | -0.26 |
| AT1G79760.2 | downstream target of AGL15-4 | DTA4 | Sampling day | -0.27 |
| AT4G24240.2 | WRKY DNA-binding protein 7 | WRKY7 | Sampling day | -0.64 |
| AT4G12240.1 | zinc finger (C2H2 type) family protein | - | Sampling day | -0.14 |
| AT3G44800.1 | Meprin and TRAF (MATH) homology domain-containing protein | - | Sampling day | -0.08 |
| AT2G14330.1 | transposable element gene | - | Sampling day | -0.20 |
| AT1G16070.1 | tubby like protein 8 | TLP8 | Sampling day | -0.15 |
| AT2G42560.1 | late embryogenesis abundant domain-containing protein / LEA domain-containing protein | - | Sampling day | -0.12 |
| AT4G09035.1 | | - | Sampling day | -0.17 |
| AT5G19200.1 | NAD(P)-binding Rossmann-fold superfamily protein | TSC10B | Sampling day | -0.23 |
| AT2G07085.1 | | - | Sampling day | -0.11 |
| AT3G27095.1 | NULL | - | Sampling day | -0.14 |
| AT1G62610.4 | NAD(P)-binding Rossmann-fold superfamily protein | - | Sampling day | -0.17 |
| AT1G34790.1 | C2H2 and C2HC zinc fingers superfamily protein | TT1 | Sampling day | -0.13 |
| AT1G01335.1 | | - | Sampling day | -0.19 |
| AT2G10560.1 | NULL | - | Sampling day | -0.31 |
| AT1G22220.1 | F-box family protein | AUF2 | Sampling day | -0.28 |
| AT2G45135.2 | RING/U-box superfamily protein | - | Sampling day | -0.16 |
| AT1G65342.1 | NULL | - | Sampling day | -0.17 |
| AT3G32240.1 | transposable element gene | - | Sampling day | -0.15 |
| AT5G14520.1 | pescadillo-related | - | Sampling day | -0.19 |
| AT4G03250.1 | Homeodomain-like superfamily protein | - | Sampling day | -0.09 |
| AT1G02560.1 | nuclear encoded CLP protease 5 | CLPP5 | Sampling day | -0.20 |
| AT5G35341.1 | transposable element gene | - | Sampling day | -0.12 |
| AT1G44020.1 | Cysteine/Histidine-rich C1 domain family protein | - | Sampling day | -0.15 |
| AT3G24225.1 | CLAVATA3/ESR-RELATED 19 | CLE19 | Sampling day | -0.14 |
| AT4G28110.2 | myb domain protein 41 | MYB41 | Sampling day | -0.25 |
| AT1G50160.1 | NULL | - | Sampling day | -0.46 |
| AT1G06043.1 | | - | Sampling day | -0.19 |
| AT4G14190.1 | Pentatricopeptide repeat (PPR) superfamily protein | - | Sampling day | -0.22 |
| AT1G35610.1 | Cysteine/Histidine-rich C1 domain family protein | - | Sampling day | -0.22 |
| AT3G46640.2 | Homeodomain-like superfamily protein | PCL1 | Sampling day | -0.18 |
| AT3G54420.1 | homolog of carrot EP3-3 chitinase | EP3 | Sampling day | -0.23 |
| AT3G17480.1 | F-box and associated interaction domains-containing protein | - | Sampling day | -0.13 |
| AT4G06040.1 | | - | Sampling day | -0.16 |
| AT1G71100.1 | Ribose 5-phosphate isomerase, type A protein | RSW10 | Sampling day | -0.07 |
| AT1G78270.1 | UDP-glucosyl transferase 85A4 | UGT85A4 | Sampling day | -0.36 |
| AT2G43160.1 | ENTH/VHS family protein | - | Sampling day | -0.12 |

| | | | | |
|---|---|---|---|---|
| AT2G02860.1 | sucrose transporter 2 | SUT2 | Sampling day | -0.08 |
| AT3G26050.1 | TPX2 (targeting protein for Xklp2) protein family | - | Sampling day | -0.22 |
| AT4G29060.1 | elongation factor Ts family protein | emb2726 | Sampling day | -0.17 |
| AT3G47380.1 | Plant invertase/pectin methylesterase inhibitor superfamily protein | - | Sampling day | -0.36 |
| AT5G57290.1 | 60S acidic ribosomal protein family | - | Sampling day | -0.16 |
| AT3G32116.1 | transposable element gene | - | Sampling day | -0.17 |
| AT3G16770.1 | ethylene-responsive element binding protein | EBP | Sampling day | -0.40 |
| AT3G46183.1 | transposable element gene | - | Sampling day | -0.19 |
| AT2G46840.1 | DOMAIN OF UNKNOWN FUNCTION 724 4 | DUF4 | Sampling day | -0.23 |
| AT3G61270.2 | Arabidopsis thaliana protein of unknown function (DUF821) | - | Sampling day | -0.05 |
| AT2G45690.1 | shrunken seed protein (SSE1) | SSE1 | Sampling day | -0.26 |
| AT2G17080.1 | Arabidopsis protein of unknown function (DUF241) | - | Sampling day | -0.32 |
| AT1G79790.4 | Haloacid dehalogenase-like hydrolase (HAD) superfamily protein | FHY1 | Sampling day | -0.25 |
| AT5G28440.1 | unknown protein | - | Sampling day | -0.20 |
| AT1G12490.1 | F-box associated ubiquitination effector family protein | - | Sampling day | -0.10 |
| AT3G52440.2 | Dof-type zinc finger DNA-binding family protein | - | Sampling day | -0.10 |
| AT5G32306.1 | transposable element gene | - | Sampling day | -0.17 |
| AT1G34030.1 | Ribosomal protein S13/S18 family | - | Sampling day | -0.20 |
| AT5G58380.1 | SOS3-interacting protein 1 | SIP1 | Sampling day | -0.24 |
| AT2G24150.1 | heptahelical protein 3 | HHP3 | Sampling day | -0.14 |
| AT1G59880.1 | pre-tRNA | - | Sampling day | -0.11 |
| AT4G13555.1 | MIR397B; miRNA | MIR397B | Sampling day | -0.12 |
| AT5G22560.1 | Plant protein of unknown function (DUF247) | - | Sampling day | -0.07 |
| AT5G39785.3 | Protein of unknown function (DUF1666) | - | Sampling day | -0.21 |
| AT4G20490.1 | transposable element gene | - | Sampling day | -0.36 |
| AT5G12060.1 | Plant self-incompatibility protein S1 family | - | Sampling day | -0.13 |
| AT1G71340.1 | PLC-like phosphodiesterases superfamily protein | GDPD4 | Sampling day | -0.10 |
| AT4G26680.1 | Tetratricopeptide repeat (TPR)-like superfamily protein | - | Sampling day | -0.26 |
| AT1G27480.1 | alpha/beta-Hydrolases superfamily protein | - | Sampling day | -0.21 |
| AT1G28140.1 | NULL | - | Sampling day | -0.11 |
| AT5G17790.1 | zinc finger (Ran-binding) family protein | VAR3 | Sampling day | -0.07 |
| AT3G52075.1 | | - | Sampling day | -0.08 |
| AT5G36658.1 | ECA1 gametogenesis related family protein | - | Sampling day | -0.09 |
| AT5G14990.1 | NULL | - | Sampling day | -0.07 |
| AT1G07357.1 | | - | Sampling day | -0.08 |
| AT4G30067.1 | low-molecular-weight cysteine-rich 63 | LCR63 | Sampling day | -0.17 |
| AT3G32898.1 | transposable element gene | - | Sampling day | -0.10 |
| AT2G22121.1 | low-molecular-weight cysteine-rich 35 | LCR35 | Sampling day | -0.19 |
| AT5G23410.1 | | - | Sampling day | -0.19 |
| AT3G21950.2 | S-adenosyl-L-methionine-dependent methyltransferases superfamily | - | Sampling day | -0.10 |
| AT3G16090.1 | RING/U-box superfamily protein | Hrd1A | Sampling day | -0.06 |
| AT1G77310.1 | NULL | - | Sampling day | -0.13 |
| AT2G27140.1 | HSP20-like chaperones superfamily protein | - | Sampling day | -0.14 |
| AT1G43090.1 | Pectin lyase-like superfamily protein | - | Sampling day | -0.21 |
| AT2G38420.1 | Pentatricopeptide repeat (PPR) superfamily protein | - | Sampling day | -0.21 |
| AT1G25425.1 | CLAVATA3/ESR-RELATED 43 | CLE43 | Sampling day | -0.42 |
| AT1G74090.1 | desulfo-glucosinolate sulfotransferase 18 | SOT18 | Sampling day | -0.13 |
| AT1G09812.1 | NULL | - | Sampling day | -0.08 |
| AT2G24110.1 | NULL | - | Sampling day | -0.21 |
| AT5G19485.1 | transferases;nucleotidyltransferases | - | Sampling day | -0.11 |
| AT5G22799.1 | | - | Sampling day | -0.24 |
| AT5G24550.1 | beta glucosidase 32 | BGLU32 | Sampling day | -0.20 |
| AT5G58560.1 | Phosphatidate cytidylyltransferase family protein | FOLK | Sampling day | -0.12 |
| AT3G15280.1 | NULL | - | Sampling day | -0.19 |
| AT3G44425.1 | transposable element gene | - | Sampling day | -0.07 |
| AT4G39050.1 | Kinesin motor family protein | - | Sampling day | -0.14 |
| AT2G10400.1 | transposable element gene | - | Sampling day | -0.20 |
| AT3G31358.1 | transposable element gene | - | Sampling day | -0.08 |
| AT5G42370.2 | Calcineurin-like metallo-phosphoesterase superfamily protein | - | Sampling day | -0.14 |
| AT3G33058.1 | transposable element gene | - | Sampling day | -0.12 |
| AT1G80850.1 | DNA glycosylase superfamily protein | - | Sampling day | -0.09 |
| AT3G14500.1 | transposable element gene | - | Sampling day | -0.17 |
| AT2G19820.1 | LOB domain-containing protein 9 | LBD9 | Sampling day | -0.10 |
| AT3G59590.1 | jacalin lectin family protein | - | Sampling day | -0.13 |
| AT5G47630.2 | mitochondrial acyl carrier protein 3 | mtACP3 | Sampling day | -0.27 |
| AT5G14440.1 | Surfeit locus protein 2 (SURF2) | - | Sampling day | -0.18 |
| AT2G31920.1 | Plant protein of unknown function (DUF936) | - | Sampling day | -0.09 |
| AT3G61050.2 | Calcium-dependent lipid-binding (CaLB domain) family protein | NTMC2T4 | Sampling day | -0.10 |
| AT4G20920.2 | double-stranded RNA-binding domain (DsRBD)-containing protein | - | Sampling day | -0.19 |
| AT5G35160.4 | Endomembrane protein 70 protein family | - | Sampling day | -0.27 |
| AT1G51230.1 | Plant self-incompatibility protein S1 family | - | Sampling day | -0.07 |
| AT3G08700.1 | ubiquitin-conjugating enzyme 12 | UBC12 | Sampling day | -0.15 |
| AT5G14530.1 | Transducin/WD40 repeat-like superfamily protein | - | Sampling day | -0.29 |
| AT5G62830.1 | F-box associated ubiquitination effector family protein | - | Sampling day | -0.13 |
| AT5G36300.2 | Tetratricopeptide repeat (TPR)-like superfamily protein | - | Sampling day | -0.14 |
| AT1G32050.1 | SCAMP family protein | SCAMP5 | Sampling day | -0.20 |

| | | | | |
|---|---|---|---|---|
| AT1G09740.1 | Adenine nucleotide alpha hydrolases-like superfamily protein | - | Sampling day | -0.16 |
| AT4G16230.1 | GDSL-like Lipase/Acylhydrolase superfamily protein | - | Sampling day | -0.21 |
| AT5G33441.1 | NULL | - | Sampling day | -0.06 |
| AT1G07500.1 | NULL | - | Sampling day | -0.14 |
| AT1G54560.2 | Myosin family protein with Dil domain | XIE | Sampling day | -0.44 |
| AT4G30975.2 | other RNA | - | Sampling day | -0.11 |
| AT2G20835.1 | NULL | - | Sampling day | -0.14 |
| AT1G09685.2 | | - | Sampling day | -0.15 |
| AT2G01330.1 | nucleotide binding | - | Sampling day | -0.23 |
| AT5G49610.1 | F-box family protein | - | Sampling day | -0.05 |
| AT5G20660.1 | Zn-dependent exopeptidases superfamily protein | - | Sampling day | -0.34 |
| AT5G00425.1 | | - | Sampling day | -0.21 |
| AT4G27430.1 | COP1-interacting protein 7 | CIP7 | Sampling day | -0.27 |
| AT2G27145.1 | low-molecular-weight cysteine-rich 9 | LCR9 | Sampling day | -0.37 |
| AT2G15012.1 | NULL | - | Sampling day | -0.10 |
| AT1G69252.1 | other RNA | - | Sampling day | -0.35 |
| AT3G20670.1 | histone H2A 13 | HTA13 | Sampling day | -0.11 |
| AT2G40116.1 | Phosphoinositide-specific phospholipase C family protein | - | Sampling day | -0.15 |
| AT5G01265.1 | | - | Sampling day | -0.06 |
| AT1G55460.1 | DNA/RNA-binding protein Kin17, conserved region | - | Sampling day | -0.11 |
| AT2G18900.1 | Transducin/WD40 repeat-like superfamily protein | - | Sampling day | -0.09 |
| AT5G40560.1 | DegP protease 13 | DEG13 | Sampling day | -0.05 |
| AT4G25100.4 | Fe superoxide dismutase 1 | FSD1 | Sampling day | -0.18 |
| AT3G08455.1 | | - | Sampling day | -0.12 |
| AT3G01680.1 | NULL | SEOR1 | Sampling day | -0.11 |
| AT1G66440.1 | Cysteine/Histidine-rich C1 domain family protein | - | Sampling day | -0.16 |
| AT5G28993.1 | transposable element gene | - | Sampling day | -0.18 |
| AT1G50130.1 | NULL | - | Sampling day | -0.26 |
| AT3G13065.1 | STRUBBELIG-receptor family 4 | SRF4 | Sampling day | -0.25 |
| AT3G48220.1 | NULL | - | Sampling day | -0.21 |
| AT3G18200.2 | nodulin MtN21 /EamA-like transporter family protein | UMAMIT4 | Sampling day | -0.08 |
| AT5G61740.1 | ABC2 homolog 14 | ABCA10 | Sampling day | -0.13 |
| AT2G46200.3 | NULL | - | Sampling day | -0.10 |
| AT1G26800.1 | RING/U-box superfamily protein | - | Sampling day | -0.11 |
| AT4G38010.1 | Pentatricopeptide repeat (PPR-like) superfamily protein | - | Sampling day | -0.16 |
| AT1G07200.2 | Double Clp-N motif-containing P-loop nucleoside triphosphate hydrolases superfamily protein | - | Sampling day | -0.10 |
| AT1G53820.1 | RING/U-box superfamily protein | - | Sampling day | -0.27 |
| AT1G51900.1 | Regulator of Vps4 activity in the MVB pathway protein | - | Sampling day | -0.29 |
| AT1G32020.1 | F-box family protein | - | Sampling day | -0.20 |
| AT1G20710.1 | WUSCHEL related homeobox 10 | WOX10 | Sampling day | -0.17 |
| AT1G56570.2 | Tetratricopeptide repeat (TPR)-like superfamily protein | PGN | Sampling day | -0.15 |
| AT3G14010.5 | CTC-interacting domain 4 | CID4 | Sampling day | -0.20 |
| AT3G42471.1 | transposable element gene | - | Sampling day | -0.20 |
| AT4G02020.3 | SET domain-containing protein | SWN | Sampling day | -0.13 |
| AT3G60805.1 | pre-tRNA | - | Sampling day | -0.15 |
| AT2G14990.1 | transposable element gene | - | Sampling day | -0.15 |
| AT1G57280.1 | pre-tRNA | - | Sampling day | -0.09 |
| AT1G63220.1 | Calcium-dependent lipid-binding (CaLB domain) family protein | - | Sampling day | -0.17 |
| AT2G03340.1 | WRKY DNA-binding protein 3 | WRKY3 | Sampling day | -0.07 |
| AT5G50870.1 | ubiquitin-conjugating enzyme 27 | UBC27 | Sampling day | -0.13 |
| AT3G49832.2 | NULL | - | Sampling day | -0.22 |
| AT5G34850.1 | purple acid phosphatase 26 | PAP26 | Sampling day | -0.20 |
| AT1G42745.1 | transposable element gene | - | Sampling day | -0.14 |
| AT1G32928.1 | NULL | - | Sampling day | -0.10 |
| AT3G29820.1 | transposable element gene | - | Sampling day | -0.14 |
| AT5G28785.1 | transposable element gene | - | Sampling day | -0.07 |
| AT4G38213.1 | | - | Sampling day | -0.07 |
| AT2G41040.1 | S-adenosyl-L-methionine-dependent methyltransferases superfamily | - | Sampling day | -0.28 |
| AT4G13010.1 | Oxidoreductase, zinc-binding dehydrogenase family protein | - | Sampling day | -0.21 |
| AT5G26950.1 | AGAMOUS-like 93 | AGL93 | Sampling day | -0.08 |
| AT5G65600.1 | Concanavalin A-like lectin protein kinase family protein | - | Sampling day | -0.38 |
| AT5G19740.1 | Peptidase M28 family protein | - | Sampling day | -0.08 |
| AT5G62580.1 | ARM repeat superfamily protein | - | Sampling day | -0.30 |
| AT3G30837.1 | transposable element gene | - | Sampling day | -0.11 |
| AT2G34925.1 | CLAVATA3/ESR-RELATED 42 | CLE42 | Sampling day | -0.10 |
| AT3G49200.1 | O-acyltransferase (WSD1-like) family protein | - | Sampling day | -0.38 |
| AT5G56820.1 | F-box/RNI-like/FBD-like domains-containing protein | - | Sampling day | -0.23 |
| AT5G56490.1 | D-arabinono-1,4-lactone oxidase family protein | GulLO4 | Sampling day | -0.15 |
| AT3G46050.1 | Galactose oxidase/kelch repeat superfamily protein | - | Sampling day | -0.18 |
| AT1G29590.1 | Eukaryotic initiation factor 4E protein | eIF4E3 | Sampling day | -0.11 |
| AT2G16740.1 | ubiquitin-conjugating enzyme 29 | UBC29 | Sampling day | -0.25 |
| AT4G06526.1 | nucleic acid binding;zinc ion binding | - | Sampling day | -0.19 |
| AT4G38780.1 | Pre-mRNA-processing-splicing factor | - | Sampling day | -0.17 |
| AT5G56670.1 | Ribosomal protein S30 family protein | - | Sampling day | -0.24 |
| AT2G39950.3 | NULL | - | Sampling day | -0.19 |

| | | | | |
|---|---|---|---|---|
| AT2G02020.1 | Major facilitator superfamily protein | PTR4 | Sampling day | -0.17 |
| AT5G00675.1 | | - | Sampling day | -0.31 |
| AT2G04265.1 | | - | Sampling day | -0.33 |
| AT1G68960.2 | Protein of unknown function (DUF295) | - | Sampling day | -0.25 |
| AT1G52087.1 | transposable element gene | - | Sampling day | -0.05 |
| AT4G12930.1 | NULL | - | Sampling day | -0.14 |
| AT2G17060.2 | Disease resistance protein (TIR-NBS-LRR class) family | - | Sampling day | -0.34 |
| AT4G28890.1 | RING/U-box superfamily protein | - | Sampling day | -0.09 |
| AT5G10870.1 | chorismate mutase 2 | CM2 | Sampling day | -0.07 |
| AT4G09143.1 | NULL | - | Sampling day | -0.16 |
| AT1G51540.1 | Galactose oxidase/kelch repeat superfamily protein | - | Sampling day | -0.08 |
| AT5G02495.1 | | - | Sampling day | -0.36 |
| AT3G29764.1 | NULL | - | Sampling day | -0.15 |
| AT4G01520.1 | NAC domain containing protein 67 | NAC067 | Sampling day | -0.22 |
| AT5G62162.1 | MIR399C; miRNA | MIR399C | Sampling day | -0.16 |
| AT5G22210.2 | NULL | - | Sampling day | -0.14 |
| AT1G02920.1 | glutathione S-transferase 7 | GSTF7 | Sampling day | -0.24 |
| AT4G32760.3 | ENTH/VHS/GAT family protein | - | Sampling day | -0.10 |
| AT5G07355.1 | | - | Sampling day | -0.12 |
| AT5G59090.1 | subtilase 4.12 | SBT4.12 | Sampling day | -0.07 |
| AT4G09452.1 | NULL | - | Sampling day | -0.12 |
| AT5G43340.1 | phosphate transporter 1;6 | PHT1%3B6 | Sampling day | -0.35 |
| AT1G61880.1 | pre-tRNA | - | Sampling day | -0.28 |
| AT5G27390.2 | Mog1/PsbP/DUF1795-like photosystem II reaction center PsbP family | - | Sampling day | -0.19 |
| AT1G73240.1 | NULL | - | Sampling day | -0.43 |
| AT4G06835.1 | | - | Sampling day | -0.27 |
| AT5G10270.1 | cyclin-dependent kinase C;1 | CDKC%3B1 | Sampling day | -0.18 |
| AT4G08475.1 | | - | Sampling day | -0.09 |
| AT5G24590.2 | TCV-interacting protein | TIP | Sampling day | -0.15 |
| AT3G06960.2 | pigment defective 320 | PDE320 | Sampling day | -0.14 |
| AT1G40105.1 | transposable element gene | - | Sampling day | -0.19 |
| AT4G05585.1 | transposable element gene | - | Sampling day | -0.11 |
| AT5G35270.1 | transposable element gene | - | Sampling day | -0.14 |
| AT3G55512.1 | MIR172D; miRNA | MIR172D | Sampling day | -0.17 |
| AT3G53300.1 | cytochrome P450, family 71, subfamily B, polypeptide 31 | CYP71B31 | Sampling day | -0.19 |
| AT3G63270.1 | NULL | - | Sampling day | -0.10 |
| AT3G15111.1 | NULL | - | Sampling day | -0.16 |
| AT4G20800.1 | FAD-binding Berberine family protein | - | Sampling day | -0.16 |
| AT4G06440.1 | | - | Sampling day | -0.19 |
| AT1G32585.1 | VQ motif-containing protein-related | - | Sampling day | -0.18 |
| AT4G20200.1 | Terpenoid cyclases/Protein prenyltransferases superfamily protein | - | Sampling day | -0.06 |
| AT5G37640.1 | ubiquitin 9 | UBQ9 | Sampling day | -0.18 |
| AT2G06885.1 | transposable element gene | - | Sampling day | -0.18 |
| AT1G71780.1 | NULL | - | Sampling day | -0.13 |
| AT5G20720.3 | chaperonin 20 | CPN20 | Sampling day | -0.16 |
| AT2G04450.1 | nudix hydrolase homolog 6 | NUDT6 | Sampling day | -0.20 |
| AT5G40220.1 | AGAMOUS-like 43 | AGL43 | Sampling day | -0.40 |
| AT2G13570.1 | nuclear factor Y, subunit B7 | NF-YB7 | Sampling day | -0.07 |
| AT2G29550.1 | tubulin beta-7 chain | TUB7 | Sampling day | -0.14 |
| AT1G77540.1 | Acyl-CoA N-acyltransferases (NAT) superfamily protein | - | Sampling day | -0.11 |
| AT4G26300.1 | Arginyl-tRNA synthetase, class Ic | emb1027 | Sampling day | -0.14 |
| AT5G23230.1 | nicotinamidase 2 | NIC2 | Sampling day | -0.10 |
| AT3G32060.1 | transposable element gene | - | Sampling day | -0.26 |
| AT3G01655.1 | | - | Sampling day | -0.48 |
| AT3G22234.1 | NULL | - | Sampling day | -0.11 |
| AT2G18030.1 | Peptide methionine sulfoxide reductase family protein | MSRA5 | Sampling day | -0.13 |
| AT2G08210.1 | | - | Sampling day | -0.10 |
| AT4G29560.1 | NULL | - | Sampling day | -0.27 |
| AT1G51530.2 | RNA-binding (RRM/RBD/RNP motifs) family protein | - | Sampling day | -0.14 |
| AT5G45570.2 | Ulp1 protease family protein | - | Sampling day | -0.11 |
| AT5G60880.2 | breaking of asymmetry in the stomatal lineage | BASL | Sampling day | -0.17 |
| AT4G34460.2 | GTP binding protein beta 1 | AGB1 | Sampling day | -0.15 |
| AT2G18550.1 | homeobox protein 21 | HB21 | Sampling day | -0.25 |
| AT4G23515.2 | Toll-Interleukin-Resistance (TIR) domain family protein | - | Sampling day | -0.05 |
| AT1G31460.1 | NULL | - | Sampling day | -0.13 |
| AT1G28860.1 | pre-tRNA | - | Sampling day | -0.06 |
| AT5G57220.1 | cytochrome P450, family 81, subfamily F, polypeptide 2 | CYP81F2 | Sampling day | -0.14 |
| AT5G01650.3 | Tautomerase/MIF superfamily protein | - | Sampling day | -0.17 |
| AT4G06474.1 | transposable element gene | - | Sampling day | -0.12 |
| AT3G19360.2 | Zinc finger (CCCH-type) family protein | - | Sampling day | -0.16 |
| AT3G26910.3 | hydroxyproline-rich glycoprotein family protein | - | Sampling day | -0.06 |
| AT4G10690.1 | transposable element gene | - | Sampling day | -0.14 |
| AT5G21910.1 | NULL | - | Sampling day | -0.20 |
| AT3G18260.1 | Reticulon family protein | - | Sampling day | -0.05 |
| AT1G35550.1 | elongation factor Tu C-terminal domain-containing protein | - | Sampling day | -0.07 |
| AT3G45770.1 | Polyketide synthase, enoylreductase family protein | - | Sampling day | -0.28 |

| | | | | |
|---|---|---|---|---|
| AT3G51270.1 | protein serine/threonine kinases;ATP binding;catalytics | - | Sampling day | -0.17 |
| AT3G60965.1 | transposable element gene | - | Sampling day | -0.13 |
| AT5G36775.1 | transposable element gene | - | Sampling day | -0.16 |
| AT3G55890.2 | Yippee family putative zinc-binding protein | - | Sampling day | -0.06 |
| AT5G37040.1 | F-box family protein | - | Sampling day | -0.13 |
| AT1G54530.1 | Calcium-binding EF hand family protein | - | Sampling day | -0.13 |
| AT5G61530.4 | small G protein family protein / RhoGAP family protein | - | Sampling day | -0.22 |
| AT3G52040.1 | NULL | - | Sampling day | -0.06 |
| AT3G26630.1 | Tetratricopeptide repeat (TPR)-like superfamily protein | - | Sampling day | -0.16 |
| AT3G17330.1 | evolutionarily conserved C-terminal region 6 | ECT6 | Sampling day | -0.15 |
| AT3G12670.1 | CTP synthase family protein | emb2742 | Sampling day | -0.21 |
| AT2G20190.1 | CLIP-associated protein | CLASP | Sampling day | -0.09 |
| AT4G39030.1 | MATE efflux family protein | EDS5 | Sampling day | -0.13 |
| AT1G74310.1 | heat shock protein 101 | HSP101 | Sampling day | -0.18 |
| AT5G18840.1 | Major facilitator superfamily protein | - | Sampling day | -0.11 |
| AT5G54340.1 | C2H2 and C2HC zinc fingers superfamily protein | - | Sampling day | -0.06 |
| AT1G57200.1 | pre-tRNA | - | Sampling day | -0.21 |
| AT4G09470.1 | NULL | - | Sampling day | -0.11 |
| AT3G13670.1 | Protein kinase family protein | - | Sampling day | -0.20 |
| AT1G78160.2 | pumilio 7 | PUM7 | Sampling day | -0.09 |
| AT2G32140.1 | transmembrane receptors | - | Sampling day | -0.15 |
| AT3G46484.1 | transposable element gene | - | Sampling day | -0.21 |
| AT5G15745.1 | | - | Sampling day | -0.37 |
| AT2G25340.1 | vesicle-associated membrane protein 712 | VAMP712 | Sampling day | -0.10 |
| AT5G03715.1 | | - | Sampling day | -0.06 |
| AT1G23590.2 | Domain of unknown function DUF220 | - | Sampling day | -0.19 |
| AT1G76990.3 | ACT domain repeat 3 | ACR3 | Sampling day | -0.15 |
| AT3G03065.1 | | - | Sampling day | -0.11 |
| AT5G02760.1 | Protein phosphatase 2C family protein | - | Sampling day | -0.20 |
| AT1G72930.1 | toll/interleukin-1 receptor-like | TIR | Sampling day | -0.22 |
| AT1G30600.1 | Subtilase family protein | - | Sampling day | -0.23 |
| AT2G26460.1 | RED family protein | SMU2 | Sampling day | -0.16 |
| AT3G43490.1 | Zinc knuckle (CCHC-type) family protein | - | Sampling day | -0.29 |
| AT3G30340.1 | nodulin MtN21 /EamA-like transporter family protein | UMAMIT32 | Sampling day | -0.16 |
| AT1G15940.1 | Tudor/PWWP/MBT superfamily protein | - | Sampling day | -0.10 |
| AT3G23820.1 | UDP-D-glucuronate 4-epimerase 6 | GAE6 | Sampling day | -0.26 |
| AT4G30310.4 | FGGY family of carbohydrate kinase | - | Sampling day | -0.16 |
| AT3G19940.1 | Major facilitator superfamily protein | - | Sampling day | -0.15 |
| AT2G09290.1 | | - | Sampling day | -0.20 |
| AT1G78810.3 | NULL | - | Sampling day | -0.11 |
| AT1G51120.1 | AP2/B3 transcription factor family protein | - | Sampling day | -0.11 |
| AT2G23530.1 | Zinc-finger domain of monoamine-oxidase A repressor R1 | - | Sampling day | -0.31 |
| AT5G42340.1 | Plant U-Box 15 | PUB15 | Sampling day | -0.50 |
| AT3G61690.2 | nucleotidyltransferases | - | Sampling day | -0.47 |
| AT1G49010.1 | Duplicated homeodomain-like superfamily protein | - | Sampling day | -0.18 |
| AT3G11690.1 | NULL | - | Sampling day | -0.08 |
| AT5G27882.1 | transposable element gene | - | Sampling day | -0.08 |
| AT1G14220.1 | Ribonuclease T2 family protein | - | Sampling day | -0.24 |
| AT1G67210.3 | Proline-rich spliceosome-associated (PSP) family protein / zinc knuckle (CCHC-type) family protein | - | Sampling day | -0.17 |
| AT4G36980.2 | NULL | - | Sampling day | -0.31 |
| AT5G33381.1 | transposable element gene | - | Sampling day | -0.27 |
| AT5G51600.1 | Microtubule associated protein (MAP65/ASE1) family protein | PLE | Sampling day | -0.24 |
| AT5G19800.1 | hydroxyproline-rich glycoprotein family protein | - | Sampling day | -0.39 |
| AT2G29280.3 | NULL | - | Sampling day | -0.15 |
| AT1G63500.1 | Protein kinase protein with tetratricopeptide repeat domain | BSK7 | Sampling day | -0.16 |
| AT4G16807.1 | NULL | - | Sampling day | -0.62 |
| AT4G09420.1 | Disease resistance protein (TIR-NBS class) | - | Sampling day | -0.13 |
| AT1G52010.1 | transposable element gene | - | Sampling day | -0.51 |
| AT5G41560.2 | NULL | - | Sampling day | -0.22 |
| AT2G42890.2 | MEI2-like 2 | ML2 | Sampling day | -0.19 |
| AT5G47077.1 | low-molecular-weight cysteine-rich 6 | LCR6 | Sampling day | -0.26 |
| AT4G24840.1 | NULL | - | Sampling day | -0.07 |
| AT5G44250.1 | Protein of unknown function DUF829, transmembrane 53 | - | Sampling day | -0.12 |
| AT5G17640.1 | Protein of unknown function (DUF1005) | ASG1 | Sampling day | -0.16 |
| AT1G31330.1 | photosystem I subunit F | PSAF | Sampling day | -0.10 |
| AT3G04120.1 | glyceraldehyde-3-phosphate dehydrogenase C subunit 1 | GAPC1 | Sampling day | -0.14 |
| AT3G08235.1 | | - | Sampling day | -0.15 |
| AT5G32950.1 | transposable element gene | - | Sampling day | -0.09 |
| AT5G23630.1 | phosphate deficiency response 2 | PDR2 | Sampling day | -0.20 |
| AT5G61580.1 | phosphofructokinase 4 | PFK4 | Sampling day | -0.16 |
| AT2G12465.1 | low-molecular-weight cysteine-rich 50 | LCR50 | Sampling day | -0.15 |
| AT5G09555.1 | | - | Sampling day | -0.10 |
| AT1G42610.1 | transposable element gene | - | Sampling day | -0.07 |
| AT5G23590.1 | DNAJ heat shock N-terminal domain-containing protein | - | Sampling day | -0.06 |
| ATMG09440.1 | | - | Sampling day | -0.13 |

| | | | | |
|---|---|---|---|---|
| AT2G44650.1 | chloroplast chaperonin 10 | CHL-CPN10 | Sampling day | -0.19 |
| AT4G03039.1 | MIR826a; miRNA | MIR826A | Sampling day | -0.17 |
| AT5G50780.2 | Histidine kinase-, DNA gyrase B-, and HSP90-like ATPase family | - | Sampling day | -0.18 |
| AT1G70410.3 | beta carbonic anhydrase 4 | BCA4 | Sampling day | -0.15 |
| AT1G05065.1 | CLAVATA3/ESR-RELATED 20 | CLE20 | Sampling day | -0.12 |
| AT2G47100.1 | pre-tRNA | - | Sampling day | -0.21 |
| AT5G03725.1 | | - | Sampling day | -0.12 |
| AT1G28930.1 | pre-tRNA | - | Sampling day | -0.18 |
| AT2G43930.2 | Protein kinase superfamily protein | - | Sampling day | -0.16 |
| AT5G27760.2 | Hypoxia-responsive family protein | - | Sampling day | -0.17 |
| AT5G07740.1 | actin binding | - | Sampling day | -0.19 |
| AT2G09640.1 | | - | Sampling day | -0.12 |
| AT3G02775.1 | | - | Sampling day | -0.17 |
| AT5G62180.1 | carboxyesterase 20 | CXE20 | Sampling day | -0.06 |
| AT1G10865.1 | NULL | - | Sampling day | -0.16 |
| AT2G15890.1 | maternal effect embryo arrest 14 | MEE14 | Sampling day | -0.14 |
| AT5G64060.1 | NAC domain containing protein 103 | NAC103 | Sampling day | -0.14 |
| AT5G50510.1 | Molecular chaperone Hsp40/DnaJ family protein | - | Sampling day | -0.09 |
| AT1G13270.1 | methionine aminopeptidase 1B | MAP1C | Sampling day | -0.23 |
| AT4G18860.1 | NULL | - | Sampling day | -0.36 |
| AT4G02735.1 | | - | Sampling day | -0.06 |
| AT5G09790.1 | ARABIDOPSIS TRITHORAX-RELATED PROTEIN 5 | ATXR5 | Sampling day | -0.16 |
| AT3G62120.3 | Class II aaRS and biotin synthetases superfamily protein | - | Sampling day | -0.17 |
| AT2G47460.1 | myb domain protein 12 | MYB12 | Sampling day | -0.12 |
| AT1G66940.3 | protein kinase-related | - | Sampling day | -0.08 |
| AT1G74290.1 | alpha/beta-Hydrolases superfamily protein | - | Sampling day | -0.07 |
| AT2G08845.1 | | - | Sampling day | -0.11 |
| AT1G04907.1 | | U1-10 | Sampling day | -0.16 |
| AT1G50180.1 | NB-ARC domain-containing disease resistance protein | - | Sampling day | -0.27 |
| AT5G35555.1 | transposable element gene | - | Sampling day | -0.07 |
| AT1G06130.1 | glyoxalase 2-4 | GLX2-4 | Sampling day | -0.21 |
| AT3G30830.1 | transposable element gene | - | Sampling day | -0.15 |
| AT5G56440.1 | F-box/RNI-like/FBD-like domains-containing protein | - | Sampling day | -0.16 |
| AT1G58684.1 | Ribosomal protein S5 family protein | - | Sampling day | -0.17 |
| AT3G12960.1 | NULL | SMP1 | Sampling day | -0.24 |
| AT2G03480.3 | QUASIMODO2 LIKE 2 | QUL2 | Sampling day | -0.09 |
| AT5G44780.1 | NULL | MORF4 | Sampling day | -0.13 |
| AT2G35570.1 | NULL | - | Sampling day | -0.34 |
| AT1G10500.1 | chloroplast-localized ISCA-like protein | CPISCA | Sampling day | -0.18 |
| AT5G17230.3 | PHYTOENE SYNTHASE | PSY | Sampling day | -0.20 |
| AT1G50670.1 | OTU-like cysteine protease family protein | - | Sampling day | -0.17 |
| AT5G40440.2 | mitogen-activated protein kinase kinase 3 | MKK3 | Sampling day | -0.15 |
| AT2G30260.2 | U2 small nuclear ribonucleoprotein B | U2B" | Sampling day | -0.17 |
| AT4G05097.1 | | - | Sampling day | -0.24 |
| AT1G06937.1 | | - | Sampling day | -0.10 |
| AT5G08300.1 | Succinyl-CoA ligase, alpha subunit | - | Sampling day | -0.16 |
| AT5G28020.1 | cysteine synthase D2 | CYSD2 | Sampling day | -0.14 |
| AT1G23620.1 | NULL | - | Sampling day | -0.10 |
| AT5G66640.3 | DA1-related protein 3 | DAR3 | Sampling day | -0.23 |
| AT3G26390.1 | NULL | - | Sampling day | -0.15 |
| AT2G35280.1 | F-box family protein | - | Sampling day | -0.18 |
| AT4G36890.1 | Nucleotide-diphospho-sugar transferases superfamily protein | IRX14 | Sampling day | -0.25 |
| AT5G50320.1 | radical SAM domain-containing protein / GCN5-related N-acetyltransferase (GNAT) family protein | ELO3 | Sampling day | -0.50 |
| AT4G21070.1 | breast cancer susceptibility1 | BRCA1 | Sampling day | -0.25 |
| AT3G03900.2 | adenosine-5'-phosphosulfate (APS) kinase 3 | APK3 | Sampling day | -0.08 |
| AT3G62040.1 | Haloacid dehalogenase-like hydrolase (HAD) superfamily protein | - | Sampling day | -0.12 |
| AT2G18560.1 | UDP-Glycosyltransferase superfamily protein | - | Sampling day | -0.24 |
| AT1G17180.1 | glutathione S-transferase TAU 25 | GSTU25 | Sampling day | -0.09 |
| AT3G57645.1 | U2.2; snRNA | U2.2 | Sampling day | -0.06 |
| AT1G05673.1 | | U6atac2 | Sampling day | -0.14 |
| AT5G07455.1 | | - | Sampling day | -0.07 |
| AT2G07737.1 | transposable element gene | - | Sampling day | -0.06 |
| AT4G17850.1 | NULL | - | Sampling day | -0.24 |
| AT4G07585.1 | | - | Sampling day | -0.11 |
| AT4G10320.1 | tRNA synthetase class I (I, L, M and V) family protein | - | Sampling day | -0.06 |
| AT1G08523.1 | | - | Sampling day | -0.21 |
| AT2G00650.1 | | - | Sampling day | -0.09 |
| AT4G34570.2 | thymidylate synthase 2 | THY-2 | Sampling day | -0.09 |
| AT3G48300.2 | cytochrome P450, family 71, subfamily A, polypeptide 23 | CYP71A23 | Sampling day | -0.10 |
| AT5G02890.1 | HXXXD-type acyl-transferase family protein. Overexpression causes a glossy phenotype and defects in cuticular wax deposition. OE plants are blocked in very long chain fatty acid elongation (C28 and higher). | - | Sampling day | -0.30 |
| AT5G07820.1 | Plant calmodulin-binding protein-related | - | Sampling day | -0.11 |
| AT3G07730.1 | NULL | - | Sampling day | -0.07 |
| AT1G26180.2 | NULL | - | Sampling day | -0.18 |

| | | | | |
|---|---|---|---|---|
| AT1G78940.3 | Protein kinase protein with adenine nucleotide alpha hydrolases-like | - | Sampling day | -0.05 |
| AT5G42825.1 | NULL | - | Sampling day | -0.30 |
| AT3G11440.2 | myb domain protein 65 | MYB65 | Sampling day | -0.18 |
| AT3G15940.2 | UDP-Glycosyltransferase superfamily protein | - | Sampling day | -0.10 |
| AT5G27160.1 | transposable element gene | - | Sampling day | -0.08 |
| AT1G37110.1 | transposable element gene | - | Sampling day | -0.19 |
| AT5G28260.1 | transposable element gene | - | Sampling day | -0.11 |
| AT4G05073.1 | transposable element gene | - | Sampling day | -0.13 |
| AT2G29570.1 | proliferating cell nuclear antigen 2 | PCNA2 | Sampling day | -0.18 |
| AT4G03125.1 | | - | Sampling day | -0.13 |
| AT5G43170.1 | zinc-finger protein 3 | ZF3 | Sampling day | -0.28 |
| AT1G66245.1 | NULL | - | Sampling day | -0.25 |
| AT5G34856.1 | transposable element gene | - | Sampling day | -0.09 |
| AT1G51520.1 | RNA-binding (RRM/RBD/RNP motifs) family protein | - | Sampling day | -0.13 |
| AT3G21450.1 | Protein kinase superfamily protein | - | Sampling day | -0.30 |
| AT2G35650.2 | cellulose synthase like | CSLA07 | Sampling day | -0.33 |
| AT2G39230.1 | LATERAL ORGAN JUNCTION | LOJ | Sampling day | -0.12 |
| AT2G06550.1 | transposable element gene | - | Sampling day | -0.16 |
| AT1G04540.1 | Calcium-dependent lipid-binding (CaLB domain) family protein | - | Sampling day | -0.06 |
| AT1G70000.1 | myb-like transcription factor family protein | - | Sampling day | -0.10 |
| AT1G76260.1 | DWD (DDB1-binding WD40 protein) hypersensitive to ABA 2 | DWA2 | Sampling day | -0.22 |
| AT3G06325.1 | | - | Sampling day | -0.17 |
| AT1G06360.2 | Fatty acid desaturase family protein | - | Sampling day | -0.18 |
| AT1G07513.1 | | - | Sampling day | -0.09 |
| AT1G05603.1 | | - | Sampling day | -0.13 |
| AT4G04500.3 | cysteine-rich RLK (RECEPTOR-like protein kinase) 37 | CRK37 | Sampling day | -0.18 |
| AT5G42310.1 | Pentatricopeptide repeat (PPR-like) superfamily protein | - | Sampling day | -0.18 |
| AT1G09647.1 | | - | Sampling day | -0.18 |
| AT2G14350.1 | transposable element gene | - | Sampling day | -0.19 |
| AT1G50050.2 | CAP (Cysteine-rich secretory proteins, Antigen 5, and Pathogenesis-related 1 protein) superfamily protein | - | Sampling day | -0.39 |
| AT3G03995.1 | | - | Sampling day | -0.11 |
| AT2G04560.1 | transferases, transferring glycosyl groups | LpxB | Sampling day | -0.28 |
| AT1G73480.1 | alpha/beta-Hydrolases superfamily protein | - | Sampling day | -0.07 |
| AT3G13677.2 | NULL | - | Sampling day | -0.19 |
| AT3G05060.1 | NOP56-like pre RNA processing ribonucleoprotein | - | Sampling day | -0.37 |
| AT1G15530.1 | Concanavalin A-like lectin protein kinase family protein | - | Sampling day | -0.17 |
| AT4G39400.1 | Leucine-rich receptor-like protein kinase family protein | BRI1 | Sampling day | -0.23 |
| AT3G33011.1 | transposable element gene | - | Sampling day | -0.19 |
| AT3G62610.1 | myb domain protein 11 | MYB11 | Sampling day | -0.18 |
| AT4G09310.1 | SPla/RYanodine receptor (SPRY) domain-containing protein | - | Sampling day | -0.37 |
| AT3G30820.1 | Arabidopsis retrotransposon ORF-1 protein | - | Sampling day | -0.10 |
| AT2G16410.1 | transposable element gene | - | Sampling day | -0.08 |
| AT1G51840.2 | protein kinase-related | - | Sampling day | -0.30 |
| AT5G64040.1 | photosystem I reaction center subunit PSI-N, chloroplast, putative / PSI-N, putative (PSAN) | PSAN | Sampling day | -0.19 |
| AT3G16850.1 | Pectin lyase-like superfamily protein | - | Sampling day | -0.08 |
| AT3G54770.3 | RNA-binding (RRM/RBD/RNP motifs) family protein | ARP1 | Sampling day | -0.06 |
| AT5G17380.1 | Thiamine pyrophosphate dependent pyruvate decarboxylase family | - | Sampling day | -0.28 |
| AT1G16445.1 | S-adenosyl-L-methionine-dependent methyltransferases superfamily | - | Sampling day | -0.28 |
| AT3G06085.1 | | - | Sampling day | -0.19 |
| AT1G11560.1 | Oligosaccharyltransferase complex/magnesium transporter family | - | Sampling day | -0.21 |
| AT1G63960.2 | Copper transport protein family | - | Sampling day | -0.09 |
| AT5G09795.1 | NULL | - | Sampling day | -0.18 |
| AT5G42530.1 | NULL | - | Sampling day | -0.11 |
| AT2G08100.1 | | - | Sampling day | -0.21 |
| AT5G66607.1 | NULL | - | Sampling day | -0.14 |
| AT3G01740.1 | Mitochondrial ribosomal protein L37 | - | Sampling day | -0.06 |
| AT4G31650.2 | Transcriptional factor B3 family protein | - | Sampling day | -0.10 |
| AT5G09995.3 | NULL | - | Sampling day | -0.21 |
| AT1G66740.1 | ASF1 like histone chaperone | SGA2 | Sampling day | -0.20 |
| AT5G22570.1 | WRKY DNA-binding protein 38 | WRKY38 | Sampling day | -0.35 |
| AT4G12270.1 | Copper amine oxidase family protein | - | Sampling day | -0.21 |
| AT1G04773.1 | | - | Sampling day | -0.13 |
| AT2G46680.1 | homeobox 7 | HB-7 | Sampling day | -0.26 |
| AT4G02485.2 | 2-oxoglutarate (2OG) and Fe(II)-dependent oxygenase superfamily | - | Sampling day | -0.09 |
| AT5G44290.3 | Protein kinase superfamily protein | - | Sampling day | -0.18 |
| AT4G08825.1 | | - | Sampling day | -0.07 |
| AT2G21160.1 | Translocon-associated protein (TRAP), alpha subunit | - | Sampling day | -0.22 |
| AT5G14760.1 | L-aspartate oxidase | AO | Sampling day | -0.20 |
| AT4G36970.1 | Remorin family protein | - | Sampling day | -0.23 |
| AT5G53830.1 | VQ motif-containing protein | - | Sampling day | -0.06 |
| AT4G12230.1 | alpha/beta-Hydrolases superfamily protein | - | Sampling day | -0.36 |
| AT5G11530.2 | embryonic flower 1 (EMF1) | EMF1 | Sampling day | -0.11 |
| AT4G14510.1 | CRM family member 3B | CFM3B | Sampling day | -0.25 |
| AT1G12040.1 | leucine-rich repeat/extensin 1 | LRX1 | Sampling day | -0.19 |

| | | | | |
|---|---|---|---|---|
| AT5G09360.1 | laccase 14 | LAC14 | Sampling day | -0.38 |
| AT3G14050.1 | RELA/SPOT homolog 2 | RSH2 | Sampling day | -0.27 |
| AT1G31210.1 | transposable element gene | - | Sampling day | -0.13 |
| AT3G51420.1 | strictosidine synthase-like 4 | SSL4 | Sampling day | -0.11 |
| AT2G24060.1 | Translation initiation factor 3 protein | - | Sampling day | -0.17 |
| AT1G45904.1 | NULL | - | Sampling day | -0.10 |
| AT5G20430.1 | Mob1/phocein family protein | - | Sampling day | -0.21 |
| AT5G23650.1 | Homeodomain-like transcriptional regulator | - | Sampling day | -0.14 |
| AT4G08975.1 | | - | Sampling day | -0.12 |
| AT2G29925.1 | | - | Sampling day | -0.46 |
| AT3G62528.1 | NULL | - | Sampling day | -0.15 |
| AT1G42210.1 | transposable element gene | - | Sampling day | -0.06 |
| AT1G12150.2 | Plant protein of unknown function (DUF827) | - | Sampling day | -0.06 |
| AT1G60930.1 | RECQ helicase L4B | RECQ4B | Sampling day | -0.17 |
| AT3G60240.3 | eukaryotic translation initiation factor 4G | EIF4G | Sampling day | -0.14 |
| AT1G08857.1 | | - | Sampling day | -0.08 |
| AT3G08940.2 | light harvesting complex photosystem II | LHCB4.2 | Sampling day | -0.65 |
| AT5G28100.1 | transposable element gene | - | Sampling day | -0.18 |
| AT3G11960.3 | Cleavage and polyadenylation specificity factor (CPSF) A subunit | - | Sampling day | -0.17 |
| ATCG00750.1 | ribosomal protein S11 | RPS11 | Sampling day | -0.10 |
| AT1G69670.1 | cullin 3B | CUL3B | Sampling day | -0.08 |
| AT3G06180.1 | Ribosomal protein L34e superfamily protein | - | Sampling day | -0.19 |
| AT4G11230.1 | Riboflavin synthase-like superfamily protein | - | Sampling day | -0.21 |
| AT1G61390.2 | S-locus lectin protein kinase family protein | - | Sampling day | -0.19 |
| AT1G43330.1 | Homeodomain-like superfamily protein | - | Sampling day | -0.06 |
| AT4G21280.1 | photosystem II subunit QA | PSBQA | Sampling day | -0.16 |
| AT5G67265.1 | NULL | - | Sampling day | -0.07 |
| AT3G18220.1 | Phosphatidic acid phosphatase (PAP2) family protein | LPP4 | Sampling day | -0.24 |
| AT5G20165.3 | NULL | - | Sampling day | -0.17 |
| AT3G30250.1 | transposable element gene | - | Sampling day | -0.14 |
| AT1G26976.1 | NULL | - | Sampling day | -0.20 |
| AT3G43850.2 | NULL | - | Sampling day | -0.18 |
| AT5G09925.1 | | - | Sampling day | -0.25 |
| AT1G65365.1 | NULL | - | Sampling day | -0.20 |
| ATMG00080.1 | ribosomal protein L16 | RPL16 | Sampling day | -0.13 |
| AT3G50925.1 | Defensin-like (DEFL) family protein | - | Sampling day | -0.24 |
| AT1G02820.1 | Late embryogenesis abundant 3 (LEA3) family protein | LEA3 | Sampling day | -0.09 |
| AT2G34980.1 | phosphatidylinositolglycan synthase family protein | SETH1 | Sampling day | -0.12 |
| AT2G10555.1 | transposable element gene | - | Sampling day | -0.06 |
| AT5G32293.1 | transposable element gene | - | Sampling day | -0.22 |
| AT3G59415.1 | pre-tRNA | - | Sampling day | -0.21 |
| AT1G16380.1 | Cation/hydrogen exchanger family protein | ATCHX1 | Sampling day | -0.36 |
| AT1G30400.2 | multidrug resistance-associated protein 1 | ABCC1 | Sampling day | -0.25 |
| AT5G65370.1 | ENTH/ANTH/VHS superfamily protein | - | Sampling day | -0.19 |
| AT5G24870.2 | RING/U-box superfamily protein | - | Sampling day | -0.07 |
| AT3G17600.1 | indole-3-acetic acid inducible 31 | IAA31 | Sampling day | -0.21 |
| AT5G55400.1 | Actin binding Calponin homology (CH) domain-containing protein | - | Sampling day | -0.16 |
| AT5G17140.1 | Cysteine proteinases superfamily protein | - | Sampling day | -0.08 |
| AT3G16030.3 | lectin protein kinase family protein | CES101 | Sampling day | -0.10 |
| AT5G38660.1 | acclimation of photosynthesis to environment | APE1 | Sampling day | -0.24 |
| AT5G55610.2 | NULL | - | Sampling day | -0.13 |
| AT1G28400.1 | NULL | - | Sampling day | -0.08 |
| AT3G61430.1 | plasma membrane intrinsic protein 1A | PIP1A | Sampling day | -0.09 |
| AT5G50020.3 | DHHC-type zinc finger family protein | - | Sampling day | -0.15 |
| AT1G05127.1 | | - | Sampling day | -0.08 |
| AT4G06742.1 | transposable element gene | - | Sampling day | -0.13 |
| AT4G34120.1 | Cystathionine beta-synthase (CBS) family protein | LEJ1 | Sampling day | -0.09 |
| AT1G67240.1 | transposable element gene | - | Sampling day | -0.15 |
| AT1G08370.1 | decapping 1 | DCP1 | Sampling day | -0.24 |
| AT5G01940.2 | eukaryotic translation initiation factor 2B family protein / eIF-2B family | - | Sampling day | -0.12 |
| AT3G26230.1 | cytochrome P450, family 71, subfamily B, polypeptide 24 | CYP71B24 | Sampling day | -0.16 |
| AT4G02800.1 | NULL | - | Sampling day | -0.13 |
| AT4G28480.1 | DNAJ heat shock family protein | - | Sampling day | -0.09 |
| AT3G47150.1 | F-box and associated interaction domains-containing protein | - | Sampling day | -0.07 |
| AT4G11470.2 | cysteine-rich RLK (RECEPTOR-like protein kinase) 31 | CRK31 | Sampling day | -0.08 |
| AT1G36770.1 | transposable element gene | - | Sampling day | -0.13 |
| AT5G06310.1 | Nucleic acid-binding, OB-fold-like protein | AtPOT1b | Sampling day | -0.09 |
| AT1G48095.1 | NULL | - | Sampling day | -0.20 |
| AT4G06815.1 | | - | Sampling day | -0.21 |
| AT5G45560.2 | Pleckstrin homology (PH) domain-containing protein / lipid-binding START domain-containing protein | - | Sampling day | -0.10 |
| AT2G45890.1 | RHO guanyl-nucleotide exchange factor 4 | ROPGEF4 | Sampling day | -0.09 |
| AT3G13432.1 | NULL | - | Sampling day | -0.20 |
| AT5G08375.1 | | - | Sampling day | -0.20 |
| AT1G09745.1 | | - | Sampling day | -0.18 |
| AT4G18170.1 | WRKY DNA-binding protein 28 | WRKY28 | Sampling day | -0.22 |

| | | | | |
|---|---|---|---|---|
| AT3G01545.1 | | - | Sampling day | -0.15 |
| AT3G06045.1 | | - | Sampling day | -0.14 |
| AT2G19802.1 | NULL | - | Sampling day | -0.22 |
| AT3G17540.1 | F-box and associated interaction domains-containing protein | - | Sampling day | -0.18 |
| AT3G06070.1 | NULL | - | Sampling day | -0.33 |
| AT2G34208.1 | MIR399F; miRNA | MIR399F | Sampling day | -0.31 |
| AT1G50810.1 | transposable element gene | - | Sampling day | -0.15 |
| AT5G41680.2 | Protein kinase superfamily protein | - | Sampling day | -0.26 |
| AT3G06750.1 | hydroxyproline-rich glycoprotein family protein | - | Sampling day | -0.07 |
| AT3G31406.1 | transposable element gene | - | Sampling day | -0.21 |
| AT4G08385.1 | | - | Sampling day | -0.11 |
| AT5G42840.1 | Cysteine/Histidine-rich C1 domain family protein | - | Sampling day | -0.10 |
| AT3G44713.1 | NULL | - | Sampling day | -0.17 |
| AT4G40030.4 | Histone superfamily protein | - | Sampling day | -0.15 |
| AT2G17040.1 | NAC domain containing protein 36 | NAC036 | Sampling day | -0.22 |
| AT4G39020.1 | SH3 domain-containing protein | - | Sampling day | -0.23 |
| AT1G30140.2 | NULL | - | Sampling day | -0.19 |
| AT2G15960.1 | NULL | - | Sampling day | -0.21 |
| AT1G52060.1 | Mannose-binding lectin superfamily protein | - | Sampling day | -0.21 |
| AT2G42390.1 | protein kinase C substrate, heavy chain-related | - | Sampling day | -0.18 |
| AT3G46950.1 | Mitochondrial transcription termination factor family protein | - | Sampling day | -0.07 |
| AT3G06910.1 | UB-like protease 1A | ULP1A | Sampling day | -0.19 |
| AT5G01005.1 | | - | Sampling day | -0.11 |
| AT2G07320.1 | transposable element gene | - | Sampling day | -0.10 |
| AT5G17970.1 | Disease resistance protein (TIR-NBS-LRR class) family | - | Sampling day | -0.08 |
| AT4G06618.1 | transposable element gene | - | Sampling day | -0.11 |
| AT2G04440.1 | MutT/nudix family protein | - | Sampling day | -0.15 |
| AT1G08885.1 | | - | Sampling day | -0.11 |
| AT3G08740.1 | elongation factor P (EF-P) family protein | - | Sampling day | -0.07 |
| AT2G09680.1 | | - | Sampling day | -0.13 |
| AT4G19640.3 | Ras-related small GTP-binding family protein | ARA7 | Sampling day | -0.20 |
| AT1G45150.1 | NULL | - | Sampling day | -0.11 |
| AT3G16310.1 | mitotic phosphoprotein N' end (MPPN) family protein | - | Sampling day | -0.57 |
| AT5G56600.1 | profilin 3 | PRF3 | Sampling day | -0.21 |
| AT2G37555.1 | other RNA | - | Sampling day | -0.24 |
| AT3G08695.1 | | U2-6 | Sampling day | -0.14 |
| AT2G21360.1 | pre-tRNA | - | Sampling day | -0.15 |
| AT4G18820.1 | AAA-type ATPase family protein | - | Sampling day | -0.18 |
| AT3G59480.1 | pfkB-like carbohydrate kinase family protein | - | Sampling day | -0.21 |
| AT2G02740.1 | ssDNA-binding transcriptional regulator | WHY3 | Sampling day | -0.09 |
| AT5G25390.2 | Integrase-type DNA-binding superfamily protein | SHN3 | Sampling day | -0.21 |
| AT4G34280.1 | transducin family protein / WD-40 repeat family protein | - | Sampling day | -0.14 |
| AT2G47935.1 | | - | Sampling day | -0.11 |
| AT4G31248.1 | other RNA | - | Sampling day | -0.14 |
| AT4G12735.1 | NULL | - | Sampling day | -0.21 |
| AT5G19470.2 | nudix hydrolase homolog 24 | NUDT24 | Sampling day | -0.21 |
| AT5G04980.1 | DNAse I-like superfamily protein | - | Sampling day | -0.25 |
| AT4G19940.1 | F-box and associated interaction domains-containing protein | - | Sampling day | -0.08 |
| AT2G11115.1 | transposable element gene | - | Sampling day | -0.13 |
| AT2G24220.1 | purine permease 5 | PUP5 | Sampling day | -0.17 |
| AT2G34560.2 | P-loop containing nucleoside triphosphate hydrolases superfamily | CCP1 | Sampling day | -0.11 |
| AT1G70020.1 | Protein of unknown function (DUF1163) | - | Sampling day | -0.16 |
| AT1G06925.1 | NULL | - | Sampling day | -0.12 |
| AT3G25570.2 | Adenosylmethionine decarboxylase family protein | - | Sampling day | -0.08 |
| AT3G13226.1 | regulatory protein RecX family protein | - | Sampling day | -0.14 |
| AT4G29650.1 | Cytidine/deoxycytidylate deaminase family protein | - | Sampling day | -0.21 |
| AT3G58890.1 | RNI-like superfamily protein | - | Sampling day | -0.14 |
| AT5G03730.1 | Protein kinase superfamily protein | CTR1 | Sampling day | -0.14 |
| AT2G44680.3 | casein kinase II beta subunit 4 | CKB4 | Sampling day | -0.08 |
| AT2G15420.1 | myosin heavy chain-related | - | Sampling day | -0.20 |
| AT5G58310.1 | methyl esterase 18 | MES18 | Sampling day | -0.21 |
| AT4G07990.1 | Chaperone DnaJ-domain superfamily protein | - | Sampling day | -0.18 |
| AT1G32340.1 | NDR1/HIN1-like 8 | NHL8 | Sampling day | -0.14 |
| AT3G28216.1 | NULL | - | Sampling day | -0.06 |
| AT2G32080.2 | purin-rich alpha 1 | PUR ALPHA-1 | Sampling day | -0.07 |
| AT3G06380.2 | tubby-like protein 9 | TLP9 | Sampling day | -0.16 |
| AT4G20030.1 | RNA-binding (RRM/RBD/RNP motifs) family protein | - | Sampling day | -0.10 |
| AT5G38035.1 | transposable element gene | - | Sampling day | -0.11 |
| AT1G18070.1 | Translation elongation factor EF1A/initiation factor IF2gamma family | - | Sampling day | -0.15 |
| AT3G18165.1 | modifier of snc1,4 | MOS4 | Sampling day | -0.10 |
| AT1G68585.1 | NULL | - | Sampling day | -0.10 |
| AT2G25305.1 | Putative membrane lipoprotein | - | Sampling day | -0.08 |
| AT5G23100.1 | Protein of unknown function, DUF617 | - | Sampling day | -0.19 |
| AT4G35680.1 | Arabidopsis protein of unknown function (DUF241) | - | Sampling day | -0.07 |
| AT2G28490.1 | RmlC-like cupins superfamily protein | - | Sampling day | -0.14 |
| AT3G14920.1 | Peptide-N4-(N-acetyl-beta-glucosaminyl)asparagine amidase A protein | - | Sampling day | -0.20 |

| | | | | |
|---|---|---|---|---|
| AT1G09943.1 | | - | 1day perior to sampling | 0.18 |
| AT2G07719.1 | Putative membrane lipoprotein | - | 1day perior to sampling | 0.06 |
| AT2G32950.1 | Transducin/WD40 repeat-like superfamily protein | COP1 | 1day perior to sampling | 0.22 |
| AT4G34970.1 | actin depolymerizing factor 9 | ADF9 | 1day perior to sampling | 0.12 |
| AT3G51880.1 | high mobility group B1 | HMGB1 | 1day perior to sampling | 0.31 |
| AT4G05583.2 | transposable element gene | - | 1day perior to sampling | 0.24 |
| AT3G27480.1 | Cysteine/Histidine-rich C1 domain family protein | - | 1day perior to sampling | 0.16 |
| AT4G30060.1 | Core-2/I-branching beta-1,6-N-acetylglucosaminyltransferase family | - | 1day perior to sampling | 0.10 |
| AT1G41798.1 | transposable element gene | - | 1day perior to sampling | 0.29 |
| AT2G09500.1 | | - | 1day perior to sampling | 0.21 |
| AT5G46150.1 | LEM3 (ligand-effect modulator 3) family protein / CDC50 family | - | 1day perior to sampling | 0.22 |
| AT2G24090.1 | Ribosomal protein L35 | PRPL35 | 1day perior to sampling | 0.07 |
| AT3G42190.1 | transposable element gene | - | 1day perior to sampling | 0.21 |
| AT2G47680.1 | zinc finger (CCCH type) helicase family protein | - | 1day perior to sampling | 0.09 |
| AT1G09823.1 | | - | 1day perior to sampling | 0.11 |
| AT2G05335.1 | SCR-like 15 | SCRL15 | 1day perior to sampling | 0.10 |
| AT3G09100.2 | mRNA capping enzyme family protein | - | 1day perior to sampling | 0.13 |
| AT2G23445.1 | | - | 1day perior to sampling | 0.28 |
| AT4G01510.2 | Arv1-like protein | ARV2 | 1day perior to sampling | 0.25 |
| AT4G06728.1 | transposable element gene | - | 1day perior to sampling | 0.21 |
| AT2G37450.3 | nodulin MtN21 /EamA-like transporter family protein | UMAMIT13 | 1day perior to sampling | 0.20 |
| AT3G59960.2 | histone-lysine N-methyltransferase ASHH4 | ASHH4 | 1day perior to sampling | 0.29 |
| AT4G27550.1 | trehalose-6-phosphatase synthase S4 | TPS4 | 1day perior to sampling | 0.08 |
| AT3G62740.2 | beta glucosidase 7 | BGLU7 | 1day perior to sampling | 0.15 |
| AT2G06240.1 | transposable element gene | - | 1day perior to sampling | 0.27 |
| AT2G04290.1 | transposable element gene | - | 1day perior to sampling | 0.40 |
| AT1G13210.1 | autoinhibited Ca2+/ATPase II | ACA.l | 1day perior to sampling | 0.12 |
| AT3G08055.1 | | - | 1day perior to sampling | 0.28 |
| AT5G55850.3 | RPM1-interacting protein 4 (RIN4) family protein | NOI | 1day perior to sampling | 0.09 |
| AT5G47360.1 | Tetratricopeptide repeat (TPR)-like superfamily protein | - | 1day perior to sampling | 0.08 |
| AT3G10572.2 | 3-phosphoinositide-dependent protein kinase-1, putative | APEM9 | 1day perior to sampling | 0.16 |
| AT5G36730.1 | F-box and associated interaction domains-containing protein | - | 1day perior to sampling | 0.29 |
| AT1G24180.1 | Thiamin diphosphate-binding fold (THDP-binding) superfamily protein | IAR4 | 1day perior to sampling | 0.29 |
| AT5G32017.1 | pre-tRNA | - | 1day perior to sampling | 0.22 |
| AT2G09150.1 | | - | 1day perior to sampling | 0.14 |
| AT2G33780.1 | VQ motif-containing protein | - | 1day perior to sampling | 0.32 |
| AT4G22217.1 | Arabidopsis defensin-like protein | - | 1day perior to sampling | 0.24 |
| AT5G51050.1 | Mitochondrial substrate carrier family protein | APC2 | 1day perior to sampling | 0.29 |
| AT1G61820.1 | beta glucosidase 46 | BGLU46 | 1day perior to sampling | 0.16 |
| AT2G46950.2 | cytochrome P450, family 709, subfamily B, polypeptide 2 | CYP709B2 | 1day perior to sampling | 0.17 |
| AT2G00370.1 | | - | 1day perior to sampling | 0.20 |
| AT5G63135.1 | NULL | - | 1day perior to sampling | 0.06 |
| AT1G09233.1 | | - | 1day perior to sampling | 0.30 |
| AT1G08377.1 | | - | 1day perior to sampling | 0.17 |
| AT4G25900.1 | Galactose mutarotase-like superfamily protein | - | 1day perior to sampling | 0.20 |
| AT1G49700.3 | Plant protein 1589 of unknown function | - | 1day perior to sampling | 0.26 |
| AT4G01525.1 | transposable element gene | SADHU5-1 | 1day perior to sampling | 0.26 |
| AT2G31215.1 | basic helix-loop-helix (bHLH) DNA-binding superfamily protein | - | 1day perior to sampling | 0.13 |
| AT1G51740.2 | syntaxin of plants 81 | SYP81 | 1day perior to sampling | 0.26 |
| AT4G06684.1 | transposable element gene | - | 1day perior to sampling | 0.17 |
| AT5G14320.1 | Ribosomal protein S13/S18 family | EMB3137 | 1day perior to sampling | 0.20 |
| AT1G21475.1 | Protein of unknown function (DUF506) | - | 1day perior to sampling | 0.06 |
| AT3G23440.1 | embryo sac development arrest 6 | EDA6 | 1day perior to sampling | 0.08 |
| AT3G60610.1 | NULL | - | 1day perior to sampling | 0.18 |
| AT5G35870.1 | NULL | - | 1day perior to sampling | 0.30 |
| AT1G32760.1 | Glutaredoxin family protein | - | 1day perior to sampling | 0.22 |
| AT3G00540.1 | | - | 1day perior to sampling | 0.29 |
| AT1G28870.1 | pre-tRNA | - | 1day perior to sampling | 0.18 |
| AT1G22690.1 | Gibberellin-regulated family protein | - | 1day perior to sampling | 0.13 |
| AT5G18450.1 | Integrase-type DNA-binding superfamily protein | - | 1day perior to sampling | 0.12 |
| AT5G37730.1 | NULL | - | 1day perior to sampling | 0.09 |
| AT5G33255.1 | transposable element gene | - | 1day perior to sampling | 0.06 |
| AT2G32800.1 | protein kinase family protein | AP4.3A | 1day perior to sampling | 0.21 |
| AT4G37410.1 | cytochrome P450, family 81, subfamily F, polypeptide 4 | CYP81F4 | 1day perior to sampling | 0.16 |
| AT4G18203.1 | | - | 1day perior to sampling | 0.24 |
| AT1G17410.3 | Nucleoside diphosphate kinase family protein | - | 1day perior to sampling | 0.27 |
| AT1G05443.1 | | - | 1day perior to sampling | 0.22 |
| AT1G14010.1 | emp24/gp25L/p24 family/GOLD family protein | - | 1day perior to sampling | 0.21 |
| AT2G09325.1 | | - | 1day perior to sampling | 0.08 |
| AT1G02400.2 | gibberellin 2-oxidase 6 | GA2OX6 | 1day perior to sampling | 0.08 |
| AT3G20840.1 | Integrase-type DNA-binding superfamily protein | PLT1 | 1day perior to sampling | 0.17 |
| AT2G01818.1 | PLATZ transcription factor family protein | - | 1day perior to sampling | 0.19 |
| AT4G06547.1 | transposable element gene | - | 1day perior to sampling | 0.14 |
| AT5G05930.3 | guanylyl cyclase 1 | GC1 | 1day perior to sampling | 0.08 |
| AT4G07840.1 | transposable element gene | - | 1day perior to sampling | 0.36 |
| AT5G05570.2 | transducin family protein / WD-40 repeat family protein | - | 1day perior to sampling | 0.22 |

| | | | | |
|---|---|---|---|---|
| AT3G48490.1 | NULL | - | 1day perior to sampling | 0.34 |
| AT2G25070.2 | Protein phosphatase 2C family protein | - | 1day perior to sampling | 0.08 |
| AT2G30830.1 | 2-oxoglutarate (2OG) and Fe(II)-dependent oxygenase superfamily | - | 1day perior to sampling | 0.31 |
| AT3G22220.3 | hAT transposon superfamily | - | 1day perior to sampling | 0.28 |
| AT3G25225.1 | NULL | - | 1day perior to sampling | 0.25 |
| AT5G05980.2 | DHFS-FPGS homolog B | DFB | 1day perior to sampling | 0.09 |
| AT3G18610.1 | nucleolin like 2 | NUC-L2 | 1day perior to sampling | 0.08 |
| AT1G33980.1 | Smg-4/UPF3 family protein | UPF3 | 1day perior to sampling | 0.22 |
| AT5G51195.1 | NULL | - | 1day perior to sampling | 0.42 |
| AT5G45540.1 | Protein of unknown function (DUF594) | - | 1day perior to sampling | 0.22 |
| AT3G61750.2 | Cytochrome b561/ferric reductase transmembrane with DOMON related domain | - | 1day perior to sampling | 0.22 |
| AT2G09910.1 | transposable element gene | - | 1day perior to sampling | 0.07 |
| AT2G20770.1 | GCR2-like 2 | GCL2 | 1day perior to sampling | 0.26 |
| AT5G08645.1 | | - | 1day perior to sampling | 0.27 |
| AT1G66490.1 | F-box and associated interaction domains-containing protein | - | 1day perior to sampling | 0.28 |
| AT2G10760.1 | transposable element gene | - | 1day perior to sampling | 0.18 |
| AT5G08775.1 | | - | 1day perior to sampling | 0.20 |
| AT4G08078.1 | transposable element gene | - | 1day perior to sampling | 0.32 |
| AT5G10100.2 | Haloacid dehalogenase-like hydrolase (HAD) superfamily protein | TPPI | 1day perior to sampling | 0.09 |
| AT5G08805.1 | | - | 1day perior to sampling | 0.10 |
| AT1G56310.1 | Polynucleotidyl transferase, ribonuclease H-like superfamily protein | - | 1day perior to sampling | 0.26 |
| AT5G54110.2 | membrane-associated mannitol-induced | MAMI | 1day perior to sampling | 0.11 |
| AT4G23440.2 | Disease resistance protein (TIR-NBS class) | - | 1day perior to sampling | 0.12 |
| AT4G05215.1 | | - | 1day perior to sampling | 0.17 |
| AT2G37035.1 | NULL | - | 1day perior to sampling | 0.26 |
| AT5G04485.1 | | - | 1day perior to sampling | 0.16 |
| AT2G35090.1 | Protein of unknown function (DUF1640) | - | 1day perior to sampling | 0.15 |
| AT4G03215.1 | | - | 1day perior to sampling | 0.26 |
| AT4G12350.1 | myb domain protein 42 | MYB42 | 1day perior to sampling | 0.09 |
| AT5G04405.1 | | - | 1day perior to sampling | 0.12 |
| AT3G05480.1 | cell cycle checkpoint control protein family | RAD9 | 1day perior to sampling | 0.19 |
| AT5G18440.2 | NULL | NUFIP | 1day perior to sampling | 0.20 |
| AT5G07080.1 | HXXXD-type acyl-transferase family protein | - | 1day perior to sampling | 0.25 |
| AT5G42965.1 | Polynucleotidyl transferase, ribonuclease H-like superfamily protein | - | 1day perior to sampling | 0.15 |
| AT1G66170.1 | RING/FYVE/PHD zinc finger superfamily protein | MMD1 | 1day perior to sampling | 0.07 |
| AT4G21470.1 | riboflavin kinase/FMN hydrolase | FMN/FHY | 1day perior to sampling | 0.27 |
| AT2G03300.1 | Toll-Interleukin-Resistance (TIR) domain family protein | - | 1day perior to sampling | 0.12 |
| AT2G08165.1 | | - | 1day perior to sampling | 0.25 |
| AT4G30740.1 | NULL | - | 1day perior to sampling | 0.30 |
| AT3G16470.3 | Mannose-binding lectin superfamily protein | JR1 | 1day perior to sampling | 0.27 |
| AT5G47590.2 | Heat shock protein HSP20/alpha crystallin family | - | 1day perior to sampling | 0.18 |
| AT4G38440.1 | NULL | IYO | 1day perior to sampling | 0.11 |
| AT2G12770.1 | transposable element gene | - | 1day perior to sampling | 0.26 |
| AT3G07510.1 | NULL | - | 1day perior to sampling | 0.20 |
| AT3G50440.1 | methyl esterase 10 | MES10 | 1day perior to sampling | 0.11 |
| AT2G43510.1 | trypsin inhibitor protein 1 | TI1 | 1day perior to sampling | 0.19 |
| AT5G09475.1 | | - | 1day perior to sampling | 0.13 |
| AT4G04410.1 | transposable element gene | - | 1day perior to sampling | 0.25 |
| AT5G47080.3 | casein kinase II beta chain 1 | CKB1 | 1day perior to sampling | 0.20 |
| AT2G40280.1 | S-adenosyl-L-methionine-dependent methyltransferases superfamily | - | 1day perior to sampling | 0.36 |
| AT2G02640.1 | Cysteine/Histidine-rich C1 domain family protein | - | 1day perior to sampling | 0.28 |
| AT5G22640.3 | MORN (Membrane Occupation and Recognition Nexus) repeat-containing protein | emb1211 | 1day perior to sampling | 0.09 |
| AT2G38170.3 | cation exchanger 1 | CAX1 | 1day perior to sampling | 0.25 |
| AT3G44730.2 | kinesin-like protein 1 | KP1 | 1day perior to sampling | 0.17 |
| AT5G67250.1 | SKP1/ASK1-interacting protein 2 | SKIP2 | 1day perior to sampling | 0.19 |
| AT1G22400.1 | UDP-Glycosyltransferase superfamily protein | UGT85A1 | 1day perior to sampling | 0.26 |
| AT2G42560.1 | late embryogenesis abundant domain-containing protein / LEA domain-containing protein | - | 1day perior to sampling | 0.11 |
| AT3G42760.1 | transposable element gene | - | 1day perior to sampling | 0.20 |
| AT1G05297.1 | | - | 1day perior to sampling | 0.30 |
| AT3G46640.2 | Homeodomain-like superfamily protein | PCL1 | 1day perior to sampling | 0.14 |
| AT4G29090.1 | Ribonuclease H-like superfamily protein | - | 1day perior to sampling | 0.09 |
| AT3G16770.1 | ethylene-responsive element binding protein | EBP | 1day perior to sampling | 0.41 |
| AT5G58380.1 | SOS3-interacting protein 1 | SIP1 | 1day perior to sampling | 0.20 |
| AT5G22560.1 | Plant protein of unknown function (DUF247) | - | 1day perior to sampling | 0.13 |
| AT5G39785.3 | Protein of unknown function (DUF1666) | - | 1day perior to sampling | 0.10 |
| AT2G12940.1 | Basic-leucine zipper (bZIP) transcription factor family protein | UNE4 | 1day perior to sampling | 0.23 |
| AT3G00820.1 | | - | 1day perior to sampling | 0.14 |
| AT3G31358.1 | transposable element gene | - | 1day perior to sampling | 0.05 |
| AT4G27530.1 | NULL | - | 1day perior to sampling | 0.09 |
| AT1G32050.1 | SCAMP family protein | SCAMP5 | 1day perior to sampling | 0.08 |
| AT2G11370.1 | transposable element gene | - | 1day perior to sampling | 0.05 |
| AT1G55460.1 | DNA/RNA-binding protein Kin17, conserved region | - | 1day perior to sampling | 0.15 |
| AT1G26800.1 | RING/U-box superfamily protein | - | 1day perior to sampling | 0.09 |

| | | | | |
|---|---|---|---|---|
| AT3G42471.1 | transposable element gene | - | 1day perior to sampling | 0.11 |
| AT1G64130.1 | Polyketide cyclase/dehydrase and lipid transport superfamily protein | - | 1day perior to sampling | 0.13 |
| AT5G19740.1 | Peptidase M28 family protein | - | 1day perior to sampling | 0.15 |
| AT2G34925.1 | CLAVATA3/ESR-RELATED 42 | CLE42 | 1day perior to sampling | 0.10 |
| AT3G49200.1 | O-acyltransferase (WSD1-like) family protein | - | 1day perior to sampling | 0.30 |
| AT5G56490.1 | D-arabinono-1,4-lactone oxidase family protein | GulLO4 | 1day perior to sampling | 0.10 |
| AT5G56670.1 | Ribosomal protein S30 family protein | - | 1day perior to sampling | 0.09 |
| AT5G00675.1 | | - | 1day perior to sampling | 0.25 |
| AT1G67000.1 | Protein kinase superfamily protein | - | 1day perior to sampling | 0.28 |
| AT5G02495.1 | | - | 1day perior to sampling | 0.47 |
| AT1G21550.1 | Calcium-binding EF-hand family protein | - | 1day perior to sampling | 0.33 |
| AT3G01655.1 | | - | 1day perior to sampling | 0.30 |
| AT4G15215.3 | pleiotropic drug resistance 13 | ABCG41 | 1day perior to sampling | 0.05 |
| AT4G19550.2 | zinc ion binding;transcription regulators | - | 1day perior to sampling | 0.28 |
| AT5G62900.1 | NULL | - | 1day perior to sampling | 0.42 |
| AT1G05660.1 | Pectin lyase-like superfamily protein | - | 1day perior to sampling | 0.13 |
| AT3G61690.2 | nucleotidyltransferases | - | 1day perior to sampling | 0.23 |
| AT1G52010.1 | transposable element gene | - | 1day perior to sampling | 0.34 |
| AT2G01460.3 | P-loop containing nucleoside triphosphate hydrolases superfamily | - | 1day perior to sampling | 0.31 |
| AT2G47100.1 | pre-tRNA | - | 1day perior to sampling | 0.12 |
| AT3G10870.1 | methyl esterase 17 | MES17 | 1day perior to sampling | 0.16 |
| AT1G04803.1 | | - | 1day perior to sampling | 0.08 |
| AT1G08953.1 | | - | 1day perior to sampling | 0.12 |
| AT4G37570.1 | transposable element gene | - | 1day perior to sampling | 0.10 |
| AT1G73885.1 | NULL | - | 1day perior to sampling | 0.42 |
| AT1G51030.1 | NULL | - | 1day perior to sampling | 0.32 |
| AT5G05740.2 | ethylene-dependent gravitropism-deficient and yellow-green-like 2 | EGY2 | 1day perior to sampling | 0.07 |
| AT5G50320.1 | radical SAM domain-containing protein / GCN5-related N-acetyltransferase (GNAT) family protein | ELO3 | 1day perior to sampling | 0.23 |
| AT4G10010.2 | Protein kinase superfamily protein | - | 1day perior to sampling | 0.16 |
| AT3G11440.2 | myb domain protein 65 | MYB65 | 1day perior to sampling | 0.12 |
| AT5G67580.1 | Homeodomain-like/winged-helix DNA-binding family protein | TRB2 | 1day perior to sampling | 0.25 |
| AT1G29410.2 | phosphoribosylanthranilate isomerase 3 | PAI3 | 1day perior to sampling | 0.07 |
| AT3G58800.1 | NULL | - | 1day perior to sampling | 0.24 |
| AT3G03995.1 | | - | 1day perior to sampling | 0.14 |
| AT1G03290.2 | NULL | - | 1day perior to sampling | 0.19 |
| AT4G12230.1 | alpha/beta-Hydrolases superfamily protein | - | 1day perior to sampling | 0.23 |
| AT2G09160.1 | | - | 1day perior to sampling | 0.08 |
| AT5G20430.1 | Mob1/phocein family protein | - | 1day perior to sampling | 0.13 |
| AT5G17140.1 | Cysteine proteinases superfamily protein | - | 1day perior to sampling | 0.08 |
| AT5G43930.3 | Transducin family protein / WD-40 repeat family protein | - | 1day perior to sampling | 0.06 |
| AT5G24000.1 | Protein of unknown function (DUF819) | - | 1day perior to sampling | 0.11 |
| AT2G02200.1 | transposable element gene | - | 1day perior to sampling | 0.38 |
| AT4G06618.1 | transposable element gene | - | 1day perior to sampling | 0.20 |
| AT1G10310.1 | NAD(P)-binding Rossmann-fold superfamily protein | - | 1day perior to sampling | 0.10 |
| AT1G63450.1 | root hair specific 8 | RHS8 | 1day perior to sampling | 0.07 |
| AT5G02650.1 | NULL | - | 1day perior to sampling | 0.10 |
| AT5G50290.1 | NULL | - | 1day perior to sampling | 0.12 |
| AT5G62260.2 | AT hook motif DNA-binding family protein | - | 1day perior to sampling | -0.05 |
| AT3G33555.1 | transposable element gene | - | 1day perior to sampling | -0.23 |
| AT3G07115.1 | pre-tRNA | - | 1day perior to sampling | -0.06 |
| AT5G20300.4 | Avirulence induced gene (AIG1) family protein | Toc90 | 1day perior to sampling | -0.10 |
| AT5G34839.1 | transposable element gene | - | 1day perior to sampling | -0.06 |
| AT1G60787.2 | | - | 1day perior to sampling | -0.07 |
| AT3G19830.1 | Calcium-dependent lipid-binding (CaLB domain) family protein | NTMC2T5.2 | 1day perior to sampling | -0.09 |
| AT3G42556.1 | transposable element gene | - | 1day perior to sampling | -0.05 |
| AT4G22400.1 | NULL | - | 1day perior to sampling | -0.06 |
| AT1G02570.1 | NULL | - | 1day perior to sampling | -0.06 |
| AT1G26790.1 | Dof-type zinc finger DNA-binding family protein | - | 1day perior to sampling | -0.05 |
| AT1G01350.1 | Zinc finger (CCCH-type/C3HC4-type RING finger) family protein | - | 1day perior to sampling | -0.06 |
| AT2G30290.1 | VACUOLAR SORTING RECEPTOR 2 | VSR2 | 1day perior to sampling | -0.07 |
| AT1G06570.2 | phytoene desaturation 1 | PDS1 | 1day perior to sampling | -0.14 |
| AT3G06240.1 | F-box family protein | - | 1day perior to sampling | -0.15 |
| AT2G12083.1 | transposable element gene | - | 1day perior to sampling | -0.06 |
| AT2G05195.1 | | - | 1day perior to sampling | -0.13 |
| AT1G16080.1 | NULL | - | 1day perior to sampling | -0.08 |
| AT2G08205.1 | | - | 1day perior to sampling | -0.06 |
| AT1G47340.1 | F-box and associated interaction domains-containing protein | - | 1day perior to sampling | -0.11 |
| AT5G35140.1 | transposable element gene | - | 1day perior to sampling | -0.07 |
| AT2G04920.1 | F-box and associated interaction domains-containing protein | - | 1day perior to sampling | -0.07 |
| AT1G08717.1 | | - | 1day perior to sampling | -0.23 |
| AT3G07110.1 | Ribosomal protein L13 family protein | - | 1day perior to sampling | -0.14 |
| AT2G39420.1 | alpha/beta-Hydrolases superfamily protein | - | 1day perior to sampling | -0.05 |
| AT1G07900.1 | LOB domain-containing protein 1 | LBD1 | 1day perior to sampling | -0.14 |
| AT3G21160.2 | alpha-mannosidase 2 | MNS2 | 1day perior to sampling | -0.08 |
| AT1G04993.1 | | - | 1day perior to sampling | -0.13 |

| | | | | |
|---|---|---|---|---|
| AT3G54800.1 | Pleckstrin homology (PH) and lipid-binding START domains- | - | 1day perior to sampling | -0.08 |
| AT5G04190.1 | phytochrome kinase substrate 4 | PKS4 | 1day perior to sampling | -0.23 |
| AT3G29460.1 | transposable element gene | - | 1day perior to sampling | -0.36 |
| AT5G26910.4 | NULL | TRM8 | 1day perior to sampling | -0.27 |
| AT1G13730.1 | Nuclear transport factor 2 (NTF2) family protein with RNA binding (RRM-RBD-RNP motifs) domain | - | 1day perior to sampling | -0.22 |
| ATMG00340.1 | NULL | TRNY.1 | 1day perior to sampling | -0.05 |
| AT3G03435.1 | | - | 1day perior to sampling | -0.09 |
| ATCG00200.1 | NULL | TRNC | 1day perior to sampling | -0.06 |
| AT2G25010.1 | Aminotransferase-like, plant mobile domain family protein | - | 1day perior to sampling | -0.29 |
| AT2G20362.1 | NULL | - | 1day perior to sampling | -0.09 |
| AT1G21870.1 | golgi nucleotide sugar transporter 5 | GONST5 | 1day perior to sampling | -0.26 |
| AT4G22415.1 | transposable element gene | - | 1day perior to sampling | -0.31 |
| AT1G68765.1 | Putative membrane lipoprotein | IDA | 1day perior to sampling | -0.07 |
| AT3G31460.1 | transposable element gene | - | 1day perior to sampling | -0.12 |
| AT5G41491.1 | NULL | - | 1day perior to sampling | -0.05 |
| AT1G22885.1 | NULL | - | 1day perior to sampling | -0.10 |
| AT1G13755.1 | Defensin-like (DEFL) family protein | - | 1day perior to sampling | -0.25 |
| AT4G06722.1 | transposable element gene | - | 1day perior to sampling | -0.12 |
| AT5G07240.1 | IQ-domain 24 | IQD24 | 1day perior to sampling | -0.08 |
| AT2G17620.1 | Cyclin B2;1 | CYCB2%3B1 | 1day perior to sampling | -0.10 |
| AT2G36580.1 | Pyruvate kinase family protein | - | 1day perior to sampling | -0.06 |
| AT2G07791.1 | transposable element gene | - | 1day perior to sampling | -0.09 |
| AT1G44040.1 | NULL | - | 1day perior to sampling | -0.09 |
| AT2G05605.1 | | - | 1day perior to sampling | -0.06 |
| AT2G01031.1 | transposable element gene | - | 1day perior to sampling | -0.08 |
| AT3G43950.1 | Protein kinase superfamily protein | - | 1day perior to sampling | -0.25 |
| AT5G23940.1 | HXXXD-type acyl-transferase family protein | PEL3 | 1day perior to sampling | -0.06 |
| AT4G04595.1 | | - | 1day perior to sampling | -0.24 |
| AT1G67630.1 | DNA polymerase alpha 2 | POLA2 | 1day perior to sampling | -0.17 |
| AT5G28770.3 | bZIP transcription factor family protein | BZO2H3 | 1day perior to sampling | -0.08 |
| AT3G22110.1 | 20S proteasome alpha subunit C1 | PAC1 | 1day perior to sampling | -0.05 |
| AT4G30710.4 | Family of unknown function (DUF566) | QWRF8 | 1day perior to sampling | -0.06 |
| AT3G61590.1 | Galactose oxidase/kelch repeat superfamily protein | HWS | 1day perior to sampling | -0.20 |
| AT1G53220.1 | pre-tRNA | - | 1day perior to sampling | -0.06 |
| AT1G48760.2 | delta-adaptin | delta-ADR | 1day perior to sampling | -0.17 |
| AT1G74050.1 | Ribosomal protein L6 family protein | - | 1day perior to sampling | -0.06 |
| AT1G32505.1 | transposable element gene | - | 1day perior to sampling | -0.11 |
| AT1G61850.1 | phospholipases;galactolipases | - | 1day perior to sampling | -0.08 |
| AT4G03113.1 | NULL | - | 1day perior to sampling | -0.21 |
| AT1G09140.3 | SERINE-ARGININE PROTEIN 30 | SR30 | 1day perior to sampling | -0.05 |
| AT5G19290.1 | alpha/beta-Hydrolases superfamily protein | - | 1day perior to sampling | -0.15 |
| AT1G08663.1 | | - | 1day perior to sampling | -0.21 |
| AT4G07565.1 | | - | 1day perior to sampling | -0.12 |
| AT3G55110.1 | ABC-2 type transporter family protein | ABCG18 | 1day perior to sampling | -0.12 |
| AT5G10530.1 | Concanavalin A-like lectin protein kinase family protein | - | 1day perior to sampling | -0.12 |
| AT3G33133.1 | transposable element gene | - | 1day perior to sampling | -0.19 |
| AT4G07655.1 | | - | 1day perior to sampling | -0.06 |
| AT1G56760.1 | pre-tRNA | - | 1day perior to sampling | -0.25 |
| AT4G15150.1 | glycine-rich protein | - | 1day perior to sampling | -0.06 |
| AT2G13940.1 | transposable element gene | - | 1day perior to sampling | -0.34 |
| AT3G01395.1 | | - | 1day perior to sampling | -0.09 |
| AT1G04743.1 | | - | 1day perior to sampling | -0.24 |
| AT1G09863.1 | | - | 1day perior to sampling | -0.22 |
| AT3G07210.1 | NULL | - | 1day perior to sampling | -0.06 |
| AT3G25880.1 | NAD(P)-binding Rossmann-fold superfamily protein | - | 1day perior to sampling | -0.14 |
| AT5G63160.3 | BTB and TAZ domain protein 1 | BT1 | 1day perior to sampling | -0.05 |
| AT3G60340.2 | alpha/beta-Hydrolases superfamily protein | - | 1day perior to sampling | -0.08 |
| AT5G37160.1 | P-loop containing nucleoside triphosphate hydrolases superfamily | - | 1day perior to sampling | -0.06 |
| AT1G19030.1 | transposable element gene | - | 1day perior to sampling | -0.06 |
| AT3G48920.1 | myb domain protein 45 | MYB45 | 1day perior to sampling | -0.18 |
| AT1G05487.1 | | - | 1day perior to sampling | -0.17 |
| AT4G31805.1 | WRKY family transcription factor | - | 1day perior to sampling | -0.06 |
| AT5G08270.1 | NULL | - | 1day perior to sampling | -0.07 |
| AT1G51402.1 | NULL | - | 1day perior to sampling | -0.06 |
| AT4G19680.1 | iron regulated transporter 2 | IRT2 | 1day perior to sampling | -0.18 |
| AT5G04480.1 | UDP-Glycosyltransferase superfamily protein | - | 1day perior to sampling | -0.27 |
| AT5G43770.1 | proline-rich family protein | - | 1day perior to sampling | -0.05 |
| AT5G13890.1 | Family of unknown function (DUF716) | - | 1day perior to sampling | -0.16 |
| AT5G66010.2 | RNA-binding (RRM/RBD/RNP motifs) family protein | - | 1day perior to sampling | -0.08 |
| AT4G24020.1 | NIN like protein 7 | NLP7 | 1day perior to sampling | -0.23 |
| AT5G63490.1 | CBS / octicosapeptide/Phox/Bemp1 (PB1) domains-containing protein | - | 1day perior to sampling | -0.20 |
| AT5G50650.1 | Transducin/WD40 repeat-like superfamily protein | - | 1day perior to sampling | -0.26 |
| AT2G09445.1 | | - | 1day perior to sampling | -0.08 |
| AT2G34000.1 | RING/U-box superfamily protein | - | 1day perior to sampling | -0.25 |
| AT4G16350.1 | calcineurin B-like protein 6 | CBL6 | 1day perior to sampling | -0.07 |

| | | | | |
|---|---|---|---|---|
| AT2G05460.1 | transposable element gene | - | 1day perior to sampling | -0.11 |
| AT1G23930.1 | transposable element gene | - | 1day perior to sampling | -0.22 |
| AT3G04330.1 | Kunitz family trypsin and protease inhibitor protein | - | 1day perior to sampling | -0.15 |
| AT4G20070.1 | allantoate amidohydrolase | AAH | 1day perior to sampling | -0.07 |
| AT1G09075.1 | | - | 1day perior to sampling | -0.07 |
| AT2G38770.1 | P-loop containing nucleoside triphosphate hydrolases superfamily | EMB2765 | 1day perior to sampling | -0.10 |
| AT3G44935.1 | NULL | - | 1day perior to sampling | -0.28 |
| AT2G47570.1 | Ribosomal protein L18e/L15 superfamily protein | - | 1day perior to sampling | -0.23 |
| AT1G76780.2 | HSP20-like chaperones superfamily protein | - | 1day perior to sampling | -0.05 |
| AT2G33470.1 | glycolipid transfer protein 1 | GLTP1 | 1day perior to sampling | -0.27 |
| AT5G17070.1 | NULL | - | 1day perior to sampling | -0.29 |
| AT3G20090.1 | cytochrome P450, family 705, subfamily A, polypeptide 18 | CYP705A18 | 1day perior to sampling | -0.06 |
| AT3G25013.2 | Synaptobrevin family protein | - | 1day perior to sampling | -0.19 |
| AT1G26515.1 | F-box and associated interaction domains-containing protein | - | 1day perior to sampling | -0.13 |
| AT4G27900.1 | CCT motif family protein | - | 1day perior to sampling | -0.08 |
| AT3G05695.1 | | - | 1day perior to sampling | -0.09 |
| AT3G42305.1 | transposable element gene | - | 1day perior to sampling | -0.13 |
| AT3G43684.1 | transposable element gene | - | 1day perior to sampling | -0.09 |
| AT4G29260.1 | HAD superfamily, subfamily IIIB acid phosphatase | - | 1day perior to sampling | -0.22 |
| AT1G74650.2 | myb domain protein 31 | MYB31 | 1day perior to sampling | -0.14 |
| ATMG09450.1 | | - | 1day perior to sampling | -0.09 |
| AT3G14510.1 | Polyprenyl synthetase family protein | - | 1day perior to sampling | -0.28 |
| AT3G04860.1 | Plant protein of unknown function (DUF868) | - | 1day perior to sampling | -0.15 |
| AT1G12480.1 | C4-dicarboxylate transporter/malic acid transport protein | OZS1 | 1day perior to sampling | -0.08 |
| ATCG00590.1 | electron carriers | ORF31 | 1day perior to sampling | -0.06 |
| AT3G11435.1 | MIR172C; miRNA | MIR172C | 1day perior to sampling | -0.06 |
| AT3G15530.1 | S-adenosyl-L-methionine-dependent methyltransferases superfamily | - | 1day perior to sampling | -0.07 |
| AT4G35480.1 | RING-H2 finger A3B | RHA3B | 1day perior to sampling | -0.18 |
| AT5G48860.1 | NULL | - | 1day perior to sampling | -0.29 |
| AT4G08830.1 | transposable element gene | - | 1day perior to sampling | -0.22 |
| ATCG00600.1 | PETG | PETG | 1day perior to sampling | -0.19 |
| AT5G09695.1 | | - | 1day perior to sampling | -0.06 |
| AT5G16870.1 | Peptidyl-tRNA hydrolase II (PTH2) family protein | - | 1day perior to sampling | -0.07 |
| AT3G55370.3 | OBF-binding protein 3 | OBP3 | 1day perior to sampling | -0.13 |
| AT5G00810.1 | | - | 1day perior to sampling | -0.08 |
| AT5G38880.1 | NULL | 5-Aug | 1day perior to sampling | -0.26 |
| AT3G05105.1 | | - | 1day perior to sampling | -0.10 |
| AT2G45180.1 | Bifunctional inhibitor/lipid-transfer protein/seed storage 2S albumin superfamily protein | - | 1day perior to sampling | -0.06 |
| AT5G29624.1 | Cysteine/Histidine-rich C1 domain family protein | - | 1day perior to sampling | -0.07 |
| AT3G44770.1 | Protein of unknown function (DUF626) | - | 1day perior to sampling | -0.16 |
| AT5G09450.1 | Tetratricopeptide repeat (TPR)-like superfamily protein | - | 1day perior to sampling | -0.08 |
| AT3G44267.1 | transposable element gene | - | 1day perior to sampling | -0.05 |
| AT1G59500.1 | Auxin-responsive GH3 family protein | GH3.4 | 1day perior to sampling | -0.15 |
| AT1G74210.1 | PLC-like phosphodiesterases superfamily protein | GDPD5 | 1day perior to sampling | -0.12 |
| AT1G52440.1 | alpha/beta-Hydrolases superfamily protein | - | 1day perior to sampling | -0.06 |
| AT5G65690.4 | phosphoenolpyruvate carboxykinase 2 | PCK2 | 1day perior to sampling | -0.23 |
| AT2G05465.1 | | - | 1day perior to sampling | -0.09 |
| AT4G15955.2 | alpha/beta-Hydrolases superfamily protein | - | 1day perior to sampling | -0.14 |
| AT2G25190.1 | PPPDE putative thiol peptidase family protein | - | 1day perior to sampling | -0.07 |
| AT4G33585.1 | NULL | - | 1day perior to sampling | -0.08 |
| AT1G05980.1 | pre-tRNA | - | 1day perior to sampling | -0.11 |
| AT3G43352.1 | transposable element gene | - | 1day perior to sampling | -0.25 |
| AT5G01945.1 | | - | 1day perior to sampling | -0.06 |
| AT2G09990.1 | Ribosomal protein S5 domain 2-like superfamily protein | - | 1day perior to sampling | -0.15 |
| AT2G14910.2 | NULL | - | 1day perior to sampling | -0.22 |
| AT2G07220.1 | transposable element gene | - | 1day perior to sampling | -0.23 |
| AT3G55840.1 | Hs1pro-1 protein | - | 1day perior to sampling | -0.32 |
| AT5G08540.1 | NULL | - | 1day perior to sampling | -0.26 |
| AT4G07640.1 | transposable element gene | - | 1day perior to sampling | -0.20 |
| AT2G12130.1 | transposable element gene | - | 1day perior to sampling | -0.06 |
| AT3G14540.2 | Terpenoid cyclases/Protein prenyltransferases superfamily protein | - | 1day perior to sampling | -0.06 |
| AT4G28610.1 | phosphate starvation response 1 | PHR1 | 1day perior to sampling | -0.27 |
| AT3G55400.2 | methionyl-tRNA synthetase / methionine--tRNA ligase / MetRS | OVA1 | 1day perior to sampling | -0.05 |
| AT5G17050.1 | UDP-glucosyl transferase 78D2 | UGT78D2 | 1day perior to sampling | -0.29 |
| AT1G53360.2 | F-box associated ubiquitination effector family protein | - | 1day perior to sampling | -0.05 |
| AT5G42232.1 | Defensin-like (DEFL) family protein | - | 1day perior to sampling | -0.06 |
| AT5G08710.2 | Regulator of chromosome condensation (RCC1) family protein | RUG1 | 1day perior to sampling | -0.11 |
| AT3G06650.2 | ATP-citrate lyase B-1 | ACLB-1 | 1day perior to sampling | -0.09 |
| AT4G05300.1 | transposable element gene | - | 1day perior to sampling | -0.32 |
| AT1G44020.1 | Cysteine/Histidine-rich C1 domain family protein | - | 1day perior to sampling | -0.25 |
| AT5G00390.1 | | - | 1day perior to sampling | -0.36 |
| AT2G43680.5 | IQ-domain 14 | IQD14 | 1day perior to sampling | -0.28 |
| AT4G24615.1 | | - | 1day perior to sampling | -0.23 |
| AT2G07689.1 | NADH-Ubiquinone/plastoquinone (complex I) protein | - | 1day perior to sampling | -0.06 |
| AT5G60335.1 | Thioesterase superfamily protein | - | 1day perior to sampling | -0.08 |

| | | | | |
|---|---|---|---|---|
| AT1G14770.2 | RING/FYVE/PHD zinc finger superfamily protein | - | 1day perior to sampling | -0.07 |
| AT4G20090.1 | Pentatricopeptide repeat (PPR) superfamily protein | EMB1025 | 1day perior to sampling | -0.07 |
| AT3G49370.1 | Calcium-dependent protein kinase (CDPK) family protein | - | 1day perior to sampling | -0.25 |
| AT2G34860.1 | DnaJ/Hsp40 cysteine-rich domain superfamily protein | EDA3 | 1day perior to sampling | -0.07 |
| AT4G23960.2 | F-box family protein | - | 1day perior to sampling | -0.21 |
| AT3G52075.1 | | - | 1day perior to sampling | -0.07 |
| AT1G60980.1 | gibberellin 20-oxidase 4 | GA20OX4 | 1day perior to sampling | -0.07 |
| AT1G64990.2 | GPCR-type G protein 1 | GTG1 | 1day perior to sampling | -0.17 |
| AT1G51745.1 | Tudor/PWWP/MBT superfamily protein | - | 1day perior to sampling | -0.07 |
| AT3G18200.2 | nodulin MtN21 /EamA-like transporter family protein | UMAMIT4 | 1day perior to sampling | -0.06 |
| AT5G16640.1 | Pentatricopeptide repeat (PPR) superfamily protein | - | 1day perior to sampling | -0.30 |
| AT3G33025.1 | NULL | - | 1day perior to sampling | -0.19 |
| AT3G18670.2 | Ankyrin repeat family protein | - | 1day perior to sampling | -0.07 |
| AT5G19020.1 | mitochondrial editing factor 18 | MEF18 | 1day perior to sampling | -0.06 |
| AT5G28515.1 | transposable element gene | - | 1day perior to sampling | -0.30 |
| AT2G39610.1 | pre-tRNA | - | 1day perior to sampling | -0.25 |
| AT2G42920.1 | Pentatricopeptide repeat (PPR-like) superfamily protein | - | 1day perior to sampling | -0.08 |
| AT1G03310.2 | debranching enzyme 1 | DBE1 | 1day perior to sampling | -0.24 |
| AT4G20200.1 | Terpenoid cyclases/Protein prenyltransferases superfamily protein | - | 1day perior to sampling | -0.06 |
| AT1G70250.1 | receptor serine/threonine kinase, putative | - | 1day perior to sampling | -0.05 |
| AT3G24620.1 | RHO guanyl-nucleotide exchange factor 8 | ROPGEF8 | 1day perior to sampling | -0.22 |
| AT3G43682.1 | NULL | - | 1day perior to sampling | -0.28 |
| AT1G52980.1 | GTP-binding family protein | AtNug2 | 1day perior to sampling | -0.22 |
| AT2G02023.1 | NULL | - | 1day perior to sampling | -0.28 |
| AT4G17070.1 | peptidyl-prolyl cis-trans isomerases | - | 1day perior to sampling | -0.08 |
| AT1G14640.1 | SWAP (Suppressor-of-White-APricot)/surp domain-containing protein | - | 1day perior to sampling | -0.22 |
| AT4G21120.2 | amino acid transporter 1 | AAT1 | 1day perior to sampling | -0.05 |
| AT5G54320.1 | Protein of unknown function (DUF295) | - | 1day perior to sampling | -0.07 |
| AT3G28685.1 | pre-tRNA | - | 1day perior to sampling | -0.32 |
| AT5G61580.1 | phosphofructokinase 4 | PFK4 | 1day perior to sampling | -0.20 |
| AT3G03920.1 | H/ACA ribonucleoprotein complex, subunit Gar1/Naf1 protein | - | 1day perior to sampling | -0.19 |
| AT1G52150.3 | Homeobox-leucine zipper family protein / lipid-binding START domain-containing protein | ATHB-15 | 1day perior to sampling | -0.18 |
| AT1G26340.1 | cytochrome B5 isoform A | CB5-A | 1day perior to sampling | -0.06 |
| AT4G04050.1 | transposable element gene | - | 1day perior to sampling | -0.06 |
| AT1G09835.1 | | - | 1day perior to sampling | -0.15 |
| AT3G24550.1 | proline extensin-like receptor kinase 1 | PERK1 | 1day perior to sampling | -0.16 |
| AT5G19890.1 | Peroxidase superfamily protein | - | 1day perior to sampling | -0.06 |
| AT5G20830.1 | sucrose synthase 1 | SUS1 | 1day perior to sampling | -0.18 |
| AT1G77690.1 | like AUX1 3 | LAX3 | 1day perior to sampling | -0.11 |
| AT3G16540.1 | DegP protease 11 | DEG11 | 1day perior to sampling | -0.10 |
| AT3G18535.2 | tubulin-tyrosine ligases | - | 1day perior to sampling | -0.08 |
| AT1G61680.2 | terpene synthase 14 | TPS14 | 1day perior to sampling | -0.07 |
| AT2G17950.1 | Homeodomain-like superfamily protein | WUS | 1day perior to sampling | -0.21 |
| AT3G42850.2 | Mevalonate/galactokinase family protein | - | 1day perior to sampling | -0.10 |
| AT2G29735.1 | | - | 1day perior to sampling | -0.13 |
| AT4G11240.1 | Calcineurin-like metallo-phosphoesterase superfamily protein | TOPP7 | 1day perior to sampling | -0.21 |
| AT3G47150.1 | F-box and associated interaction domains-containing protein | - | 1day perior to sampling | -0.06 |
| AT1G72440.1 | CCAAT-binding factor | EDA25 | 1day perior to sampling | -0.08 |
| AT4G38510.3 | ATPase, V1 complex, subunit B protein | VAB2 | 1day perior to sampling | -0.18 |
| AT5G50470.1 | nuclear factor Y, subunit C7 | NF-YC7 | 1day perior to sampling | -0.05 |
| AT1G11610.2 | cytochrome P450, family 71, subfamily A, polypeptide 18 | CYP71A18 | 1day perior to sampling | -0.09 |
| AT4G04530.1 | transposable element gene | - | 1day perior to sampling | -0.09 |
| AT5G65290.2 | LMBR1-like membrane protein | - | 1day perior to sampling | -0.09 |
| AT5G08605.1 | | - | 1day perior to sampling | -0.27 |
| AT1G37012.1 | transposable element gene | - | 1day perior to sampling | -0.18 |
| AT4G06630.1 | transposable element gene | - | 1day perior to sampling | -0.21 |
| AT1G67410.2 | Exostosin family protein | - | 1day perior to sampling | -0.17 |
| AT3G12110.1 | actin-11 | ACT11 | 1day perior to sampling | -0.09 |
| AT4G24400.1 | CBL-interacting protein kinase 8 | CIPK8 | 1day perior to sampling | -0.10 |
| AT4G35680.1 | Arabidopsis protein of unknown function (DUF241) | - | 1day perior to sampling | -0.07 |
| AT4G31354.1 | NULL | - | 1day perior to sampling | -0.08 |
| AT3G09505.1 | pre-tRNA | - | 2day perior to sampling | 0.34 |
| AT1G09467.1 | | - | 2day perior to sampling | 0.24 |
| AT5G48070.1 | xyloglucan endotransglucosylase/hydrolase 20 | XTH20 | 2day perior to sampling | 0.24 |
| AT3G46650.1 | UDP-Glycosyltransferase superfamily protein | - | 2day perior to sampling | 0.25 |
| AT5G48760.1 | Ribosomal protein L13 family protein | - | 2day perior to sampling | 0.39 |
| AT1G71830.1 | somatic embryogenesis receptor-like kinase 1 | SERK1 | 2day perior to sampling | 0.05 |
| AT1G11220.2 | Protein of unknown function (DUF761) | - | 2day perior to sampling | 0.08 |
| AT3G07685.1 | | - | 2day perior to sampling | 0.22 |
| AT1G28050.1 | B-box type zinc finger protein with CCT domain | BBX13 | 2day perior to sampling | 0.11 |
| AT3G09100.2 | mRNA capping enzyme family protein | - | 2day perior to sampling | 0.13 |
| AT2G23445.1 | | - | 2day perior to sampling | 0.19 |
| AT5G16990.1 | Zinc-binding dehydrogenase family protein | - | 2day perior to sampling | 0.18 |
| AT1G75394.1 | NULL | - | 2day perior to sampling | 0.18 |
| AT2G35520.1 | Defender against death (DAD family) protein | DAD2 | 2day perior to sampling | 0.06 |

| | | | | |
|---|---|---|---|---|
| AT3G08600.1 | Protein of unknown function (DUF1191) | - | 2day perior to sampling | 0.16 |
| AT3G45500.1 | NULL | - | 2day perior to sampling | 0.28 |
| AT1G75120.1 | Nucleotide-diphospho-sugar transferase family protein | RRA1 | 2day perior to sampling | 0.26 |
| AT5G56280.1 | COP9 signalosome subunit 6A | CSN6A | 2day perior to sampling | 0.27 |
| AT2G04290.1 | transposable element gene | - | 2day perior to sampling | 0.28 |
| AT5G03970.1 | F-box associated ubiquitination effector family protein | - | 2day perior to sampling | 0.19 |
| AT1G52320.5 | NULL | - | 2day perior to sampling | 0.20 |
| AT5G61605.1 | NULL | - | 2day perior to sampling | 0.11 |
| AT3G08105.1 | | - | 2day perior to sampling | 0.30 |
| AT4G28050.1 | tetraspanin7 | TET7 | 2day perior to sampling | 0.33 |
| AT4G17914.1 | NULL | - | 2day perior to sampling | 0.12 |
| AT4G09146.1 | transposable element gene | - | 2day perior to sampling | 0.16 |
| AT4G11213.1 | pre-tRNA | - | 2day perior to sampling | 0.09 |
| AT1G24180.1 | Thiamin diphosphate-binding fold (THDP-binding) superfamily protein | IAR4 | 2day perior to sampling | 0.23 |
| AT1G58300.1 | heme oxygenase 4 | HO4 | 2day perior to sampling | 0.26 |
| AT5G56452.1 | FBD-like domain family protein | - | 2day perior to sampling | 0.08 |
| AT2G40500.1 | Protein kinase superfamily protein | - | 2day perior to sampling | 0.10 |
| AT5G50690.2 | hydroxysteroid dehydrogenase 7 | HSD7 | 2day perior to sampling | 0.29 |
| AT5G18870.1 | Inosine-uridine preferring nucleoside hydrolase family protein | NSH5 | 2day perior to sampling | 0.12 |
| AT2G39170.1 | NULL | - | 2day perior to sampling | 0.23 |
| AT3G29792.1 | transposable element gene | - | 2day perior to sampling | 0.09 |
| AT1G09233.1 | | - | 2day perior to sampling | 0.31 |
| AT5G02502.1 | Oligosaccaryltransferase | - | 2day perior to sampling | 0.07 |
| AT4G25900.1 | Galactose mutarotase-like superfamily protein | - | 2day perior to sampling | 0.23 |
| AT1G10320.2 | Zinc finger C-x8-C-x5-C-x3-H type family protein | - | 2day perior to sampling | 0.25 |
| AT4G35700.1 | zinc finger (C2H2 type) family protein | DAZ3 | 2day perior to sampling | 0.23 |
| AT1G06483.1 | | - | 2day perior to sampling | 0.07 |
| AT5G58595.1 | snoRNA | - | 2day perior to sampling | 0.23 |
| AT3G60610.1 | NULL | - | 2day perior to sampling | 0.17 |
| AT3G43090.1 | transposable element gene | - | 2day perior to sampling | 0.33 |
| AT5G38410.1 | Ribulose bisphosphate carboxylase (small chain) family protein | RBCS3B | 2day perior to sampling | 0.19 |
| AT4G07516.1 | transposable element gene | - | 2day perior to sampling | 0.10 |
| AT3G43251.1 | NULL | - | 2day perior to sampling | 0.27 |
| AT5G37730.1 | NULL | - | 2day perior to sampling | 0.08 |
| AT5G13870.2 | xyloglucan endotransglucosylase/hydrolase 5 | XTH5 | 2day perior to sampling | 0.21 |
| AT2G32800.1 | protein kinase family protein | AP4.3A | 2day perior to sampling | 0.21 |
| AT3G56650.1 | Mog1/PsbP/DUF1795-like photosystem II reaction center PsbP family | PPD6 | 2day perior to sampling | 0.06 |
| AT3G20540.1 | polymerase gamma 1 | POLGAMMA1 | 2day perior to sampling | 0.06 |
| AT5G48480.1 | Lactoylglutathione lyase / glyoxalase I family protein | - | 2day perior to sampling | 0.13 |
| AT5G18300.1 | NAC domain containing protein 88 | NAC088 | 2day perior to sampling | 0.23 |
| AT3G07755.1 | | - | 2day perior to sampling | 0.26 |
| AT2G05710.1 | aconitase 3 | ACO3 | 2day perior to sampling | 0.09 |
| AT3G22142.1 | Bifunctional inhibitor/lipid-transfer protein/seed storage 2S albumin superfamily protein | - | 2day perior to sampling | 0.08 |
| AT4G06698.1 | transposable element gene | - | 2day perior to sampling | 0.14 |
| AT5G50580.2 | SUMO-activating enzyme 1B | SAE1B | 2day perior to sampling | 0.24 |
| AT2G04785.1 | | - | 2day perior to sampling | 0.06 |
| AT1G35537.1 | Defensin-like (DEFL) family protein | - | 2day perior to sampling | 0.18 |
| AT2G19590.1 | ACC oxidase 1 | ACO1 | 2day perior to sampling | 0.18 |
| AT4G23380.1 | Protein of Unknown Function (DUF239) | - | 2day perior to sampling | 0.12 |
| AT3G61010.1 | Ferritin/ribonucleotide reductase-like family protein | - | 2day perior to sampling | 0.16 |
| AT1G03610.2 | Protein of unknown function (DUF789) | - | 2day perior to sampling | 0.25 |
| AT4G04400.1 | transposable element gene | - | 2day perior to sampling | 0.06 |
| AT4G08078.1 | transposable element gene | - | 2day perior to sampling | 0.23 |
| AT4G23260.2 | cysteine-rich RLK (RECEPTOR-like protein kinase) 18 | CRK18 | 2day perior to sampling | 0.13 |
| AT2G17180.1 | C2H2-like zinc finger protein | DAZ1 | 2day perior to sampling | 0.16 |
| AT5G24120.3 | sigma factor E | SIGE | 2day perior to sampling | 0.21 |
| AT4G26420.1 | S-adenosyl-L-methionine-dependent methyltransferases superfamily | GAMT1 | 2day perior to sampling | 0.26 |
| AT3G15200.1 | Tetratricopeptide repeat (TPR)-like superfamily protein | - | 2day perior to sampling | 0.30 |
| AT1G35590.1 | transposable element gene | - | 2day perior to sampling | 0.24 |
| AT2G02870.1 | Galactose oxidase/kelch repeat superfamily protein | - | 2day perior to sampling | 0.19 |
| AT2G28720.1 | Histone superfamily protein | - | 2day perior to sampling | 0.34 |
| AT5G46330.1 | Leucine-rich receptor-like protein kinase family protein | FLS2 | 2day perior to sampling | 0.31 |
| AT3G46460.2 | ubiquitin-conjugating enzyme 13 | UBC13 | 2day perior to sampling | 0.07 |
| AT5G04485.1 | | - | 2day perior to sampling | 0.16 |
| AT5G24370.1 | Plant invertase/pectin methylesterase inhibitor superfamily protein | - | 2day perior to sampling | 0.13 |
| AT3G04115.1 | | - | 2day perior to sampling | 0.32 |
| AT1G15320.1 | NULL | - | 2day perior to sampling | 0.24 |
| AT5G60820.1 | RING/U-box superfamily protein | - | 2day perior to sampling | 0.11 |
| AT2G08960.1 | | - | 2day perior to sampling | 0.14 |
| AT1G38194.1 | transposable element gene | - | 2day perior to sampling | 0.33 |
| AT3G28680.1 | Serine carboxypeptidase S28 family protein | - | 2day perior to sampling | 0.09 |
| AT4G15910.1 | drought-induced 21 | DI21 | 2day perior to sampling | 0.23 |
| AT5G11870.1 | Alkaline phytoceramidase (aPHC) | - | 2day perior to sampling | 0.16 |
| AT5G24490.1 | 30S ribosomal protein, putative | - | 2day perior to sampling | 0.23 |
| AT5G09490.1 | Ribosomal protein S19 family protein | - | 2day perior to sampling | 0.21 |

| | | | | |
|---|---|---|---|---|
| AT4G14170.1 | Pentatricopeptide repeat (PPR) superfamily protein | - | 2day perior to sampling | 0.06 |
| AT4G21210.1 | PPDK regulatory protein | RP1 | 2day perior to sampling | 0.22 |
| AT5G07110.1 | prenylated RAB acceptor 1.B6 | PRA1.B6 | 2day perior to sampling | 0.08 |
| AT5G19210.2 | P-loop containing nucleoside triphosphate hydrolases superfamily | - | 2day perior to sampling | 0.22 |
| AT5G64395.1 | NULL | - | 2day perior to sampling | 0.10 |
| AT3G48320.1 | cytochrome P450, family 71, subfamily A, polypeptide 21 | CYP71A21 | 2day perior to sampling | 0.16 |
| AT3G32050.1 | NULL | - | 2day perior to sampling | 0.07 |
| AT3G59530.3 | Calcium-dependent phosphotriesterase superfamily protein | LAP3 | 2day perior to sampling | 0.09 |
| AT3G52730.2 | ubiquinol-cytochrome C reductase UQCRX/QCR9-like family protein | - | 2day perior to sampling | 0.05 |
| AT2G07535.1 | | - | 2day perior to sampling | 0.06 |
| AT4G09784.1 | NULL | - | 2day perior to sampling | 0.06 |
| AT4G26080.1 | Protein phosphatase 2C family protein | ABI1 | 2day perior to sampling | 0.19 |
| AT5G47175.1 | low-molecular-weight cysteine-rich 3 | LCR3 | 2day perior to sampling | 0.11 |
| AT3G01985.1 | | - | 2day perior to sampling | 0.06 |
| AT1G15260.1 | NULL | - | 2day perior to sampling | 0.06 |
| AT1G64253.1 | | - | 2day perior to sampling | 0.07 |
| AT3G31360.1 | transposable element gene | - | 2day perior to sampling | 0.06 |
| AT5G51780.2 | basic helix-loop-helix (bHLH) DNA-binding superfamily protein | - | 2day perior to sampling | 0.05 |
| AT2G05950.1 | transposable element gene | - | 2day perior to sampling | 0.05 |
| AT1G23250.2 | Caleosin-related family protein | - | 2day perior to sampling | 0.19 |
| AT3G54910.2 | RNI-like superfamily protein | - | 2day perior to sampling | 0.18 |
| AT3G19270.2 | cytochrome P450, family 707, subfamily A, polypeptide 4 | CYP707A4 | 2day perior to sampling | 0.06 |
| AT1G11655.1 | NULL | - | 2day perior to sampling | 0.16 |
| AT3G20395.1 | RING/U-box superfamily protein | - | 2day perior to sampling | 0.26 |
| AT2G33070.3 | nitrile specifier protein 2 | NSP2 | 2day perior to sampling | 0.10 |
| AT1G56340.1 | calreticulin 1a | CRT1a | 2day perior to sampling | 0.28 |
| AT2G02640.1 | Cysteine/Histidine-rich C1 domain family protein | - | 2day perior to sampling | 0.24 |
| AT5G42232.1 | Defensin-like (DEFL) family protein | - | 2day perior to sampling | 0.05 |
| AT3G50130.1 | Plant protein of unknown function (DUF247) | - | 2day perior to sampling | 0.29 |
| AT2G27290.1 | Protein of unknown function (DUF1279) | - | 2day perior to sampling | 0.20 |
| AT4G01990.1 | Tetratricopeptide repeat (TPR)-like superfamily protein | - | 2day perior to sampling | 0.06 |
| AT3G42760.1 | transposable element gene | - | 2day perior to sampling | 0.19 |
| AT1G22250.1 | NULL | - | 2day perior to sampling | 0.12 |
| AT4G14342.2 | Splicing factor 3B subunit 5/RDS3 complex subunit 10 | - | 2day perior to sampling | 0.25 |
| AT5G54300.1 | Protein of unknown function (DUF761) | - | 2day perior to sampling | 0.25 |
| AT3G43920.3 | dicer-like 3 | DCL3 | 2day perior to sampling | 0.22 |
| AT3G07125.1 | | - | 2day perior to sampling | 0.30 |
| AT3G16090.1 | RING/U-box superfamily protein | Hrd1A | 2day perior to sampling | 0.09 |
| AT2G01260.3 | Protein of unknown function (DUF789) | - | 2day perior to sampling | 0.23 |
| AT4G00520.4 | Acyl-CoA thioesterase family protein | - | 2day perior to sampling | 0.10 |
| AT5G02990.1 | Galactose oxidase/kelch repeat superfamily protein | - | 2day perior to sampling | 0.25 |
| AT4G30840.1 | Transducin/WD40 repeat-like superfamily protein | - | 2day perior to sampling | 0.21 |
| AT5G49570.1 | peptide-N-glycanase 1 | PNG1 | 2day perior to sampling | 0.13 |
| AT4G07738.1 | transposable element gene | - | 2day perior to sampling | 0.35 |
| AT2G36325.1 | GDSL-like Lipase/Acylhydrolase superfamily protein | - | 2day perior to sampling | 0.08 |
| AT3G02242.1 | NULL | - | 2day perior to sampling | 0.06 |
| AT1G54920.4 | NULL | - | 2day perior to sampling | 0.25 |
| AT3G13870.1 | Root hair defective 3 GTP-binding protein (RHD3) | RHD3 | 2day perior to sampling | 0.16 |
| AT4G07090.1 | | - | 2day perior to sampling | 0.09 |
| AT3G63088.1 | ROTUNDIFOLIA like 14 | RTFL14 | 2day perior to sampling | 0.22 |
| AT4G00885.1 | MIR165/MIR165B; miRNA | MIR165B | 2day perior to sampling | 0.07 |
| AT2G09370.1 | | - | 2day perior to sampling | 0.33 |
| AT3G18215.2 | Protein of unknown function, DUF599 | - | 2day perior to sampling | 0.30 |
| AT4G38775.1 | | - | 2day perior to sampling | 0.23 |
| AT3G06450.1 | HCO3- transporter family | - | 2day perior to sampling | 0.13 |
| AT5G18840.1 | Major facilitator superfamily protein | - | 2day perior to sampling | 0.09 |
| AT4G25300.2 | 2-oxoglutarate (2OG) and Fe(II)-dependent oxygenase superfamily | - | 2day perior to sampling | 0.11 |
| AT3G31908.2 | NULL | - | 2day perior to sampling | 0.28 |
| AT2G15325.1 | Bifunctional inhibitor/lipid-transfer protein/seed storage 2S albumin superfamily protein | - | 2day perior to sampling | 0.12 |
| AT3G20680.1 | Domain of unknown function (DUF1995) | - | 2day perior to sampling | 0.08 |
| AT5G62180.1 | carboxyesterase 20 | CXE20 | 2day perior to sampling | 0.06 |
| AT1G23200.1 | Plant invertase/pectin methylesterase inhibitor superfamily | - | 2day perior to sampling | 0.10 |
| AT3G14670.3 | NULL | - | 2day perior to sampling | 0.07 |
| AT2G25490.1 | EIN3-binding F box protein 1 | EBF1 | 2day perior to sampling | 0.12 |
| AT3G02555.2 | NULL | - | 2day perior to sampling | 0.18 |
| AT5G09575.1 | | - | 2day perior to sampling | 0.07 |
| AT3G11440.2 | myb domain protein 65 | MYB65 | 2day perior to sampling | 0.12 |
| AT2G17140.1 | Pentatricopeptide repeat (PPR) superfamily protein | - | 2day perior to sampling | 0.16 |
| AT4G37680.3 | heptahelical protein 4 | HHP4 | 2day perior to sampling | 0.16 |
| AT2G21700.1 | pre-tRNA | - | 2day perior to sampling | 0.25 |
| AT1G64800.1 | DNA binding;sequence-specific DNA binding transcription factors | - | 2day perior to sampling | 0.25 |
| AT2G02550.1 | PIN domain-like family protein | - | 2day perior to sampling | 0.25 |
| AT3G54960.1 | PDI-like 1-3 | PDIL1-3 | 2day perior to sampling | 0.10 |
| AT3G52200.2 | Dihydrolipoamide acetyltransferase, long form protein | LTA3 | 2day perior to sampling | 0.21 |
| AT3G49770.1 | NULL | - | 2day perior to sampling | -0.18 |

| | | | | |
|---|---|---|---|---|
| AT1G05597.1 | | - | 2day perior to sampling | -0.07 |
| AT1G65830.1 | pre-tRNA | - | 2day perior to sampling | -0.12 |
| AT3G55980.2 | salt-inducible zinc finger 1 | SZF1 | 2day perior to sampling | -0.06 |
| AT3G60870.1 | AT-hook motif nuclear-localized protein 18 | AHL18 | 2day perior to sampling | -0.26 |
| AT2G09960.1 | transposable element gene | - | 2day perior to sampling | -0.20 |
| AT1G49370.1 | NULL | - | 2day perior to sampling | -0.05 |
| AT2G04465.1 | | - | 2day perior to sampling | -0.11 |
| AT1G30260.1 | NULL | - | 2day perior to sampling | -0.10 |
| AT1G69550.1 | disease resistance protein (TIR-NBS-LRR class) | - | 2day perior to sampling | -0.24 |
| AT1G56460.2 | HIT zinc finger ;PAPA-1-like conserved region | - | 2day perior to sampling | -0.11 |
| AT1G60050.1 | Nodulin MtN21 /EamA-like transporter family protein | UMAMIT35 | 2day perior to sampling | -0.20 |
| AT3G16550.1 | DEGP protease 12 | DEG12 | 2day perior to sampling | -0.30 |
| AT5G13800.1 | pheophytinase | PPH | 2day perior to sampling | -0.29 |
| AT3G52800.2 | A20/AN1-like zinc finger family protein | - | 2day perior to sampling | -0.24 |
| AT4G06950.1 | | - | 2day perior to sampling | -0.13 |
| AT3G29460.1 | transposable element gene | - | 2day perior to sampling | -0.29 |
| AT2G40165.1 | | - | 2day perior to sampling | -0.23 |
| AT4G22330.1 | Alkaline phytoceramidase (aPHC) | ATCES1 | 2day perior to sampling | -0.20 |
| AT1G64618.1 | other RNA | - | 2day perior to sampling | -0.25 |
| AT4G04435.1 | | - | 2day perior to sampling | -0.09 |
| AT1G61280.2 | Phosphatidylinositol N-acetylglucosaminyltransferase, GPI19/PIG-P | - | 2day perior to sampling | -0.08 |
| AT4G22415.1 | transposable element gene | - | 2day perior to sampling | -0.45 |
| AT3G31460.1 | transposable element gene | - | 2day perior to sampling | -0.15 |
| AT5G08205.1 | | - | 2day perior to sampling | -0.09 |
| AT1G07303.1 | | - | 2day perior to sampling | -0.32 |
| AT1G61000.1 | NULL | - | 2day perior to sampling | -0.34 |
| AT2G14285.1 | Small nuclear ribonucleoprotein family protein | - | 2day perior to sampling | -0.19 |
| AT4G01040.1 | Glycosyl hydrolase superfamily protein | - | 2day perior to sampling | -0.25 |
| AT5G09365.1 | | - | 2day perior to sampling | -0.23 |
| AT2G23100.1 | Cysteine/Histidine-rich C1 domain family protein | - | 2day perior to sampling | -0.20 |
| AT4G14630.1 | germin-like protein 9 | GLP9 | 2day perior to sampling | -0.15 |
| AT5G05100.1 | Single-stranded nucleic acid binding R3H protein | - | 2day perior to sampling | -0.23 |
| AT2G04940.1 | scramblase-related | - | 2day perior to sampling | -0.20 |
| AT5G26236.1 | transposable element gene | - | 2day perior to sampling | -0.10 |
| AT2G32905.1 | Domain of unknown function (DUF313) | - | 2day perior to sampling | -0.16 |
| AT4G29380.1 | protein kinase family protein / WD-40 repeat family protein | VPS15 | 2day perior to sampling | -0.11 |
| AT3G48860.2 | NULL | - | 2day perior to sampling | -0.18 |
| AT1G67630.1 | DNA polymerase alpha 2 | POLA2 | 2day perior to sampling | -0.20 |
| AT3G61590.1 | Galactose oxidase/kelch repeat superfamily protein | HWS | 2day perior to sampling | -0.18 |
| AT4G08028.1 | Molecular chaperone Hsp40/DnaJ family protein | - | 2day perior to sampling | -0.06 |
| AT2G08975.1 | | - | 2day perior to sampling | -0.20 |
| AT1G48760.2 | delta-adaptin | delta-ADR | 2day perior to sampling | -0.10 |
| AT1G25250.2 | indeterminate(ID)-domain 16 | IDD16 | 2day perior to sampling | -0.24 |
| AT2G08190.1 | | - | 2day perior to sampling | -0.20 |
| AT1G32280.1 | Bifunctional inhibitor/lipid-transfer protein/seed storage 2S albumin superfamily protein | - | 2day perior to sampling | -0.21 |
| AT5G56555.1 | | - | 2day perior to sampling | -0.09 |
| AT5G66755.1 | pre-tRNA | - | 2day perior to sampling | -0.07 |
| AT1G45215.1 | Protein of unknown function (DUF784) | - | 2day perior to sampling | -0.23 |
| AT5G04785.1 | | - | 2day perior to sampling | -0.16 |
| AT5G40680.1 | Galactose oxidase/kelch repeat superfamily protein | - | 2day perior to sampling | -0.11 |
| AT5G19290.1 | alpha/beta-Hydrolases superfamily protein | - | 2day perior to sampling | -0.16 |
| AT5G61630.1 | NULL | - | 2day perior to sampling | -0.11 |
| AT1G25500.2 | Plasma-membrane choline transporter family protein | - | 2day perior to sampling | -0.07 |
| AT1G75070.1 | pre-tRNA | - | 2day perior to sampling | -0.25 |
| AT3G21090.1 | ABC-2 type transporter family protein | ABCG15 | 2day perior to sampling | -0.25 |
| AT4G21100.1 | damaged DNA binding protein 1B | DDB1B | 2day perior to sampling | -0.22 |
| AT3G48740.1 | Nodulin MtN3 family protein | SWEET11 | 2day perior to sampling | -0.19 |
| AT1G40097.1 | transposable element gene | - | 2day perior to sampling | -0.18 |
| AT3G33133.1 | transposable element gene | - | 2day perior to sampling | -0.25 |
| AT5G35340.1 | transposable element gene | - | 2day perior to sampling | -0.14 |
| AT2G13940.1 | transposable element gene | - | 2day perior to sampling | -0.32 |
| AT2G38920.4 | SPX (SYG1/Pho81/XPR1) domain-containing protein / zinc finger (C3HC4-type RING finger) protein-related | - | 2day perior to sampling | -0.24 |
| AT5G37352.1 | NULL | - | 2day perior to sampling | -0.20 |
| AT1G04743.1 | | - | 2day perior to sampling | -0.25 |
| AT3G05395.1 | | - | 2day perior to sampling | -0.17 |
| AT1G43883.1 | transposable element gene | - | 2day perior to sampling | -0.09 |
| AT3G13500.1 | NULL | - | 2day perior to sampling | -0.31 |
| AT5G44568.1 | NULL | - | 2day perior to sampling | -0.24 |
| AT1G50420.1 | scarecrow-like 3 | SCL3 | 2day perior to sampling | -0.29 |
| AT1G53470.1 | mechanosensitive channel of small conductance-like 4 | MSL4 | 2day perior to sampling | -0.25 |
| AT3G08545.1 | | - | 2day perior to sampling | -0.19 |
| AT3G60980.1 | Tetratricopeptide repeat (TPR)-like superfamily protein | - | 2day perior to sampling | -0.13 |
| AT3G48920.1 | myb domain protein 45 | MYB45 | 2day perior to sampling | -0.27 |
| AT5G07235.1 | | - | 2day perior to sampling | -0.06 |

| | | | | |
|---|---|---|---|---|
| AT5G04480.1 | UDP-Glycosyltransferase superfamily protein | - | 2day perior to sampling | -0.25 |
| AT1G53900.2 | Eukaryotic translation initiation factor 2B (eIF-2B) family protein | - | 2day perior to sampling | -0.20 |
| AT1G46624.1 | transposable element gene | - | 2day perior to sampling | -0.27 |
| AT5G05360.1 | NULL | - | 2day perior to sampling | -0.22 |
| AT5G13890.1 | Family of unknown function (DUF716) | - | 2day perior to sampling | -0.20 |
| AT5G03553.1 | | - | 2day perior to sampling | -0.27 |
| AT5G27620.2 | cyclin H;1 | CYCH%3B1 | 2day perior to sampling | -0.12 |
| AT3G17720.1 | Pyridoxal phosphate (PLP)-dependent transferases superfamily protein | - | 2day perior to sampling | -0.23 |
| AT1G21630.2 | Calcium-binding EF hand family protein | - | 2day perior to sampling | -0.24 |
| AT3G20400.1 | F-box associated ubiquitination effector family protein | EMB2743 | 2day perior to sampling | -0.16 |
| AT3G28720.2 | NULL | - | 2day perior to sampling | -0.22 |
| AT5G39750.1 | AGAMOUS-like 81 | AGL81 | 2day perior to sampling | -0.36 |
| AT1G23930.1 | transposable element gene | - | 2day perior to sampling | -0.22 |
| AT3G04330.1 | Kunitz family trypsin and protease inhibitor protein | - | 2day perior to sampling | -0.12 |
| AT3G09162.1 | NULL | - | 2day perior to sampling | -0.14 |
| AT3G44326.1 | F-box family protein | - | 2day perior to sampling | -0.16 |
| AT3G44935.1 | NULL | - | 2day perior to sampling | -0.25 |
| AT5G23210.5 | serine carboxypeptidase-like 34 | SCPL34 | 2day perior to sampling | -0.24 |
| AT1G53635.1 | NULL | - | 2day perior to sampling | -0.15 |
| AT1G68460.1 | isopentenyltransferase 1 | IPT1 | 2day perior to sampling | -0.07 |
| AT4G16920.2 | Disease resistance protein (TIR-NBS-LRR class) family | - | 2day perior to sampling | -0.27 |
| AT2G33470.1 | glycolipid transfer protein 1 | GLTP1 | 2day perior to sampling | -0.32 |
| AT4G13600.1 | Carbohydrate-binding X8 domain superfamily protein | - | 2day perior to sampling | -0.21 |
| AT3G46387.1 | transposable element gene | - | 2day perior to sampling | -0.19 |
| AT5G44973.1 | NULL | - | 2day perior to sampling | -0.22 |
| AT3G25013.2 | Synaptobrevin family protein | - | 2day perior to sampling | -0.22 |
| AT3G42305.1 | transposable element gene | - | 2day perior to sampling | -0.12 |
| AT5G26320.1 | TRAF-like family protein | - | 2day perior to sampling | -0.09 |
| AT5G11720.1 | Glycosyl hydrolases family 31 protein | - | 2day perior to sampling | -0.24 |
| AT5G59620.1 | transposable element gene | - | 2day perior to sampling | -0.26 |
| AT3G61130.1 | galacturonosyltransferase 1 | GAUT1 | 2day perior to sampling | -0.13 |
| AT1G60600.2 | UbiA prenyltransferase family protein | ABC4 | 2day perior to sampling | -0.08 |
| AT1G72910.1 | Toll-Interleukin-Resistance (TIR) domain-containing protein | - | 2day perior to sampling | -0.20 |
| AT5G39693.1 | MIR869a; miRNA | MIR869A | 2day perior to sampling | -0.21 |
| AT1G36406.1 | transposable element gene | - | 2day perior to sampling | -0.13 |
| AT3G01605.1 | | - | 2day perior to sampling | -0.18 |
| AT3G14510.1 | Polyprenyl synthetase family protein | - | 2day perior to sampling | -0.27 |
| AT1G18871.1 | NULL | - | 2day perior to sampling | -0.10 |
| AT3G04860.1 | Plant protein of unknown function (DUF868) | - | 2day perior to sampling | -0.22 |
| AT1G54420.1 | NULL | - | 2day perior to sampling | -0.24 |
| AT4G38480.1 | Transducin/WD40 repeat-like superfamily protein | - | 2day perior to sampling | -0.20 |
| AT1G75520.1 | SHI-related sequence 5 | SRS5 | 2day perior to sampling | -0.19 |
| AT5G24830.1 | Tetratricopeptide repeat (TPR)-like superfamily protein | - | 2day perior to sampling | -0.18 |
| AT2G09555.1 | | - | 2day perior to sampling | -0.26 |
| AT3G51150.2 | ATP binding microtubule motor family protein | - | 2day perior to sampling | -0.10 |
| AT2G36750.1 | UDP-glucosyl transferase 73C1 | UGT73C1 | 2day perior to sampling | -0.26 |
| AT5G36662.1 | ECA1 gametogenesis related family protein | - | 2day perior to sampling | -0.19 |
| AT5G48860.1 | NULL | - | 2day perior to sampling | -0.23 |
| AT5G65535.1 | pre-tRNA | - | 2day perior to sampling | -0.09 |
| AT5G38880.1 | NULL | - | 2day perior to sampling | -0.22 |
| AT5G03095.1 | | - | 2day perior to sampling | -0.33 |
| AT3G05105.1 | | - | 2day perior to sampling | -0.10 |
| AT5G28580.2 | transposable element gene | - | 2day perior to sampling | -0.35 |
| AT1G77010.1 | Pentatricopeptide repeat (PPR) superfamily protein | - | 2day perior to sampling | -0.13 |
| AT5G65690.4 | phosphoenolpyruvate carboxykinase 2 | PCK2 | 2day perior to sampling | -0.24 |
| AT3G30350.2 | NULL | RGF4 | 2day perior to sampling | -0.09 |
| AT5G16650.2 | Chaperone DnaJ-domain superfamily protein | - | 2day perior to sampling | -0.11 |
| AT3G61710.4 | AUTOPHAGY 6 | ATG6 | 2day perior to sampling | -0.28 |
| AT2G09990.1 | Ribosomal protein S5 domain 2-like superfamily protein | - | 2day perior to sampling | -0.09 |
| AT2G32550.2 | Cell differentiation, Rcd1-like protein | - | 2day perior to sampling | -0.08 |
| AT4G04560.1 | transposable element gene | - | 2day perior to sampling | -0.23 |
| AT2G07220.1 | transposable element gene | - | 2day perior to sampling | -0.27 |
| AT3G55840.1 | Hs1pro-1 protein | - | 2day perior to sampling | -0.25 |
| AT4G28220.1 | NAD(P)H dehydrogenase B1 | NDB1 | 2day perior to sampling | -0.09 |
| AT3G24575.1 | | - | 2day perior to sampling | -0.24 |
| AT5G08540.1 | NULL | - | 2day perior to sampling | -0.27 |
| AT4G16020.2 | transposable element gene | - | 2day perior to sampling | -0.11 |
| AT3G08920.1 | Rhodanese/Cell cycle control phosphatase superfamily protein | - | 2day perior to sampling | -0.30 |
| AT3G46980.4 | phosphate transporter 4;3 | PHT4%3B3 | 2day perior to sampling | -0.08 |
| AT5G17050.1 | UDP-glucosyl transferase 78D2 | UGT78D2 | 2day perior to sampling | -0.30 |
| AT5G37072.3 | NULL | - | 2day perior to sampling | -0.19 |
| AT4G08040.1 | 1-aminocyclopropane-1-carboxylate synthase 11 | ACS11 | 2day perior to sampling | -0.11 |
| AT5G08710.2 | Regulator of chromosome condensation (RCC1) family protein | RUG1 | 2day perior to sampling | -0.14 |
| AT3G06650.2 | ATP-citrate lyase B-1 | ACLB-1 | 2day perior to sampling | -0.12 |
| ATCG00710.1 | photosystem II reaction center protein H | PSBH | 2day perior to sampling | -0.07 |
| AT1G62610.4 | NAD(P)-binding Rossmann-fold superfamily protein | - | 2day perior to sampling | -0.14 |

| | | | | |
|---|---|---|---|---|
| AT1G44020.1 | Cysteine/Histidine-rich C1 domain family protein | - | 2day perior to sampling | -0.24 |
| AT5G19151.1 | NULL | - | 2day perior to sampling | -0.12 |
| AT5G00390.1 | | - | 2day perior to sampling | -0.37 |
| AT2G43680.5 | IQ-domain 14 | IQD14 | 2day perior to sampling | -0.28 |
| AT1G06970.1 | cation/hydrogen exchanger 14 | CHX14 | 2day perior to sampling | -0.27 |
| AT1G01725.1 | NULL | - | 2day perior to sampling | -0.22 |
| AT5G60335.1 | Thioesterase superfamily protein | - | 2day perior to sampling | -0.07 |
| AT5G39520.1 | Protein of unknown function (DUF1997) | - | 2day perior to sampling | -0.24 |
| AT1G54230.1 | Winged helix-turn-helix transcription repressor DNA-binding | - | 2day perior to sampling | -0.31 |
| AT4G23960.2 | F-box family protein | - | 2day perior to sampling | -0.25 |
| AT3G06990.1 | Cysteine/Histidine-rich C1 domain family protein | - | 2day perior to sampling | -0.38 |
| AT5G38210.1 | Protein kinase family protein | - | 2day perior to sampling | -0.11 |
| AT5G47630.2 | mitochondrial acyl carrier protein 3 | mtACP3 | 2day perior to sampling | -0.25 |
| AT5G22796.1 | | - | 2day perior to sampling | -0.21 |
| AT2G30130.1 | Lateral organ boundaries (LOB) domain family protein | ASL5 | 2day perior to sampling | -0.21 |
| AT4G27310.1 | B-box type zinc finger family protein | BBX28 | 2day perior to sampling | -0.37 |
| AT5G16640.1 | Pentatricopeptide repeat (PPR) superfamily protein | - | 2day perior to sampling | -0.34 |
| AT3G33025.1 | NULL | - | 2day perior to sampling | -0.19 |
| AT5G56720.1 | Lactate/malate dehydrogenase family protein | c-NAD-MDH3 | 2day perior to sampling | -0.17 |
| AT5G06255.1 | | - | 2day perior to sampling | -0.21 |
| AT5G35270.1 | transposable element gene | - | 2day perior to sampling | -0.10 |
| AT3G32060.1 | transposable element gene | - | 2day perior to sampling | -0.13 |
| AT5G45670.1 | GDSL-like Lipase/Acylhydrolase superfamily protein | - | 2day perior to sampling | -0.25 |
| AT1G27045.2 | Homeobox-leucine zipper protein family | ATHB54 | 2day perior to sampling | -0.22 |
| AT2G25340.1 | vesicle-associated membrane protein 712 | VAMP712 | 2day perior to sampling | -0.10 |
| AT5G36210.1 | alpha/beta-Hydrolases superfamily protein | - | 2day perior to sampling | -0.18 |
| AT3G09735.1 | S1FA-like DNA-binding protein | - | 2day perior to sampling | -0.19 |
| AT5G17640.1 | Protein of unknown function (DUF1005) | ASG1 | 2day perior to sampling | -0.21 |
| AT5G32950.1 | transposable element gene | - | 2day perior to sampling | -0.10 |
| AT1G09020.1 | homolog of yeast sucrose nonfermenting 4 | SNF4 | 2day perior to sampling | -0.24 |
| AT5G48640.4 | Cyclin family protein | - | 2day perior to sampling | -0.31 |
| AT3G32912.1 | transposable element gene | - | 2day perior to sampling | -0.14 |
| AT1G31993.1 | transposable element gene | - | 2day perior to sampling | -0.20 |
| AT5G56440.1 | F-box/RNI-like/FBD-like domains-containing protein | - | 2day perior to sampling | -0.23 |
| AT1G52150.3 | Homeobox-leucine zipper family protein / lipid-binding START domain-containing protein | ATHB-15 | 2day perior to sampling | -0.25 |
| AT1G65680.1 | expansin B2 | EXPB2 | 2day perior to sampling | -0.17 |
| AT2G12160.1 | NULL | - | 2day perior to sampling | -0.19 |
| AT1G66245.1 | NULL | - | 2day perior to sampling | -0.28 |
| AT3G24550.1 | proline extensin-like receptor kinase 1 | PERK1 | 2day perior to sampling | -0.14 |
| AT5G38830.1 | Cysteinyl-tRNA synthetase, class Ia family protein | - | 2day perior to sampling | -0.27 |
| AT1G77690.1 | like AUX1 3 | LAX3 | 2day perior to sampling | -0.14 |
| AT1G06037.1 | | - | 2day perior to sampling | -0.11 |
| AT5G64040.1 | photosystem I reaction center subunit PSI-N, chloroplast, putative / PSI-N, putative (PSAN) | PSAN | 2day perior to sampling | -0.26 |
| AT1G16445.1 | S-adenosyl-L-methionine-dependent methyltransferases superfamily | - | 2day perior to sampling | -0.40 |
| ATMG00080.1 | ribosomal protein L16 | RPL16 | 2day perior to sampling | -0.09 |
| AT2G36470.1 | Plant protein of unknown function (DUF868) | - | 2day perior to sampling | -0.27 |
| AT2G06925.1 | Phospholipase A2 family protein | PLA2-ALPHA | 2day perior to sampling | -0.26 |
| AT5G35965.1 | transposable element gene | - | 2day perior to sampling | -0.16 |
| AT4G25860.2 | OSBP(oxysterol binding protein)-related protein 4A | ORP4A | 2day perior to sampling | -0.25 |
| AT1G77730.1 | Pleckstrin homology (PH) domain superfamily protein | - | 2day perior to sampling | -0.23 |
| AT5G08605.1 | | - | 2day perior to sampling | -0.13 |
| AT4G06630.1 | transposable element gene | - | 2day perior to sampling | -0.17 |
| AT5G30470.1 | transposable element gene | - | 2day perior to sampling | -0.22 |
| AT5G53770.1 | Nucleotidyltransferase family protein | - | 2day perior to sampling | -0.05 |
| AT4G12680.1 | NULL | - | 2day perior to sampling | -0.09 |
| AT2G25125.1 | | - | 2day perior to sampling | -0.09 |
| AT3G18830.1 | polyol/monosaccharide transporter 5 | PMT5 | 2day perior to sampling | -0.05 |
| AT1G66490.1 | F-box and associated interaction domains-containing protein | - | 3day perior to sampling | 0.24 |
| AT5G66570.1 | PS II oxygen-evolving complex 1 | PSBO1 | 3day perior to sampling | 0.25 |
| AT2G17140.1 | Pentatricopeptide repeat (PPR) superfamily protein | - | 3day perior to sampling | 0.17 |
| AT4G14630.1 | germin-like protein 9 | GLP9 | 3day perior to sampling | -0.19 |
| AT5G13890.1 | Family of unknown function (DUF716) | - | 3day perior to sampling | -0.22 |
| AT3G20400.1 | F-box associated ubiquitination effector family protein | EMB2743 | 3day perior to sampling | -0.16 |
| AT1G19485.2 | Transducin/WD40 repeat-like superfamily protein | - | 3day perior to sampling | -0.30 |
| AT2G34860.1 | DnaJ/Hsp40 cysteine-rich domain superfamily protein | EDA3 | 3day perior to sampling | -0.09 |

**Table S3 Summary of the false-assignment rates reported by previous studies**

| Experimental procedure | Rate (%) | Literature |
|---|---|---|
| false-assignment rate estimated from sequencing error | <0.001 | Kircher et al., 2012 (1) |
| false-assignment rate due to pooled PCR | 0.4 | Kircher et al., 2012 (1) |
| Mistake in cluster assign during sequencing | 0.01 - 0.03 | Kircher et al., 2012 (1) |
| Index hopping in exclusion PCR | 0.1 - 2 | Illumina (2) |

(1) Kircher, M., Sawyer, S. and Meyer, M. (2012) Double indexing overcomes inaccuracies in multiplex sequencing on the Illumina platform. Nucleic Acids Res 40: e3.

(2) https://jp.illumina.com/science/education/minimizing-index-hopping.html
